# Discovery and Synthesis of GS-7682, a Novel Prodrug of a 4′-CN-4-Aza-7,9-Dideazaadenosine *C*-Nucleoside with Broad-Spectrum Potency Against Pneumo- and Picornaviruses and Efficacy in RSV-Infected African Green Monkeys

**DOI:** 10.1101/2024.04.17.589937

**Authors:** Dustin S. Siegel, Hon C. Hui, Jared Pitts, Meghan S. Vermillion, Kazuya Ishida, Davin Rautiola, Michael Keeney, Hammad Irshad, Lijun Zhang, Kwon Chun, Gregory Chin, Bindu Goyal, Edward Doerffler, Hai Yang, Michael O. Clarke, Chris Palmiotti, Arya Vijjapurapu, Nicholas C. Riola, Kirsten Stray, Eisuke Murakami, Bin Ma, Ting Wang, Xiaofeng Zhao, Yili Xu, Gary Lee, Bruno Marchand, Minji Seung, Arabinda Nayak, Adrian Tomkinson, Nani Kadrichu, Scott Ellis, Ona Barauskas, Joy Y. Feng, Jason K. Perry, Michel Perron, John P. Bilello, Philip J. Kuehl, Raju Subramanian, Tomas Cihlar, Richard L. Mackman

## Abstract

Acute respiratory viral infections (ARVI), such as pneumovirus and respiratory picornavirus infections, exacerbate disease in COPD and asthma patients. A research program targeting respiratory syncytial virus (RSV) led to the discovery of GS-7682 (**1**) a novel phosphoramidate prodrug of a 4′-CN-4-aza-7,9-dideazaadenosine *C*-nucleoside GS-646089 (**2**) with broad antiviral activity against RSV EC_50_ = 3-46 nM, human metapneumovirus (hMPV) EC_50_ = 210 ± 50 nM, human rhinovirus (RV) EC_50_ = 54-61 nM, and enterovirus (EV) EC_50_ = 83-90 nM. Prodrug optimization for cellular potency and lung cell metabolism identified the 5’-methyl((*S*)-hydroxy(phenoxy)phosphoryl)-L-alaninate in combination with 2’,3’-diisobutyrate promoieties as optimal for high intracellular triphosphate formation in vitro and in vivo. **1** demonstrated significant reductions of viral loads in the lower respiratory tract of RSV-infected African green monkeys when administered once daily via intratracheal nebulized aerosol. Together these finding support additional evaluation of **1** and its analogs as a potential therapeutic for pneumo- and picornaviruses.

## Introduction

Acute respiratory viral infections (ARVI) represent a significant cause of morbidity and mortality worldwide. In the wake of the COVID-19 global health emergency, increased awareness of the global disease burden, heightened community-based surveillance, and the expanded use of molecular multiplex diagnostic assays for respiratory viruses have greatly increased our ability to rapidly diagnose ARVIs, which is essential for effective antiviral intervention.^1^ Recent analysis shows the pneumoviruses (RSV and hMPV), and the respiratory picornaviruses (RV and EVs) have high prevalence.^2^ In pediatrics, the elderly (especially with comorbidities), and those who are immunocompromised, these infections can progress to the lower respiratory tract (LRT) resulting in severe disease associated with airway inflammation, bronchiolitis or bronchitis, pneumonia, and respiratory failure in extreme cases. In particular, adults with underlying chronic respiratory disease, namely chronic obstructive pulmonary disease (COPD) and asthma, are disproportionally affected by these ARVIs. A key clinical feature of COPD and asthma are the occurrences of exacerbations characterized as acute events of increased symptomology, airflow obstruction, and progressive decline in lung function. ARVIs, in particular RV and RSV,^3–6^ are identified as a major cause of these exacerbations which contribute significantly to morbidity, mortality, and reduction in overall quality of life.^3,6,7^ Among adults, the viruses in these families account for up to 40% of non-influenza ARVI illnesses requiring hospitalization^8,9^ with a substantial burden to the economy estimated at over $40 billion in the US alone.^10^ Advances in prophylaxis for RSV have been recently realized with the approvals of vaccines Arexvy^®^ and Abrysvo^®^ for adults.^11,12^ However, there are currently no approved treatments that are both safe and effective for RSV, hMPV, RV, or EV.

The human pneumoviruses, RSV and hMPV, are enveloped single-stranded negative-sense ribonucleic acid (RNA) viruses that each have two subtypes A and B. The human respiratory picornaviruses, RV and EV, are non-enveloped single-stranded positive-sense RNA viruses with RV distinguished by three species A, B, and C. The viral RNA-dependent RNA polymerases (RdRp) encoded by these viruses represent salient molecular targets for ARVI treatment as they are essential for the replication and transcription of the viral genome. Ribonucleoside(tide) analogues that mimic natural RNA substrates and inhibit the respiratory viral polymerase have been reported (Figure 1).^13^ Ribavirin (**3**), a broad-spectrum inhaled antiviral nucleoside, has been approved for the treatment of hospitalized infants with severe RSV infection, but has limited efficacy and significant concerns for safety.^14–16^ Lumicitabine (**4**), an oral RSV antiviral nucleoside, was advanced into Phase 2 clinical trials in infants hospitalized with RSV; however, it was discontinued due to safety concerns associated with neutropenia.^17,18^ Molnupiravir (**5**), a broad-spectrum oral nucleoside, has received emergency use authorization by the FDA for treatment of mild to moderate COVID-19 in adults at risk for progression to severe disease where alternative treatment options are not accessible or appropriate. **5** is also currently being evaluated in a Phase 2 clinical trial for RSV.^19–21^ Remdesivir (RDV, **6**), an IV administered broad-spectrum nucleotide prodrug^22^ of GS-441524 (**7**), was initially discovered as an inhibitor of RSV,^23^ then advanced into clinical development for treatment of Ebola,^24–26^ and ultimately was the first approved treatment of COVID-19 in adults and pediatric patients at high risk for severe disease.^27–30^ Concurrent with the studies reported herein, a nebulized inhaled formulation for **6** was also developed and advanced into Phase 2 clinical trials for treatment of COVID-19.^31–32^ Obeldesivir (ODV, **8**), an oral prodrug of **7**, was explored in Phase 3 clinical trials for COVID-19,^33^ and the parent **7** is reported to have potent RSV activity.^23^

**Figure 1.**
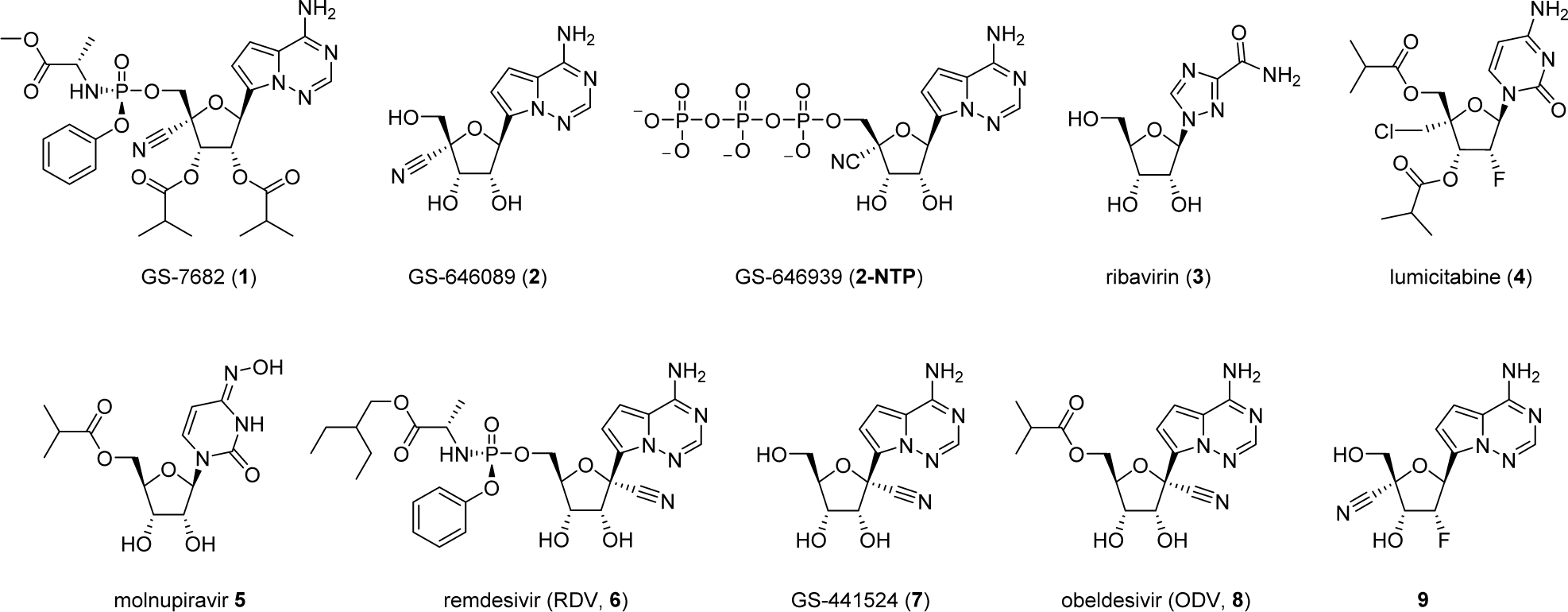
Structures of nucleoside and nucleotide inhibitors of respiratory viruses.

Following the initial discovery of **6**,^23^ a further focus on RdRp inhibitors resulted in the discovery of **9**,^34^ a 4’CN-2’d2’F 4-aza-7,9-dideazaadenosine *C*-nucleoside that inhibits RSV. However, 2’d2’F nucleosides including **9** can mimic both ribonucleoside as well as 2’-deoxyribonucleosides and have been reported to be incorporated by both human DNA and RNA polymerases.^35,36^ Herein, is reported the continued structure-activity relationship (SAR) efforts that combined 4’-substitutions with the *C*-nucleoside 2’-OH ribose core targeting high selectivity against incorporation by human polymerases, which led to the discovery of **1** (Figure 1). **1** is a novel phosphoramidate prodrug of 4’-cyano *C*-ribonucleoside analog **2** that delivers high levels of the active triphosphate GS-646939 (**2-NTP**) in cells. Most notable is that **1** exhibits a broad-spectrum profile of potent activity against both pneumoviruses (RSV, hMPV) and respiratory picornaviruses (RV, EV) with excellent selectivity. Oral delivery of phosphoramidate nucleoside prodrugs is invariably subject to high first-pass hepatic extraction, thereby limiting lung exposure of the intact prodrug. Furthermore, we achieved efficient delivery of **1** by way of pulmonary administration directly to the primary sites of infection in an African green monkey (AGM) model of RSV resulting in strong in vivo efficacy. As such, inhaled **1** is a novel and promising agent for the treatment of multiple respiratory viral infections.^37^

## Results and Discussion

The syntheses of 4’-substituted *C*-ribonucleoside analogs are shown in Scheme 1 starting from the previously described ribonucleoside **10**.^24^ The key step in the synthetic strategy was the introduction of the additional 4’-carbon modification through an oxidation/aldol sequence. In the first step, reduction of the 1’-hydroxyl with BF_3_•OEt_2_ and triethylsilane afforded **11** with high yield and selectivity for the desired β-anomer. Global removal of the benzyl groups yielded **12**.

Acetonide protection of the 2′,3′-hydroxyls followed by selective silylation of the 5’-hydroxyl, 6*N* BOC protection, and desilylation afforded the protected *C*-nucleoside **13**. Oxidation of the 5′-hydroxyl, aldol reaction, and reduction afforded the desired diol **14** in good yield. Through a known sequence for *N*-nucleosides,^38^ a TBS group was selectively installed in high yield on the 5’-hydroxyl generating **15**, a providing the key intermediate for 4’-analogue synthesis via the single unprotected 4’-CH_2_OH handle.

The 4’-methyl analog **20** was prepared through conversion of the 4’-CH_2_OH to the iodide, reduction, and then removal of the protecting groups. The 4’-vinyl **21** was prepared through oxidation of the 4’-CH_2_OH to the aldehyde followed by Wittig olefination and removal of the protecting groups. The 4’-cyano **2** was prepared through oxidation of the 4’-CH_2_OH to the aldehyde followed by oxime formation, CDI mediated nitrile formation and removal of the protecting groups. The 4’-CH_2_Cl **22** was prepared through 4’-CH_2_OH triflation and chloride displacement followed by removal of the protecting groups. An alternative synthetic strategy was required for the installation of the azide group in **26**. The synthesis of **26** began with **11** where 6*N*-benzoylation and removal of the three benzyl groups through hydrogenations afforded **23**. Selective tritylation of the 5’-hydroxyl followed by TBS protection of the 2’,3’-hydroxyls and trityl deprotection yielded **24**. The 5’-hydroxyl was then converted to the iodide with (PhO)_3_MeI and elimination with *t-*BuOK afforded the exocyclic olefin **25**. A sequence of DMDO olefin epoxidation, selective addition of the azide to afford the desired 4’-β-anomer, and removal of the protecting groups resulted in **26**. The nucleotide triphosphates (NTP) were prepared from the parent nucleosides via the general procedure shown in Scheme 2. Treatment of the 4’-substituted ribo-*C*-nucleoside analogues with POCl_3_ followed by pyrophosphate afforded the corresponding NTPs.

**Scheme 1.**
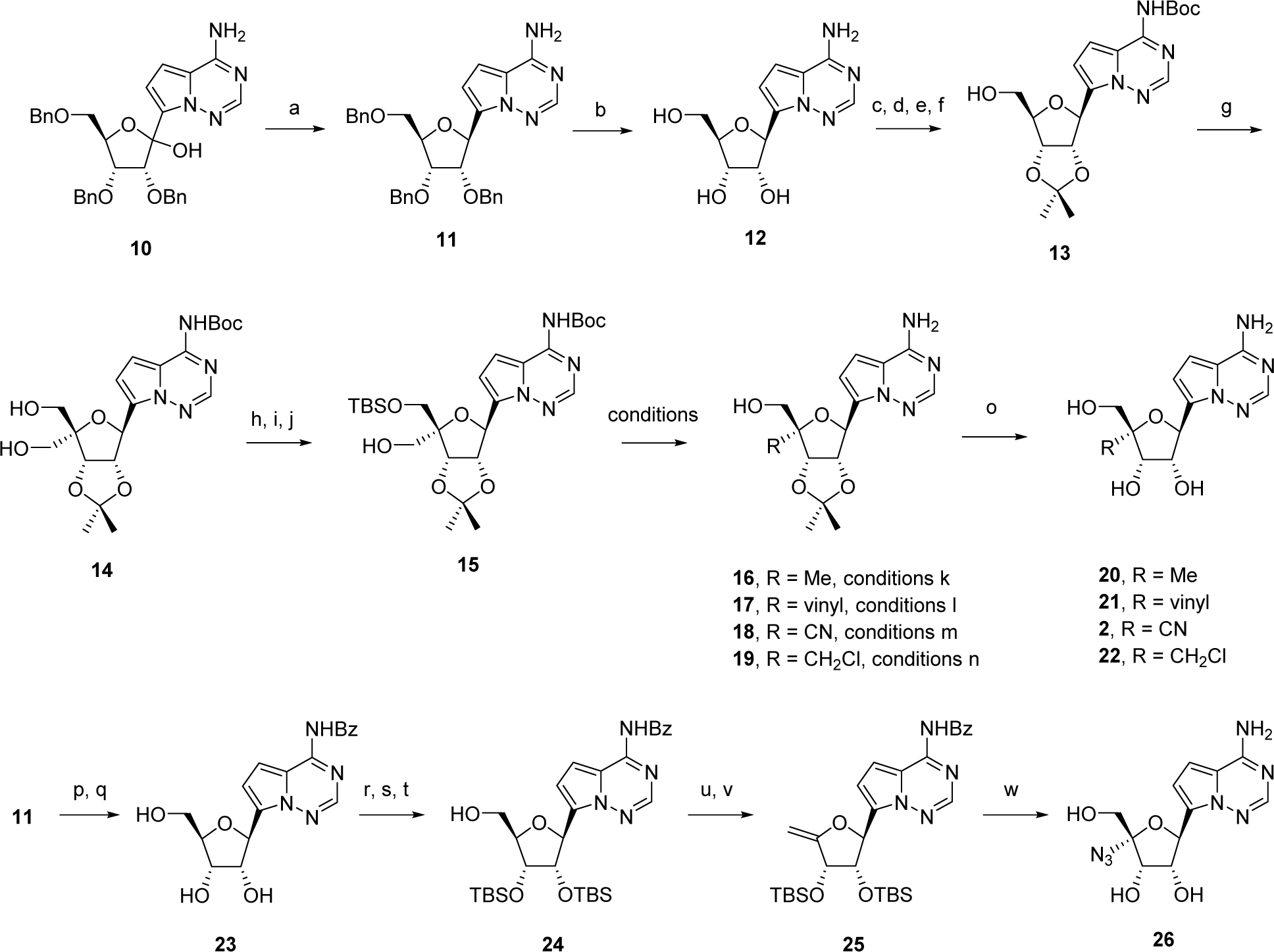
Synthesis of 4’-substituted ribo-*C*-nucleosides.*^a^* *^a^*Reagents and conditions: (a) BF_3_•OEt_2_, Et_3_SiH, CH_2_Cl_2_, 86%; (b) H_2_, Pd/C, AcOH, 99%; (c) (EtO)_3_CH, TsOH, Acetone, 71%; (d) TBSCl, imid., CH_2_Cl_2_, 74%; (e) BOC_2_O, Et_3_N, DMAP, THF, 88%; (f) TBAF, THF, 86%; (g) EDCI, py, TFA, DMSO; H_2_CO, NaOH, dioxane; NaBH_4_, EtOH, 68%; (h) DMTrCl, Et_3_N, CH_2_Cl_2_, 79%; (i) TBSCl, imid., DMF, 78%; (j) TsOH, CHCl_3_, MeOH, 94%; (k) I_2_, PPh_3_, imid., PhMe; H_2_, Pd/C, Et_3_N, MeOH; dioxane, H_2_O, 120 °C, 7% (3-steps); TBAF, THF, 76%; (l) SO_3_•py, DMSO, Et_2_N, CH_2_Cl_2_, 88%; MePPh_3_Br, nBuLi, THF, 78%; TBAF, THF, 92%; dioxane, H_2_O, 120 °C, 98%; (m) EDCI, py, TFA, DMSO, PhMe; NH_2_OH•HCl, py; CDI, MeCN, 87% (3-steps); ZnBr_2_, CH_2_Cl_2_, 99%; TBAF, THF, 93%; (n) TfCl, py; LiCl, DMF, 42% (2-steps); dioxane, H_2_O, 120 °C; TBAF, THF, 87% (2-steps); (o) HCl, MeCN, **20** 82%, **21** 97%, **2** 99%, or TFA, MeCN, H_2_O, **22** 86%. (p) BzCl, py, 99%; (q) HCO_2_H, Pd/C, EtOH, 95%; (r) DMTrCl, py 92%; (s) TBSCl, imid., DMF, 98%; (t) TsOH, MeOH, CH_2_Cl_2_, 92%; (u) (PhO)_3_MeI, DMF, 72%; (v) *t-*BuOK, py, 98%; (w) DMDO, acetone; TMSN_3_, InBr_3_, CH_2_Cl_2_; CsF, DMF; NH_4_OH, MeOH, 29% (4-steps).

**Scheme 2.**
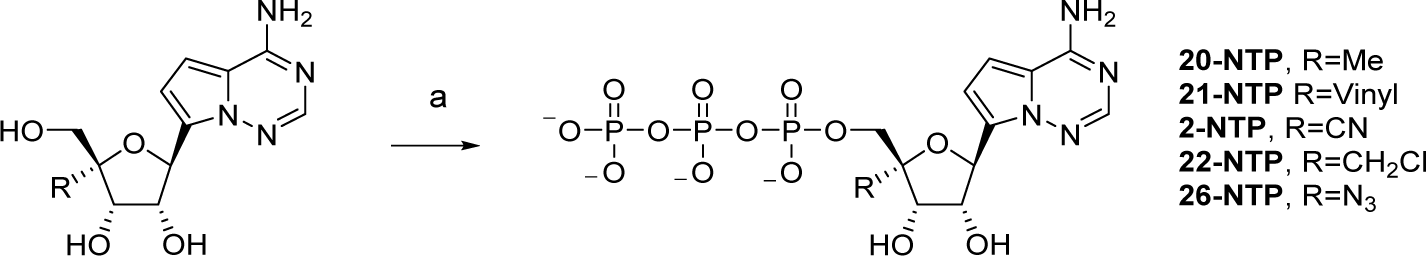
General NTP synthesis of 4’-substituted ribo-*C*-nucleosides.*^a^* *^a^*Reagents and conditions: (a) POCl_3_, [*n*Bu_3_NH]_2_P_2_O_7_H_2_

A variety of phosphoramidate prodrugs **28** were prepared in good yield by coupling of the phosphoramidate reagent **27** in the presence of MgCl_2_ to **18** (Scheme 3). The phosphoramidate reagents **27** contained either pentafluorophenol or *p*-nitrophenol leaving groups and were either coupled as the single *S*p isomer at phosphorus to afford the corresponding *S*p isomers **28** or as a diastereomeric mixture (mix) at phosphorus followed by preparatory chromatography separation to afford the individual *S*p or *R*p isomers of **28**. The corresponding 2’,3’-ester *S*p-phosphoramidates **29** were prepared in good yield through *N*,*N*′-diisopropylcarbodiimide (DIC) mediate carboxylate coupling, or anhydride coupling, and preparatory chromatographic separation if **28-mix** was employed. The structure of **1** was confirmed by small-molecule X-ray crystallography shown in Figure 2.

**Figure 2.**
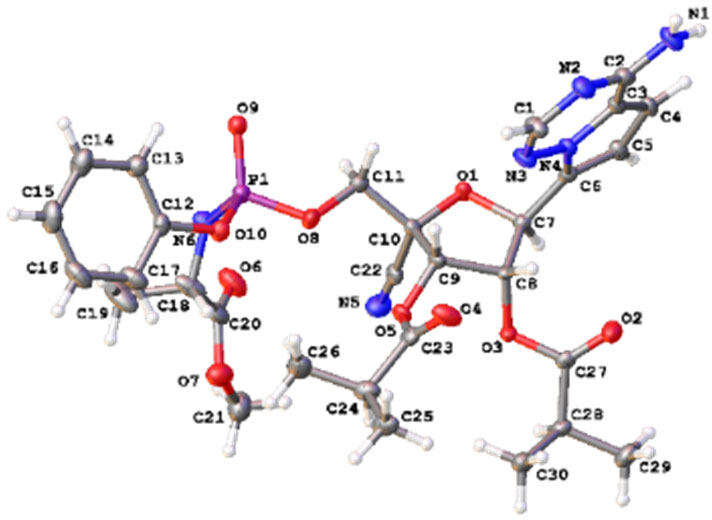
Small molecule X-ray structure thermal ellipsoid representation of **1**.

**Scheme 3.**
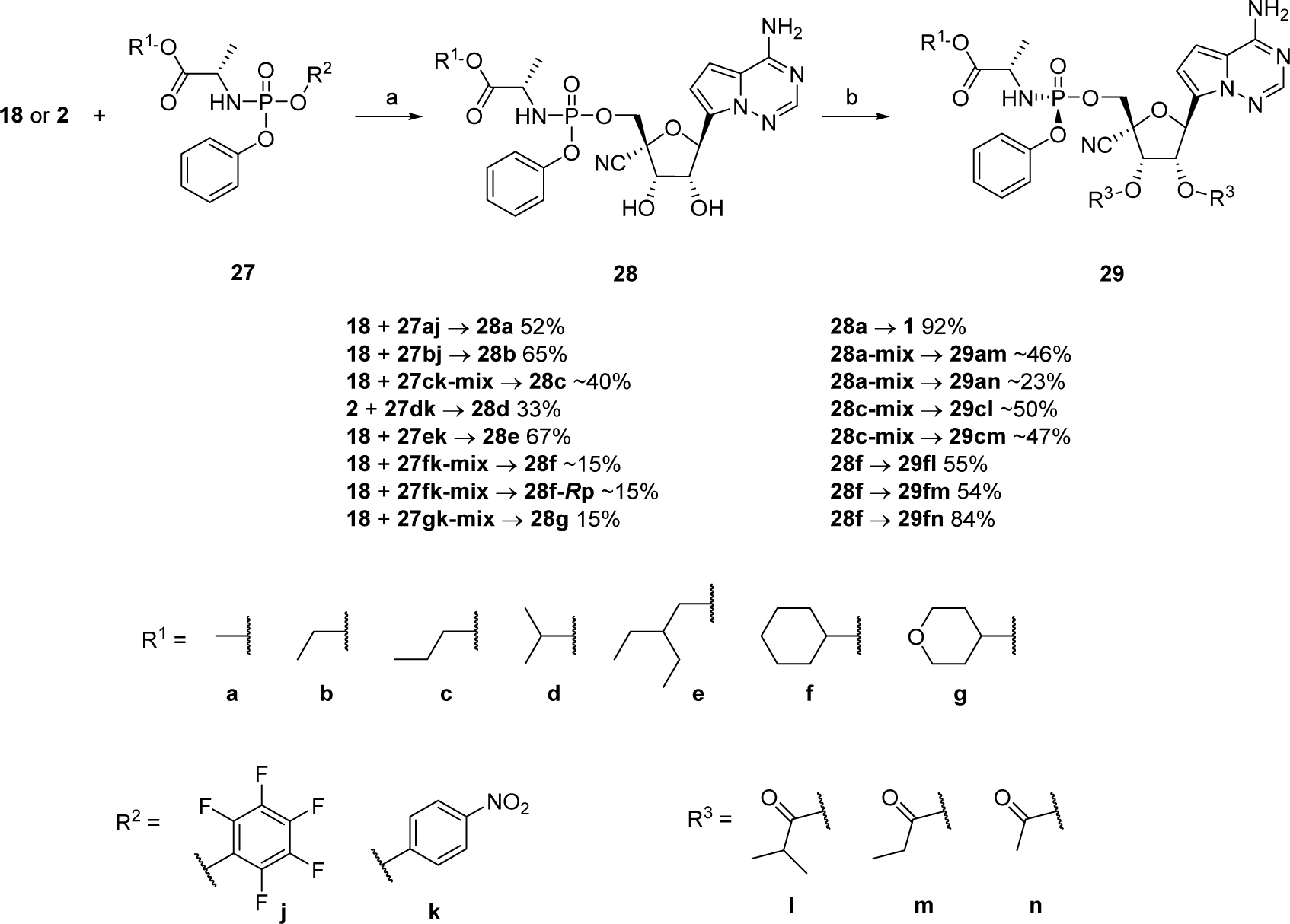
Phosphoramidate prodrug syntheses on nucleoside 2.*^a,b^* *^a^*Reagents and conditions: (a) MgCl_2_, *i*Pr_2_NEt, THF; if **18** then HCl; chromatographic isomer separation from **27**-**mix**. (b) R^3^OH, DIC, DMAP (**29am**, **29an**, **29fl**, **29fm**, **29fn**) or (R^3^)_2_O, DMAP (**1**, **29cl**, **29cm**); chromatographic isomer separation from **28**-**mix**. *^b^*Compounds **27**-**29** are single *S*p isomers at phosphorus unless denoted as single ***R*p** isomer or isomer mixture (**mix**).

The novel 4’-substituted *C-*ribonucleosides **2**, **20-22** and **26** were assayed for RSV inhibition in primary normal human bronchial epithelial (NHBE) cells and in the HEp-2 cell line (Table 1). These two cell types provided a broad assessment of metabolism in physiologically relevant primary lung cells and homogenous immortalized cells. Cytotoxicity was evaluated in HEp-2 cells and the more sensitive, rapidly proliferating MT4 T-cell line. The 4’-substituted analogues overall displayed similar antiviral potency in the NHBE and HEp-2 cells. The unsubstituted *C*-ribonucleoside analogue **12** was highly cytotoxic in HEp-2 and MT4 cells at low nanomolar concentrations. The introduction of ribose modifications rendered selectivity and maintained cell viability at high micromolar concentrations. Notably, the 4’-N_3_ **26** and 4’-CNs **9** and **2** conveyed appreciable antiviral potency against RSV in the 2-7 µM range versus the other 4’-carbon linked substitutions and were in a comparable range to the 1’-CN analog **7**. Furthermore, **26**, **9**, and **2** were not cytotoxic in HEp-2 or MT4 cells at concentrations up to 50 µM.

**Table 1.**
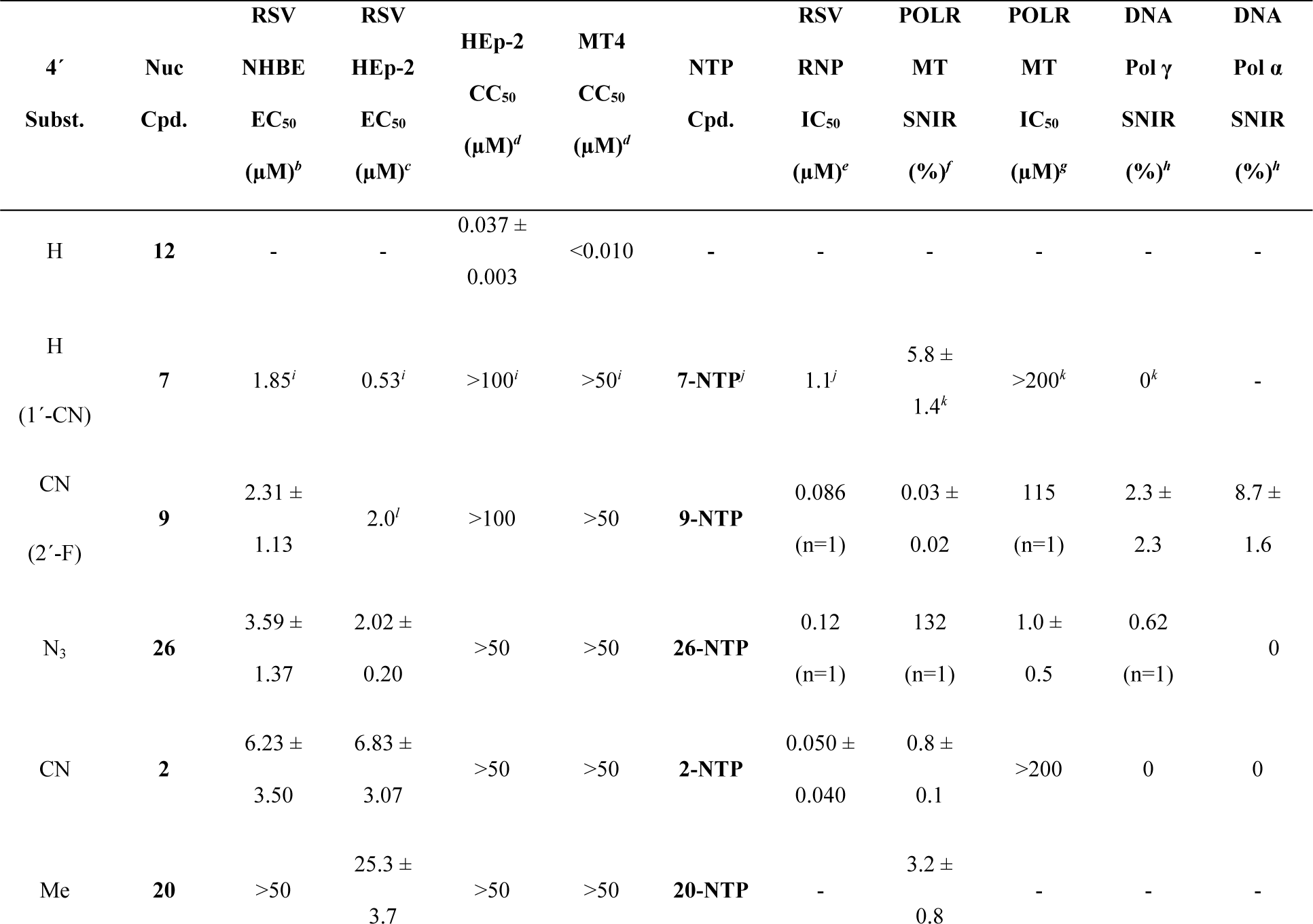
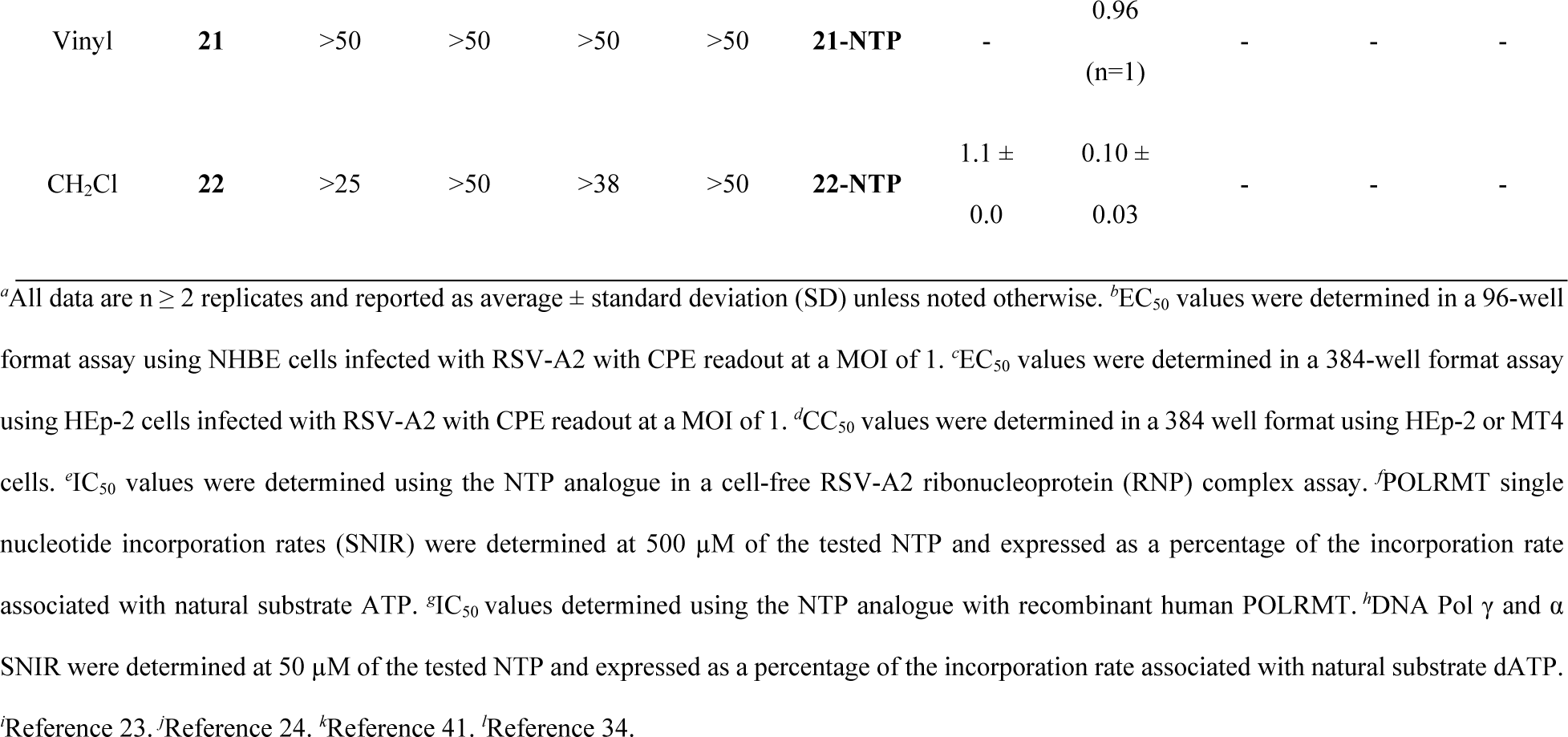
4’-Substituted adenine *C*-nucleoside(tide) SAR.*^a^*.

Nucleoside analogs require activation by intracellular nucleoside(tide) kinases to generate their respective active NTP metabolites, that in turn compete with endogenous nucleotides for incorporation into nascent viral RNA strands (e.g., **2→2-NTP**, Figure 3). The first phosphorylation step, affording the nucleoside 5’-monophosphate (NMP) **2-NMP** in the intracellular anabolism pathway to the NTP, can often be rate-limiting. This can lead to inactive nucleosides in cell culture even though the corresponding NTP may be a highly potent inhibitor of the target RdRp. Therefore, the active NTP analogues were profiled in biochemical assays for their intrinsic potency against the viral polymerases and selectivity against human polymerases to identify the optimal RdRp inhibitor from the series. The RSV activity was evaluated with the RSV-A2 RdRp-containing ribonucleoprotein (RNP) complex^24^ and the human RNA and DNA polymerases were evaluated using single nucleotide incorporation rate assays (SNIR) and inhibition (IC_50_) assays.^39,40^

**Figure 3.**
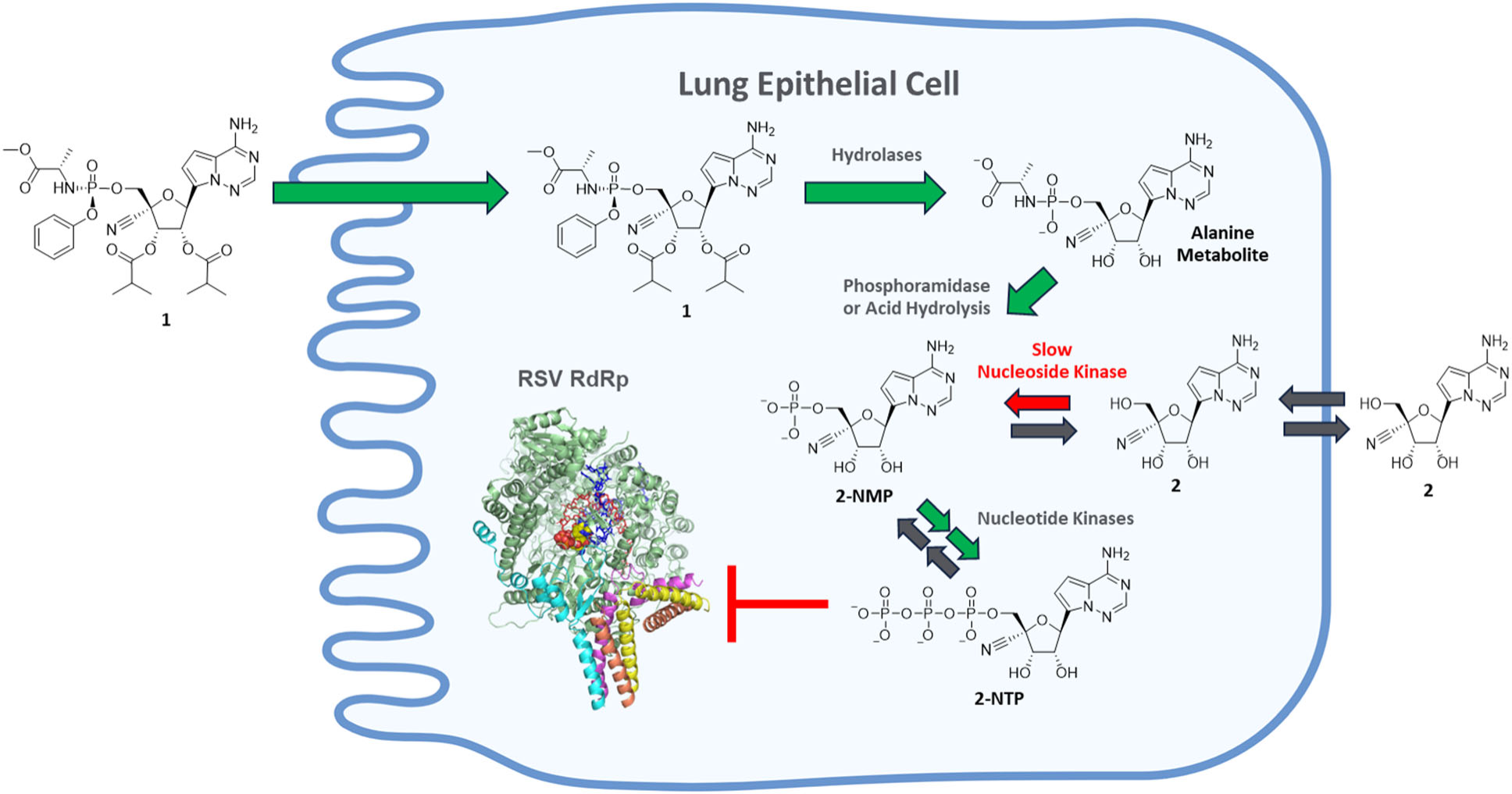
Proposed intracellular metabolism of **1** and **2** in lung epithelial cells.

Notably, the 4’-N_3_ **26-NTP** and the 4’-CN modified analogues, **9-NTP** and **2-NTP**, displayed highly potent inhibition of the RSV RNP and were >10 fold improved as compared to the 1’-CN **7-NTP** and 4’CH_2_Cl **22-NTP** (Table 1). The 4’-N_3_ NTP **26-NTP** showed high incorporation and inhibition of human mitochondrial RNA polymerase (POLRMT) indicating a significant safety risk consistent with a historical 4’-N_3_ nucleoside balapiravir, which was discontinued due to safety findings.^42^ Optimally, the 4’-CN **2-NTP** exhibited the most potent intrinsic inhibition of both the RSV RNP (IC_50_ = 0.050 ± 0.04 µM) and RV RdRp (IC_50_ = 0.043 µM (n=1), data not shown in Table 1) and the highest selectivity against POLRMT (SNIR = 0.8%) and human DNA polymerases (SNIR = 0%). Additionally, the IC_50_ values for **2-NTP** against DNA Pols α and γ were >200 µM (data not shown in Table 1).

Early homology models suggested the possibility of a 4’-pocket.^24^ This hypothesis was later confirmed with the publication of a cryo-electron microscopy structure for apo RSV L-protein (PDB 6PZK).^43^ Using this structure in combination with a ternary structure of Lassa L-protein (PDB 7OJN),^44^ we constructed a model of **2-NTP** as it is positioned in the active site of the RSV polymerase prior to incorporation into the nascent RNA strand (Figure 4). The 4’-CN of **2-NTP** occupies a pocket formed from the palm residues F704, N705, Q782, W785, T786 and G810. Further structural analysis will be included in a broader mechanism of action study published elsewhere.^45^ The model, together with the **2-NTP** data reported in Table 1, supports the proposed mechanism of selective polymerase binding followed by the incorporation of **2-NTP** into viral RNA.

**Figure 4.**
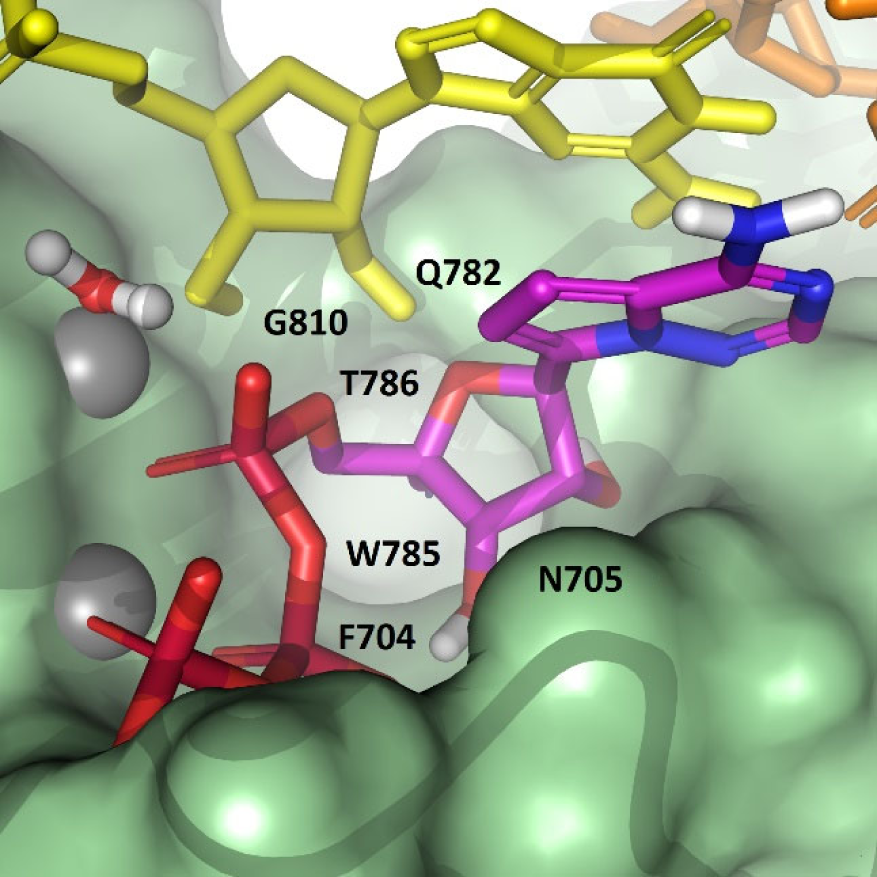
Model of TSV and an elongating RNA with **2-NTP** in its pre-incorporation state. The model was constructed from a cryo-EM structure of the apo RSV L-protein (6PZK), guided by the ternary structure for Lassa (7OJN). A pocket under the inhibitor ribose accommodates the 4’-CN substitution.

The intrinsic antiviral potency of **2-NTP** in contrast to the modest antiviral activity of **2** in cells suggested that the metabolism of **2** was limited in the cell types evaluated. The key question was which phosphorylation step for **2-NTP** formation was rate limiting as other 4’-subsituted nucleosides (e.g., **4**) have been reported to have a slow diphosphate formation step.^46^ Thus, **2** was profiled with phosphoramidate prodrugs to determine if improved antiviral activity could be achieved through direct NMP delivery inside cells. Gratifyingly, the prodrugs exhibited marked improvements in potency (Table 2) implicating the NMP formation step as rate limiting. Since administration by inhalation was targeted for direct delivery to the respiratory epithelium, further phosphoramidate prodrug optimization focused primarily on achieving high potency and rapid intracellular NTP formation in lung cells.

**Table 2.** In Vitro Profiles of Phosphoramidate and 2’,3’-Ester Phosphoramidate Prodrugs.*^a^*.

| Cpd. | R<sup>1</sup> | R<sup>3</sup> | RSV NHBE | RSV HEp-2 | HEp-2 | MT4 | LogD<sup>e</sup> | 2-NTP Conc. |
| --- | --- | --- | --- | --- | --- | --- | --- | --- |
| | | | EC<sub>50</sub> (μM)<sup>b</sup> | EC<sub>50</sub> (μM)<sup>c</sup> | CC<sub>50</sub> (μM)<sup>d</sup> | CC<sub>50</sub> (μM)<sup>d</sup> | | NHBE<br>(pmol/$10^6$ cells)<sup>f</sup> |
| **28a** | Me | - | 0.295 ± 0.133 | 0.122 ± 0.080 | >50 | >38 | 1.1 | - |
| **28b** | Et | - | 0.932 (n=1) | 0.095 ± 0.027 | >100 | 28.7 ± 4.5 | 1.1 | - |
| **28c** | *n*Pr | - | 0.168 ± 0.044 | 0.055 ± 0.019 | >50 | 10.3 ± 1.5 | 1.2 | - |
| **28d** | *i*Pr | - | 0.933 ± 0.340 | 0.391 ± 0.254 | >50 | >57 | 1.3 | - |
| **28e** | 2EtBu | - | 0.054 ± 0.006 | 0.029 ± 0.011 | >95 | 40.1 ± 8.8 | 2.1 | - |
| **28f** | cHex | - | 0.141 ± 0.117 | 0.148 ± 0.167 | >100 | >50 | 2.2 | 13 ± 4 |
| **28f-Rp** | cHex | - | 0.630 (n=1) | 3.11 ± 1.43 | >50 | >50 | - | - |
| **28g** | 4-THP | - | >4.3 | 1.49 ± 0.06 | >50 | >57 | 1.3 | - |
| **1** | Me | isobutyrate | 0.015 ± 0.007 | 0.046 ± 0.019 | 22.7 ± 4.6 | 3.24 ± 0.51 | 3.0 | 916 ± 324 |
| **29am** | Me | propionate | 0.019 ± 0.004 | 0.092 ± 0.006 | 30.0 ± 2.6 | 3.33 ± 0.21 | 2.2 | 384 ± 113 |
| **29an** | Me | acetate | 0.051 ± 0.019 | 0.127 ± 0.074 | >42 | 10.6 ± 2.2 | 2.0 | - |
| **29cl** | nPr | isobutyrate | 0.046 ± 0.009 | 0.036 ± 0.021 | 15.7 ± 4.3 | 4.92 ± 1.91 | 3.3 | 402 ± 154 |
| **29cm** | nPr | propionate | 0.056 ± 0.024 | 0.041 ± 0.019 | 27.3 ± 2.8 | 7.65 ± 1.88 | 2.8 | 165 (n=1) |
| **29fl** | cHex | isobutyrate | 0.017 ± 0.005 | 0.060 ± 0.019 | 18.1 ± 3.1 | 8.70 ± 2.92 | 4.0 | 170 (n=1) |
| **29fm** | cHex | propionate | 0.041 ± 0.021 | 0.060 ± 0.013 | 33.9 ± 3.0 | 17.1 ± 5.4 | 3.5 | - |
| **29fn** | cHex | acetate | 0.023 ± 0.006 | 0.206 ± 0.064 | >49 | 29.8 ± 2.5 | 2.9 | 40 (n=1) |

<sup>a</sup>All data are n ≥ 2 replicates and reported as average ± SD unless noted otherwise. <sup>b</sup>EC<sub>50</sub> values were determined in a 96-well format assay using NHBE cells infected with RSV-A2 with CPE readout at a MOI of 1. <sup>c</sup>EC<sub>50</sub> values were determined in a 384-well format assay using HEp-2 cells infected with RSV-A2 with CPE readout at a MOI of 1. <sup>d</sup>CC<sub>50</sub> values were determined in a 384 well format using HEp-2 or MT4 cells. <sup>e</sup>(n=1). <sup>f</sup>Intracellular **2-NTP** concentration in NHBE cells at 2 h from 10 μM incubation.

Phosphoramidate prodrugs **28a-28g** were profiled for anti-RSV potency in NHBE and HEp-2 cells, and cytotoxicity in the HEp-2 and MT4 cell lines (Table 2). Compared to **2**, substantial potency gains were realized from the 5’-phosphoramidate prodrug series while maintaining low cytotoxicity. The Me **28a** and *n*Pr **28c** esters exhibited improved potency compared to Et ester **28b**, but the higher LogD 2EtBu **28e** exhibited the optimal potency in the series. For the proximally branched esters, improved potency was also observed for the higher LogD *c*Hex **28f** as compared to the *i*Pr **28d**. Furthermore, the *S*p isomer of the *c*Hex ester prodrug **28f** exhibited improved potency compared to the *R*p isomer **28f-*R*p**. The low LogD 4-tetrahydropyran (THP) ester **28g** was not tolerated and lost significant potency. Moving forward we also masked the 2’,3’-hydroxyls with esters to further evaluate lipophilicity across a LogD range and remove hydrogen bond donors to promote cellular uptake.

A series of 2’,3’-diisobutyrate, -dipropionate, or -diacetate esters **29fl-29cm** provided further improvements in potency compared to their respective 2’,3’-hydroxyl counterparts attributed, in part, to enhanced cell permeability (Table 2). The 2’,3’-diester substitution enabled tuning of the physiochemical properties with the 2’,3’-diisobutyrate generally exhibiting the optimal potency in the series. The most effective prodrug **1** in NHBE cells exhibited high potency (EC_50_ = 0.015 ± 0.007 µM) with corresponding low cytotoxicity (NHBE CC_50_ = 8.3 µM, data not shown in Table 2) thereby achieving a 640-fold selectivity index (CC_50_/EC_50_). The anti-RSV activity of the prodrug **1** in NHBE cells represents a 400-fold improvement compared to the corresponding parent nucleoside **2** and a three-fold improvement over **6** (EC_50_ = 0.049 µM^23^). Similarly, **1** exhibited potent activity (EC_50_ = 0.046 ± 0.019 µM) and a 500-fold selectivity index in HEp-2 cells.

To maximize delivery of the **2-NMP** into lung cells via inhalation administration, rapid cellular uptake, and efficient first-pass pulmonary metabolism of the prodrugs were desirable.^47,48^ Toward this end, the levels of intracellular NTP were evaluated at an early timepoint of 2-h following incubation of prodrugs at 10 µM in NHBE cell culture as a part of the screening paradigm (Table 2). This assay was designed to understand whether NTP can form rapidly in lung cells from a “pulse-like” inhaled dose in contrast to the cellular antiviral activity assays that are conducted with continuous compound incubation over 4 d. Among the series of prodrugs, **1** rapidly delivered the highest intracellular NTP (916 ± 324 pmol/10^6^ cells). The proposed **1** metabolic pathway is described in Figure 3, where the **1** prodrug moieties putatively facilitate cellular uptake followed by global hydrolase ester cleavage to afford the alanine metabolite. Phosphoramidase or lysosomal acid-mediated cleavage of the P-N bond generates the **2-NMP**, which is then anabolized via nucleotide kinases to the active **2-NTP** leading to RSV RdRp inhibition. The parent **2** is likely limited by a slow first phosphorylation step to generate **2-NMP** resulting in lower levels of intracellular **2-NTP**.

The anti-RSV activity of **1** was further profiled in the primary 3D mucociliary tissue culture of the human airway epithelium (HAE) shown in Figure 5. Treatment was conducted from either the basolateral- or apical-facing surfaces, the latter to recapitulate topical delivery from inhalation, and the RSV RNA levels were measured by RT-qPCR in the apical supernatants at 72 h.^49^ Comparable concentration-dependent declines of RSV levels were observed from incubations with 0.01, 0.05, and 0.25 µM of **1**, where the 0.25 µM-treatment group reached median viral load reductions of 2.5log_10_ (basolateral) and 1.9log_10_ (apical). The cumulative data in primary NHBE cultures and HAE model supported **1** as a potent inhibitor of RSV replication in lung cells.

**Figure 5.**
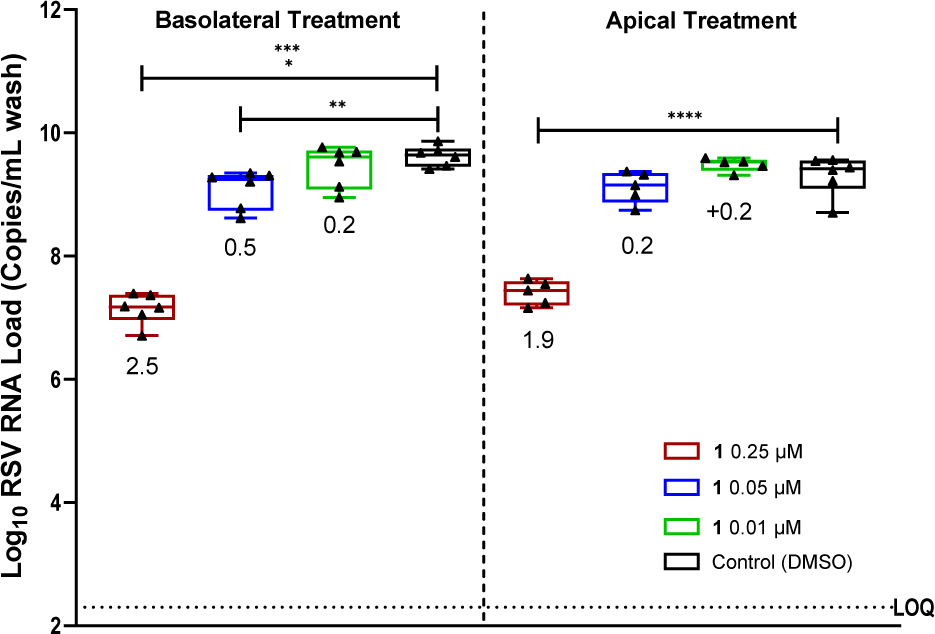
RSV viral RNA loads in HAE cultures following daily treatment with fresh media and compound on the basolateral (24-h incubation) or apical (3-h incubation) compartment with **1** or DMSO control. Compound treatment was initiated concurrently with infection at a MOI=0.1. Presented are box and whisker plots of the median and range of six technical replicates for each treatment. Values below the box and whisker plot represent the mean log reduction of the treatment group compared to the respective DMSO control. ** 0.01 < *p* < 0.001 **** *p* < 0.0001; *p*-Values were determined by the ordinary One-Way ANOVA with multiple comparisons using Dunnett’s correction.

To assess the spectrum of antiviral activity, the in vitro activity of **1** was evaluated against representative respiratory viruses from the *Pneumoviridae*, *Picornaviridae*, *Orthomyxoviridae*, and *Coronaviridae* families in various cell lines (Table 3). **1** exhibited potent activity against pneumoviruses RSV and hMPV, and also potently inhibited picornavirus RV-A, RV-B, EV-D68, and EV-71. In particular, the anti-RV activity of the prodrug **1** in H1 HeLa cells represents a several-fold improvement over the reported potency of **6** (RV-A 16 EC_50_ = 0.75 µM and RV-B 14 EC_50_ = 0.385 ± 0.318 µM^22^). **1** was inactive against influenza A (at 2 µM) or had low potency against SARS-CoV-2 (EC_50_ = 4.75 ± 1.35 µM). Importantly, **1** demonstrated broad-spectrum potent activity against pneumo- and picornaviruses commonly associated with disease exacerbation for people with asthma or COPD.

**Table 3.** Antiviral activities 1 against respiratory viruses in cell lines.*^a^*.

| Virus family | Virus | Cells | EC<sub>50</sub> (μM) | CC<sub>50</sub> (μM) |
| --- | --- | --- | --- | --- |
| *Pneumo* | RSV A2<sup>b</sup> | HEp-2 | 0.046 ± 0.019 | 22 |
|  | hMPV A1<sup>c</sup> | LLC-MK2 | 0.210 ± 0.050 | >2 (n=1) |
| *Picorna* | RV-A 16<sup>c</sup> | H1 HeLa | 0.061 ± 0.006 | 22.2 ± 6.3 |
|  | RV-B 14<sup>c</sup> | H1 HeLa | 0.054 ± 0.002 | 22.2 ± 6.3 |
|  | EV-D68<sup>c</sup> | RD | 0.090 ± 0.060 | >2 |
|  | EV-71<sup>c</sup> | RD | 0.083 ± 0.035 | >2 |
| *Orthomyxo* | FluA H1N1<sup>c</sup> | MDCK | >2 | >2 (n=1) |
| *Corona* | SARS-CoV-2<sup>c</sup> | A549-hACE2 | 4.75 ± 1.35 | 21.1 ± 4.4 |

<sup>a</sup>All data are n ≥ 2 replicates and reported as average ± SD unless noted otherwise. <sup>b</sup>Table 2. <sup>c</sup>Assay protocols provided in the SI.

The in vitro intracellular metabolism of **1** was also evaluated in respiratory cells to correlate the RSV antiviral activity to the concentrations of **2-NTP** formed and determine the half-life (Figure 6, Table 4). Primary human bronchial epithelial cells (HBE) from normal and diseased (DHBE) COPD or asthma donors treated with 1 µM of **1** yielded high concentrations of intracellular **2-NTP**. No meaningful differences in **2-NTP** levels were observed between NHBE and DHBE cells incubated with either compound, suggesting metabolic pathways are not disturbed in individuals with COPD or asthma (Figure 6A). The **2-NTP** half-life in NHBEs was assessed via a 2-h pulse incubation with 200 nM of **1** followed by media replacement and monitoring intracellular **2-NTP** concentration for 72 h. The terminal half-life of **2-NTP** was 21 h (Figure 6B), reflecting a potential for once-daily dosing in vivo. Metabolism from a 48-h continuous incubation of 200 nM of **1** was also determined in the H1 HeLa cells permissive to RV infection (Figure 6C). From these NTP formation assays, an estimate of the average intracellular concentration of **2-NTP** present in cells when 50% of the virus replication is inhibited (intracellular IC_50_), can be determined. This provides an physiologically relevant cellular readout to corroborate the biochemical IC_50_ assay. The RSV antiviral potency for **1** was evaluated in the HBE cell types with an RSV firefly luciferase (Fluc) reporter assay (Table 4). **1** was potent against RSV Fluc with EC_50_ values of <7 nM across donors. The calculated intracellular IC_50_ values were comparable across normal and diseased HBEs at <100 nM for **1** (Table 4) with good agreement to the RSV biochemical data (IC_50_ = 0.05 ± 0.04 µM, Table 1). Furthermore, an intracellular IC_50_ of 84 nM was calculated for RV based on the antiviral potency of **1** in H1 HeLa cells, which also aligned with the RV biochemical data (IC_50_ = 43 nM, data not shown). From this dataset, the intrinsic inhibition of **2-NTP** was determined to be similar for both RSV and RV providing confidence that both viruses could be effectively inhibited in vivo.

**Figure 6.**
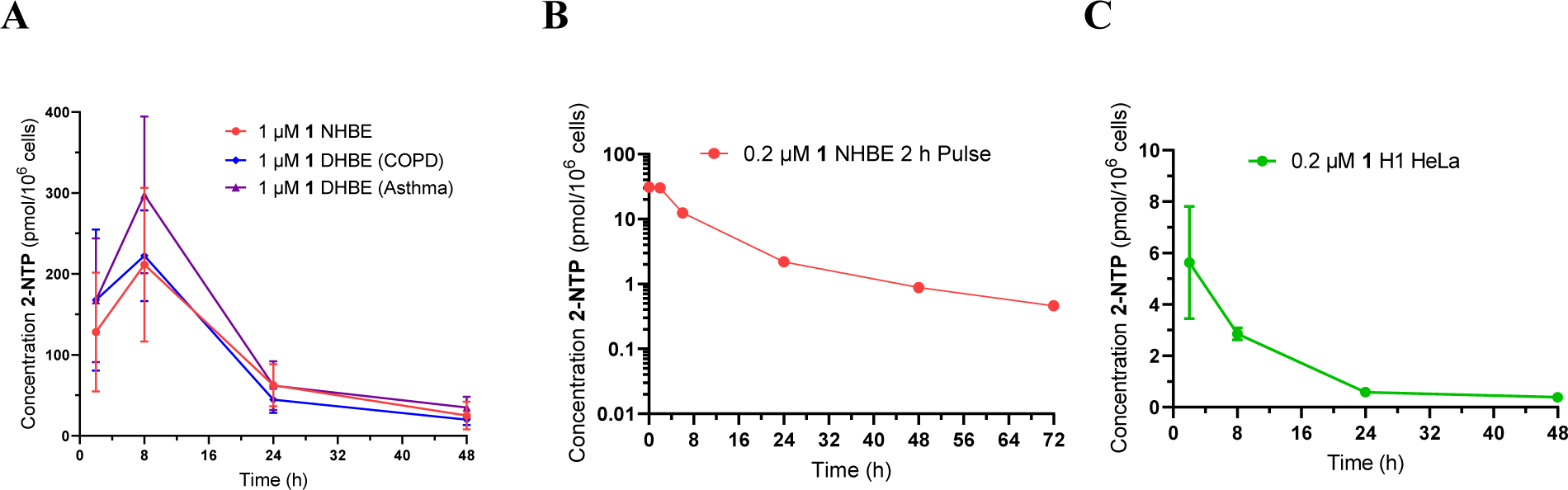
Intracellular **2-NTP** concentrations from incubation with **1**. (A) **1** (1 µM) continuous incubation over 48 h in normal and diseased, COPD and asthma, human bronchial epithelial cells. (B) **1** (0.2 µM) 2-h pulse incubation normal human bronchial epithelial cells. (C) **1** (0.2 µM) continuous incubation over 48 h in H1 HeLa cells.

**Table 4.** In vitro 1 antiviral activity and metabolism in primary respiratory cells.

| Cell Type | RSV Fluc<br>EC<sub>50</sub> (μM)<sup>a</sup> | RVA-16<br>EC<sub>50</sub> (μM)<sup>b</sup> | Incubation<br>Conc. **1** (μM) | [**2**-NTP]<br>(pmol/10<sup>6</sup> cells)<sup>c</sup> | Intracellular<br>IC<sub>50</sub> (μM)<sup>d</sup> |
| --- | --- | --- | --- | --- | --- |
| NHBE | <0.0031<sup>e</sup> | - | 1 | 88.8 | <0.038 |
| COPD HBE | <0.0066<sup>e</sup> | - | 1 | 85.0 | <0.085 |
| Asthma HBE | 0.0058 ± 0.0014 | - | 1 | 113.3 | 0.097 |
| H1 HeLa | - | 0.061 ± 0.006 | 0.2 | 0.84 | 0.084 |

<sup>a</sup>EC<sub>50</sub> data represent the average ± SD of two experiments determined in a 96-well format assay using NHBE and DHBE in an RSV Fluc reporter assay at a MOI of 0.1. <sup>b</sup>Table 3. <sup>c</sup>Average intracellular **2**-NTP following continuous incubation **1** at 1 μM in HBE or 0.2 μM in H1 HeLa cells over 48 h. <sup>d</sup>Estimated **2**-NTP concentrations in cells at EC<sub>50</sub> concentrations (intracellular IC<sub>50</sub>) calculated using the average cell volumes of 7 pL/cell for HBEs and 3 pL/cell for H1 HeLa, calculated via the equation: intracellular IC<sub>50</sub> = [(Mean **2**-NTP (pmol/10<sup>6</sup> cells) / cell volume (pL/cell)] / [incubation conc. ( μM))] × [EC<sub>50</sub> conc. ( μM)]. <sup>e</sup>One replicate was below the lowest point in the concentration curve (0.0023 μM) thus average represented as < value of the other replicate.

Inhalation administration was targeted for topical delivery directly to the respiratory epithelium, the primary site of infection for respiratory viruses. Oral dosing of **1** in cynomolgus monkey (cyno) resulted in low oral bioavailability (<1%, data not shown) as expected due to high first-pass intestinal and/or hepatic extraction, and intravenous (IV) dosing is not practical for treatment in the outpatient setting. To establish proof-of-concept for pharmacokinetics (PK) and NTP lung levels following inhalation delivery, pulmonary administration of **1** was performed using intratracheal (IT) dosing to anesthetized, ventilated cynos and AGMs. A aqueous suspension formulation of crystalline **1** was aerosolized with a vibrating mesh nebulizer and presented to the LRT through an endotracheal tube. Target lung deposited doses were achieved by varying the exposure duration, and were calculated from ex-vivo deposition of **1** on collection filters with the assumption of a 100% deposition fraction. The median mass aerodynamic diameter of the aerosol was 1.34 µm ± 1.81 µm (cyno) or 1.55 ± 1.65 µm (AGM), predicted to attain uniform deposition across the large and small airways.^50,51^ The cyno plasma and lung tissue PK profiles for **1** IV and IT administration are shown in Figures 7a and 7b and Table 5. From a 30-min IV infusion at 5 mg/kg in cyno, **1** exhibits a short half-life (t_1/2_ = 0.34 ± 0.17 h) attributed to the breakdown of **1** in tissue and plasma resulting in persistent plasma exposures of the parent nucleoside **2** (t_1/2_ = 8.71 ± 3.01 h) and concentrations of **2-NTP** in lung tissue of 4.28 ± 1.32 nmol/g tissue at 24 h post-dose. In comparison to IV dosing, the plasma concentrations of **1** following IT dosing in cyno (lung deposited dose: 4.0 mg/kg) persisted with a longer half-life (t_1/2_ = 13.0 ± 0.6 h) relative to IV, which was attributed to dissolution-limited absorption of the lung deposited dose of **1**. IT delivery of aerosolized **1** directly to the lung resulted in approximately 6-fold higher **2-NTP** levels at 24 h compared to IV infusion in cynos. IV and IT dose-normalized exposures (AUC_0-24h_) were similar for both **1** and **2**; therefore, the higher lung **2-NTP** levels arising from IT dosing are attributed to first-pass metabolism as **1** penetrates the lung before reaching the plasma. IT administration of **1** in AGMs at a similar lung deposited dose level of 5.9 mg/kg provided comparable plasma concentration profiles of **1** and **2** as well as high lung tissue concentrations of **2-NTP** (Figure 7c, Table 5). The exposures achieved from the IT delivered dose of **1** were sufficient to support moving forward in the RSV AGM efficacy model.

**Figure 7.**
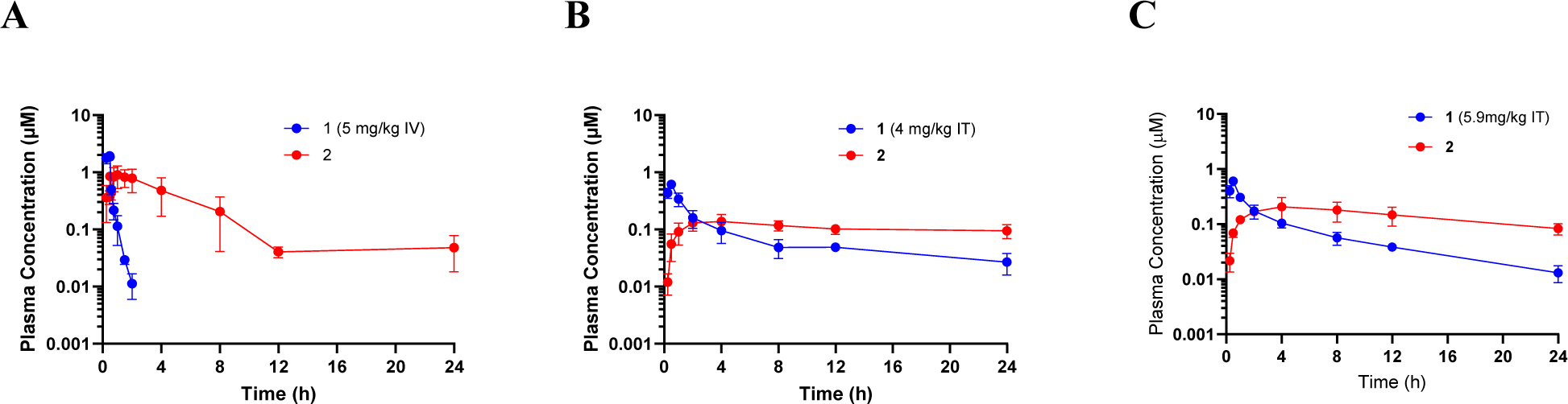
In vivo monkey plasma PK of **1** (blue) and **2** (red) following (A) 30 min IV infusion of 5 mg/kg **1** in cyno, (B) IT aerosol exposure at 4 mg/kg lung deposited dose of **1** in cyno, (C) IT aerosol exposure at 5.9 mg/kg lung deposited dose of **1** in AGM.

**Table 5.** In vivo cyno and AGM plasma and lung tissue PK parameters for IV infusion or IT aerosol administration of 1.*^a^*.

| Route | Species | Dose (mg/kg) | Analyte | $t_{1/2}$ (h) | $C_{max}$ (µM) | $AUC_{0-24h}$ (µM·h) | Lung 2-NTP<br>(nmol/g tissue)<sup>b</sup> |
| --- | --- | --- | --- | --- | --- | --- | --- |
| IV<sup>c</sup> | cyno<sup>d</sup> | 5 | **1** | 0.34 ± 0.17 | 1.99 ± 0.24 | 0.962 ± 0.127 | 4.28 ± 1.32 |
|  |  |  | **2** | 8.71 ± 3.01 | 0.961 ± 0.379 | 6.03 ± 3.75 |  |
| IT | cyno | 4.0<sup>e</sup> | **1** | 13.0 ± 0.6 | 0.62 ± 0.08 | 1.86 ± 0.44 | 20.5 ± 3.0 |
|  |  |  | **2** | 38.8 ± 4.0 | 0.14 ± 0.05 | 2.55 ± 0.61 |  |
| IT | AGM | 5.9<sup>e</sup> | **1** | 7.01 ± 0.49 | 0.61 ± 0.08 | 1.74 ± 0.21 | 24.6 ± 5.7 |
|  |  |  | **2** | 14.7 ± 2.6 | 0.22 ± 0.09 | 3.39 ± 1.11 |  |

<sup>a</sup>All values are the average ± SD from three monkeys unless otherwise noted. <sup>b</sup>Lung **2-NTP** was measured from the lower lung lobe tissue homogenate at 24 h post-dose. <sup>c</sup>30 min infusion. <sup>d</sup>n=6 cynos. <sup>e</sup>Lung deposited dose. Parameters: terminal half-life ( $t_{1/2}$ ; h); maximum concentration ( $C_{max}$ ; µM); area under the concentration–time curve from time 0 to 24 h ( $AUC_{0-24h}$ ; µM·h).

AGMs are semi-permissive for RSV replication and have been used to evaluate the efficacy of both vaccine candidates and small molecule inhibitors including nucleosides.^52,53^ We have previously reported that AGMs administered IV **6** at 10 mg/kg initiated 4 h prior to RSV infection (once daily for 6 days) resulted in a >2-log_10_ reduction of peak viral loads in the upper and lower respiratory tract.^31^ Here we conducted studies where AGMs were infected with RSV on day 0 and administered with either a nebulized vehicle or **1** crystalline suspension at 1.6 mg/kg or 7.6 mg/kg lung-deposited doses via intratracheal aerosol approximately 1 h post-infection (Figure 8). Dosing was continued once daily for 5 additional days (6 days total). Bronchoalveolar lavage (BAL) fluid and nasal swabs were collected at baseline, on study Day 1, and every other day thereafter until 15 days post-infection. On the days in which **1** was administered, the BAL and nasal sampling were performed approximately 1 h prior to dosing. RT-qPCR was subsequently performed on all samples to determine viral loads over the course of the infection. Compared to vehicle control, inhaled **1**, at both doses, resulted in significant reductions in RSV RNA copies in BALF samples collected between 3- and 11-days post-infection. At Day 5 post-infection, the final day of **1** treatment, an average 3.6 log_10_ reduction in viral load was observed for both the 1.6 and 7.6 mg/kg **1** lung deposited doses. A moderate rebound in BAL viral loads were observed 2 to 4 days after the final dose of **1**. Viral RNA levels then declined with similar kinetics in animals treated with **1** or vehicle; however, the viral loads remained lower in **1**-treated AGMs out to Day 11 of the study. With IT delivery, the nasopharyngeal epithelium was not directly exposed to aerosolized **1**. Thus, as anticipated, RSV viral load reductions in nasal swabs did not reach statistical significance in AGMs treated with **1** compared to vehicle (data not shown). Taken together, these results demonstrate that aerosolized **1** delivered to the LRT significantly reduced RSV replication in BAL samples (>99.9%), supporting that **1** was efficiently delivered to and metabolized by the LRT epithelial cells, the primary site of RSV infection associated with severe disease. Thus, we provide proof-of-concept in vivo efficacy to support inhaled **1** as an effective treatment for RSV in a nonhuman primate infection model.

**Figure 8.**
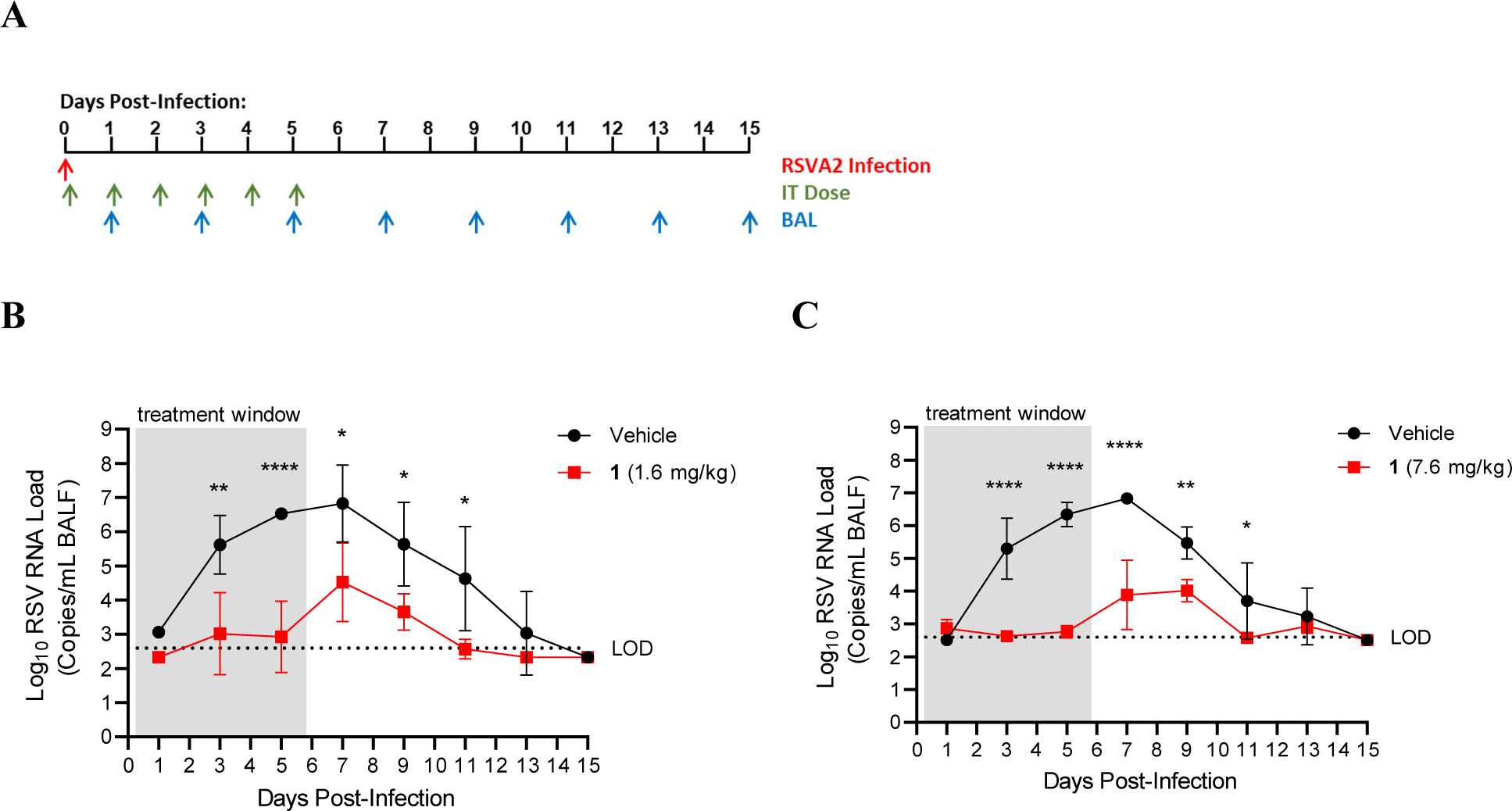
Efficacy in RSV A2-infected AGMs treated with **1** (red) or vehicle (black) by once daily IT aerosol beginning 1-h post-infection for 6 days (n=3-4 per treatment group). (A) Schematic of study design. (B) RSV viral RNA loads in bronchoalveolar lavage fluid (BALF) samples accessed by RT-qPCR for **1** (1.6 mg/kg lung deposited dose). (C) RSV viral RNA loads in BALF samples accessed by RT-qPCR for **1** (7.6 mg/kg lung deposited dose). Samples below the assay limit of detection (LOD, stripped lines) were assigned a value equal to ½ the LOD then log transformed for analysis. The average ± SD for each treatment group is shown. \**p* < 0.05, \*\**p* < 0.01, \*\*\**p* < 0.001, \*\*\*\**p* < 0.0001; data were analyzed by repeated measures two-way ANOVA with Bonferroni post-hoc correction for multiple comparisons.

## Conclusion

ARVIs are an important cause of morbidity and mortality worldwide. A SAR campaign identified a 4’-CN adenine *C*-ribonucleoside **2** with RSV antiviral activity. Its corresponding 5’-triphosphate metabolite **2-NTP** exhibited 1) potent intrinsic activity against the RSV RNP and RV RdRp, 2) excellent selectivity as it was neither an inhibitor nor substrate for human RNA or DNA polymerases, and 3) a long in vitro intracellular half-life of 21 h. Evaluation of phosphoramidate prodrugs of nucleoside **2** resulted in the discovery of **1** as a potent and selective inhibitor of pneumoviruses RSV and hMPV, and picornaviruses RV and EV in vitro. Intracellular metabolism analyses of **1** in primary lung cell cultures demonstrated rapid generation of high concentrations of **2-NTP**. Intratracheal aerosol administration of **1** to AGMs led to levels of active triphosphate **2-NTP** in respiratory tissues that resulted in robust in vivo efficacy in the RSV AGM challenge model. The overall profile supports further development of inhaled **1**, or alternate prodrugs of **2**, as a potential therapeutic for pneumo- and picornaviruses.

## EXPERIMENTAL SECTION

All organic compounds were synthesized at Gilead Sciences, Inc (Foster City, CA) unless otherwise noted. Commercially available solvents and reagents were used as received without further purification. Nuclear magnetic resonance (NMR) spectra were recorded on a Varian Mercury Plus 400 MHz at rt, with tetramethylsilane as an internal standard. Proton nuclear magnetic resonance spectra are reported in parts per million (ppm) on the *δ* scale and are referenced from the residual protium in the NMR solvent (CDCl_3_: *δ* 7.26, CD_3_OD: *δ* 3.31, DMSO-*d_6_*: *δ* 2.50, CD_3_CN: δ 1.94, D_2_O: δ 4.79). Data is reported as follows: chemical shift [multiplicity (s = singlet, d = doublet, t = triplet, q = quartet, p = pentet, sep = septet, m = multiplet, br = broad, app = apparent), coupling constants (*J*) in Hertz, integration. Phosphorus-31 nuclear magnetic resonance spectra are reported in parts per million on the *δ* scale. Data is reported as follows: chemical shift [multiplicity (s = singlet, d = doublet, t = triplet), coupling constants (*J*) in Hertz. No special nomenclature is used for equivalent phosphorus resonances. LC/MS was conducted on a Thermo Finnigan MSQ Std using electrospray positive and negative [M + 1]^+^ and [M – 1]^−^, and a Dionex Summit HPLC System (model: P680A HPG) equipped with a Gemini 5 u C18 110A column (30 mm × 4.60 mm), eluting with 0.05% formic acid in 1% acetonitrile/water and 0.05% formic acid in 99% acetonitrile/water. Preparative normal phase silica gel chromatography was conducted using a Teledyne ISCO CombiFlash Companion instrument with silica gel cartridges. All compounds are >95% pure by HPLC analysis. HPLC conditions to assess purity were as follows: Agilent 1100 Series HPLC, Phenominex Kinetex C18, 2.6 µm 100Å, 100 × 4.6 mm column; 2-98% gradient of 0.1% trifluoroacetic acid in water and 0.1% trifluoroacetic acid in acetonitrile; flow rate, 1.5 mL/min; acquisition time, 8.5 min; wavelength, UV 214 and 254 nm.

### 7-((2S,3S,4R,5R)-3,4-Bis(benzyloxy)-5-((benzyloxy)methyl)tetrahydrofuran-2-yl)pyrrolo[2,1-f][1,2,4]triazin-4-amine (11)

To a solution of **10**, previously synthesized,^24^ (4.74 g, 8.58 mmol) and triethylsilane (3.56 mL, 22.3 mmol), in DCM (43 mL) was added boron trifluoride diethyl etherate (1.59 mL, 12.9 mmol) slowly at 0 °C under an argon atmosphere. After 2 h, the reaction mixture was slowly diluted with saturated aqueous sodium bicarbonate solution (100 mL), and the resulting mixture was extracted with ethyl acetate (2 × 150 mL), was dried over anhydrous sodium sulfate, and was concentrated under reduced pressure. The crude residue was purified via silica gel chromatography (0–100% ethyl acetate/hexanes) to afford the title compound (3.94 g, 86%). ^1^H NMR (400 MHz, CDCl_3_) δ 7.88 (s, 1H), 7.37 – 7.22 (m, 15H), 6.73 (d, *J* = 4.6 Hz, 1H), 6.71 (d, *J* = 4.6 Hz, 1H), 5.66 (d, *J* = 4.2 Hz, 1H), 4.71 (s, 2H), 4.60 (d, *J* = 12.0 Hz, 1H), 4.54 (s, 2H), 4.45 (d, *J* = 11.9 Hz, 1H), 4.39 (dt, *J* = 7.1, 3.6 Hz, 1H), 4.25 (t, *J* = 4.6 Hz, 1H), 4.14 – 4.10 (m, 1H), 3.78 (dd, *J* = 10.7, 3.4 Hz, 1H), 3.65 (dd, *J* = 10.7, 4.0 Hz, 1H). LC/MS *m/z* = 537.41 [M+1].

### (2S,3R,4S,5R)-2-(4-aminopyrrolo[2,1-f][1,2,4]triazin-7-yl)-5 (hydroxymethyl)tetrahydrofuran-3,4-diol (12)

Compound **11** (2.64 g, 4.91 mmol) was dissolved in acetic acid (50 mL). The flask was purged with argon and 10% Pd/C (1.05 g, 0.982 mmol) was added. The flask was evacuated and backfilled with H_2(g)_ three times. The reaction mixture was stirred under an atmosphere of H_2(g)_. After 1h, the flask was purged with nitrogen and the reaction mixture was filtered through a celite pad with methanol washings. The filtrate was concentrated under reduced pressure and then co-evaporated with ethyl acetate followed by hexanes. The residue was placed under high vacuum to afford the title compound (1.31 g, 99%). Caution! **12** is cytotoxic. ^1^H NMR (400 MHz, DMSO-*d*_6_) δ 7.80 (s, 1H), 7.66 (s, 2H), 6.82 (d, *J* = 4.4 Hz, 1H), 6.66 (d, *J* = 4.4 Hz, 1H), 5.09 (d, *J* = 6.5 Hz, 1H), 5.06-4.56 (m, 3H), 4.21 (t, *J* = 5.9 Hz, 1H), 3.93 (t, *J* = 4.9 Hz, 1H), 3.77 (q, *J* = 4.5 Hz, 1H), 3.48 (ddd, *J* = 38.9, 11.8, 4.4 Hz, 2H). LC/MS *m/z* = 267.13 [M+1].

### ((3a*R*,4*R*,6*S*,6a*S*)-6-(4-Aminopyrrolo[2,1-*f*][1,2,4]triazin-7-yl)-2,2-dimethyltetrahydrofuro[3,4-*d*][1,3]dioxol-4-yl)methanol

Compound **12** (3.13 g, 11.7 mmol) was dissolved in acetone (80 mL) and TsOH (6.00 g, 31.5 mmol) was added. Triethylorthoformate (6.0 mL, 36.1 mmol) was slowly added over 10 min. The resulting mixture was allowed to stir at ambient temperature overnight. Saturated aqueous sodium carbonate solution was added until the reaction mixture was pH=8. The solids were removed by filtration and the filtrate was concentrated under reduced pressure. The crude residue was partitioned between ethyl acetate and brine. The phases were split and the organics were dried over anhydrous sodium sulfate, were filtered, and were concentrated under reduced pressure. The crude was purified by silica gel chromatography (60-100% ethyl acetate in hexanes to 20% methanol in ethyl acetate) to afford the title compound (2.55 g, 71%). ^1^H NMR (400 MHz, DMSO-*d*_6_) δ 7.83 (s, 1H), 7.71 (s, 2H), 6.83 (d, *J* = 4.4 Hz, 1H), 6.73 (d, *J* = 4.5 Hz, 1H), 5.21 (d, *J* = 4.9 Hz, 1H), 5.01 (dd, *J* = 6.6, 4.9 Hz, 1H), 4.84 (t, *J* = 5.7 Hz, 1H), 4.71 (dd, *J* = 6.7, 3.7 Hz, 1H), 3.99 – 3.85 (m, 1H), 3.46 (t, *J* = 5.5 Hz, 2H), 1.48 (s, 3H), 1.29 (s, 3H). LC/MS *m/z* = 307.21 [M+1].

### 7-((3aS,4S,6R,6aR)-6-(((tert-Butyldimethylsilyl)oxy)methyl)-2,2-dimethyltetrahydrofuro[3,4-d][1,3]dioxol-4-yl)pyrrolo[2,1-f][1,2,4]triazin-4-amine

((3a*R*,4*R*,6*S*,6a*S*)-6-(4-Aminopyrrolo[2,1-*f*][1,2,4]triazin-7-yl)-2,2-dimethyltetrahydrofuro[3,4-*d*][1,3]dioxol-4-yl)methanol (2.55 g, 8.32 mmol) was dissolved in DCM (50 mL) and the mixture was cooled to 0 °C. Imidazole (1.70 g, 24.9 mmol) was added followed by TBSCl (1.88 g, 12.5 mmol). After 16 h, the reaction was quenched with methanol. The resulting mixture was concentrated under reduced pressure and the crude residue was partitioned between water and ethyl acetate. The organics were dried over anhydrous sodium sulfate, were filtered, and were concentrated under reduced pressure. The crude residue was purified by silica gel chromatography (50-100% ethyl acetate in hexanes) to afford the title compound (2.60 g, 74%). ^1^H NMR (400 MHz, DMSO-*d*_6_) δ 7.83 (s, 1H), 7.74 (s, 2H), 6.82 (d, *J* = 4.4 Hz, 1H), 6.68 (d, *J* = 4.4 Hz, 1H), 5.26 (d, *J* = 4.4 Hz, 1H), 5.00 (dd, *J* = 6.5, 4.5 Hz, 1H), 4.71 (dd, *J* = 6.5, 3.7 Hz, 1H), 3.97 (td, *J* = 5.1, 3.6 Hz, 1H), 3.64 (d, *J* = 5.2 Hz, 2H), 1.48 (s, 3H), 1.28 (s, 3H), 0.83 (s, 9H), −0.02 (s, 6H). LC/MS *m/z* = 421.60 [M+1].

### *tert-*Butyl (7-((3a*S*,4*S*,6*R*,6a*R*)-6-(((*tert-*butyldimethylsilyl)oxy)methyl)-2,2-dimethyltetrahydrofuro[3,4-*d*][1,3]dioxol-4-yl)pyrrolo[2,1-*f*][1,2,4]triazin-4-yl)carbamate

7-((3a*S*,4*S*,6*R*,6a*R*)-6-(((*tert-*Butyldimethylsilyl)oxy)methyl)-2,2-dimethyltetrahydrofuro[3,4-*d*][1,3]dioxol-4-yl)pyrrolo[2,1-*f*][1,2,4]triazin-4-amine (2.59 g, 6.16 mmol) was dissolved in THF (60 mL) and the resulting solution was cooled to 0 °C. BOC_2_O (2.69 g, 12.3 mmol) and DMAP (0.3 g, 2.46 mmol) were then added. Triethylamine (2.56 mL, 18.3 mmol) was slowly added and the reaction mixture was allowed to warm to rt. After 3 h, the reaction mixture was cooled to 0 °C and methanol (10 mL) was added followed conc. NH_4_OH_(aq)_ (50 mL). The resulting mixture was allowed to warm to rt and was stirred overnight. The reaction mixture was concentrated under reduced pressure and the crude residue was partitioned between ethyl acetate and water. The layers were separated and the organic layer was dried over anhydrous sodium sulfate, was filtered, and was concentrated under reduced pressure. The crude residue was purified by silica gel chromatography (0-100% ethyl acetate in hexanes) to afford the title compound (2.82 g, 88%). ^1^H NMR (400 MHz, DMSO-*d*_6_) δ 10.46 (s, 1H), 8.20 (s, 1H), 7.19 (d, *J* = 4.6 Hz, 1H), 6.90 (d, *J* = 4.6 Hz, 1H), 5.33 (d, *J* = 4.2 Hz, 1H), 5.02 (dd, *J* = 6.5, 4.3 Hz, 1H), 4.72 (dd, *J* = 6.5, 3.6 Hz, 1H), 4.01 (q, *J* = 5.0 Hz, 1H), 3.64 (d, *J* = 5.1 Hz, 2H), 1.49 (s, 9H), 1.32 (d, *J* = 22.7 Hz, 6H), 0.82 (s, 9H), −0.03 (s, 6H). LC/MS *m/z* = 521.27 [M+1].

### *tert-*Butyl (7-((3a*S*,4*S*,6*R*,6a*R*)-6-(hydroxymethyl)-2,2-dimethyltetrahydrofuro[3,4-*d*][1,3]dioxol-4-yl)pyrrolo[2,1-*f*][1,2,4]triazin-4-yl)carbamate (13)

*tert-*Butyl (7-((3a*S*,4*S*,6*R*,6a*R*)-6-(((*tert-*butyldimethylsilyl)oxy)methyl)-2,2-dimethyltetrahydrofuro[3,4-*d*][1,3]dioxol-4-yl)pyrrolo[2,1-*f*][1,2,4]triazin-4-yl)carbamate (2.8 g, 5.4 mmol) was dissolved in THF (50 mL), and TBAF (1.0M in THF, 5.92 mL, 5.92 mmol) was added. After 30 min, additional TBAF (1.0M in THF, 5.92 mL, 5.92 mmol) was added. After another 30 min, the reaction mixture was quenched with water and the resulting mixture was extracted with ethyl acetate (2 ×). The combine the organic layers were washed with brine, were dried over anhydrous sodium sulfate, were filtered, and were concentrated under reduced pressure. The crude residue was purified by silica gel chromatography (10-100% ethyl acetate in hexanes) to afford the title compound (2.19 g, 86%). ^1^H NMR (400 MHz, DMSO-*d*_6_) δ 10.46 (s, 1H), 8.22 (s, 1H), 7.20 (s, 1H), 6.95 (s, 1H), 5.29 (d, *J* = 4.6 Hz, 1H), 5.03 (dd, *J* = 6.6, 4.7 Hz, 1H), 4.85 (t, *J* = 5.7 Hz, 1H), 4.72 (dd, *J* = 6.6, 3.6 Hz, 1H), 4.05 – 3.90 (m, 1H), 3.46 (t, *J* = 5.6 Hz, 2H), 1.50 (s, 12H), 1.29 (s, 3H). LC/MS *m/z* = 407.05 [M+1].

### *tert-*Butyl (7-((3a*S*,4*S*,6a*S*)-6,6-bis(hydroxymethyl)-2,2-dimethyltetrahydrofuro[3,4-*d*][1,3]dioxol-4-yl)pyrrolo[2,1-*f*][1,2,4]triazin-4-yl)carbamate (14)

Compound **13** (1.78 g, 4.38 mmol) was dissolved in DMSO (20 mL) and toluene (15mL). Pyridine (0.35 mL, 4.38 mmol) and EDCI (1.26 g, 6.56 mmol) were added followed by TFA (0.178 mL, 2.39 mmol). After 90 min, additional pyridine (0.35 mL, 4.38 mmol) and EDCI (1.26 g, 6.56 mmol) were added and the reaction mixture was stirred for an additional 30 min. The reaction was quenched with water and the resulting mixture was extracted with DCM. The aqueous was back extracted with DCM. The organic layers were combined and were washed with brine, were dried over anhydrous sodium sulfate, were filtered, and were concentrated under reduced pressure. The crude was put under high vacuum for 15 min then was used as is for the next step. The crude residue was dissolved in dioxane (15 mL) and formaldehyde (37% in water, 5.0 mL, 37.2 mmol) and 2N NaOH (5.34 mL, 10.7 mmol) were added sequentially. After 10 min, the reaction was quenched with AcOH and the resulting mixture was partitioned between saturated aqueous sodium bicarbonate solution and DCM. The aqueous layer was back extracted with DCM. The organic layers were combined and were wash with brine, were dried over anhydrous sodium sulfate, were filtered, and were concentrated under reduced pressure. The crude was placed under high vacuum for 15 min then were taken directly into the next reaction. The crude residue was dissolved in EtOH (50 mL), and NaBH_4_ (0.324 g, 8.76 mmol) was added in small portions. After 20 min, the reaction mixture was quenched with AcOH and was concentrated under reduced pressure. The crude residue was partitioned between with ethyl acetate and saturated aqueous sodium bicarbonate solution. The organic layer was split, was dried over anhydrous sodium sulfate, was filtered, and was concentrated under reduced pressure. The crude residue was purified by silica gel chromatography (50-100% ethyl acetate in hexanes) to afford the title compound (1.91 g, 68%). ^1^H NMR (400 MHz, DMSO-*d*_6_) δ 10.45 (s, 1H), 8.20 (s, 1H), 7.19 (d, *J* = 4.3 Hz, 1H), 6.95 (d, *J* = 4.7 Hz, 1H), 5.35 (d, *J* = 5.2 Hz, 1H), 5.06 (t, *J* = 5.7 Hz, 1H), 4.79-4.74 (m, 2H), 4.45 (t, *J* = 5.8 Hz, 1H), 3.73 – 3.46 (m, 3H), 3.40 – 3.30 (m, 1H), 1.50 (s, 12H), 1.27 (s, 3H). LC/MS *m/z* = 437.09 [M+1].

### *tert-*Butyl (7-((3a*S*,4*S*,6*S*,6a*S*)-6-((bis(4-methoxyphenyl)(phenyl)methoxy)methyl)-6-(hydroxymethyl)-2,2-dimethyltetrahydrofuro[3,4-*d*][1,3]dioxol-4-yl)pyrrolo[2,1-*f*][1,2,4]triazin-4-yl)carbamate

Compound **14** (1.15 g, 2.63 mmol) was dissolved in DCM (50 mL) and TEA (0.73 mL, 5.27 mmol) was added. The resulting solution was cooled to 0 °C and DMTrCl (1.35 g, 3.95 mmol) was added. After 10 min, the reaction mixture was quenched with methanol as was then diluted with DCM. The resulting mixture was washed with saturated aqueous sodium bicarbonate solution and brine. The organic layer was dried over anhydrous sodium sulfate, was filtered, and was concentrated under reduced pressure. The crude residue was purified by silica gel chromatography (0-100% ethyl acetate in hexanes) to afford to afford the title compound (1.95 g, 79%). ^1^H NMR (400 MHz, DMSO-*d*_6_) δ10.46 (s, 1H), 8.23 (s, 1H), 7.56 – 7.07 (m, 10H), 7.07 – 6.70 (m, 5H), 5.24 (d, *J* = 5.2 Hz, 1H), 5.04 (t, *J* = 5.9 Hz, 1H), 4.93 – 4.71 (m, 2H), 3.80 – 3.59 (m, 7H), 3.52 (dd, *J* = 10.9, 4.8 Hz, 1H), 3.25 (d, *J* = 9.9 Hz, 1H), 3.09 (d, *J* = 9.9 Hz, 1H), 1.50 (s, 9H), 1.25 (s, 3H), 1.21 (s, 3H). LC/MS *m/z* = 739.28 [M+1].

### *tert-*Butyl (7-((3a*S*,4*S*,6*R*,6a*S*)-6-((bis(4-methoxyphenyl)(phenyl)methoxy)methyl)-6-(((*tert-* butyldimethylsilyl)oxy)methyl)-2,2-dimethyltetrahydrofuro[3,4-*d*][1,3]dioxol-4-yl)pyrrolo[2,1-*f*][1,2,4]triazin-4-yl)carbamate

*tert-*Butyl (7-((3a*S*,4*S*,6*S*,6a*S*)-6-((bis(4-methoxyphenyl)(phenyl)methoxy)methyl)-6-(hydroxymethyl)-2,2-dimethyltetrahydrofuro[3,4-*d*][1,3]dioxol-4-yl)pyrrolo[2,1-*f*][1,2,4]triazin-4-yl)carbamate (1.53 g, 2.08 mmol) was dissolved in DMF (10 mL) and imidazole (0.42 g, 6.23 mmol) was added followed by TBSCl (0.47 g, 3.11 mmol). After 1 h, the reaction was quenched with methanol and partitioned between ethyl acetate and 5% LiCl solution. The phases were split and the organic layer was washed with brine, was dried over anhydrous sodium sulfate, was filtered, and was concentrated under reduced pressure. The crude residue was purified by silica gel chromatography (0-50% ethyl acetate in hexanes) to afford the title compound (1.77 g, 78%). ^1^H NMR (400 MHz, DMSO-*d*_6_) δ 10.47 (s, 1H), 8.23 (s, 1H), 7.56 – 6.66 (m, 15H), 5.31 (d, *J* = 4.9 Hz, 1H), 5.14 (dd, *J* = 6.5, 4.9 Hz, 1H), 4.73 (d, *J* = 6.5 Hz, 1H), 3.87 (d, *J* = 9.7 Hz, 1H), 3.72 (s, 6H), 3.53 (d, *J* = 9.7 Hz, 1H), 3.31 (m, 1H), 3.08 (d, *J* = 9.8 Hz, 1H), 1.50 (s, 9H), 1.25 (s, 3H), 1.22 (s, 3H), 0.75 (s, 9H), −0.04 (s, 3H), −0.08 (s, 3H). LC/MS *m/z* = 853.50 [M+1].

### *tert-*Butyl (7-((3a*S*,4*S*,6*R*,6a*S*)-6-(((*tert-*butyldimethylsilyl)oxy)methyl)-6-(hydroxymethyl)-2,2-dimethyltetrahydrofuro[3,4-*d*][1,3]dioxol-4-yl)pyrrolo[2,1-*f*][1,2,4]triazin-4-yl)carbamate (15)

*tert-*Butyl (7-((3a*S*,4*S*,6*R*,6a*S*)-6-((bis(4-methoxyphenyl)(phenyl)methoxy)methyl)-6-(((*tert-*butyldimethylsilyl)oxy)methyl)-2,2-dimethyltetrahydrofuro[3,4-*d*][1,3]dioxol-4-yl)pyrrolo[2,1-*f*][1,2,4]triazin-4-yl)carbamate (1.38 g, 1.62 mmol) was dissolved in chloroform (20 mL) and the resulting solution was cooled to 0 °C. A solution of TsOH (0.34 g, 1.78 mmol) in methanol (16 mL) was then add slowly. After 30 min, the reaction was quenched with saturated aqueous sodium bicarbonate solution, and the resulting mixture was partitioned between ethyl acetate and brine. The layers were split and the organic layer was dried over anhydrous sodium sulfate, was filtered, and was concentrated under reduced pressure. The crude residue was purified by silica gel chromatography (0-100% ethyl acetate in hexanes) to afford the title compound (0.84 g, 94%). ^1^H NMR (400 MHz, DMSO-*d*_6_) δ 10.42 (s, 1H), 8.20 (s, 1H), 7.19 (s, 1H), 6.89 (s, 1H), 5.37 (d, *J* = 4.8 Hz, 1H), 5.05 (dd, *J* = 6.2, 4.8 Hz, 1H), 4.71 (d, *J* = 6.2 Hz, 1H), 4.51 (t, *J* = 5.5 Hz, 1H), 3.70 (d, *J* = 10.2 Hz, 1H), 3.59 (d, *J* = 5.5 Hz, 2H), 3.49 (d, *J* = 10.2 Hz, 1H), 1.49 (s, 12H), 1.28 (s, 3H), 0.82 (s, 9H), −0.01 (s, 3H), −0.02 (s, 3H). LC/MS *m/z* = 551.25 [M+1].

### 7-((3aS,4S,6R,6aS)-6-(((tert-Butyldimethylsilyl)oxy)methyl)-2,2,6-trimethyltetrahydrofuro[3,4-d][1,3]dioxol-4-yl)pyrrolo[2,1-f][1,2,4]triazin-4-amine

To a solution of **15** (660 mg, 1.2 mmol), PPh_3_ (1.26 g, 4.8 mmol), and imidazole (327 mg, 4.8 mmol) in toluene (12 mL) was added I_2_ (610 mg, 2.4 mmol) at rt, and the resulting mixture was heated to 100 °C. After 24 h, the reaction mixture was allowed to cool to rt, diluted with ethyl acetate (50 mL), and washed with saturated aqueous sodium thiosulfate solution (2 × 50 mL), water (50 mL) and brine (50 mL). The organic layer was dried over anhydrous sodium sulfate and concentrated under reduced pressure. The crude residue was purified by silica gel chromatography (0-100% ethyl acetate in hexanes). The fractions containing the desired intermediate were collected and concentrated under reduced pressure. The residue was dissolved in methanol (12 mL) and triethylamine (0.2 mL) and 10% Pd/C (200 mg) were added at rt. The resulting mixture was stirred under a hydrogen gas atmosphere for 5 h, at which point the mixture was filtered through a pad of celite and concentrated under reduced pressure. The residue was dissolved in ethyl acetate (50 mL), washed with brine (2 × 50 mL), dried over anhydrous sodium sulfate, and concentrated under reduced pressure. The residue was dissolved in dioxane (8 mL) and water (2 mL) and was heated to 120 °C for 2 h. The mixture was then concentrated under reduced pressure, and the crude residue was purified by silica gel chromatography (0-50% ethyl acetate in hexanes) to afford the title compound (39 mg, 7%). ^1^H NMR (400 MHz, CDCl_3_) δ 7.95 (s, 1H), 6.71 (d, J = 4.6 Hz, 1H), 6.57 (d, J = 4.5 Hz, 1H), 5.77 (bs, 2H), 5.50 (d, J = 5.0 Hz, 1H), 5.10 (dd, J = 6.4, 5.0 Hz, 1H), 4.79 – 4.71 (m, 1H), 3.61 – 3.44 (m, 2H), 1.61 (s, 3H), 1.40 (s, 3H), 1.37 (s, 3H), 0.88 (s, 9H), 0.023 (s, 3H), 0.018 (s, 3H). LCMS *m/z* = 435.4 [M+1].

### ((3a*S*,4*R*,6*S*,6a*S*)-6-(4-Aminopyrrolo[2,1-*f*][1,2,4]triazin-7-yl)-2,2,4-trimethyltetrahydrofuro[3,4-*d*][1,3]dioxol-4-yl)methanol (16)

To a solution of 7-((3a*S*,4*S*,6*R*,6a*S*)-6-(((*tert-*butyldimethylsilyl)oxy)methyl)-2,2,6-trimethyltetrahydrofuro[3,4-*d*][1,3]dioxol-4-yl)pyrrolo[2,1-*f*][1,2,4]triazin-4-amine (90 mg, 0.20 mmol) in THF (5 mL) was added TBAF (1M in THF, 0.6 mL, 0.6 mmol) at 0 °C. The reaction mixture was allowed to warm to rt and was stirred for 16 h. The resulting mixture was diluted with ethyl acetate (20 mL), washed with brine (5 × 20 mL), dried over anhydrous sodium sulfate, and concentrated under reduced pressure. The crude residue was purified by silica gel chromatography (0-100% ethyl acetate in hexanes) to afford the title compound (49 mg, 76%). ^1^H NMR (400 MHz, CDCl_3_) δ 7.87 (s, 1H), 6.64 (d, J = 4.5 Hz, 1H), 6.56 (d, J = 4.4 Hz, 1H), 6.15 (bs, 2H), 5.27 (t, J = 6.5 Hz, 1H), 5.21 (bs, 1H), 5.09 (d, J = 6.9 Hz, 1H), 4.97 (d, J = 6.2 Hz, 1H), 3.68 (d, J = 11.7 Hz, 1H), 3.59 (d, J = 11.9 Hz, 1H), 1.62 (s, 3H), 1.35 (s, 3H), 1.32 (s, 3H). LCMS *m/z* = 321.2 [M+1].

### (2R,3S,4R,5S)-5-(4-Aminopyrrolo[2,1-f][1,2,4]triazin-7-yl)-2-(hydroxymethyl)-2-methyltetrahydrofuran-3,4-diol (20)

To a solution of **16** (25 mg, 0.08 mmol) in MeCN (2 mL) was added concentrated aqueous hydrochloric acid solution (12N, 0.15 mL) at 0 °C. After 1 h, the resulting mixture was diluted saturated aqueous sodium bicarbonate solution to achieve a pH=8 and was directly purified by purified preparatory HPLC (Gemeni C18 5uM 110Å 100 x 30 mm column, 5-100% acetonitrile in water gradient) to afford the title compound (18 mg, 82%). ^1^H NMR (400 MHz, D_2_O) δ 7.58 (s, 1H), 6.69 – 6.59 (m, 2H), 5.10 (d, J = 8.7 Hz, 1H), 4.65 (m, 1H), 4.12 (d, J = 5.5 Hz, 1H), 3.42 (s, 2H), 1.17 (s, 3H). MS *m/z* = 281.2 [M+1].

### *tert-*Butyl (7-((3a*S*,4*S*,6*R*,6a*S*)-6-(((*tert-*butyldimethylsilyl)oxy)methyl)-2,2-dimethyl-6-vinyltetrahydrofuro[3,4-*d*][1,3]dioxol-4-yl)pyrrolo[2,1-*f*][1,2,4]triazin-4-yl)carbamate

To a solution of **15** (980 mg, 1.78 mmol) in DMSO (0.76 mL) and DCM (20 mL) was added triethylamine (1.57 mL, 11.3 mmol), and sulfur trioxide pyridine complex (850 mg, 5.34 mmol) at 0 °C. The resulting mixture was allowed to warm to rt. After 16 h, the reaction mixture was cooled to 0 °C and water (0.5 mL) and sodium bicarbonate (1.5 g) was added. The resulting mixture was concentrated under reduced pressure and the crude residue was purified by silica gel chromatography (20-40% ethyl acetate in hexanes) to afford the aldehyde intermediate (864 mg, 88%). To a suspension of methyltriphenylphosphonium bromide (1.4 g, 5.1 mmol) in THF (25 mL) was added n-butyllithium (2.5 M in hexanes, 1.9 mL, 4.8 mmol) at −78 °C. After 1 h, the resulting mixture was warmed to 0 °C and a solution of aldehyde intermediate (864 mg, 1.57 mmol) in THF (5 mL) was added. The reaction mixture was allowed to warm to rt. After 16 h, saturated aqueous ammonium chloride solution (6 mL), ethyl acetate (10 mL) and hexanes (10 mL) were added. The phases were split and the organic layer was washed with water (20 mL), dried over anhydrous magnesium sulfate, and concentrated under reduced pressure. The crude residue was purified by silica gel chromatography (15-40% ethyl acetate in hexanes) to afford the title compound (670 mg, 78%). ^1^H NMR (400 MHz, CDCl_3_) δ 8.12 (s, 1H), 6.92 (s, 1H), 6.02 (dd, J = 17.4, 10.9 Hz, 1H), 5.62 (dd, J = 17.4, 1.8 Hz, 1H), 5.53 (d, J = 5.3 Hz, 1H), 5.31 (dd, J = 11.0, 1.8 Hz, 1H), 5.04 – 4.91 (m, 2H), 3.65 (d, J = 10.5 Hz, 1H), 3.57 (d, J = 10.5 Hz, 1H), 1.59 (s, 3H), 1.57 (s, 9H), 1.37 (s, 3H), 0.89 (s, 9H), 0.04 (s, 3H), 0.03 (s, 3H).

### ((3a*S*,4*R*,6*S*,6a*S*)-6-(4-Aminopyrrolo[2,1-*f*][1,2,4]triazin-7-yl)-2,2-dimethyl-4-vinyltetrahydrofuro[3,4-*d*][1,3]dioxol-4-yl)methanol (17)

To a solution of *tert-*butyl (7-((3a*S*,4*S*,6*R*,6a*S*)-6-(((*tert-*butyldimethylsilyl)oxy)methyl)-2,2-dimethyl-6-vinyltetrahydrofuro[3,4-*d*][1,3]dioxol-4-yl)pyrrolo[2,1-*f*][1,2,4]triazin-4-yl)carbamate (670 mg, 1.23 mmol) in THF (2 mL) was added TBAF (1 M in THF, 3.68 mL, 11.3 mmol) at rt. After 6 h, the reaction mixture diluted with ethyl acetate (20 mL), washed with water (20 mL), dried over anhydrous sodium sulfate, and concentrated under reduced pressure. The crude residue was purified by silica gel chromatography (20-50% ethyl acetate in hexanes) to afford the desilylated intermediate (485 mg, 92%). The desilylated intermediate (485 mg, 1.12 mmol) was taken up into dioxane (4 mL) and water (1 mL) and was heated to 120 °C. After 4 h, the resulting mixture was concentrated under reduced pressure and the crude residue was purified by silica gel chromatography (ethyl acetate) to afford the title compound (365 mg, 98%). ^1^H NMR (400 MHz, CD_3_CN) δ 7.86 (s, 1H), 6.80 – 6.72 (m, 2H), 6.28 (s, 2H), 6.01 (dd, J = 17.4, 10.9 Hz, 1H), 5.48 (dd, J = 17.4, 2.0 Hz, 1H), 5.24 (dd, J = 10.9, 2.0 Hz, 1H), 5.21 – 5.13 (m, 2H), 5.06 – 4.99 (m, 1H), 4.55 (dd, J = 10.7, 2.4 Hz, 1H), 3.62 (dd, J = 11.6, 10.7 Hz, 1H), 3.43 (dd, J = 11.6, 2.4 Hz, 1H), 1.53 (s, 3H), 1.35 – 1.31 (m, 3H).

### (2R,3S,4R,5S)-5-(4-Aminopyrrolo[2,1-f][1,2,4]triazin-7-yl)-2-(hydroxymethyl)-2-vinyltetrahydrofuran-3,4-diol (21)

To a solution of **17** (60.0 mg, 0.181 mmol) in MeCN (2 mL) was added concentrated aqueous hydrochloric acid solution (12N, 1.0 mL) at 0 °C. After 16 h, the resulting mixture was diluted saturated aqueous sodium bicarbonate solution to achieve a pH=8 and was directly purified by purified preparatory HPLC (Gemeni C18 5uM 110Å 100 x 30 mm column, 0-30% acetonitrile in water gradient) to afford the title compound (51.2 mg, 97%). ^1^H NMR (400 MHz, CD_3_CN) δ 7.86 (s, 1H), 6.78 (d, *J* = 4.5 Hz, 1H), 6.75 (d, *J* = 4.4 Hz, 1H), 6.27 (s, 2H), 6.00 (dd, *J* = 17.4, 10.9 Hz, 1H), 5.42 (dd, *J* = 17.4, 2.1 Hz, 1H), 5.22 (dd, *J* = 11.0, 2.2 Hz, 1H), 5.01 (d, *J* = 8.4 Hz, 1H), 4.78 – 4.69 (m, 1H), 4.57 (dd, *J* = 10.6, 2.7 Hz, 1H), 4.35 – 4.29 (m, 1H), 3.58 (t, *J* = 11.2 Hz, 1H), 3.44 – 3.36 (m, 2H), 3.33 (d, *J* = 4.4 Hz, 1H). LC/MS *m/z* = 293.19 [M+1].

### *tert-*Butyl (7-((3a*S*,4*S*,6*R*,6a*S*)-6-(((*tert-*butyldimethylsilyl)oxy)methyl)-6-cyano-2,2-dimethyltetrahydrofuro[3,4-*d*][1,3]dioxol-4-yl)pyrrolo[2,1-*f*][1,2,4]triazin-4-yl)carbamate

Compound **15** (0.838 g, 1.52 mmol) was dissolved in DMSO (5 mL) and toluene (3 mL). Pyridine (0.14 mL, 1.67 mmol) and EDCI (0.438 g, 2.28 mmol) were added followed by TFA (0.057 mL, 0.761 mmol). After 30 min, additional pyridine (0.14 mL, 1.67 mmol) and EDCI (0.438 g, 2.28 mmol) were added. After 1 h, additional pyridine (0.14 mL, 1.67 mmol) and EDCI (0.438 g, 2.28 mmol) were added. After 2 h, the reaction mixture was quenched with saturated aqueous sodium bicarbonate solution and was partitioned between ethyl acetate and saturated aqueous sodium bicarbonate solution. The layers were separated and the organic layer was dried over anhydrous sodium sulfate, was filtered, and was concentrated under reduced pressure. The residue was dissolved in DCM was concentrated under high vacuum for 1 h to afford a residue that was used directly in the next step. The residue was dissolved in pyridine (8 mL) and hydroxylamine hydrochloride (0.159 g, 2.28 mmol) was added in one portion. After 15 min, the reaction mixture was concentrated under reduced pressure and was partitioned between ethyl acetate and water. The organic layer was dried over anhydrous sodium sulfate, was filtered, and was concentrated under reduced pressure. The crude residue was placed under high vacuum for 30 min and used as is for the third step. The crude residue was dissolved in MeCN (8 mL). CDI (0.37 g, 2.28 mmol) was added in one portion. After 45 min, additional CDI (0.37 g, 2.28 mmol) was added. After 1 h, the reaction was quenched with saturated aqueous sodium bicarbonate solution. The crude was partitioned between ethyl acetate and saturated aqueous sodium bicarbonate solution. The layers were separated and the organic layer was dried over anhydrous sodium sulfate, was filtered, and was concentrated under reduced pressure. The crude residue was purified by silica gel chromatography (0-50% ethyl acetate in hexanes) to afford the title compound (0.72 g, 87%). ^1^H NMR (400 MHz, DMSO-*d*_6_) δ 10.53 (s, 1H), 8.25 (s, 1H), 7.21 (s, 1H), 7.00 (d, *J* = 4.6 Hz, 1H), 5.62 (d, *J* = 3.6 Hz, 1H), 5.28 (dd, *J* = 6.6, 3.7 Hz, 1H), 4.93 (d, *J* = 6.6 Hz, 1H), 3.83 (s, 2H), 1.62 (s, 3H), 1.50 (s, 9H), 1.33 (s, 3H), 0.83 (s, 9H), 0.00 (s, 6H). LC/MS *m/z* = 546.15 [M+1].

### (3aS,4R,6S,6aS)-6-(4-Aminopyrrolo[2,1-f][1,2,4]triazin-7-yl)-4-(((tert-butyldimethylsilyl)oxy)methyl)-2,2-dimethyltetrahydrofuro[3,4-d][1,3]dioxole-4-carbonitrile

*tert-*Butyl (7-((3a*S*,4*S*,6*R*,6a*S*)-6-(((*tert-*butyldimethylsilyl)oxy)methyl)-6-cyano-2,2-dimethyltetrahydrofuro[3,4-*d*][1,3]dioxol-4-yl)pyrrolo[2,1-*f*][1,2,4]triazin-4-yl)carbamate (0.688 g, 1.26 mmol) was dissolved in DCM (15 mL). Zinc bromide (0.567 g, 2.52 mmol) was added in one portion and the reaction mixture was stirred at ambient temperature. After 3 h, the reaction mixture was added to a silica load cartridge and was purified by silica gel chromatography (40-100% ethyl acetate in hexanes) to afford the title compound (0.56 g, 99%). ^1^H NMR (400 MHz, DMSO-*d_6_*) δ 7.86 (s, 1H), 7.80 (s, 2H), 6.85 (d, J = 4.5 Hz, 1H), 6.79 (d, J = 4.5 Hz, 1H), 5.55 (d, J = 3.7 Hz, 1H), 5.25 (dd, J = 6.6, 3.8 Hz, 1H), 4.92 (d, J = 6.6 Hz, 1H), 3.82 (s, 2H), 1.61 (s, 3H), 1.33 (s, 3H), 0.83 (s, 9H), −0.13 (s, 6H). LC/MS m/z = 446.68 [M+1].

### (3aS,4R,6S,6aS)-6-(4-Aminopyrrolo[2,1-f][1,2,4]triazin-7-yl)-4-(hydroxymethyl)-2,2-dimethyltetrahydrofuro[3,4-d][1,3]dioxole-4-carbonitrile (18)

(3a*S*,4*R*,6*S*,6a*S*)-6-(4-Aminopyrrolo[2,1-*f*][1,2,4]triazin-7-yl)-4-(((*tert-*butyldimethylsilyl)oxy)methyl)-2,2-dimethyltetrahydrofuro[3,4-*d*][1,3]dioxole-4-carbonitrile (8.41 g, 18.87 mmol) was taken up in THF (100 mL). Added TBAF 1.0 M in THF (28.31 mL, 28.31 mmol) in one portion at ambient temperature. Allowed to stir at ambient temperature for 10 min. The reaction mixture was quenched with water and the organics were removed under reduced pressure. The crude was partitioned between ethyl acetate and water. The layers were separated and the aqueous was washed with ethyl acetate. The organics were combined and dried over anhydrous sodium sulfate. The solids were filtered and the solvent removed under reduced pressure. The crude was purified by silica gel chromatography (0-10% methanol in DCM) to afford the title compound (5.83 g, 93%). ^1^H NMR (400 MHz, DMSO-*d*_6_) δ 7.87-7.80 (m, 3H), 6.85 (d, J = 4.5Hz, 1H), 6.82 (d, J = 4.5Hz, 1H), 5.74 (t, *J* = 5.8 Hz, 1H), 5.52 (d, *J* = 4.2 Hz, 1H), 5.24 (dd, *J* = 6.8, 4.2 Hz, 1H), 4.92 (d, *J* = 6.8 Hz, 1H), 3.65 (dd, *J* = 6.1, 1.7 Hz, 2H), 1.61 (s, 3H), 1.33 (s, 3H). LC/MS *m/z* = 332.14 [M+1].

### (2R,3S,4R,5S)-5-(4-Aminopyrrolo[2,1-f][1,2,4]triazin-7-yl)-3,4-dihydroxy-2-(hydroxymethyl)tetrahydrofuran-2-carbonitrile (2)

To a solution of **18** (81.0 mg, 0.245 mmol) in MeCN (1 mL) was added concentrated aqueous hydrochloric acid solution (12N, 1.0 mL) at rt. After 16 h, thre reaction mixture was directly purified by purified preparatory HPLC (Phenominex Synergi 4u Hydro-RR 80Å 150 x 30 mm column, 0-100% acetonitrile in water gradient) to afford the title compound (82 mg, 99%) as the HCl salt. ^1^H NMR (400 MHz, CD_3_OD) δ 8.08 (s, 1H), 7.45 (d, *J* = 4.8 Hz, 1H), 7.05 (d, *J* = 4.8 Hz, 1H), 5.56 (d, *J* = 5.5 Hz, 1H), 4.48 (t, *J* = 5.5 Hz, 1H), 4.39 (d, *J* = 5.5 Hz, 1H), 3.90 (d, *J* = 12.0 Hz, 1H), 3.83 (d, *J* = 12.0 Hz, 1H). LC/MS m/z = 292.16 [M+1].

### *tert-*Butyl (7-((3a*S*,4*S*,6*R*,6a*S*)-6-(((*tert-*butyldimethylsilyl)oxy)methyl)-6-(chloromethyl)-2,2-dimethyltetrahydrofuro[3,4-*d*][1,3]dioxol-4-yl)pyrrolo[2,1-*f*][1,2,4]triazin-4-yl)carbamate

Compound **15** (200 mg, 0.36 mmol) was dissolved in anhydrous pyridine (5 mL). Trifluoromethanesulfonyl chloride (58 µL, 0.54 mmol) was added in one portion and the reaction mixture was stirred for 60 min at rt. Additional trifluoromethanesulfonyl chloride (100 µL) was then added. After 30 min, additional trifluoromethanesulfonyl chloride (100 µL) was added. After an additional 30 min, the reaction mixture was concentrated under reduced pressure. The crude residue was dissolved in anhydrous DMF (5 mL) and lithium chloride (308 mg, 7.26 mmol) was then added in one portion. The resulting mixture was stirred at rt for 16 h. The reaction mixture was diluted with ethyl acetate (50 mL) and was washed with saturated aqueous sodium chloride solution (3 x 20 mL). The organic layer was dried over anhydrous sodium sulfate and was concentrated under reduced pressure. The crude residue was purified with silica gel chromatography (0-20% ethyl acetate in hexanes) to afford the title compound (72 mg, 42%). LCMS *m/z* = 569.0 [M+1].

### ((3a*S*,4*R*,6*S*,6a*S*)-6-(4-Aminopyrrolo[2,1-*f*][1,2,4]triazin-7-yl)-4-(chloromethyl)-2,2-dimethyltetrahydrofuro[3,4-*d*][1,3]dioxol-4-yl)methanol (19)

*tert-*Butyl (7-((3a*S*,4*S*,6*R*,6a*S*)-6-(((*tert-*butyldimethylsilyl)oxy)methyl)-6-(chloromethyl)-2,2-dimethyltetrahydrofuro[3,4-*d*][1,3]dioxol-4-yl)pyrrolo[2,1-*f*][1,2,4]triazin-4-yl)carbamate (72 mg 0.15 mmol) was dissolved in dioxane (8 mL) and water (2 mL) was added. The reaction mixture was heated to 120 °C for 2 h at which point the reaction was allowed to cool to rt and was concentrated under reduced pressure. The residue was dissolved in THF (5 mL) and TBAF (97 mg, 0.307 mmol) was added. After 1 h, the resulting mixture was diluted with ethyla acetate (20 mL) was washed with saturated aqueous sodium chloride solution (3 x 20 mL). The organic layer was dried over anhydrous sodium sulfate and was concentrated under reduced pressure. The crude residue was purified with silica gel chromatography (0-20% ethyl acetate in hexanes) to afford the title compound (47 mg, 87%). ^1^H NMR (400 MHz, CDCl_3_) δ 7.87 (s, 1H), 6.65 (d, J = 4.5 Hz, 1H), 6.54 (d, J = 4.5 Hz, 1H), 6.12 (bs, 2H), 5.33 (dd, J = 6.9, 5.9 Hz, 1H), 5.14 (d, J = 6.9 Hz, 1H), 5.05 (d, J = 5.9 Hz, 1H), 3.98 – 3.71 (m, 4H), 1.63 (s, 3H), 1.36 (s, 3H). MS *m/z* = 355.3 [M+1].

### (2R,3S,4R,5S)-5-(4-Aminopyrrolo[2,1-f][1,2,4]triazin-7-yl)-2-(chloromethyl)-2-(hydroxymethyl)tetrahydrofuran-3,4-diol (22)

To a solution of **19** (26 mg, 0.07 mmol) in MeCN (2 mL) was added a mixture of TFA and water (1:1, 5 mL). After 16 h, the reaction mixture was then concentrated under reduced pressure. The crude residue was dissolved in aqueous sodium bicarbonate solution and acetonitrile and was purified with prep HPLC to afford the title compound (20 mg, 86%). ^1^H NMR (400 MHz, D_2_O) δ 7.61 (s, 1H), 6.71 – 6.66 (m, 3H), 5.19 (d, *J* = 9.1 Hz, 1H), 4.72 – 4.65 (m, 1H), 4.28 (d, *J* = 5.2 Hz, 1H), 3.82 – 3.75 (m, 2H), 3.69 (d, *J* = 12.2 Hz, 1H), 3.61 (d, *J* = 12.1 Hz, 1H). MS *m/z* = 315.3 [M+1].

### *N*-(7-((2*S*,3*S*,4*R*,5*R*)-3,4-Bis(benzyloxy)-5-((benzyloxy)methyl)tetrahydrofuran-2-yl)pyrrolo[2,1-*f*][1,2,4]triazin-4-yl)benzamide

To a solution of compound **11** (3.94 g, 7.34 mmol) in pyridine (36.7 mL) was added benzoyl chloride (1.69 ml, 14.68 mmol) slowly at rt under an argon atmosphere. After 1 h, additional benzoyl chloride (1.69 ml, 14.68 mmol) was added slowly. After 19 h, water (20 mL) was added slowly. Ammonium hydroxide (∼10 mL) was then added slowly until the reaction mixture was basic at pH=10. After 1 h, water (150 mL) was added dropwise via addition funnel and white solids slowly began to precipitate from the reaction mixture over the course of the addition. The resulting mixture was stirred for 24 h and the white solids were collected by vacuum filtration and were dried azeotropically from toluene to afford the title compound (4.8 g, 99%). ^1^H NMR (400 MHz, CDCl_3_) δ 8.23 (br s, 1H), 7.62 (t, *J* = 7.8 Hz, 1H), 7.53 (t, *J* = 7.6 Hz, 2H), 7.38 – 7.21 (m, 18H), 7.17 (d, *J* = 7.6 Hz, 1H), 5.69 (d, *J* = 4.1 Hz, 1H), 4.71 (s, 2H), 4.63 – 4.44 (m, 4H), 4.43 – 4.39 (m, 1H), 4.22 (t, *J* = 4.5 Hz, 1H), 4.15 – 4.10 (m, 1H), 3.79 (dd, *J* = 10.8, 3.2 Hz, 1H), 3.65 (dd, *J* = 10.7, 3.7 Hz, 1H). LC/MS *m/z* = 641.18 [M+1].

### *N*-(7-((2*S*,3*R*,4*S*,5*R*)-3,4-Dihydroxy-5-(hydroxymethyl)tetrahydrofuran-2-yl)pyrrolo[1,2-*f*][1,2,4]triazin-4-yl)benzamide (23)

Ethanol (68.5 mL) and formic acid (51.7 mL, 1.37 mol) were added sequentially to a mixture of *N*-(7-((2*S*,3*S*,4*R*,5*R*)-3,4-bis(benzyloxy)-5-((benzyloxy)methyl)tetrahydrofuran-2-yl)pyrrolo[2,1-*f*][1,2,4]triazin-4-yl)benzamide (4.39 g, 6.85 mmol) and palladium on carbon (10% by wt, 2.2 g) at RT under an argon atmosphere. After 3 d, the reaction mixture was filtered through a pad of celite, and the filtrate was concentrated under reduced pressure. The crude residue was azeotroped with toluene (3 x 20 mL) to afford the title compound (2.4 g 95%), which was used directly in the next step without further purification. ^1^H NMR (400 MHz, CD_3_OD) δ 8.15 (s, 1H), 7.67 – 7.40 (m, 5H), 7.23 (d, *J* = 4.7 Hz, 1H), 7.00 (d, *J* = 4.7 Hz, 1H), 5.40 (d, *J* = 6.0 Hz, 1H), 4.44 (t, *J* = 5.7 Hz, 1H), 4.17 (t, *J* = 5.1 Hz, 1H), 4.03 (q, *J* = 4.3 Hz, 1H), 3.81 (dd, *J* = 12.1, 3.5 Hz, 1H), 3.71 (dd, *J* = 12.0, 4.5 Hz, 1H). LC/MS *m/z* = 371.15 [M+1].

### *N*-(7-((2*S*,3*R*,4*S*,5*R*)-5-((Bis(4-methoxyphenyl)(phenyl)methoxy)methyl)-3,4-dihydroxytetrahydrofuran-2-yl)pyrrolo[2,1-*f*][1,2,4]triazin-4-yl)benzamide

4,4’-Dimethoxytrityl chloride (2.23 g, 6.59 mmol) was added as a solid in one portion to a solution of **23** (2.44 g, 6.59 mmol) in pyridine (32.5 mL) at rt. After 5.5 h, the reaction mixture was diluted with ethyl acetate (300 mL) and the resulting mixture was washed with brine (3 × 200 mL). The organic layer was concentrated under reduced pressure, and the crude residue was purified via silica gel chromatography (0–100% ethyl acetate/hexanes) to afford the title compound (4.09 g, 92%). LC/MS *m/z* = 673.22 [M+1].

### *N*-(7-((2*S*,3*S*,4*R*,5*R*)-5-((Bis(4-methoxyphenyl)(phenyl)methoxy)methyl)-3,4-bis((*tert*-butyldimethylsilyl)oxy)tetrahydrofuran-2-yl)pyrrolo[2,1-*f*][1,2,4]triazin-4-yl)benzamide

*tert-*Butyldimethylsilyl chloride (2.47 g, 16.4 mmol) was added to a solution of *N*-(7-((2*S*,3*R*,4*S*,5*R*)-5-((bis(4-methoxyphenyl)(phenyl)methoxy)methyl)-3,4-dihydroxytetrahydrofuran-2-yl)pyrrolo[2,1-*f*][1,2,4]triazin-4-yl)benzamide (1.84 g, 2.74 mmol) and imidazole (2.23 g, 32.8 mmol) in *N*,*N*-dimethylformamide (28.2 mL) at rt. After 17 h, saturated aqueous sodium bicarbonate solution (500 mL) was added slowly to the reaction mixture. The resulting mixture was extracted with ethyl acetate (500 mL), and the organic layer was washed with brine (2 × 400 mL), was dried over anhydrous sodium sulfate, and was concentrated under reduced pressure. The crude residue was purified via silica gel chromatography (0–100% ethyl acetate/hexanes) to afford the title compound (2.41 g 98%). LC/MS *m/z* = 901.37 [M+1].

### *N*-(7-((2*S*,3*S*,4*R*,5*R*)-3,4-Bis((*tert*-butyldimethylsilyl)oxy)-5-(hydroxymethyl)tetrahydrofuran-2-yl)pyrrolo[2,1-*f*][1,2,4]triazin-4-yl)benzamide (24)

A solution of *p-*toluenesulfonic acid monohydrate (509 mg, 2.67 mmol) in methanol (3.7 mL) was slowly added to a solution of *N*-(7-((2*S*,3*S*,4*R*,5*R*)-5-((bis(4-methoxyphenyl)(phenyl)methoxy)methyl)-3,4-bis((*tert*-butyldimethylsilyl)oxy)tetrahydrofuran-2-yl)pyrrolo[2,1-*f*][1,2,4]triazin-4-yl)benzamide (2.41 g, 2.67 mmol) in DCM (22.3 mL) at 0 °C. After 1.5 h, the reaction mixture was diluted with saturated aqueous bicarbonate solution (100 mL), and the resulting mixture was extracted with DCM (2 × 100 mL). The combined organic extracts were dried over anhydrous sodium sulfate and were concentrated under reduced pressure. The crude residue was purified via silica gel chromatography (0–100% ethyl acetate/hexanes) to afford the title compound (1.48 g, 92%). ^1^H NMR (400 MHz, CDCl_3_) δ 8.72 (s, 1H), 8.16 (t, *J* = 7.1 Hz, 2H), 8.07 (t, *J* = 7.7 Hz, 3H), 7.49 – 7.43 (m, 1H), 5.75 (d, *J* = 8.2 Hz, 1H), 5.28 (dd, *J* = 8.1, 4.7 Hz, 1H), 4.81 (d, *J* = 5.0 Hz, 1H), 4.70 – 4.63 (m, 1H), 4.44 (d, *J* = 12.3 Hz, 1H), 4.24 (d, *J* = 12.4 Hz, 1H), 1.48 (s, 9H), 1.30 (s, 9H), 0.65 (s, 3H), 0.64 (s, 3H), 0.41 (s, 3H), 0.00 (s, 3H). LC/MS *m/z* = 599.19 [M+1].

### *N*-(7-((2*S*,3*S*,4*R*,5*S*)-3,4-Bis((*tert*-butyldimethylsilyl)oxy)-5-(iodomethyl)tetrahydrofuran-2-yl)pyrrolo[2,1-*f*][1,2,4]triazin-4-yl)benzamide

Compund **24** (0.77 g, 1.29 mmol) was added to a solution of methyltriphenoxyphosphonium iodide (0.64 g, 1.41 mmol) in DMF (6.4 mL) at rt. After 3 h, an additional portion of methyltriphenoxyphosphonium iodide (0.64 g, 1.41 mmol) was added. After 1 h, the reaction mixture was diluted with ethyl acetate (200 mL) and was washed with brine (3 × 100 mL). The organic layer was dried over anhydrous sodium sulfate and was concentrated under reduced pressure. The crude residue was purified via silica gel chromatography (0–100% ethyl acetate/hexanes) to afford the title compound (0.67 g, 73%). ^1^H NMR (400 MHz, CDCl_3_) δ 8.21 (br s, 1H), 7.61 (br t, *J* = 7.2 Hz, 1H), 7.53 (br t, *J* = 7.5 Hz, 3H), 7.05 (br s, 1H), 5.44 (d, *J* = 4.5 Hz, 1H), 4.52 (t, *J* = 4.3 Hz, 1H), 4.08 – 3.99 (m, 2H), 3.55 (dd, *J* = 10.7, 5.2 Hz, 1H), 3.38 (dd, *J* = 10.7, 5.0 Hz, 1H), 0.93 (s, 9H), 0.85 (s, 9H), 0.16 (s, 3H), 0.11 (s, 3H), −0.01 (s, 3H), −0.11 (s, 1H). LC/MS *m/z* = 709.16 [M+1].

### *N*-(7-((2*S*,3*S*,4*S*)-3,4-Bis((*tert*-butyldimethylsilyl)oxy)-5-methylenetetrahydrofuran-2-yl)pyrrolo[2,1-*f*][1,2,4]triazin-4-yl)benzamide (25)

Potassium *t*-butoxide (700 mg, 6.24 mmol) was added to a solution of *N*-(7-((2*S*,3*S*,4*R*,5*S*)-3,4-bis((*tert*-butyldimethylsilyl)oxy)-5-(iodomethyl)tetrahydrofuran-2-yl)pyrrolo[2,1-*f*][1,2,4]triazin-4-yl)benzamide (1.77 g, 2.5 mmol) in pyridine (25 mL) at rt. After 2 h, the reaction mixture was diluted with saturated aqueous sodium bicarbonate solution (25 mL) and brine (200 mL). The resulting mixture was extracted with ethyl acetate (300 mL). The organic layer was then washed with brine (200 mL), was dried over anhydrous sodium sulfate, and was concentrated under reduced pressure. The crude residue was purified via silica gel chromatography (0–100% ethyl acetate/hexanes) to afford the title compound (1.43 mg, 98%). LC/MS *m/z* = 581.37 [M+1].

### (2R,3S,4R,5S)-5-(4-Aminopyrrolo[2,1-f][1,2,4]triazin-7-yl)-2-azido-2-(hydroxymethyl)tetrahydrofuran-3,4-diol (26)

DMDO (0.07M solution in acetone, 13.8 mL, 0.964 mmol) was added to a solution of intermediate **25** (560 mg, 0.964 mmol) in acetone (4.82 mL) at 0 °C. After 10 min, the reaction mixture was concentrated under reduced pressure was dried azeotropically with toluene (2 × 1 mL) to afford the oxidized intermediate that was used immediately in the next step without further purification. To a solution the oxidized intermediate and azidotrimethysilane (242 µL, 1.84 mmol) in DCM (1.5 mL) was added indium (III) bromide (130 mg, 0.369 mmol) at rt under an argon atmosphere. After 1 h, the reaction mixture was quenched with saturated aqueous sodium bicarbonate solution (1 mL). The resulting mixture was partitioned between DCM (20 mL) and saturated aqueous sodium bicarbonate solution (20 mL). The phases were split and the aqueous layer was extracted with DCM (20 mL). The combined organic layers were dried over anhydrous sodium sulfate and were concentrated under reduced pressure. The residue was dissolved in DMF (5 mL) and cesium fluoride (256 mg, 1.68 mmol) was added at rt. After 25 h, the reaction mixture was diluted with brine (100 mL), and the resulting mixture was extracted with ethyl acetate (3 × 100 mL). The combined organic layers were dried over anhydrous sodium sulfate and were concentrated under reduced pressure. The residue was dissolved in methanol (1 mL) and concentrated ammonium hydroxide (1 mL) was added at rt. After 2 d, the reaction mixture was concentrated under reduced pressure and was directly purified by preparatory HPLC (Phenominex Synergi 4u Hydro-RR 80Å 150 x 30 mm column, 5-100% acetonitrile in water gradient). The fractions containing the desired were combined and were concentrated under reduced pressure. The residue was repurified via silica gel chromatography (0– 20% methanol in DCM) to afford the title compound (15.2 29%, over 4-steps). ^1^H NMR (400 MHz, CD_3_OD) δ 7.79 (s, 1H), 6.86 (d, *J* = 4.5 Hz, 1H), 6.77 (d, *J* = 4.5 Hz, 1H), 5.51 (d, *J* = 6.0 Hz, 1H), 4.63 (t, *J* = 5.8 Hz, 1H), 4.37 (d, *J* = 5.7 Hz, 1H), 3.69 (d, *J* = 12.0 Hz, 1H), 3.59 (d, *J* = 12.0 Hz, 1H). LC/MS *m/z* = 308.08 [M+1].

### Methyl ((*S*)-(((2*R*,3*S*,4*R*,5*S*)-5-(4-aminopyrrolo[2,1-*f*][1,2,4]triazin-7-yl)-2-cyano-3,4-dihydroxytetrahydrofuran-2-yl)methoxy)(phenoxy)phosphoryl)-L-alaninate (28a)

*N*,*N*-Diisopropylethylamine (0.12 mL, 0.68 mmol) and magnesium chloride (38.8 mg, 0.41 mmol) were added to **18** (100 mg, 0.30 mmol) and **27aj** (141 mg, 0.33 mmol) in tetrahydrofuran (3 mL) at rt. The mixture was heated to 50 °C. After 1 h, the reaction mixture was allowed to cool to rt, diluted with ethyl acetate (25 mL) and the resulting mixture was washed with water (5 × 10 mL) and brine (10 mL). The organic layer was dried over anhydrous sodium sulfate and concentrated under reduced pressure. Concentrated aqueous hydrochloric acid solution (0.2 mL) was added dropwise to the crude residue in acetonitrile (3 mL) at 0 °C. The mixture was warmed to rt. After 3 h, the reaction mixture was diluted with ethyl acetate (25 mL) and the resulting mixture was washed with saturated aqueous sodium carbonate solution (20 mL) and brine (20 mL). The organic layer was dried over anhydrous sodium sulfate and concentrated under reduced pressure. The crude residue was subjected to silica gel chromatography (0-20% methanol in DCM) to afford the title compound (150 mg, 93%). ^1^H NMR (400 MHz, CD_3_OD) δ 7.80 (s, 1H), 7.38 – 7.28 (m, 2H), 7.26 – 7.13 (m, 3H), 6.84 (d, *J* = 4.5 Hz, 1H), 6.73 (d, *J* = 4.5 Hz, 1H), 5.49 (d, *J* = 5.1 Hz, 1H), 4.63 (dd, *J* = 5.6, 5.1 Hz, 1H), 4.47 (d, *J* = 5.6 Hz, 1H), 4.40 (dd, *J* = 10.9, 6.3 Hz, 1H), 4.33 (dd, *J* = 10.9, 5.6 Hz, 1H), 3.95 – 3.85 (m, 1H), 3.60 (s, 3H), 1.26 (dd, *J* = 7.1, 1.1 Hz, 3H). ^31^P NMR (162 MHz, CD_3_OD) δ 3.24. LCMS: MS *m/z* = 533.15 [M+1].

### Ethyl ((*S*)-(((2*R*,3*S*,4*R*,5*S*)-5-(4-aminopyrrolo[2,1-*f*][1,2,4]triazin-7-yl)-2-cyano-3,4-dihydroxytetrahydrofuran-2-yl)methoxy)(phenoxy)phosphoryl)-L-alaninate (28b)

To **18** (150 mg, 0.45 mmol), **27bj** (298 mg, 0.68 mmol), and MgCl_2_ (65 mg, 0.68 mmol) in THF (6 mL) was added *N,N*-diisopropylethylamine (0.20 mL, 1.13 mmol) dropwise. The resulting mixture was stirred at 50 °C for 2 h, cooled to rt, diluted with ethyl acetate (150 mL), washed with brine (2 × 50 mL), dried, and concentrated under reduced pressure. The residue was dissolved in acetonitrile (6 mL) and concentrated aqueous hydrochloric acid solution (0.3 mL) added at 0 °C. The resulting mixture was stirred for 1 h allowed to warm to rt. After 1 h, saturated aqueous sodium bicarbonate solution (2 mL) was added and the mixture was purified by HPLC (Phenomenex Gemini-NX 10µ C18 110°A 250 x 30 mm column, 5-70% acetonitrile in water gradient) to afford the title compound (160 mg, 65%). ^1^H NMR (400 MHz, CD_3_OD) δ 7.80 (s, 1H), 7.31 (d, *J* = 7.7 Hz, 2H), 7.25 – 7.14 (m, 3H), 6.84 (d, *J* = 4.5 Hz, 1H), 6.73 (d, *J* = 4.6 Hz, 1H), 5.49 (d, *J* = 5.1 Hz, 1H), 4.62 (t, *J* = 5.3 Hz, 1H), 4.46 (d, *J* = 5.6 Hz, 1H), 4.40 (dd, *J* = 10.9, 6.2 Hz, 1H), 4.33 (dd, *J* = 10.9, 5.4 Hz, 1H), 4.11 – 3.98 (m, 2H), 3.87 (dd, *J* = 9.9, 7.1 Hz, 1H), 1.25 (dd, *J* = 7.1, 1.0 Hz, 3H), 1.16 (t, *J* = 7.1 Hz, 3H). ^31^P NMR (162 MHz, CD_3_OD) δ 3.26. LC/MS *m/z* = 547.12 [M+1].

### Propyl ((*S*)-(((2*R*,3*S*,4*R*,5*S*)-5-(4-aminopyrrolo[2,1-*f*][1,2,4]triazin-7-yl)-2-cyano-3,4-dihydroxytetrahydrofuran-2-yl)methoxy)(phenoxy)phosphoryl)-L-alaninate (28c)

*N*,*N*-Diisopropylethylamine (0.33 mL, 1.9 mmol) and magnesium chloride (108 mg, 1.13 mmol) were added to a mixture of **18** (250.0 mg, 0.76 mmol) and **27ck-mix** (462 mg, 1.13 mmol) in tetrahydrofuran (7.5 mL) at rt. The mixture was heated to 55 °C. After 2 h, the reaction mixture was allowed to cool to rt, diluted with ethyl acetate (30 mL) and the resulting mixture was washed with water (5 × 20 mL) and brine (20 mL). The organic layer was dried over anhydrous sodium sulfate and concentrated under reduced pressure. Concentrated aqueous hydrochloric acid solution (0.53 mL) was added dropwise to the crude residue in acetonitrile (7.5 mL) at 0 °C. The mixture was warmed to rt. After 2 h, the reaction mixture was diluted with ethyl acetate (100 mL) and the resulting mixture was washed with saturated aqueous sodium carbonate solution (75 mL) and brine (75 mL). The organic layer was dried over anhydrous sodium sulfate and concentrated under reduced pressure. The crude residue was subjected to silica gel chromatography (0-20% methanol in DCM) to afford **28c-mix** (305 mg, 65%, ∼2:1 diastereomeric mixture). ^1^H NMR (400 MHz, CD_3_OD) δ 7.80 (d, *J* = 7.2 Hz, 1H), 7.37 – 7.27 (m, 2H), 7.26 – 7.13 (m, 3H), 6.85 (dd, *J* = 4.5, 2.9 Hz, 1H), 6.74 (dd, *J* = 4.6, 2.1 Hz, 1H), 5.50 (t, *J* = 5.3 Hz, 1H), 4.63 (q, *J* = 5.3 Hz, 1H), 4.54 – 4.31 (m, 3H), 4.07 – 3.82 (m, 3H), 1.68 – 1.49 (m, 2H), 1.31 – 1.26 (m, 3H), 0.90 (dt, *J* = 9.9, 7.4 Hz, 3H). ^31^P NMR (162 MHz, CD_3_OD) δ 3.27. LC/MS *m/z* = 561.20 [M+1]. Resolution of the *S*p and *R*p diastereomers was conducted via chiral SFC (Chiralpak AD-H, 5um, 21 x 250 mm, Isopropyl alcohol 30%) from 40 mg of the mixture. First eluting *R*p diastereomer **28c-*R*p** (14.5 mg): ^1^H NMR (400 MHz, CD_3_OD) δ 7.79 (s, 1H), 7.33 – 7.26 (m, 2H), 7.20 – 7.12 (m, 3H), 6.85 (d, *J* = 4.5 Hz, 1H), 6.73 (d, *J* = 4.5 Hz, 1H), 5.51 (d, *J* = 5.0 Hz, 1H), 4.68 – 4.60 (m, 1H), 4.53 (d, *J* = 5.6 Hz, 1H), 4.48 (dd, *J* = 10.9, 6.0 Hz, 1H), 4.36 (dd, *J* = 10.9, 5.1 Hz, 1H), 4.06 – 3.95 (m, 2H), 3.88 (dq, *J* = 9.4, 7.1 Hz, 1H), 1.62 (h, *J* = 7.3 Hz, 2H), 1.26 (dd, *J* = 7.1, 1.3 Hz, 3H), 0.91 (t, *J* = 7.5 Hz, 3H). ^31^P NMR (162 MHz, CD_3_OD) δ 3.26. LCMS *m/z* = 561.21 [M+1]. Second eluting *S*p diastereomer **28c** (22.4 mg): ^1^H NMR (400 MHz, CD_3_OD) δ 7.80 (s, 1H), 7.37 – 7.29 (m, 2H), 7.26 – 7.14 (m, 3H), 6.84 (d, *J* = 4.5 Hz, 1H), 6.74 (d, *J* = 4.5 Hz, 1H), 5.49 (d, *J* = 5.0 Hz, 1H), 4.62 (dd, *J* = 5.6, 5.1 Hz, 1H), 4.47 (d, *J* = 5.6 Hz, 1H), 4.41 (dd, *J* = 10.9, 6.3 Hz, 1H), 4.33 (dd, *J* = 10.9, 5.5 Hz, 1H), 4.02 – 3.85 (m, 3H), 1.57 (dtd, *J* = 14.0, 7.4, 6.6 Hz, 2H), 1.27 (dd, *J* = 7.2, 1.1 Hz, 3H), 0.87 (t, *J* = 7.5 Hz, 3H). ^31^P NMR (162 MHz, CD_3_OD) δ 3.27. LCMS *m/z* = 561.26 [M+1].

### Isopropyl ((*S*)-(((2*R*,3*S*,4*R*,5*S*)-5-(4-aminopyrrolo[2,1-*f*][1,2,4]triazin-7-yl)-2-cyano-3,4-dihydroxytetrahydrofuran-2-yl)methoxy)(phenoxy)phosphoryl)-L-alaninate (28d)

Compound **2** (50 mg, 0.172 mmol) and **27dk** (previously synthesized,^54^ 84 mg, 0.206 mmol) were taken up into anhydrous *N*,*N*-dimethylformamide (2 mL). Magnesium chloride (36 mg, 0.378 mmol) was added in one portion. The reaction mixture was heated at 50 °C. *N*,*N*-Diisopropylethylamine (75 uL, 0.43 mmol) was added, and the reaction was stirred for 4.5 h at 50 °C. The reaction mixture was cooled to rt, diluted with ethyl acetate (30 mL), and washed with 5% aqueous citric acid solution (10 mL) and then brine (10 mL). The organic layer was dried over anhydrous sodium sulfate and concentrated under reduced pressure. The crude was purified via silica gel chromatography (0-5% methanol in DCM) to afford the title compound (32 mg, 33%). ^1^H NMR (400 MHz, CD_3_OD) δ 7.79 (s, 1H), 7.36 – 7.25 (m, 2H), 7.25 – 7.12 (m, 3H), 6.84 (d, *J* = 4.5 Hz, 1H), 6.73 (d, *J* = 4.5 Hz, 1H), 5.49 (d, *J* = 5.1 Hz, 1H), 4.91 – 4.84 (m, 1H), 4.62 (dd, *J* = 5.6, 5.0 Hz, 1H), 4.47 (d, *J* = 5.6 Hz, 1H), 4.45 – 4.30 (m, 2H), 3.85 (dq, *J* = 10.0, 7.1 Hz, 1H), 1.25 (d, *J* = 7.2 Hz, 3H), 1.15 (t, *J* = 6.4 Hz, 6H). ^31^P NMR (162 MHz, CD_3_OD) δ 3.31. MS *m/z* = 561.0 [M+1], 559.0 [M-1].

### 2-Ethylbutyl ((*S*)-(((2*R*,3*S*,4*R*,5*S*)-5-(4-aminopyrrolo[2,1-*f*][1,2,4]triazin-7-yl)-2-cyano-3,4-dihydroxytetrahydrofuran-2-yl)methoxy)(phenoxy)phosphoryl)-L-alaninate (28e)

To a mixture of **18** (700 mg, 2.11 mmol), **27ek** (previously synthesized,^24^ 998 mg, 2.22 mmol), and magnesium chloride (302 mg, 3.17 mmol) was added tetrahydrofuran (8.5 mL) at rt followed by the addition of *N,N*-Diisopropylethylamine (0.92 mL, 5.282 mmol). The resulting mixture was stirred at 50 °C for 3 h. The reaction mixture was then concentrated under reduced pressure and the residue obtained was diluted with saturated sodium chloride solution and DCM. The layers were split and the organic layer was dried over anhydrous sodium sulfate, filtered, and concentrated under reduced pressure. The crude residue was purified via silica gel chromatography (0-14% methanol in DCM). The fractions containing desired product were combined and concentration under reduced pressure. The residue was dissolved in an acetonitrile (10 mL) and was cooled in an ice bath followed by the dropwise addition of concentrated hydrochloric acid (4 mL, 48 mmol). The reaction mixture was then stirred at rt for 1 h, was cooled in an ice bath and was diluted with water. The resulting mixture was neutralized the solution with 3N sodium hydroxide to pH = 7 and extracted with DCM. Organic layer was dried over anhydrous sodium sulfate, filtered, and concentrated under reduced pressure. The residue was purified by silica gel chromatography (0-20% methanol in DCM) to afford the title compound (0.85 g, 67%). ^1^H NMR (400 MHz, CD_3_OD) δ 7.80 (s, 1H), 7.38 – 7.29 (m, 2H), 7.27 – 7.13 (m, 3H), 6.84 (d, *J* = 4.5 Hz, 1H), 6.74 (d, *J* = 4.5 Hz, 1H), 5.49 (d, *J* = 5.0 Hz, 1H), 4.61 (t, *J* = 5.3 Hz, 1H), 4.49 – 4.29 (m, 3H), 4.04 – 3.82 (m, 3H), 1.43 (dq, *J* = 12.5, 6.1 Hz, 1H), 1.37 – 1.23 (m, 7H), 0.84 (td, *J* = 7.5, 1.1 Hz, 6H). ^31^P NMR (162 MHz, CD_3_OD) δ 3.27. MS *m/z* = 603 [M+1].

### Cyclohexyl ((*S*)-(((2*R*,3*S*,4*R*,5*S*)-5-(4-aminopyrrolo[2,1-*f*][1,2,4]triazin-7-yl)-2-cyano-3,4-dihydroxytetrahydrofuran-2-yl)methoxy)(phenoxy)phosphoryl)-L-alaninate (28f) and cyclohexyl ((*R*)-(((2*R*,3*S*,4*R*,5*S*)-5-(4-aminopyrrolo[2,1-*f*][1,2,4]triazin-7-yl)-2-cyano-3,4-dihydroxytetrahydrofuran-2-yl)methoxy)(phenoxy)phosphoryl)-L-alaninate (28f-*R*p)

To a mixture of **18** (99 mg, 0.30 mmol), **27fk-mix** (201 mg, 0.45 mmol), and MgCl_2_ (43 mg, 0.45 mmol) in DMF (4 mL) was added *N,N*-diisopropylethylamine (0.13 mL, 0.75 mmol) dropwise at rt. The resulting mixture was stirred at rt for 15 h and purified by directly by preparative HPLC (Phenominex Synergi 4u Hydro-RR 80Å 150 x 30 mm column, 10-100% acetonitrile in water gradient) afford the acetonide intermediate (130 mg, 68%). The acetonide intermediate was dissolved in acetonitrile (3 mL) and concentrate hydrochloride acide solution (0.1 mL) was added. The resulting mixture was stirred at 50 °C for 2 h and was purified by preparative HPLC (Phenominex Synergi 4u Hydro-RR 80Å 150 x 30 mm column, 10-80% acetonitrile in water gradient) to give **28f-mix** (55 mg, 31%, 1:1 diastereomeric mixture). ^1^H NMR (400 MHz, CD_3_OD) δ 7.80 (s, 0.5H), 7.78 (s, 0.5H), 7.42 – 7.05 (m, 5H), 6.84 (m, 1H), 6.73 (m, 1H), 5.50 (m, 1H), 4.64 (m, 2H), 4.57 – 4.25 (m, 3H), 3.86 (m, 1H), 1.91 – 1.61 (m, 4H), 1.61 – 1.09 (m, 9H). ^31^P NMR (162 MHz, CD_3_OD) δ 3.3. LC/MS *m/z* = 601 [M+1]. Resolution of the *S*p and *R*p diastereomers was conducted via chiral preparatory HPLC (Chiralpak IA,150 x 4.6 mm, Heptane 70% Ethanol 30%) from 15 mg of the mixture. First eluting *R*p diastereomer **28f-*R*p** (6.7 mg): ^1^H NMR (400 MHz, CD_3_OD) δ 7.78 (s, 1H), 7.34 – 7.23 (m, 2H), 7.19 – 7.10 (m, 3H), 6.85 (d, *J* = 4.5 Hz, 1H), 6.73 (d, *J* = 4.5 Hz, 1H), 5.51 (d, *J* = 5.0 Hz, 1H), 4.69 (td, *J* = 8.8, 4.2 Hz, 1H), 4.62 (t, *J* = 5.3 Hz, 1H), 4.53 – 4.44 (m, 2H), 4.36 (dd, *J* = 10.9, 5.2 Hz, 1H), 3.86 (dq, *J* = 9.4, 7.1 Hz, 1H), 1.85 – 1.62 (m, 4H), 1.58 – 1.20 (m, 9H). ^31^P NMR (162 MHz, CD_3_OD) δ 3.31. LC/MS *m/z* = 601 [M+1]. Second eluting *S*p diastereomer **28f** (6.8 mg): ^1^H NMR (400 MHz, CD_3_OD) δ 7.80 (s, 1H), 7.37 – 7.27 (m, 2H), 7.26 – 7.13 (m, 3H), 6.84 (d, *J* = 4.5 Hz, 1H), 6.73 (d, *J* = 4.5 Hz, 1H), 5.49 (d, *J* = 5.0 Hz, 1H), 4.71 – 4.56 (m, 2H), 4.46 (d, *J* = 5.6 Hz, 1H), 4.45 – 4.30 (m, 2H), 3.87 (dq, *J* = 10.0, 7.1 Hz, 1H), 1.80 – 1.61 (m, 4H), 1.55 – 1.21 (m, 9H). ^31^P NMR (162 MHz, CD_3_OD) δ 3.30. LC/MS *m/z* = 601 [M+1].

### Tetrahydro-2*H*-pyran-4-yl ((*S*)-(((2*R*,3*S*,4*R*,5*S*)-5-(4-aminopyrrolo[2,1-*f*][1,2,4]triazin-7-yl)-2-cyano-3,4-dihydroxytetrahydrofuran-2-yl)methoxy)(phenoxy)phosphoryl)-L-alaninate (28g)

To a mixture of **18** (90 mg, 0.27 mmol), **27gk-mix** (184 mg, 3 mmol), and MgCl_2_ (39 mg, 0.40 mmol) in tetrahydrofuran (3 mL) was added *N,N*-diisopropylethylamine (88 mg, 0.70 mmol) drop wise. The resulting mixture was stirred at 50 °C for 2 h, and the reaction mixture was diluted with ethyl acetate (40 mL), washed with water (40 mL) and brine (40 mL), and concentrated under reduced pressure. The residue was then dissolved in acetonitrile (40 mL) and concentrated hydrochloric acid solution was added dropwise at 0 °C. The resulting mixture was stirred at rt for 2 h, cooled to 0 °C and neutralized to pH 7 by dropwise addition of 2 N NaOH. The resulting mixture was diluted with ethyl acetate (150 mL), washed with water (50 mL) and brine (50 mL). The aqueous phase was extracted with ethyl acetate (2 × 50 mL) and the combined organic layers were dried over anhydrous sodium sulfate and concentrated under reduced pressure. The residue was purified by preparative HPLC (Phenominex Synergi 4u Hydro-RR 80Å 150 x 30 mm column, 10-40% acetonitrile in water gradient) to afford the two diastereomers. First eluting *R*p diastereomer **28g-*R*p** (6 mg, 4%): ^1^H NMR (400 MHz, CD_3_OD) δ 7.78 (s, 1H), 7.29 (dd, *J* = 8.7, 7.0 Hz, 2H), 7.16 (ddd, *J* = 7.1, 2.1, 1.1 Hz, 3H), 6.85 (d, *J* = 4.5 Hz, 1H), 6.73 (d, *J* = 4.5 Hz, 1H), 5.50 (d, *J* = 5.0 Hz, 1H), 4.88 (dq, *J* = 9.4, 5.1, 4.7 Hz, 1H), 4.63 (t, *J* = 5.3 Hz, 1H), 4.55 – 4.44 (m, 2H), 4.36 (dd, *J* = 10.9, 5.2 Hz, 1H), 3.86 (m, 3H), 3.50 (dtd, *J* = 11.3, 5.4, 2.7 Hz, 2H), 1.94 – 1.76 (m, 2H), 1.60 (dtd, *J* = 12.9, 8.4, 3.9 Hz, 2H), 1.27 (dd, *J* = 7.1, 1.3 Hz, 3H). ^31^P NMR (162 MHz, CD_3_OD) δ 3.23. LC/MS *m/z* = 603 [M+1]. Second eluting *S*p diastereomer **28g** (24 mg 15%): ^1^H NMR (400 MHz, CD_3_OD) δ 7.80 (s, 1H), 7.33 (dd, *J* = 8.6, 7.2 Hz, 2H), 7.27 – 7.11 (m, 3H), 6.84 (d, *J* = 4.5 Hz, 1H), 6.74 (d, *J* = 4.5 Hz, 1H), 5.49 (d, *J* = 5.0 Hz, 1H), 4.80 (m, 1H), 4.61 (t, *J* = 5.3 Hz, 1H), 4.50 – 4.38 (m, 2H), 4.35 (dd, *J* = 10.9, 5.5 Hz, 1H), 3.90 (dq, *J* = 9.9, 7.1 Hz, 1H), 3.85 – 3.75 (m, 2H), 3.46 (dddd, *J* = 11.8, 8.9, 6.0, 3.2 Hz, 2H), 1.81 (tdd, *J* = 9.6, 4.6, 2.5 Hz, 2H), 1.57 (dtd, *J* = 12.7, 8.4, 3.9 Hz, 2H), 1.27 (dd, *J* = 7.1, 1.1 Hz, 3H). ^31^P NMR (162 MHz, CD_3_OD) δ 3.23. LC/MS *m/z* = 603 [M+1].

### (2*R*,3*S*,4*S*,5*S*)-5-(4-Aminopyrrolo[2,1-*f*][1,2,4]triazin-7-yl)-2-cyano-2-((((*S*)-(((*S*)-1-methoxy-1-oxopropan-2-yl)amino)(phenoxy)phosphoryl)oxy)methyl)tetrahydrofuran-3,4-diyl bis(2-methylpropanoate) (1)

**28a** (0.68 g, 1.3 mmol) was dissolved in anhydrous tetrahydrofuran (10 mL). 4-(Dimethylamino)pyridine (23 mg, 0.19 mmol), and isobutyric anhydride (445 µL, 2.7 mmol) were added. After 30 min, the reaction mixture was diluted with ethyl acetate (10 mL) and washed with saturated aqueous sodium bicarbonate solution (2 × 10 mL) and brine (10 mL). The organic layer was dried over anhydrous sodium sulfate and was concentrated under reduced pressure. The residue was purified by silica gel column chromatography (0 to 100% ethyl acetate in hexanes) to afford the title compound (0.79 g, 92%). ^1^H NMR (400 MHz, CD_3_OD) δ 7.83 (s, 1H), 7.34 – 7.27 (m, 2H), 7.22 – 7.13 (m, 3H), 6.84 (d, *J* = 4.5 Hz, 1H), 6.75 (d, *J* = 4.6 Hz, 1H), 5.92 (d, *J* = 5.8 Hz, 1H), 5.81 (dd, *J* = 5.9, 4.7 Hz, 1H), 5.68 (d, *J* = 4.7 Hz, 1H), 4.49 – 4.37 (m, 2H), 3.97 – 3.83 (m, 1H), 3.61 (s, 3H), 2.73 – 2.55 (m, 2H), 1.31 – 1.13 (m, 15H). ^31^P NMR (162 MHz, CD_3_OD) δ 3.02 (s). LC/MS *m/z* = 673.2 [M+1].

### (2R,3S,4S,5S)-5-(4-Aminopyrrolo[2,1-f][1,2,4]triazin-7-yl)-2-cyano-2-((((S)-(((S)-1-methoxy-1-oxopropan-2-yl)amino)(phenoxy)phosphoryl)oxy)methyl)tetrahydrofuran-3,4-diyl dipropionate (29am)

**28a-mix** (54 mg, 0.10 mmol) was dissolved in 2 mL of anhydrous tetrahydrofuran. Propionic acid (30 µL, 0.40 mmol) and *N,Ń-*diisopropylcarbodiimide (62 µL, 0.40 mmol) were added and stirred for 30 min. DMAP (12 mg, 0.10 mmol) was added and stirred for 16 h. Methanol (0.5 mL) was added and stirred for 20 min. The reaction mixture was purified directly with preparative HPLC (Phenominex Synergi 4u Hydro-RR 80Å 150 x 30 mm column, 10-40% acetonitrile in water gradient) to afford **29am-mix** (52 mg, 81%, ∼1.75:1 diastereomeric mixture). ^1^H NMR (400 MHz, CD_3_OD) δ 7.88 – 7.84 (m, 1H), 7.36 – 7.24 (m, 2H), 7.24 – 7.10 (m, 3H), 6.99 – 6.95 (m, 1H), 6.84 – 6.81 (m, 1H), 5.99 – 5.75 (m, 2H), 5.73 – 5.70 (m, 1H), 4.57 – 4.38 (m, 2H), 3.96 – 3.71 (m, 1H), 3.65 – 3.60 (m, 3H), 2.55 – 2.34 (m, 4H), 1.34 – 1.03 (m, 9H). ^31^P NMR (162 MHz, CD_3_OD) δ 3.03. LC/MS *m/z* = 645.2 [M+1]; 643.3 [M-1]. Resolution of the *S*p and *R*p diastereomers was conducted via chiral preparatory HPLC (Chiralpak IF, 150 x 4.6 mm, SFC 30% ethanol isocratic) from 46 mg of the mixture. First eluting *R*p diastereomer **29am-*R*p** (13 mg). ^1^H NMR (400 MHz, CD_3_OD) δ 7.81 (s, 1H), 7.35 – 7.25 (m, 2H), 7.21 – 7.11 (m, 3H), 6.86 (d, *J* = 4.6 Hz, 1H), 6.78 (d, *J* = 4.5 Hz, 1H), 5.99 (d, *J* = 5.8 Hz, 1H), 5.87 (dd, *J* = 5.8, 4.4 Hz, 1H), 5.70 (d, *J* = 4.4 Hz, 1H), 4.52 (dd, *J* = 11.1, 5.9 Hz, 1H), 4.42 (dd, *J* = 11.1, 4.9 Hz, 1H), 3.87 – 3.74 (m, 1H), 3.63 (s, 3H), 2.57 – 2.36 (m, 4H), 1.31 – 1.05 (m, 9H). ^31^P NMR (162 MHz, CD_3_OD) δ 3.01 (s). LC/MS *m/z* = 645.2 [M+1]. Second eluting *S*p diastereomer **29am** (26 mg). ^1^H NMR (400 MHz, CD_3_OD) δ 7.83 (s, 1H), 7.35 – 7.26 (m, 2H), 7.22 – 7.12 (m, 3H), 6.84 (d, *J* = 4.5 Hz, 1H), 6.74 (d, *J* = 4.5 Hz, 1H), 5.91 (d, *J* = 5.9 Hz, 1H), 5.82 (dd, *J* = 5.9, 4.8 Hz, 1H), 5.69 (d, *J* = 4.8 Hz, 1H), 4.49 – 4.36 (m, 2H), 3.96 – 3.84 (m, 1H), 3.61 (s, 3H), 2.53 – 2.33 (m, 4H), 1.27 (dd, *J* = 7.2, 1.1 Hz, 3H), 1.22 – 1.08 (m, 6H). ^31^P NMR (162 MHz, CD_3_OD) δ 3.02 (s). LC/MS *m/z* = 645.2 [M+1].

### (2R,3S,4S,5S)-5-(4-Aminopyrrolo[2,1-f][1,2,4]triazin-7-yl)-2-cyano-2-((((S)-(((S)-1-methoxy-1-oxopropan-2-yl)amino)(phenoxy)phosphoryl)oxy)methyl)tetrahydrofuran-3,4-diyl diacetate (29an)

*N*,*N*’-diisopropylcarbodiimide (0.307 mL, 1.97 mmol) and 4-dimethylaminopyridine (48.0 mg, 0.394 mmol) were added to a solution of **28a-mix** (210 mg, 0.394 mmol) and acetic acid (0.113 mL, 1.97 mmol) in tetrahydrofuran (2.0 mL) at rt. After 1.5 h, methanol (0.2 mL) was added and the resulting mixture was concentrated under reduced pressure. The crude residue was subjected to silica gel chromatography eluting with 0-100% ethyl acetate in hexanes to afford **29an-mix** (110 mg, 45%, ∼2:1 diastereomeric mixture). ^1^H NMR (400 MHz, CD_3_OD) δ 7.82 (s, 0.66H), 7.80 (s, 0.33H), 7.36 – 7.25 (m, 2H), 7.22 – 7.12 (m, 3H), 6.86 – 6.81 (m, 1H), 6.78 – 6.71 (m, 1H), 5.96 (d, *J =* 5.9 Hz, 0.33H), 5.89 (d, *J =* 5.9 Hz, 0.66H), 5.87 – 5.78 (m, 1H), 5.71 – 5.68 (m, 1H), 4.54 – 4.35 (m, 2H), 3.95 – 3.75 (m, 1H), 3.62 (s, 1H), 3.60 (s, 2H), 2.16 (s, 1H), 2.15 (s, 2H), 2.11 (s, 1H), 2.10 (s, 2H), 1.27 (dd, *J =* 7.2, 1.1 Hz, 2H), 1.21 (dd, *J =* 7.2, 1.3 Hz, 1H). ^31^P NMR (162 MHz, CD_3_OD) δ 3.02 (s), 3.00 (s). LC/MS *m/z* = 617.16 [M+1]. Resolution of the *S*p and *R*p diastereomers was conducted via chiral preparatory SFC (Chiralpak ADH, 30% ethanol) from 70 mg of the mixture. First eluting *R*p diastereomer **29an-*R*p** (21 mg). ^1^H NMR (400 MHz, CD_3_OD) δ 7.80 (s, 1H), 7.34 – 7.24 (m, 2H), 7.19 – 7.12 (m, 3H), 6.85 (d, *J* = 4.5 Hz, 1H), 6.77 (d, *J* = 4.5 Hz, 1H), 5.95 (d, *J* = 5.9 Hz, 1H), 5.85 (dd, *J* = 5.9, 4.6 Hz, 1H), 5.70 (d, *J* = 4.6 Hz, 1H), 4.52 (dd, *J* = 11.1, 5.9 Hz, 1H), 4.42 (dd, *J* = 11.1, 4.9 Hz, 1H), 3.80 (dq, *J* = 9.4, 7.1 Hz, 1H), 3.63 (s, 3H), 2.16 (s, 3H), 2.11 (s, 3H), 1.21 (dd, *J* = 7.2, 1.3 Hz, 3H). ^31^P NMR (162 MHz, CD_3_OD) δ 3.00 (s). LC/MS *m/z* = 617.16 [M+1]. Second eluting *S*p diastereomer **29an** (42.8 mg). ^1^H NMR (400 MHz, CD_3_OD) δ 7.83 (s, 1H), 7.34 – 7.26 (m, 2H), 7.23 – 7.11 (m, 3H), 6.83 (d, *J* = 4.5 Hz, 1H), 6.74 (d, *J* = 4.6 Hz, 1H), 5.89 (d, *J* = 5.9 Hz, 1H), 5.81 (dd, *J* = 5.9, 4.9 Hz, 1H), 5.70 (d, *J* = 4.9 Hz, 1H), 4.50 – 4.38 (m, 2H), 3.96 – 3.86 (m, 1H), 3.61 (s, 3H), 2.15 (s, 3H), 2.10 (s, 3H), 1.27 (dd, *J* = 7.1, 1.1 Hz, 4H). ^31^P NMR (162 MHz, CD_3_OD) δ 3.02 (s). LC/MS MS *m/z* = 617.16 [M+1].

### (2*R*,3*S*,4*S*,5*S*)-5-(4-Aminopyrrolo[2,1-*f*][1,2,4]triazin-7-yl)-2-cyano-2-((((*S*)-(((*S*)-1-oxo-1-propoxypropan-2-yl)amino)(phenoxy)phosphoryl)oxy)methyl)tetrahydrofuran-3,4-diyl bis(2-methylpropanoate) (29cl)

Isobutyric anhydride (30.3 uL, 0.18 mmol) was added to a solution of **28c-mix** (51.2 mg, 0.09 mmol) and 4-dimethylaminopyridine (1.8 mg, 0.1 mmol) in tetrahydrofuran (1.8 mL) at rt. After 30 min, the reaction mixture was diluted with ethyl acetate (15 mL) and the resulting mixture was washed with saturated aqueous sodium carbonate solution (10 mL) and brine (10 mL). The organic layer was dried over anhydrous sodium sulfate and concentrated under reduced pressure. The crude residue was purified by silica gel chromatography (20-100% ethyl acetate in hexanes) to afford **29cl-mix** (50 mg, 74%, ∼2:1 diastereomeric mixture) as a white solid. ^1^H NMR (400 MHz, CD_3_OD) δ 7.85 – 7.80 (m, 1H), 7.32 – 7.27 (m, 2H), 7.25 - 7.06 (m, 3H), 6.88 – 6.83 (m, 1H), 6.79 – 6.73 (m, 1H), 5.98 – 5.90 (m, 1H), 5.84 – 5.78 (m, 1H), 5.70 – 5.65 (m, 1H), 4.60 – 4.35 (m, 2H), 4.08 – 3.75 (m, 3H), 2.75 – 2.52 (m, 2H), 1.63 – 1.50 (m, 2H), 1.33 - 1.14 (m, 15H), 0.93 – 0.86 (m, 3H). ^31^P NMR (162 MHz, CD_3_OD) δ 3.06, 3.04. LC/MS *m/z* = 701.47 [M+1]. Resolution of the *S*p and *R*p diastereomers was conducted via chiral preparatory SFC (Chiralpak AD-H, 5um, 21 x 250 mm, isopropyl alcohol 30%) from 50 mg of the mixture. First eluting *R*p diastereomer **29cl-*R*p** (14.6 mg). ^1^H NMR (400 MHz, CD_3_OD) δ 7.81 (s, 1H), 7.34 – 7.26 (m, 2H), 7.20 – 7.11 (m, 3H), 6.86 (d, *J* = 4.6 Hz, 1H), 6.77 (d, *J* = 4.6 Hz, 1H), 5.97 (d, *J* = 5.9 Hz, 1H), 5.85 (dd, *J* = 5.9, 4.4 Hz, 1H), 5.68 (d, *J* = 4.4 Hz, 1H), 4.52 (dd, *J* = 11.1, 5.7 Hz, 1H), 4.42 (dd, *J* = 11.0, 4.9 Hz, 1H), 4.08 – 3.95 (m, 2H), 3.84 (dq, *J* = 9.4, 7.2 Hz, 1H), 2.72 – 2.60 (m, 2H), 1.65 – 1.57 (m, 2H), 1.26 – 1.16 (m, 15H), 0.90 (t, *J* = 7.4 Hz, 3H). ^31^P NMR (162 MHz, CD_3_OD) δ 3.07. LC/MS *m/z* = 701.48 [M+1]. Second eluting *S*p diastereomer **29cl** (21 mg). ^1^H NMR (400 MHz, CD_3_OD) δ 7.84 (s, 1H), 7.35 – 7.25 (m, 2H), 7.24 – 7.13 (m, 3H), 6.84 (d, *J* = 4.5 Hz, 1H), 6.75 (d, *J* = 4.6 Hz, 1H), 5.91 (d, *J* = 5.9 Hz, 1H), 5.80 (dd, *J* = 5.9, 4.7 Hz, 1H), 5.67 (d, *J* = 4.6 Hz, 1H), 4.50 – 4.39 (m, 2H), 4.07 – 3.82 (m, 3H), 2.73 – 2.56 (m, 2H), 1.66 – 1.52 (m, 2H), 1.29 (dd, *J* = 7.1, 1.1 Hz, 3H), 1.26 – 1.15 (m, 12H), 0.89 (t, *J* = 7.4 Hz, 3H). ^31^P NMR (162 MHz, CD_3_OD) δ 3.05. LCMS: MS *m/z* = 701.47 [M+1].

### (2R,3S,4S,5S)-5-(4-Aminopyrrolo[2,1-f][1,2,4]triazin-7-yl)-2-cyano-2-((((S)-(((S)-1-oxo-1-propoxypropan-2-yl)amino)(phenoxy)phosphoryl)oxy)methyl)tetrahydrofuran-3,4-diyl dipropionate (29cm)

Propionic anhydride (23.5 µL, 0.18 mmol) was added to a solution of **28c-mix** (51 mg, 0.090 mmol) and 4-dimethylaminopyridine (1.8 mg, 0.10 mmol) in tetrahydrofuran (1.8 mL) at rt. After 20 min, the reaction mixture was diluted with ethyl acetate (15 mL) and the resulting mixture was washed with saturated aqueous sodium carbonate solution (10 mL) and brine (10 mL). The organic layer was dried over anhydrous sodium sulfate and concentrated under reduced pressure. The crude residue was purified by silica gel chromatography (20-100% ethyl acetate in hexanes) to afford **29cm-mix** (45.0 mg, 71%, ∼2:1 diastereomeric mixture) as a white solid. ^1^H NMR (400 MHz, CD_3_OD) δ 7.84 – 7.80 (m, 1H), 7.38 – 7.26 (m, 2H), 7.25 – 7.09 (m, 3H), 6.87 – 6.82 (m, 1H), 6.79 – 6.73 (m, 1H), 6.02 – 5.87 (m, 1H), 5.88 – 5.79 (m, 1H), 5.70 – 5.67 (m, 1H), 4.57 – 4.37 (m, 2H), 4.07 – 3.76 (m, 3H), 2.55 – 2.26 (m, 4H), 1.63 – 1.57 (m, 2H), 1.31 – 1.11 (m, 9H), 0.93 – 0.86 (m, 3H). ^31^P NMR (162 MHz, CD_3_OD) δ 3.05. LCMS: MS *m/z* = 673.40 [M+1]. Resolution of the *S*p and *R*p diastereomers was conducted via chiral preparatory SFC (Chiralpak AD-H, 5um, 21 x 250 mm, isopropyl alcohol 30%) from 45 mg of the mixture. First eluting *R*p diastereomer **29cm-*R*p** (12 mg). ^1^H NMR (400 MHz, CD_3_OD) δ 7.81 (s, 1H), 7.35 – 7.26 (m, 2H), 7.20 – 7.12 (m, 3H), 6.86 (d, *J* = 4.5 Hz, 1H), 6.78 (d, *J* = 4.5 Hz, 1H), 5.97 (d, *J* = 5.9 Hz, 1H), 5.86 (dd, *J* = 5.9, 4.5 Hz, 1H), 5.70 (d, *J* = 4.5 Hz, 1H), 4.52 (dd, *J* = 11.1, 5.8 Hz, 1H), 4.43 (dd, *J* = 11.1, 5.0 Hz, 1H), 4.08 – 3.94 (m, 2H), 3.84 (dq, *J* = 9.4, 7.2 Hz, 1H), 2.55 – 2.35 (m, 4H), 1.61 (dtd, *J* = 14.1, 7.4, 6.6 Hz, 2H), 1.24 (dd, *J* = 7.1, 1.3 Hz, 3H), 1.16 (dt, *J* = 13.4, 7.5 Hz, 6H), 0.90 (t, *J* = 7.4 Hz, 3H). ^31^P NMR (162 MHz, CD_3_OD) δ 3.05. LC/MS *m/z* = 673.41 [M+1]. Second eluting *S*p diastereomer **29cm** (21 mg). ^1^H NMR (400 MHz, CD_3_OD) δ 7.83 (s, 1H), 7.35 – 7.27 (m, 2H), 7.24 – 7.14 (m, 3H), 6.84 (d, *J* = 4.6 Hz, 1H), 6.75 (d, *J* = 4.5 Hz, 1H), 5.91 (d, *J* = 5.9 Hz, 1H), 5.82 (dd, *J* = 5.9, 4.7 Hz, 1H), 5.69 (d, *J* = 4.7 Hz, 1H), 4.50 – 4.38 (m, 2H), 4.06 – 3.85 (m, 3H), 2.54 – 2.37 (m, 4H), 1.67 – 1.51 (m, 2H), 1.29 (dd, *J* = 7.1, 1.1 Hz, 3H), 1.23 – 1.10 (m, 6H), 0.89 (t, *J* = 7.4 Hz, 3H). ^31^P NMR (162 MHz, CD_3_OD) δ 3.05. LC/MS *m/z* = 673.41 [M+1].

### (2*R*,3*S*,4*S*,5*S*)-5-(4-Aminopyrrolo[2,1-*f*][1,2,4]triazin-7-yl)-2-cyano-2-((((*S*)-(((*S*)-1-(cyclohexyloxy)-1-oxopropan-2-yl)amino)(phenoxy)phosphoryl)oxy)methyl)tetrahydrofuran-3,4-diyl bis(2-methylpropanoate) (29fl)

**28f** (10 mg, 0.017 mmol) was dissolved in anhydrous DMF (1 mL). Isobutyric acid (6.2 µL, 0.070 mmol) and *N*,*N*’-diisopropylcarbodiimide (10 µL, 0.070 mmol) were added to the reaction and stirred for 20 min. 4-Dimethylaminopyridine (2 mg, 0.017 mmol) was added, and the reaction was stirred for 16 h. The reaction mixture was diluted with ethyl acetate (10 mL) and the resulting mixture was washed with saturated aqueous sodium carbonate solution (10 mL) and brine (10 mL). The organic layer was dried over anhydrous sodium sulfate and concentrated under reduced pressure. The crude residue was purified by silica gel chromatography (0-100% ethyl acetate in hexanes) to afford the title compound (6.8 mg, 55%). ^1^H NMR (400 MHz, CD_3_OD) δ 7.82 (s, 1H), 7.34 – 7.26 (m, 2H), 7.22 – 7.13 (m, 3H), 6.83 (d, *J* = 4.6 Hz, 1H), 6.73 (d, *J* = 4.6 Hz, 1H), 5.89 (d, *J* = 5.9 Hz, 1H), 5.81 – 5.75 (m, 1H), 5.67 (d, *J* = 4.7 Hz, 1H), 4.72 – 4.61 (m, 1H), 4.48 – 4.38 (m, 2H), 3.92 – 3.81 (m, 1H), 2.71 – 2.55 (m, 2H), 1.81 – 1.64 (m, 2H), 1.50 (d, *J* = 8.2 Hz, 1H), 1.46 – 1.13 (m, 22H). ^31^P NMR (162 MHz, CD_3_OD) δ 3.07 (s). LC/MS *m/z* = 741.1 [M+1].

### (2R,3S,4S,5S)-5-(4-Aminopyrrolo[2,1-f][1,2,4]triazin-7-yl)-2-cyano-2-((((S)-(((S)-1-(cyclohexyloxy)-1-oxopropan-2-yl)amino)(phenoxy)phosphoryl)oxy)methyl)tetrahydrofuran-3,4-diyl dipropionate (29fm)

**28f** (25 mg, 0.042 mmol) was dissolved in anhydrous tetrahydrofuran (1 mL). Propionic acid (16 µL, 0.21 mmol) and *N*,*N*’-diisopropylcarbodiimide (32 µL, 0.21 mmol) were added to the reaction and stirred for 20 min. 4-Dimethylaminopyridine (5.0 mg, 0.042 mmol) was added, and the reaction was stirred for 4 h. The reaction was concentrated under reduced pressure, and the residue was purified via silica gel chromatography (0-100% ethyl acetate in hexanes) to afford the title compound (16 mg, 54%). ^1^H NMR (400 MHz, CDCl_3_) δ 7.89 (s, 1H), 7.35 – 7.22 (m, 2H), 7.22 – 7.09 (m, 3H), 6.65 (d, *J* = 4.5 Hz, 1H), 6.54 (d, *J* = 4.5 Hz, 1H), 5.95 (bs, 2H), 5.87 (d, *J* = 5.8 Hz, 1H), 5.81 (dd, *J* = 5.7, 3.9 Hz, 1H), 5.68 (d, *J* = 3.9 Hz, 1H), 4.73 (m, 1H), 4.41 (d, *J* = 6.3 Hz, 2H), 4.08 – 3.92 (m, 2H), 2.49 – 2.34 (m, 4H), 1.83 – 1.61 (m, 4H), 1.56 – 1.45 (m, 1H), 1.45 – 1.21 (m, 8H), 1.17 (m, 6H). ^31^P NMR (162 MHz, CDCl_3_) δ 2.46 (s). LC/MS *m/z* = 713.1 [M+1].

### (2R,3S,4S,5S)-5-(4-Aminopyrrolo[2,1-f][1,2,4]triazin-7-yl)-2-cyano-2-((((S)-(((S)-1-(cyclohexyloxy)-1-oxopropan-2-yl)amino)(phenoxy)phosphoryl)oxy)methyl)tetrahydrofuran-3,4-diyl diacetate (29fn)

**28f** (25 mg, 0.042 mmol) was dissolved in anhydrous tetrahydrofuran (1 mL). Acetic acid (12 µL, 0.21 mmol) and *N*,*N*’-diisopropylcarbodiimide (32 µL, 0.21 mmol) were added to the reaction and stirred for 20 min. 4-Dimethylaminopyridine (5.0 mg, 0.042 mmol) was added, and the reaction was stirred for 4 h. The reaction was concentrated under reduced pressure, and the residue was purified via silica gel chromatography (0-100% ethyl acetate in hexanes) to afford the title compound (24 mg, 84%). ^1^H NMR (400 MHz, CDCl_3_) δ 7.88 (s, 1H), 7.32 – 7.23 (m, 2H), 7.23 – 7.09 (m, 3H), 6.63 (d, *J* = 4.5 Hz, 1H), 6.52 (d, *J* = 4.5 Hz, 1H), 5.93 (s, 2H), 5.85 (d, *J* = 5.7 Hz, 1H), 5.79 (dd, *J* = 5.7, 4.1 Hz, 1H), 5.68 (d, *J* = 4.1 Hz, 1H), 4.73 (m, 1H), 4.42 (d, *J* = 6.4 Hz, 2H), 4.20 – 4.01 (m, 2H), 2.13 (s, 6H), 1.82 – 1.61 (m, 4H), 1.50 (m, 1H), 1.45 – 1.28 (m, 8H). ^31^P NMR (162 MHz, CDCl_3_) δ 2.49 (s). LC/MS *m/z* = 685.3 [M+1].

### ((2*R*,3*S*,4*R*,5*S*)-5-(4-Aminopyrrolo[2,1-*f*][1,2,4]triazin-7-yl)-3,4-dihydroxy-2-methyltetrahydrofuran-2-yl)methyl triphosphate (20-NTP)

To a solution of **20** (7.0 mg, 0.025 mmol) in PO(OMe)_3_ (0.2 mL) at 0 °C was added POCl_3_ (45 mg, 0.29 mmol). The reaction mixture was stirred at 0 °C for 4 h. A solution of pyrophosphate tributylamine salts (250 mg) in acetonitrile (0.5 mL) was added, followed by tributylamine (180 mg, 0.59 mmol). The reaction mixture was stirred at 0 °C for 6 h. The reaction was quenched with triethylammonium bicarbonate buffer (1 M, 8 mL). The reaction mixture was stirred at rt for 0.5 h, then concentrated under reduced pressure and co-evaporated with water twice. The residue was dissolved in water (2 mL) and NaHCO_3_ (400 mg) was added. The mixture was concentrated under reduced pressure. The residue was dissolved in water (∼2 mL) and loaded to a C-18 column, eluted with water. The product fractions were combined, acidified with HCl (1N, 150 µL) and concentrated to ∼4 mL, loaded to an ion-exchange column, eluted with water, then 10-40% triethylammonium bicarbonate buffer (1M) in water. The product fractions were combined and concentrated to give the title compound as the tetratriethylamine salt. ^1^H NMR (400 MHz, D_2_O) δ 7.73 (s, 1H), 6.94 (d, *J* = 4.7 Hz, 1H), 6.82 (d, *J* = 4.7 Hz, 1H), 5.24 (d, *J* = 8.9 Hz, 1H), 4.76 (dd, *J* = 9.0, 5.4 Hz, 1H), 4.34 (d, *J* = 5.3 Hz, 1H), 3.81 (dd, *J* = 10.8, 6.0 Hz, 1H), 3.72 (dd, *J* = 10.8, 4.8 Hz, 1H), 3.10 – 2.99 (m, 24H), 1.23 (s, 3H), 1.19 – 1.10 (m, 36H). ^31^P NMR (162 MHz, D_2_O) δ −6.26 (d, *J* = 20.4 Hz), −11.37 (d, *J* = 19.7 Hz), −22.48 (t, *J* = 20.1 Hz). MS *m/z* = 519.1 [M-1], 521.1 [M+1].

### ((2*R*,3*S*,4*R*,5*S*)-5-(4-Aminopyrrolo[2,1-*f*][1,2,4]triazin-7-yl)-3,4-dihydroxy-2-vinyltetrahydrofuran-2-yl)methyl triphosphate (21-NTP)

To a solution of **21** (10 mg, 0.034 mmol) in PO(OMe)_3_ (0.3 mL) at 0 °C was added POCl_3_ (50 mg, 0.33 mmol). The reaction mixture was stirred at 0 °C for 4 h. A solution of pyrophosphate tributylamine salts (250 mg) in acetonitrile (0.5 mL) was added, followed by tributylamine (170 mg, 0.71 mmol). The reaction mixture was stirred at 0 °C for 1 h. The reaction was quenched with triethylammonium bicarbonate buffer (1 M, 8 mL). The reaction mixture was stirred at rt for 0.5 h, then concentrated under reduced pressure and co-evaporated with water twice. The residue was dissolved in water (2 mL) and NaHCO_3_ (400 mg) was added. The mixture was concentrated under reduced pressure. The residue was dissolved in water (∼2 mL) and loaded to a C-18 column, eluted with water. The product fractions were combined, acidified with HCl (1N, 150 µL) and concentrated to ∼4 mL, loaded to an ion-exchange column, eluted with water, then 10-40% triethylammonium bicarbonate buffer (1M) in water. The product fractions were combined and concentrated to give the title compound as the tetratriethylamine salt. ^1^H NMR (400 MHz, D_2_O) δ 7.75 (s, 1H), 6.99 (d, *J* = 4.7 Hz, 1H), 6.84 (d, *J* = 4.6 Hz, 1H), 5.95 (dd, *J* = 17.4, 11.0 Hz, 1H), 5.46 (dd, *J* = 17.5, 1.6 Hz, 1H), 5.32 – 5.23 (m, 2H), 4.74 (dd, *J* = 8.7, 5.3 Hz, 1H), 4.50 (d, *J* = 5.3 Hz, 1H), 3.96 (dd, *J* = 11.0, 6.2 Hz, 1H), 3.66 (dd, *J* = 11.0, 4.5 Hz, 1H), 3.07 – 2.98 (m, 24H), 1.19 – 1.09 (m, 36H). ^31^P NMR (162 MHz, D_2_O) δ −6.21 (d, *J* = 20.8 Hz), −11.45 (d, *J* = 19.7 Hz), −22.44 (t, *J* = 20.2 Hz). MS *m/z* = 531.1 [M-1], 533.2 [M+1].

### ((2*R*,3*S*,4*R*,5*S*)-5-(4-Aminopyrrolo[2,1-*f*][1,2,4]triazin-7-yl)-2-cyano-3,4-dihydroxytetrahydrofuran-2-yl)methyl triphosphate (2-NTP)

To a solution of **2** (5.0 mg, 0.017 mmol) in PO(OMe)_3_ (0.6 mL) at 0 °C was added POCl_3_ (45 mg, 0.29 mmol). The reaction mixture was stirred at 0 °C for 10 h. A solution of pyrophosphate tributylamine salts (250 mg) in acetonitrile (0.6 mL) was added, followed by tributylamine (110 mg, 0.59 mmol). The reaction mixture was stirred at 0 °C for 0.5 h. The reaction was quenched with triethylammonium bicarbonate buffer (1 M, 5 mL). The reaction mixture was stirred at rt for 0.5 h, then concentrated and co-evaporated with water twice. The residue was dissolved in water (5 mL) and loaded to an ion-exchange column, eluted with water, then 5-35% triethylammonium bicarbonate buffer (1M) in water. The product fractions were combined, concentrated and co-evaporated with H_2_O. The residue was purified by ion-exchange column again to give crude material, so was repurified with C-18 column (0-15% acetonitirile in water with 0.05% triethylamine), and the fractions containing product were combined and concentrated. The material was dissolved in water (1 mL) and triethylammonium bicarbonate buffer (1 M, 0.1 mL) was added. The resulting mixture was concentrated under reduced pressure and co-evaporated with H_2_O twice under reduced pressure to afford the title compound as the tetratriethylamine salt. ^1^H NMR (400 MHz, D_2_O): δ 7.78 (s, 1H), 6.85 (d, *J* = 2.4 Hz, 1H), 6.82 (d, *J* = 2.4 Hz, 1H), 5.51 (d, *J* = 3.0 Hz, 1H), 4.65 – 4.55 (m, 2H), 4.20 – 4.08 (m, 2H), 3.15 – 3.00 (m, 24H), 1.18 – 1.08 (m, 36H). ^31^P NMR (162 MHz, D_2_O): δ - 6.25 (d, *J* = 52 Hz), −12.21 (d, *J* = 52 Hz), −22.32 (t, *J* = 52 Hz). MS *m/z* = 530.2 [M-1], 532.1 [M+1].

### ((2*R*,3*S*,4*R*,5*S*)-5-(4-Aminopyrrolo[2,1-*f*][1,2,4]triazin-7-yl)-2-(chloromethyl)-3,4-dihydroxytetrahydrofuran-2-yl)methyl triphosphate (22-NTP)

**22-NTP** was prepared as the tetra-triethylamine salt in a manner similar to that described for **2-NTP** starting with **22**. ^1^H NMR (400 MHz, D_2_O) δ 7.76 (s, 1H), 6.92 (br s, 1H), 6.85 (br s, 1H), 5.32 (d, *J* = 9.6 Hz, 1H), 4.78 (dd, *J* =8, 6.4 Hz, 1H), 4.53 (d, *J* = 5.6 Hz, 1H), 4.08 (dd, *J* = 10.0, 4.0 Hz, 1H), 3.83 – 3.95 (m, 3H), 3.07 (q, *J* = 7.6 Hz, 24 H), 1.16 (t, *J* = 7.6 Hz, 36 H). ^31^P NMR (162 MHz, D_2_O) δ −9.44 (d, *J* = 45.6 Hz), −11.51 (d, *J* = 48.8 Hz), −22.95 (t, *J* = 48.4 Hz). MS *m/z* = 555.06 [M+1].

### ((2*R*,3*S*,4*R*,5*S*)-5-(4-Aminopyrrolo[2,1-*f*][1,2,4]triazin-7-yl)-2-azido-3,4-dihydroxytetrahydrofuran-2-yl)methyl triphosphate (26-NTP)

To a solution of **26** (6.0 mg, 0.019 mmol) in PO(OMe)_3_ (0.6 mL) at 0 °C was added POCl_3_ (45 mg, 0.29 mmol). The reaction mixture was stirred at 0 °C for 10 h. A solution of pyrophosphate tributylamine salts (250 mg) in acetonitrile (0.6 mL) was added, followed by tributylamine (110 mg, 0.59 mmol). The reaction mixture was stirred at 0 °C for 6 h. The reaction was quenched with triethylammonium bicarbonate buffer (1 M, 5 mL). The reaction mixture was stirred at rt for 0.5 h, then concentrated and co-evaporated with water twice. The residue was dissolved in water (5 mL) and loaded to an ion-exchange column, eluted with water, then 5-35% triethylammonium bicarbonate buffer (1M) in water. The product fractions were combined, concentrated and co-evaporated with H_2_O. The residue was purified by ion-exchange column again to give crude material, so the material was repurified with ion-exchange column again to give crude material. The material was treated with NaHCO_3_ (10 mg) and the mixture was concentrated under reduced pressure. The solid residue was dissolved in water (0.5 mL) and 40 µL of NaOH (1N) was added. The resulting mixture was purified with C-18 column, eluted with H_2_O, and the fractions containing product were combined and concentrated under reduced pressure to afford the title compound as the tetra-sodium salt. ^1^H NMR (400 MHz, D_2_O): δ 7.76 (s, 1H), 6.88 (d, *J* = 4.3 Hz, 1H), 6.81 (d, *J* = 4.6 Hz, 1H), 5.59 (d, *J* = 5.5 Hz, 1H), 4.60 (t, *J* = 5.6 Hz, 1H), 4.55 (d, *J* = 5.8 Hz, 1H), 3.99 (qd, *J* = 11.2, 5.5 Hz, 3H). ^31^P NMR (162 MHz, D_2_O): δ −8.13 (d, *J* = 19.8 Hz), −14.04 (d, *J* = 18.9 Hz), −24.00 (t, *J* = 19.3 Hz). MS *m/z* = 546.1 [M-1], 547.9 [M+1].

### Cells and Viruses for Antiviral Assays

See the Supporting information.

### RSV A2 NHBE Antiviral and Cytotoxicity Assays

Compounds were evaluated for RSV potency in NHBE cells as described previously.^23^ Compounds were evaluated for cytotoxicity in NHBE cells as described previously.^33^

### RSV A2 HEp-2 Antiviral and Cytotoxicity Assays

Compounds were evaluated for RSV potency and cytotoxicity in HEp-2 cells as described previously.^23^

### MT4 Cytotoxicity Assay

Compounds were evaluated for cytotoxicity in MT4 cells as described previously.^23^

### RSV RNP, POLRMT, DNA Pol γ, and DNA Pol α Biochemical Assays

See the Supporting information.

### LogD

See the supporting information.

### 2-NTP in vitro metabolism

See the supporting information.

### RSV HAE Assay

Prior to infection, cells were washed twice with 1 × DPBS (ThermoFisher Scientific). All compounds were diluted in DMSO to a 1 μM working concentration. Compounds were further diluted in the assay media (MatTek) to the final concentrations of 0.25, 0.05, and 0.01 μM. 2 mL/well of the compound at the indicated concentration was added to the basolateral or 200 uL/well to the apical side of the transwell HAE culture. Wells containing apical compound or DMSO were incubated for 3 h at 37 °C after which compound containing media was aspirated and cells were washed once with 1X DPBS. Cells were infected apically with RSV A2 WT virus diluted in the assay media at an approximate MOI of 0.1 in 200 μL/well. Cells were incubated with the virus at 37 °C with 5% CO_2_ for 3 h. The virus was aspirated from the wells, and cells were further incubated for 24 h at 37 °C with 5% CO_2_. Each subsequent day for 2 d, DPBS was added apically to the cells at a final volume of 200 μL/well. Cells were incubated for 30 min at 37 °C with 5% CO2. DPBS was harvested from the wells, and treatment was freshly administered on days 1 and 2 by adding 2 mL/well of freshly diluted compounds (same concentration as before) to the basolateral side of the transwell HAE culture or 200 uL/well to the apical surface as on day 0. Cells were returned to the incubator at 37 °C with 5% CO_2_. On day 3, 200 μL of the DPBS was added apically to each well and the cells were incubated for 30 min at 37 °C with 5% CO_2_. 100 µL/well of the apical wash was harvested into 200 μL of lysis buffer (ThermoFisher Scientific). Xenotropic RNA (Vetmax Xeno) internal positive control (ThermoFisher Scientific) was included in the lysis buffer as a control for RNA isolation. Viral RNA was extracted from the collected samples using the MagMAx-96 Viral RNA isolation kit (Thermo-Fisher Scientific) and Kingfisher Flex instrument (ThermoFisher Scientific) as per the manufacturer’s instruction. Viral loads in HAE cultures were determined by RT-qPCR assay as below. Quantitative RT-PCR was performed using the TaqMan RNA-to-CT 1-Step kit (ThermoFisher Scientific). Briefly, 5 μL of the extracted viral RNA was mixed with 10 μL of the buffer from the kit, 0.5 μL of the RT enzyme, the appropriate primer/probe for either RSV N or the xenotropic RNA, and RT-PCR grade water (ThermoFisher Scientific) to a final volume of 20 μL. For the RSV-specific reactions, 0.5 μL of 40× RSV N Primer Probe (Integrated DNA Technology PrimeTime qPCR Assay XL; forward primer: 5′ GCTAGTGTGCAAGCAGAAATG 3′, reverse primer: 5′ TGGAGAAGTGAGGAAATTGAGTC 3′, and probe: 5′ FAM/ATTGGGTGG/ZEN/AGAAGCAGGGTTCTAC/3IABkFQ 3′ (Coralville, Iowa)) was added per reaction. For the 20× Vetmax Xeno Internal Positive Control-VIC Assay (ThermoFisher Scientific), 0.8 μL of the primer/probe set was added. Each reaction was performed in duplicate. PCR plates were sealed with a MicroAmp optical adhesive film (ThermoFisher Scientific) and centrifuged for 1 min at 1200 RPM. The one-step PCR reaction and subsequent amplification analysis were carried out using the QuantStudio 12K Flex Real Time PCR system using the following conditions: 48 °C for 15 min, 95 °C for 10 min, followed by 40 cycles of qPCR for 15 s at 95 °C and at 60 °C for 1 min. Reactions containing 10-fold serial dilutions of RSV A2 N gene with a pTM1 backbone solution were used to generate a standard curve against which the RSV RNA content measured from test samples was quantified. Median and interquartile ranges (IQR) of six technical replicates were determined for each treatment. The p values were determined by the One-way ANOVA with Dunnett correction.

### hMPV A1 LLC-MK2, RV-A 16 H1 HeLa, RV-B 14 H1 HeLa, EV-D68 RD, EV-71 RD, FluA H1N1 MDCK, and SARS-CoV-2 A549-hACE2 Antiviral and Cytotoxicity Assays

See the Supporting information.

### RSV Fluc NHBE, COPD HBE, Asthma HBE Antiviral Assays

Human bronchial epithelial cells (NHBE donor 6517, DHBE COPD donor 29522, DHBE asthma donor 36221) (5.0 × 10^3^/well) were seeded in white wall/clear bottom 96-well plates (Corning) with culture medium to a final volume of 100 μL and incubated for 24 h at 37 °C with 5% CO_2_. On the following day, 3-fold serial dilutions (starting at 9.28 nM and ending at 2000 nM) of 1 or 4 prepared in DMSO were added to the wells using the Hewlett-Packard D300 digital dispenser with normalization to the highest concentration of DMSO in all wells (>0.1% final volume). The cells were then infected with RSV-Fluc diluted with BEGM media at an MOI of 0.1 for a final volume of 200 μL media/well. Uninfected and untreated wells were included as controls to determine compound efficacy against RSV-Fluc. Following incubation with the compound and virus for 3 d at 37 °C with 5% CO_2_, 100 μL of culture supernatant was removed from each well and replaced with 100 μL of ONE-Glo luciferase reagent (Promega, Madison, WI, Cat# E6110). The plates were gently mixed by rocking for 10 min at rt, and luminescence signal was measured using an Envision plate reader (PerkinElmer). Values were normalized to the uninfected and infected DMSO controls (0 and 100% infection, respectively). Data was fit using nonlinear regression analysis using XLfit4. EC_50_ values were then determined as the concentration reducing the firefly luciferase signal by 50%. The compiled data was generated based on at least two independent experimental replicates, each containing technical duplicates for each concentration.

### In Vivo PK and Efficacy Methods

See the Supporting information.

## ASSOCIATED CONTENT

### Supporting Information

The supporting information is available free of charge. Supporting information includes additional experimental details for compound synthesis, cell-based assays and methods, biochemical assays and methods, in vitro and in vivo PK methods, in vivo intratracheal inhalation PK and RSV efficacy methods with supporting figure and data table, small molecule X-ray structure of compound **1**, HPLC and NMR spectral data (PDF) and molecular formula strings (CSV).

### Accession Codes

The Cambridge Crystallographic Data Center (CCDC) number for the small molecule X-ray structure of compound **1** is 2348357.

## AUTHOR INFORMATION

**Corresponding Author**

**Dustin S. Siegel** – *Medicinal Chemistry, Gilead Sciences Incorporated, Foster City, California, 94404, United States;*

**Notes.** The authors declare the following competing financial interest(s): Some authors are current or former employees of Gilead Sciences and may own company stock.

## ACKNOWLEDGEMENTS

We would like to extend the following acknowledgements: The Lovelace Biomedical support teams, Albuquerque NM, for the inhaled in vivo studies. The Bioqual team, Rockville MD, and Labcorp early development Inc. (legacy Covance Laboratories) team, Madison WI, for IV in vivo PK studies. The Arnold Rheingold Laboratory, UC San Diego, for assistance solving the crystal structure. The Guy Boivin Laboratory, Laval University, for the hMPV assay. The Southern Research Institute, Frederick MD, for the EV and influenza assays. From Gilead, Yelena Zherebina for chiral chromatography, and Krista McCutcheon for early assay development. Figure 3 was adapted from “brush cell (cuboidal)” by BioRender.com (2023).

## ABBREVIATIONS

A549-hACE2; human lung carcinoma cell line expressing angiotensin-converting enzyme 2; ARVI, acute respiratory virus infection; AGM, African green monkey; ANOVA, analysis of variance; AUC, area under the concentration-time curve; BAL, bronchoalveolar lavage; BALF, bronchoalveolar lavage fluid; BEGM, bronchial epithelial cell growth medium; CC_50_, half maximal cytotoxic concentration; CCDC, Cambridge crystallographic data center; CDI, carbonyldiimidazole; C_max_, maximal concentration; Conc., concentration; COPD, chronic obstructive pulmonary disease; COVID-19, coronavirus disease 2019; Cpd., compound; CPE, cytopathic effect; Cyno, cynomolgus monkey; DHBE, disease human bronchial epithelial cell; DIC, *N*,*N*′-diisopropylcarbodiimide; DMTr, dimethoxytrityl; DPBS, Dulbecco’s phosphate buffered saline; EC_50_, half maximal effective concentration; EDCI, 1-ethyl-3-(3-dimethylaminopropyl)carbodiimide; EV, enterovirus; FDA, Food and Drug Administration; Flu, influenza virus; Fluc, firefly luciferase; H1 HeLa, human cervical adenocarcinoma cell; HAE, human airway epithelial cell; HBE, human bronchial epithelial cell; HEp-2, human epithelial type 2 cell; hMPV, human metapneumovirus; IACUC, International Animal Care and Use Committee; Imid., imidazole; IQR, interquartile range; IT, intratracheal; IV, intravenous; LLC-MK2, rhesus monkey kidney cell; LOD, limit of detection; logD, logarithm of distribution coefficient; LRT, lower respiratory tract; MDCK, Madin-Darby canine kidney cell; mix, diastereomeric mixture; MOI, multiplicity of infection; NHBE, normal human bronchial epithelial cell; NMP, nucleoside monophosphate; NTP, nucleoside triphosphate; MT4, human leukemia T-cell; ODV, obeldesivir; PBS, phosphate-buffered saline; PK, pharmacokinetics; Pol, polymerase; POLRMT, human mitochondrial RNA polymerase; RdRp, RNA-dependent RNA polymerase; RDV, remdesivir; RNP, ribonucleoprotein; RSV, respiratory syncytial virus; RT-qPCR, quantitative reverse transcription polymerase chain reaction; RV, rhinovirus; SAR, structure-activity relationship; SARS-CoV-2, severe acute respiratory coronavirus-2; SNIR, single nucleotide incorporation rate; SD, standard deviation; t_1/2_, half-life.

---

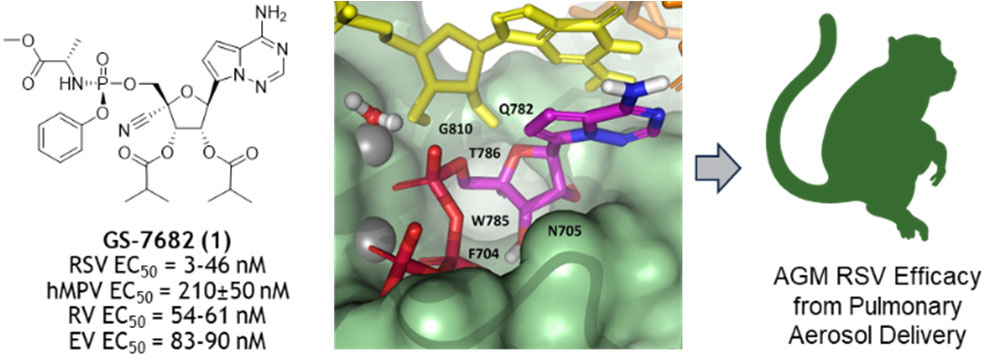
Table of Content Graphic.

