## Supporting Information for "Discovery and Synthesis of GS-7682, a Novel Prodrug of a 4′-CN-4-Aza-7,9-Dideazaadenosine *C*-Nucleoside with Broad-Spectrum Potency Against Pneumo- and Picornaviruses and Efficacy in RSV-Infected African Green Monkeys"

[¥]Lovelace Biomedical, Albuquerque, New Mexico 87108, USA.

[₸]Inspired – Pulmonary Solutions, San Carlos, California 94070, USA.

#### CONTENTS

**Compound Syntheses** ........ S2
**Cell-Based Assays and Methods** ........ S6
**Biochemical Assays and Methods** ........ S9
**Pharmacokinetic Methods** ........ S11
**In Vivo Intratracheal Inhalation PK and RSV Efficacy Methods** ........ S12
**Small Molecule X-ray Structure of Compound 1** ........ S16
**HPLC and NMR Spectral Data** ........ S17

#### COMPOUND SYNTHESES

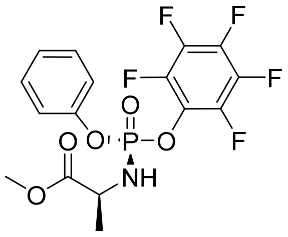

**27aj**

**Methyl ((*S*)-(perfluorophenoxy)(phenoxy)phosphoryl)-L-alaninate (27aj)**. L-Alanine methyl ester hydrochloride (14 g, 100 mmol) was mixed with 50mL of anhydrous DCM and stirred under atmospheric nitrogen in an ice bath. Phenyl dichlorophosphate (16.4 mL, 110 mmol) was added to the reaction dropwise, and the reaction mixture was stirred for 30 min. Triethylamine (29.4 mL, 210 mmol) was mixed with 20 mL anhydrous DCM and added to the reaction dropwise. The reaction was stirred for 1 h. Pentafluorophenol (18.4 g, 100 mmol) was added in one portion. Triethylamine (14.7 mL, 105 mmol) was mixed with 30 mL of anhydrous DCM and added to reaction dropwise. The reaction mixture was stirred for 16 h at rt and was diluted with DCM (50 mL) and washed with water (5 × 10 mL). The organic layer was dried over anhydrous sodium sulfate and then concentrated under reduced pressure. Isopropyl ether (130 mL) was added and then stirred for 24 h. The solids were collected, washed with isopropyl ether (30 mL) and dried under high vacuum to afford the title compound (22.2 g, 52%). $^{1}$H NMR (400 MHz, $CDCl_3$) δ 7.40 – 7.32 (m, 2H), 7.28 – 7.19 (m, 3H), 4.20 (m, 1H), 3.96 – 3.85 (m, 1H), 3.74 (s, 3H), 1.47 (d, *J* = 7.1 Hz, 3H). $^{31}$P NMR (162 MHz, $CDCl_3$) δ -1.62. $^{19}$F NMR (376 MHz, $CDCl_3$) δ -153.82 (dd, *J* = 18.5, 2.7 Hz), -159.99 (td, *J* = 21.8, 3.8 Hz), -162.65 (dd, *J* = 22.2, 17.6 Hz). LC/MS *m/z* = 425.9 [M+1], 423.9 [M-1].

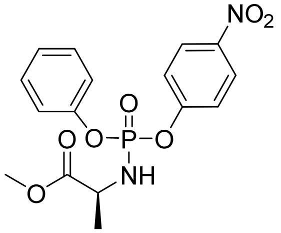

**27ak-mix**

**Methyl ((4-nitrophenoxy)(phenoxy)phosphoryl)-L-alaninate (27ak-mix).** Phenyl dichlorophosphate (2.81 mL, 18.9 mmol) and triethylamine (5.38 mL, 37.9 mmol) were sequentially added to a suspension of methyl L-alaninate hydrochloride (2.64 g, 18.9 mmol) in DCM (100 mL) at 0 °C. After 1 h, 4-nitrophenol (2.64 g, 18.9 mmol) and triethylamine (2.64 mL, 18.9 mmol) were then sequentially added at 0 °C, and the resulting mixture was then allowed to warm to rt. After 2.5 h, the reaction mixture was diluted with DCM (100 mL), washed with saturated aqueous sodium bicarbonate solution (100 mL) and brine (100 mL), dried over anhydrous sodium sulfate, and concentrated under reduced pressure. The crude residue was purified by silica gel chromatography (0-100% ethyl acetate in hexanes) to afford the title compound (2.02 g, 28%, 1:1 diastereomeric mixture). $^{1}$H NMR (400 MHz, $CDCl_3$) δ 8.25 – 8.18 (m, 2H), 7.43 – 7.29 (m, 4H), 7.29 – 7.15 (m, 3H), 4.24 – 4.07 (m, 1H), 3.97 (br q, *J* = 9.8

Hz, 1H), 3.70 (s, 3H), 1.45 – 1.35 (m, 3H). $^{31}$P NMR (162 MHz, $CDCl_3$) δ -3.12 (s), -3.17 (s). LC/MS *m/z* = 380.98 [M+1].

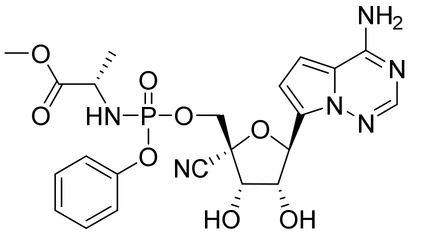

**28a-mix**

**Methyl ((((2*R*,3*S*,4*R*,5*S*)-5-(4-aminopyrrolo[2,1-*f*][1,2,4]triazin-7-yl)-2-cyano-3,4-dihydroxytetrahydrofuran-2-yl)methoxy)(phenoxy)phosphoryl)-L-alaninate (28a-mix).** Acetonitrile (5 mL) was added to a mixture of **18** (348 mg, 1.05 mmol), **27ak-mix** (399 mg, 1.05 mmol), and magnesium chloride (100 mg, 1.05 mmol) at rt. The mixture was heated to 50 °C for 10 min, and *N*,*N*-diisopropylethylamine (0.475 mL, 2.63 mmol) was added. After 2.5 h, the reaction mixture was allowed to cool to rt, and concentrated aqueous hydrochloric acid solution (0.5 mL) was added dropwise. After 1 h, the reaction mixture was diluted with ethyl acetate (100 mL) and the resulting mixture was washed with saturated aqueous sodium carbonate solution (100 mL) and brine (100 mL). The organic layer was dried over anhydrous sodium sulfate and concentrated under reduced pressure. The crude residue was subjected to silica gel chromatography eluting with 0-100% ethyl acetate in hexanes to afford the title compound (302 mg, 54%, ~1.75:1 diastereomeric mixture). $^{1}$H NMR (400 MHz, $CD_3OD$) δ 7.80 (s, 0.65H), 7.78 (s, 0.35H), 7.37 – 7.25 (m, 2H), 7.25 – 7.12 (m, 3H), 6.87 – 6.82 (m, 1H), 6.75 – 6.71 (m, 1H), 5.52 – 5.47 (m, 1H), 4.66 – 4.60 (m, 1H), 4.55 – 4.29 (m, 3H), 3.95 – 3.80 (m, 1H), 3.64 (s, 1H), 3.60 (s, 2H), 1.27 – 1.22 (m, 3H). $^{31}$P NMR (162 MHz, $CD_3OD$) δ 3.24 (s). LC/MS *m/z* = 533.13 [M+1].

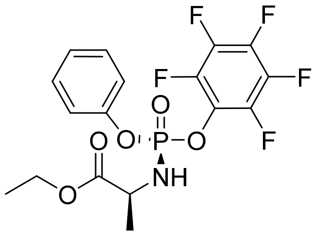

**27bj**

**Ethyl ((*S*)-(perfluorophenoxy)(phenoxy)phosphoryl)-L-alaninate (27bj).** To a solution of L-alanine ethyl ester hydrochloride (631 mg, 2.47 mmol) in DCM (15 mL) was added phenyl phosphorodichloridate (0.37 mL, 2.5 mmol) in one portion at 78 °C and triethylamine (0.68 mL, 4.93 mmol) was added dropwise over 5 min at -78 °C. After 30 min, pentafluorophenol (454 mg, 2.46 mmol) was added in one portion and triethylamine (0.34 mL, 2.465 mmol) added over 5 min at -78 °C. The dry ice bath was removed, and the resulting mixture was stirred for 1 h, then diluted with DCM, washed with brine, concentrated under reduced pressure, and the resulting residue purified by silica gel column chromatography (0-60% ethyl acetate in hexanes) to give a 1:1 diastereomeric mixture (566 mg, 49%). $^{1}$H NMR (400 MHz, $CDCl_3$) δ 7.43 – 7.30 (m, 2H), 7.32 – 7.17 (m, 3H), 4.29 – 4.11 (m, 3H), 3.94 (m, 1H), 1.52 – 1.42 (m, 3H), 1.28 (q, *J* = 7.0 Hz, 3H). Diisopropyl ether (4 mL) was added, and the suspension was sonicated and solids collected by filtration. Diisopropyl ether (5 mL) was added and the suspension was heated to 70 °C to a clear solution. The mixture was allowed to cool to rt, and the solids were collected by filtration dried under high vacuum for 30 min to afford the title compound (166 mg, 14%). $^{1}$H NMR (400 MHz, $CD_3CN$) δ 7.50 – 7.36 (m, 2H), 7.32 – 7.21 (m, 3H), 4.75 (t, *J* = 11.5 Hz, 1H), 4.17 – 3.98 (m, 3H), 1.37 (dd, *J* = 7.1, 1.1 Hz, 3H), 1.22 (t, *J* = 7.1 Hz, 3H). $^{31}$P NMR (162 MHz, $CD_3CN$) δ -0.51. $^{19}$F NMR (376 MHz, $CD_3CN$) δ -155.48 – -155.76 (m), -162.73 (td, *J* = 21.3, 3.7 Hz), -165.02 – -165.84 (m). LC/MS *m/z* = 440.5 [M-ethyl+H].

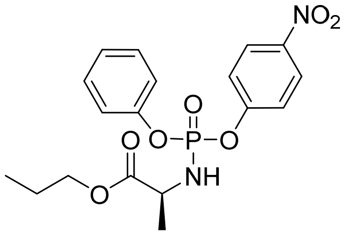


**27ck-mix**

**Propyl ((4-nitrophenoxy)(phenoxy)phosphoryl)-L-alaninate (27ck-mix).** Phenyl dichlorophosphate (0.89 ml, 5.97 mmol) in DCM (12 mL) was added dropwise over 15 min to a solution of propyl L-alaninate hydrochloride (1.0 g, 5.97 mmol) in DCM (12 mL) at 0 °C. Triethylamine (2.0 ml, 14.32 mmol) in DCM (2.5 mL) was added over 5 min. After 3.5 h, 4-nitrophenol (0.83 g, 5.97 mmol) and triethylamine (1.0 mL, 7.16 mmol) were then sequentially added at 0 °C, and the resulting mixture was then allowed to warm to rt. After 2 h, the reaction mixture was diluted with DCM (50 mL), washed with water (2 × 100 mL) and brine (50 mL), dried over anhydrous sodium sulfate, and concentrated under reduced pressure. The crude residue was purified by silica gel chromatography (0-100% ethyl acetate in hexanes) to afford the title compound (1.90 g, 76%, ~1:1 diastereomeric mixture). $^{1}$H NMR (400 MHz, $CD_3CN$) δ 8.28 - 8.20 (m, 2H), 7.49 - 7.35 (m, 4H), 7.31 - 7.19 (m, 3H), 4.72 - 4.56 (m, 1H), 4.14 - 4.02 (m, 1H), 3.99 (td, *J* = 6.6, 2.5 Hz, 2H), 1.58 (dtdd, *J* = 13.9, 7.4, 6.5, 0.9 Hz, 2H), 1.31 (ddd, *J* = 7.1, 4.2, 1.1 Hz, 3H), 0.88 (t, *J* = 7.4 Hz, 3H). $^{31}$P NMR (162 MHz, $CD_3CN$) δ -2.12, -2.22. LC/MS *m/z* = 409.12 [M+1].

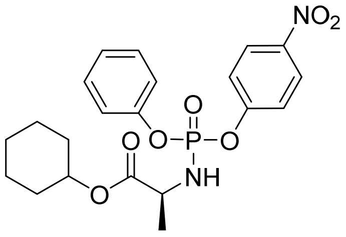


**27fk-mix**

**Cyclohexyl ((4-nitrophenoxy)(phenoxy)phosphoryl)-L-alaninate (27fk-mix).** L-Alanine, cyclohexyl ester hydrochloride (3.4 g, 16.37 mmol) was dissolved in DCM (45 mL), cooled to -78 °C, and phenyl dichlorophosphate (2.45 mL, 16.4 mmol) was added. Triethylamine (4.54 mL, 32.7 mmol) was added over 60 min at -78 °C and then 4-nitrophenol (2.3 g, 16 mmol) was added in one portion. Triethylamine (2.27 mL, 16.4 mmol) was added over 60 min at -78 °C. The resulting mixture was stirred for 2h at -78 °C and was diluted with DCM (100 mL), washed with water twice and brine, dried over anhydrous sodium sulfate, and concentrated under reduced pressure. The residue was purified by silica gel column chromatography (0-20% ethyl acetate in hexanes) to give the title compound (6.4 g, 87%, 1:1 diastereomeric mixture). $^{1}$H NMR (400 MHz, $CDCl_3$) δ 8.22 (m, 2H), 7.46 – 7.30 (m, 4H), 7.29 – 7.09 (m, 3H), 4.76 (m,

1H), 4.20 – 4.02 (m, 1H), 3.92 (m, 1H), 1.87 – 1.64 (m, 4H), 1.54 (m, 2H), 1.46 – 1.18 (m, 7H). $^{31}$P NMR (162 MHz, $CDCl_3$) δ -2.94, -3.00. LC/MS *m/z* = 449 [M+1].

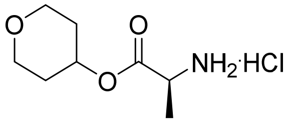

**[0001] (*S*)-tetrahydro-2*H*-pyran-4-yl 2-aminopropanoate hydrochloride.** To a mixture of L-alanine (500 mg, 5.61 mmol) and tetrahydro-2H-pyran-4-ol (5 g, 49.0 mmol) was added TMSCl (2 mL). The resulting mixture was stirred at 70 °C for 15 h and concentrated under reduced pressure. The resulting solid was triturated with 5% ethyl acetate in hexanes, filtered, and washed with 5% ethyl acetate in hexanes, and dried under reduced pressure to afford the title compound (1.33 g) that was used directly in the next step.

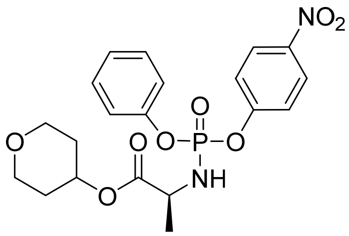

**27gk-mix**

**Tetrahydro-2*H*-pyran-4-yl ((4-nitrophenoxy)(phenoxy)phosphoryl)-L-alaninate (27gk-mix).** (*S*)-tetrahydro-2*H*-pyran-4-yl 2-aminopropanoate hydrochloride (1.33 g, 6.34 mmol) was dissolved in DCM (15 mL), cooled to -78 °C, and phenyl dichlorophosphate (1.14 mL, 7.61 mmol) was added. Triethylamine (2.2 mL, 15 mmol) was added over 30 min at -78 °C and the resulting mixture was stirred for 30 min. 4-Nitrophenol (882 mg, 6.34 mmol) was added and triethylamine (1.1 mL, 7.6 mmol) was added over 30 min at -78 °C. The mixture was stirred for 30 min and was then washed with water (20 mL) and brine (20 mL), dried over anhydrous sodium sulfate, and concentrated under reduced pressure. The residue was purified by silica gel column chromatography (0-70% ethyl acetate in hexanes) to afford the title compound (400 mg, 16%, 1:1 diastereomeric mixture). $^{1}$H NMR (400 MHz, $CDCl_3$) δ 8.22 (m, 2H), 7.49 – 7.06 (m, 7H), 4.95 (m, 1H), 4.14 (m, 1H), 4.07 – 3.80 (m, 3H), 3.52 (m, 2H), 1.95 – 1.81 (m, 2H), 1.64 m, 2H), 1.42 (m, 3H). $^{31}$P NMR (162 MHz, $CDCl_3$) δ -3.09, -3.13. LC/MS *m/z* = 451 [M+1].

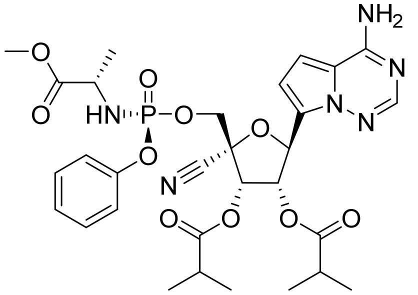

**1**

**Chiral chromatography method for (2*R*,3*S*,4*S*,5*S*)-5-(4-Aminopyrrolo[2,1-*f*][1,2,4]triazin-7-yl)-2-cyano-2-((((*S*)-(((*S*)-1-methoxy-1-oxopropan-2-**

**yl)amino)(phenoxy)phosphoryl)oxy)methyl)tetrahydrofuran-3,4-diyl bis(2-methylpropanoate) (1). 28a-mix** (54 mg, 0.10 mmol) was dissolved in 2 mL of anhydrous tetrahydrofuran. Isobutyric acid (37 µL, 0.40 mmol) and *N,N'*-diisopropylcarbodiimide (62 µL, 0.40 mmol) were added and the reaction mixture was stirred for 30 min. DMAP (12 mg, 0.1 mmol) was added and reaction mixture was stirred for 16 h. Methanol (0.5 mL) was added and the reaction mixture was stirred for 20 min. The reaction mixture was purified directly with preparative HPLC (Phenominex Synergi 4u Hydro-RR 80Å 150 x 30 mm column, 10-40% acetonitrile in water gradient) to afford **29al-mix** (46 mg, 68%, ~1.75:1 diastereomeric mixture). $^1$H NMR (400 MHz, $CD_3OD$) δ 7.90 – 7.80 (m, 1H), 7.36 – 7.24 (m, 2H), 7.24 – 7.09 (m, 3H), 7.10 – 7.00 (m, 1H), 6.86 – 6.78 (m, 1H), 5.96 – 5.74 (m, 2H), 5.73 – 5.65 (m, 1H), 4.58 – 4.38 (m, 2H), 3.96 – 3.71 (m, 2H), 3.63 (m, 3H), 2.73 – 2.54 (m, 2H), 1.33 – 1.13 (m, 12H), 1.09 (d, $J$ = 6.5 Hz, 3H). $^{31}$P NMR (162 MHz, $CD_3OD$) δ 3.04. LC/MS *m/z* = 673.2 [M+1]; 671.3 [M-1]. Resolution of the *S*p and *R*p diastereomers was conducted via chiral preparatory HPLC (Chiralpak AD-H, 150 x 4.6 mm, SFC 35% isopropyl alcohol isocratic) from 40 mg of the mixture. First eluting *R*p diastereomer **29al-*Rp*** (8 mg). $^1$H NMR (400 MHz, $CD_3OD$) δ 7.81 (s, 1H), 7.34 – 7.27 (m, 2H), 7.19 – 7.13 (m, 3H), 6.86 (d, $J$ = 4.5 Hz, 1H), 6.77 (d, $J$ = 4.6 Hz, 1H), 5.99 (d, $J$ = 5.9 Hz, 1H), 5.86 (dd, $J$ = 5.8, 4.3 Hz, 1H), 5.69 (d, $J$ = 4.3 Hz, 1H), 4.52 (dd, $J$ = 11.1, 5.8 Hz, 1H), 4.41 (dd, $J$ = 11.1, 4.8 Hz, 1H), 3.87 – 3.73 (m, 1H), 3.63 (s, 3H), 2.74 – 2.58 (m, 2H), 1.27 – 1.13 (m, 15H). $^{31}$P NMR (162 MHz, $CD_3OD$) δ 3.02 (s). LC/MS *m/z* = 673.2 [M+1]. Second eluting *S*p diastereomer **1** (19.5 mg). $^1$H NMR (400 MHz, $CD_3OD$) δ 7.83 (s, 1H), 7.34 – 7.27 (m, 2H), 7.22 – 7.13 (m, 3H), 6.84 (d, $J$ = 4.5 Hz, 1H), 6.75 (d, $J$ = 4.6 Hz, 1H), 5.92 (d, $J$ = 5.8 Hz, 1H), 5.81 (dd, $J$ = 5.9, 4.7 Hz, 1H), 5.68 (d, $J$ = 4.7 Hz, 1H), 4.49 – 4.37 (m, 2H), 3.97 – 3.83 (m, 1H), 3.61 (s, 3H), 2.73 – 2.55 (m, 2H), 1.31 – 1.13 (m, 15H). $^{31}$P NMR (162 MHz, $CD_3OD$) δ 3.02 (s). LC/MS *m/z* = 673.2 [M+1].

#### CELL-BASED ASSAYS AND METHODS

##### VIRUSES

Respiratory syncytial virus strain A2 (RSV A2) without a reporter gene or expressing the firefly luciferase transgene (RSV-Fluc) was obtained as a high-titer stock from Microbiologics (Saint Cloud, MN). The hMPV CAN99-81 (A1 genotype) was isolated and propagated in the lab a Guy Boivin.[1] EV-D68 (18952), EV-71(H), RV-A 16, and RV-B 14 viruses were obtained from ATCC (Manassas, VA).

##### CELLS

Multiple donors of normal human bronchial epithelial (NHBE) cells were purchased from Lonza (Walkersville, MD, Cat # CC-2540) and cultured in Bronchial Epithelial Growth Media (BEGM) (Lonza, Walkersville, MD, Cat # CC-3170). The cells were passaged 1-2 times per week to maintain < 80% confluency. The NHBE cells were discarded after 6 passages in culture.

Disease human bronchial epithelial (DHBE) cells were purchased from Lonza (Walkersville, MD, Cat # 00194911 Asthma Donor 36221, Cat # 00195275 COPD Donor 29522) and cultured in Bronchial Epithelial Growth Media (BEGM) (Lonza, Walkersville, MD, Cat # CC-3170). The cells were passaged 1-2 times per week to maintain < 80% confluency. The NHBE cells were discarded after 6 passages in culture.

The HEp-2 cell line was purchased from ATCC (Manassas, VA Cat # CCL-23) and maintained in DMEM (Corning, New York, NY, Cat # 15-018CM) supplemented with 10% FBS (Hyclone, Logan, UT, Cat # SH30071-03) and 1× Penicillin-Streptomycin-L-Glutamine

(Corning, New York, NY, Cat #30-009-CI). Cells were passaged 2 times per week to maintain sub-confluent densities and were used for experiments at passages 5-20.

The MT-4 cell line was obtained from the NIH AIDS Research and Reference Reagent Program and cultured in RPMI-1640 medium supplemented with 10% FBS, 100 units/mL penicillin and streptomycin, and 2 mM L-glutamine.

Human airway epithelial (HAE) cells AIR-100 were sourced from MatTek Corporation (Ashland, MA). Differentiated HAE cultures were maintained as per the manufacturer's instruction.

Rhabdomyosarcoma cells (RD; CCL-136), H1 HeLa (CRL-1958), LLC-MK2 (CCL-7), and Madin-Darby canine kidney (MDCK; CCL-34) cells were obtained from ATCC. All cell lines were grown in either Eagle's Minimum Essential Medium (EMEM) (Gibco/Thermo Fisher Scientific) or Dulbecco's Modified Eagle's Medium (DMEM) (Gibco/Thermo Fisher Scientific) supplemented with 10% FBS, 2.0 mM L-Glutamine, 100 units/mL penicillin and 100 µg/mL streptomycin. Cells were sub-cultured twice a week using standard cell culture techniques.

A549-hACE2 cells that stably express human angiotensin-converting enzyme 2 (hACE2) were established and provided by the University of Texas Medical Branch.[2] A549-hACE2 cells were grown in Dulbecco's Modified Eagle's Medium (DMEM) (Gibco/Thermo Fisher Scientific) supplemented with 10% FBS, 2.0 mM L-Glutamine, 100 units/mL penicillin and 100 µg/mL streptomycin and 10 µg/mL Blasticidin.

**HMPV A1 $EC_{50}$**

Antiviral analyses hMPV A1 was performed in the lab of Guy Boivin, Research Center in Infectious Diseases, CHU of Quebec and Laval University, Quebec City. LLC-MK2 cells were seeded in 24-well plates at 90% confluence in MEM + 10% FBS. Plates were washed with PBS and 500 µL of hMPV (strain CAN99-81 (gr. A1)), containing 40 PFU/mL with the different dilutions of the 4 compounds were added on the cells for 1 h 30 min at 37 °C, 5% $CO_2$. After the adsorption period, the medium was removed and cells were washed with PBS. An overlay of Opti-MEM 2x + methycellulose 1.6% (1:1) containing the same concentrations of compounds as in the adsorption period was added to the wells. A positive control consisting of ribavirin was also tested with concentrations ranging from 1 µM to 100 µM. The plates were incubated at 37 °C, 5% $CO_2$ for 3-5 d. After the incubation period, the cells were fixed with 7% formalin for 1 h at rt. hMPV titers were determined using a Mab specific for the F protein (clone hMPV24, ABD Serotec, Inc.) diluted 1/10,000 and a second anti-mouse IgG HRP antibody (GeneScriptCorp) diluted 1/2,500. Finally, KPL True Blue was used to reveal infected cells.

**LLC-MK2 $CC_{50}$**

LLC-MK2 cells were seeded at 7500 cells per cell and incubated overnight at 37 °C, 5% $CO_2$. The next day, compound serially diluted 5-fold titration was added to each well and the plate returned to 37 °C, 5% $CO_2$. After 5 days, LLC-MK2 cytotoxicity plates were stained with the soluble tetrazolium-based dye MTS (3- (4,5-dimethylthiazol-2-yl) -5- (3-carboxymethoxyphenyl) -2- (4-sulfophenyl) -2H-tetrazolium; CellTiter®96 Reagent, Promega) to determine cell viability and quantify compound toxicity. At termination of the assay, 20-25 µL of MTS reagent was added per well and the microtiter plates were then incubated for 2-4 h at 37 °C, 5% $CO_2$ to assess cell viability. Adhesive plate sealers were used in place of the lids, the sealed plate was inverted several times to mix the soluble formazan product and the plate was read spectrophotometrically at 490/650 nm with a Molecular Devices SpectraMax i3 plate reader.

#### RV-A 16, RV-B 14 H1 HELA $EC_{50}$

Test molecules are prepared in 100% DMSO in 384-well polypropylene plates (Greiner, Monroe, NC, Cat# 784201) with 4 replicates at 10 serially diluted concentrations (1:3). The serially diluted compounds were transferred to low dead volume Echo plates (Labcyte, Sunnyvale, CA, Cat# LP-0200). The test compounds were spotted to 384-well assay plates (Greiner, Monroe, NC, Cat# 781091) at 200 nL per well using an Echo acoustic dispenser (Labcyte, Sunnyvale, CA).
H1-HeLa cells were harvested and suspended in DMEM (supplemented with 2% FBS and 1× Penicillin-Streptomycin-L-Glutamine) and seeded to the pre-spotted assay plates at 5,000 cells per well in 30 µL. RV14 and RV16 was diluted in DMEM (supplemented with 2% FBS and 1× Penicillin-Streptomycin-L-Glutamine) at 500,000 Infectious Units (IU) per mL and 125,000 IU per mL respectively. 10 µL of virus per well was added to the assay plates containing cells and compounds, for an MOI of 1.0, and 0.25 respectively. The assay plates were incubated for 4 d at 37 °C and 5% $CO_2$. At the end of incubation, Celltiter-Glo (Promega, Madison, WI, Cat # G7573) was prepared. The assay plates and Celltiter-Glo reagent were equilibrated to rt for at least 15 min. 40 µL per well of Celltiter-Glo reagent was added and the plates were incubated at rt for 15 min before reading the luminescence signal on an EnVision multimode plate reader (Perkin Elmer, Waltham, MA). Rupintrivir was used as positive control and DMSO was used as negative control. Values were normalized to the positive and negative controls (as 0% and 100% replication, respectively) and data was fitted using four-parameter non-linear regression analysis. $EC_{50}$ was defined as the concentration reducing viral replication by 50%.

#### H1 HELA $CC_{50}$

H1 HeLa cells were plated in 96-well plates (Corning Life Sciences) at 0.1 mL/well at 0.12 × $10^6$ cells/mL in growth medium. After the cells were incubated at 37 °C in a humid atmosphere containing 5% (v/v) $CO_2$ overnight, 0.1 mL per well growth medium was added to each well along with serial diluted test compounds dispensed by a HP D300e digital dispenser (Hewlett Packard) with a final volume of 0.2 mL/well. The culture was maintained in a humidified chamber at 33 °C with 5% (v/v) $CO_2$ for 96 h. After the incubation, H1 HeLa cell viability was measured using the CellTiter-Glo Luminescent Cell Viability Assay kit (Promega) according to the manufacturer's protocol. The luminescence signals were recorded by an EnVision plate reader. The relative cell viability was calculated by normalizing the absorbance of the compound-treated wells to those of the DMSO-treated wells (set as 100%). The relative cell viability (Y-axis) versus the log10 values of compound concentration (X-axis) were plotted in software Prism (version 8). The $CC_{50}$ (the concentration of test compounds required to reduce cell viability by 50%) values were calculated using a four-parameter nonlinear regression model.

#### EV-D68, EV-71, AND FLUA H1N1 $EC_{50}$ AND $CC_{50}$

Antiviral analyses for EV-D68 (18952), EV-71(H), and Influenza (H1N1) were performed by Southern Research Institute, Frederick, MD, and were conducted in RD (enteroviruses), and MDCK cells (influenza), using cytopathic effect (CPE) assays. Virus and cells were mixed in the presence of test compound and incubated for the indicated duration. The virus was pre-titered such that control wells exhibited 85 to 95% loss of cell viability due to virus replication. Therefore, antiviral effect or cytoprotection was observed when compounds prevented virus replication. Cytoprotection and compound cytotoxicity was assessed by MTS (CellTiter®96 Reagent, Promega) dye reduction. The % reduction in viral cytopathic effects (CPE) was determined and used to calculate $EC_{50}$ (concentration inhibiting virus replication by 50%) and $CC_{50}$ (concentration resulting in 50% cell death).

#### SARS-COV-2 $EC_{50}$

SARS-CoV-2 activity was determined using a firefly reporter SARS-CoV-2 strain as previously described.[3]

#### A549-HACE2 $CC_{50}$

Compounds (200 nL) were spotted onto 384-well black plates (Greiner 781086) prior to seeding 5,000 A549-hACE2 cells per well in a volume of 40 µL of assay medium (DMEM supplemented with 2% FBS and 1× Penicillin-Streptomycin). The plates were incubated for 28 h at 37 °C with 5% $CO_2$. Following the incubation, 40 µL of CellTiter-Glo (Promega) was added and mixed 5 times. Plates were read for luminescence on an EnVision multimode plate reader. Untreated cells and cells treated with 2 µM puromycin (Sigma, St. Louis, MO) serve as 100% and 0% cell viability control, respectively. The percent of cell viability is calculated for each tested compound concentration relative to the 0% and 100% controls and the $CC_{50}$ value is determined by four-parameter non-linear regression as the compound concentration reducing the cell viability by 50%.

### BIOCHEMICAL ASSAYS AND METHODS

#### REAGENTS AND ENZYMES

All natural deoxynucleoside triphosphates (dNTPs) and nucleoside triphosphates (NTPs) were from GE Healthcare (Piscataway, NJ). The [$\alpha$-$^{33}$P]dNTPs and [$\alpha$-$^{33}$P]NTPs were purchased from PerkinElmer (Waltham, MA). All biochemical assays used radiolabeled dNTP or NTP to track DNA or RNA product formation. The products were analyzed using affinity filter-binding or electrophoresis systems and quantified using a Typhoon Trio imager and ImageQuant TL software (GE Healthcare). All concentrations refer to the final concentrations unless mentioned otherwise. Product formation in the presence of the inhibitors was expressed as a percentage of the product in water-treated controls (defined as 100%). The $IC_{50}$ was defined as the concentration at which there was a 50% decrease in product formation. The data were analyzed using GraphPad Prism (v9). $IC_{50}$ values were calculated as an average of at least three independent experiments.

RSV ribonucleoprotein (RNP) complexes were prepared according to a method modified from published methods.[4] The recombinant human mitochondrial RNA polymerase (POLRMT) was purchased from Enzymax. RNA polymerase II was purchased as part of the “HeLaScribeR Nuclear Extract in vitro Transcription System” kit from Promega. Recombinant human DNA polymerase β was purchased from ProFoldin (Hudson, MA). Recombinant human DNA polymerase γ (including both the large subunit and the small subunit) was cloned, overexpressed, and purified based on published methods.[5,6]

#### PROTEIN EXPRESSION AND PURIFICATION OF DNA POLYMERASE ALPHA CATALYTIC SUBUNIT POLA1

The POLA1 gene sequence encoding amino acid E338-E1255 was cloned into a pET24 vector in frame with N-terminal 6x-His, and SUMO tag. Expression of the recombinant His-SUMO-POLA1 fusion protein was achieved by transforming the plasmid into Rosetta2 DE3 cells and inducing *E. coli* culture in LB mediums with IPTG to a final concentration of 0.5 mM at 18 °C for 16 h. Induced culture was harvested and stored at -80 °C. For protein purification, the cell pellet was resuspended in Buffer A (50 mM Tris-HCl pH 8.0, 500 mM NaCl, 2 mM DTT, 0.1% Tween 20, 10% glycerol, 20 mM imidazole) supplemented with EDTA-free Protease inhibitor

(Roche). The cell suspension was passed through a microfluidics high-pressure homogenizer (Step-I) and the cell lysate was treated with polyethyleneimine (PEI) to a final concentration of 0.25% (v/v) at 4 °C for 30 min with constant mixing using a magnetic bar and stirrer. The lysate was centrifuged at 15,000 rpm for 30 min at 4 °C and the supernatant was treated with ammonium sulfate powder to 40% saturation. The sample was centrifuged at 15,000 rpm for 30 min and the pellet was resuspended in Buffer A to homogeneity. The resuspended sample was mixed with Ni-NTA resin for 90 min at 4 °C, washed with Buffer A, and eluted in Buffer B (50 mM Tris-HCl pH 8.0, 400 mM NaCl, 2 mM DTT, 0.1% Tween 20, 250 mM Imidazole, and 20% glycerol). The eluted samples were loaded over a Hi-Load 16/600 Superdex 200 PG column equilibrated with SEC buffer (50 mM Tris-HCl [pH 8.0], 400 mM NaCl, 2 mM DTT, 0.1% Tween-20, 20% glycerol). A major contaminant band around 70 KD was still co-purified with POLA1. The purified sample was adjusted to heparin-binding buffer (50 mM Tris-HCl [pH 8.0], 0.2 M NaCl, 2 mM DTT, 0.05% Tween-20, 20% glycerol) and loaded over a HiTrap Heparin HP column (Cytiva) equilibrated with the same buffer. The target protein was eluted using a linear gradient using a Heparin elution Buffer (50 mM Tris-HCl [pH 8.0], 1.0 M NaCl, 2 mM DTT, 0.05% Tween-20, 20% glycerol). The purest fraction was further dialyzed in the storage buffer (50 mM Tris-HCl [pH 8.0], 0.4 M NaCl, 2 mM DTT, 0.05% Tween-20, 20% glycerol), flash frozen in liquid nitrogen, and stored at -80 °C. The A260/A280 of the final prep was calculated to be 0.58 indicating negligible nucleic acid contamination. The protein prep was mass-verified by intact LC-MS analysis. In an alternative purification method, the resuspended lysate from Step-I was purified using Ni-NTA resin followed by a Hi-Trap Heparin column as described above. In the heparin purification steps, two different peaks were observed. The second peak eluted later in the salt gradient was collected and subsequently dialyzed in the storage buffer as above. The two-step purification method (Ni-NTA + Hi-Trap Heparin) showed a higher A260/A280 ratio of 0.76 although both purification methods yielded active POLA1 protein.

#### BIOCHEMICAL ASSAYS

All biochemical assays were performed as described previously.[7,8] RSV $IC_{50}$ assays was performed using crude RSV RNP complexes in the presence of 20 µM ATP, 100 µM GTP, 100 µM UTP, 100 µM CTP, and 1.5 µCi [α-$^{32}$P]ATP [3,000 Ci/mmol]) the final RNA product formation was separated on a 1.2% agarose/MOPS gel containing 2M formaldehyde and quantified using a phosphorimager. Inhibitions of human DNA alpha and gamma (mitochondria DNA) polymerase and mitochondria RNA polymerase were measured in $IC_{50}$ assays using recombinant proteins in the presence of all four natural dNTPs or NTPs. The elongated DNA or RNA products were captured on DE81 anion exchange paper and quantified using a phosphoimager. The relative incorporation rate of each analog was measured in the Single Nucleotide Incorporation (SNI) assays in the presence of the recombinant proteins and 50 µM of dATP or analog for DNA polymerases, or 500 µM ATP or analog for RNA polymerases. The rate of SNI was expressed as the percentage of the incorporation rate of the natural substrate, dATP or ATP. The reaction mixture was quenched with EDTA at selected time points from 0-60 min, run on a 20% polyacrylamide gel (8 M urea), and the product formation was quantified using Typhoon Trio Imager and Image Quant TL Software (GE Healthcare, Piscataway, NJ). The rate of SNI was calculated by fitting the product formation using the single exponential equation: $[\text{Product}] = A(1 - e^{-kt})$, where [Product] represents the amount (in nM) of the elongated product formed, $t$ represents the reaction time, $k$ represents the observed rate, and A represents the amplitude of the exponential.

#### PHARMACOKINETIC METHODS

##### LOGD

A solution of test compound in DMSO (2 μL of 10 mM) was added into a 96-well plate containing 198 μL of 1:1 acetonitrile/water. The sample plate was shaken for 30 min, and 10 μL of the resulting solution was analyzed by HPLC/UV. The instrumentation used was an Alliance 2795 HPLC coupled with a photodiode array detector 2996 (Waters, Milford, MA) using a Waters XTerra 3.5 μm 4.6 mm × 50 mm C18 column. The mobile phase consisted of solvent A (20 mM ammonium acetate aqueous solution) and solvent B (100% acetonitrile). Elution was performed using a linear gradient of solvent B from 0% to 100% in 8 min. The log D value was calculated using the retention time of test compound compared to the reference compounds. The range of log D values for reference compounds is approximately between 0.3 and 5.7.

##### 2-NTP IN VITRO METABOLISM

NHBE cells were seeded in a 6-well at 1 x $10^5$ cells/well (donor 6517) or a 12-well plate at 2.5 x $10^5$ cells/well (all other donors). DHBE COPD (donor 29522) and DHBE asthma (donor 36221) were seeded in a 12-well plate at 2.5 x $10^5$ cells/well. H1 Hela cells were seeded in a 12-well plate at 0.3 x $10^6$ cells/well. For continous treatment, the cells were continuously incubated with 0.2 μM (H1 Hela only) or 1 μM **1** over a 48-h period at 37 °C with 5% $CO_2$. For pulse treatment, compound-containing media (0.2 μM **1**) was removed 2 h post addition, washed twice with PBS, and replaced with fresh media before cultures were returned to the incubator at 37 °C. At the indicated time post-drug addition, cells were washed 3 times with ice-cold tris-buffered saline, scraped into 0.5 mL ice-cold 70% methanol and stored at -80 °C. Extracts were centrifuged at 15,000 g for 15 min and supernatants were transferred to clean tubes for evaporation in a miVac Duo concentrator (Genevac, Gardiner, NY). Dried samples were reconstituted in mobile phase A containing 3 mM ammonium formate (pH 5) with 10 mM dimethylhexylamine (DMHA) in water for analysis by LC-MS/MS, using a multi-stage linear gradient from 10% to 50% acetonitrile in mobile phase A at a flow rate of 350 μL/min. Analytes were separated using a 50 x 2 mm, 2.5 μm Luna C18(2) HST column (Phenomenex) connected to an LC-20ADXR (Shimadzu, Columbia, MD) ternary pump system and an HTS PAL autosampler (LEAP Technologies). Detection was performed on a Qtrap 6500+ (AB Sciex, Framingham, MA) mass spectrometer operating in negtive ion and multiple reaction monitoring modes. Analytes were quantified using a 7-point standard curve ranging from 0.274 to 200 pmol per million cells prepared in extracts from untreated culture wells that were counted for each time point.

##### IN VIVO INTRAVENOUS PHARMACOKINETICS

In vivo intravenous (IV) pharmacokinetic (PK) studies were performed at either Bioqual (Rockville, MD) or Covance Laboratories Inc. (Madison, WI) in accordance with local IACUC guidelines. **1** was formulated in 12% sulfobutylether-β-cyclodextrin in water (pH 3) at 2.5 mg/mL and male cynomolgus monkeys received a single IV infusion over 30 min. The rate of infusion was adjusted according to the body weight of each animal to deliver a dose of 5 mg/kg. **1** and **2** concentrations were measured in plasma collected over a 24-h sampling period. Serial blood collection was performed from 3 animals and processed to plasma. Aliquots of 25 μL of plasma were added to a mixture containing 200 μL of acetonitrile solution containing internal standard solution and centrifuged, and 200 μL of resulting supernatant was then transferred, dried under a stream of nitrogen at 40°C, and reconstituted in a mixture of 5% acetonitrile and 95% water. Plasma concentrations were determined using an 8-point calibration curve spanning at least 3 orders of magnitude with quality control samples to ensure accuracy and precision, all prepared in naiv̈e plasma. Analytes were separated on a 50 × 3.0 mm, 2.5 μm Synergi Polar-RP column (Phenomenex, Torrance, CA) using a multistage linear gradient with mobile phases containing 0.1% formic acid in either water or acetonitrile at a flow rate of 0.7 mL/min. Pharmacokinetic parameters were calculated using Phoenix (Certara, Princeton, NJ). The lung tissues were collected as a non-survival surgical procedure at approximately 24 h following the IV administration., and **2-NTP** concentrations in these lung tissues were measured. Animals were sedated with ketamine administered via an intramuscular injection, followed by intravenous fentanyl and propofol (to effect) administered to induce a deep surgical plane of anesthesia. Animals were intubated and manually ventilated with 100% oxygen to facilitate collection of a section of the lower lung from the left caudal lung lobe for immediate placement into liquid nitrogen. Frozen tissues were pulverized using a cell crusher and transferred into pre-weighed conical tubes, all performed and maintained on dry ice. Tissues were weighed, and a 4-fold volume of dry ice-cold extraction buffer (70% methanol containing 0.1% KOH, 67 mM EDTA, and 250 nM chloro-ATP as internal standard) was added. Resulting mixtures were then promptly homogenized. Undosed control tissues were used for quantification of the metabolites in lung tissue, generated by spiking an appropriate amount of metabolite standard solution into control tissues. An aliquot of the homogenate was filtered, evaporated to dryness, and reconstituted with 1 mM ammonium phosphate buffer for analysis by LCMS/MS. Analysis was performed using similar methods as previously described.[9]

#### IN VIVO INTRATRACHEAL INHALATION PK AND RSV EFFICACY METHODS

*Study Design*

In vivo intratracheal (IT) inhalation PK in cynomolgus monkeys and AGMs and RSV efficacy studies in AGMs were performed at Lovelace Biomedical Research Institute (LBRI, Albuquerque, NM) in accordance with local IACUC guidelines. **1** was aerosolized and delivered over 30 min via an endotracheal tube to cynomolgus monkeys (n=3 females) or AGMs (2 males/1 female). Plasma and lung tissue PK was assessed in cynomolgus monkeys or AGM following a single IT dose of **1** via a procedure identical to the IV dosing protocol; accordingly, **1** and **2** concentrations in plasma were measured over a 24-h sampling period, and **2-NTP** concentrations in the lung tissues were measured at approximately 24 hours post-dose.

To evaluate RSV efficacy, 14 AGMs (7 male/7 female distributed across dosing groups) were utilized in two sequential studies to compare inhaled **1** against the vehicle control (n=3 cohort 1 and n=4 cohort 2 per treatment group per study). On study day 0, animals were infected with $4 \times 10^6$ $TCID_{50}$ RSV by combined intranasal and intratracheal instillation. Approximately one hour following RSV infection, animals received treatment with either **1** or vehicle delivered by IT inhalation. Treatment was continued once daily for a total of 6 d. Bronchoalveolar lavage (BAL) fluid and nasal swabs were collected at baseline, on study day 1, and every other day thereafter through 15 d post-infection for assessment of viral load by RT-qPCR.

*Reagents and Formulations*

A suspension formulation of **1** for aerosolization was prepared as follows. Solid crystalline **1** particles in the suspension formulation in 0.1% Tween-20 + 150 mM NaCl at 15 mg/mL (cyno study) or 0.1% HPMC in 25mM Phosphate Buffer + 150 mM NaCl (AGM studies) were sized by wet milling on a Netzsch agitator bead mill to reduce the particle size below 5 μm followed by dilution with exipient to a 15 mg/mL concentration.

*Animals*

Wild-caught AGM (St. Kitts origin) were sourced through Worldwide Primates Inc. for all studies. Ages were unknown due to wild-caught sourcing but were estimated to range from three to ten years based on dentition. All protocols were reviewed by an Institutional Animal Care and Use Committee (IACUC) at LBRI. Research was conducted under an IACUC approved protocol in compliance with the Animal Welfare Act, PHS Policy, and other federal statutes and regulations relating to animals and experiments involving animals. The facilities where this research was conducted are accredited by the Association for Assessment and Accreditation of Laboratory Animal Care, International and strictly adhere to principles stated in the Guide for the Care and Use of Laboratory Animals, National Research Council, 2011 (National Academies Press, Washington, DC). All research methods and results are reported in compliance with the Animal Research: Reporting of In Vivo Experiments (ARRIVE) guidelines. AGM were housed in adjacent individual cages, in a climate-controlled room with a fixed light/dark cycle (12-h light/12-h dark). AGMs were monitored at least twice daily throughout the study. Commercial monkey chow, treats, and fruit were provided twice daily by trained personnel. Water was available ad libitum. Animals were observed a minimum of twice daily for any clinical signs of test article toxicity or RSV associated disease. A standard lexicon of pharmacologic and toxicologic observations was used for documentation of any abnormal clinical signs.

*Intratracheal Inhalation Dosing and Aerosol Characterization*

The setup shown in Figure S1 was used for aerosol generation and *in vivo* delivery. **1** was aerosolized using an Aeroneb Solo vibrating mesh nebulizer. Ventilation was controlled by a Harvard pump, and settings were standardized at a tidal volume of 12 mL/kg, 30 breaths per minute, and a 35:65 inspiratory to expiratory ratio. Animals were sedated with ketamine and anesthesia was maintained with isoflurane delivered in 100% oxygen through the breathing circuit. The aerosolized **1** was delivered to anesthetized animals through a cuffed endotracheal tube (4.5 mm inside diameter) over 30 min.

**Figure S1.** Schematic of the Anesthetized AGM IT Exposure System.

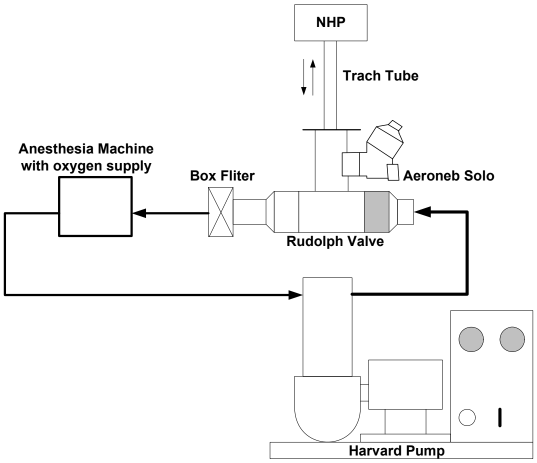


Estimation of dose for each animal was based on separate in vitro testing performed immediately prior to and after dosing the animals, during which the aerosolized **1** was collected on a pre-weighed glass fiber filter placed at the end of the ET tube connected to an identical exposure system. Filters collected during these tests were extracted using 4 mL of 0.1% trifluoroacetic acid in 1:1 water/acetonitrile. Filters were agitated for at least 30 min, and then diluted for analysis by UPLC-UV. The estimated delivered dose for each animal based on exposure duration is shown in Table S1, and the deposition fraction was assumed to be 100% for the lung deposited dose.

**Table S1.** Calculated IT Pulmaonry Delivered Doses of 1.

| Study | NHP Weight (kg) | Exposure Duration (min) | Average 1 collected on filter by chemical analysis (mg/min exposure) | Average Daily Delivered Dose (mg/kg) |
|---|---|---|---|---|
| IT Cyno PK Study | 4.32 | 30 | 0.72 | 5.0 |
| | 7.18 | 30 | 0.68 | 2.9 |
| | 5.24 | 30 | 0.67 | 3.8 |
| | **Average** | | | 3.9 |
| IT AGM PK Study | 4.53 | 30 | 0.776 | 5.1 |
| | 3.87 | 30 | 0.722 | 5.6 |
| | 3.80 | 30 | 0.876 | 6.9 |
| | **Average** | | | 5.9 |
| RSV AGM Efficacy Study (Cohort 1) | 4.96 | 16 | 0.53 | 1.6 |
| | 3.97 | 12 | 0.54 | 1.6 |
| | 3.41 | 11 | 0.50 | 1.6 |
| | **Average** | | | 1.6 |
| RSV AGM Efficacy Study (Cohort 2) | 5.80 | 30 | 1.16 | 6.0 |
| | 5.55 | 30 | 0.999 | 5.4 |
| | 3.90 | 30 | 1.14 | 8.8 |
| | 2.79 | 30 | 0.949 | 10.2 |
| | **Average** | | | 7.6 |

Particle size distribution of the aerosol was determined at the end of the endotracheal tube by APS (Model 3321, Aerodynamic Particle Sizer, TSI Inc., Shoreview, MN) in conjunction with a TSI Diluter (Model 3302A, TSI, Inc.). For these tests, aerosolization and ventilation were performed at the same nominal conditions as Estimation of Dose. The average mass median aerodynamic diameter of the **1** aerosol was 1.34 µm ± 1.81 µm (0.1% Tween-20 formulation) and 1.55 µm ± 1.65 µm (0.1% HPMC formulation).

*RSV Infections*

RSV strain A2 was sourced from Virapur (San Diego, CA). Virus was thawed and diluted to $2 \times 10^6$ $TCID_{50}$ / mL immediately prior to RSV infections. Each animal received a 2 mL total inoculum (1 mL delivered intratracheal and 1 mL delivered intranasal) for a total instillation of $4 \times 10^6$ $TCID_{50}$ RSV. AGMs were anesthetized with ketamine and isoflurane for RSV infections. For IT administration, 1 mL of the viral inoculum was administered through polyethylene (PE) tubing advanced into the trachea using a bronchoscope and deposited approximately half-way between the larynx and the carina. Approximately 1 mL sterile saline was used to flush the PE tubing followed by approximately 2 mL air. For IN administration, 0.5 mL of the viral inoculum was administered dropwise into each nostril.

*Sample Collection and Processing*

For the RSV efficacy studies, nasal swabs and BALF samples were collected from anesthetized animals at 1, 3, 5, 7, 9, 11, 13 and 15 d post-infection. On days that coincided with **1** dosing, samples were collected immediately prior to IT exposures. Nasal swabs were collected using a cotton-tipped applicator presoaked in sterile saline. Swabs were placed in a tube containing 0.5 mL sterile saline, frozen immediately on dry ice, and stored at −70 °C or below until analysis.

BALF samples from both the left and right caudal lung lobes were collected and processed as previously described.[10]

*RNA Isolation*

Real-Time Quantitative PCR (RT-qPCR) was performed on RNA extracted from BALF supernatant and nasal and throat swabs collected from each animal at each timepoint. The RNA from BALF sample mixed with lysis buffer immediately after BAL collection was extracted using ThermoFisher's MagMAX Pathogen RNA isolation kit and performed according to manufacturer instructions, including the addition of the exogenous RNA extraction control. The extraction buffer used for sample extraction was spiked with the equivalent of 20,000 copies of Xeno RNA/sample, and the combined extraction buffer + Xeno RNA was mixed with the BALF sample at the time of collection. This Internal Positive Control (IPC) RNA served as an extraction efficiency and matrix inhibition control to account for loss or inhibition of target RNA during downstream RT-qPCR analysis. A viral extraction control was included in each RNA isolation. A fixed volume of serially diluted stock virus (10 µL) was added to 290 µL of extraction buffer and was processed the same as BALF samples to act as a positive isolation control. The isolated RNA was eluted, RNA concentration was determined by nanodrop, and samples were stored at -80ºC until PCR amplification.

*RSV RT-qPCR*

Relative viral burden was measured by RT-qPCR for the RSV N gene from swabs, as well as from BALF supernatant from each animal at each timepoint. Copies per mL equivalents were calculated from a standard curve generated from RSV DNA plasmid stocks of known copy concentration. A set of RSV DNA plasmid serial dilutions to generate standard curve were added for each RT-qPCR run. Relative RSV N gene concentrations for each sample were presented as copies/mL equivalents based on the stanrdard curve run with each reaction. In addition, infectious units per mL (IU/mL) equivalents were calculated from a standard curve generated from RSV stocks of known titer that were spiked into appropriate amount of lysis buffer and extracted in parallel with the BALF samples. All samples were run in duplicate.
A fixed volume of RNA (5 µL) was added to a reaction mix of 1x TaqMan® Fast Virus 1-Step Master Mix, 1x RSV N gene Primer Probe Mix (containing 900nM of primers and 250nM of probe) and 1x Xeno™ VIC™ Primer Probe Mix for a final volume of 20µL. To control for Xeno RNA IPC amplification efficiency, 5,000 copies of Xeno RNA (5 µL) were added to a separate reaction mix which were run in parallel to test samples. The RSV N gene primers and probe sequences are:

RSV A2 F: 5' GCTCTTAGCAAAGTCAAGTTGAATGA 3'
RSV A2 R: 5' TGCTCCGTTGGATGGTGTATT 3'
RSV A2 Probe: 6FAM-ACACTCAACAAAGATCAACTTCTGTCATCCAGC-MGBNFQ

Amplification and detection were performed using an ABI 7900HT Thermal Cycler under the following cycling conditions: 50ºC for 5 min, 95ºC for 20 seconds and 40 cycles of 95ºC for 15 seconds, 60ºC for 60 seconds.

*Statistical Analysis*

Statistical analyses were performed using GraphPad Prism software. For longitudinal analyses of viral load in BALF and nasal swabs, **1** treatment was compared with its respective vehicle treatment by repeated-measures two-way ANOVA with Bonferroni post-hoc correction for multiple comparisons. For BALF samples, the viral load in the right and left lung lobes were averaged such that a single value is represented for each animal. Samples that were below the assay lower limit of quantification were assigned a value of 1/2 the lower limit of quantification before log transformation. For all analyses, a corrected $p$ value $< 0.05$ was considered statistically significant.

#### SMALL MOLECULE X-RAY STRUCTURE OF COMPOUND 1

The X-ray crystallographic coordinates and structure factor file for compound **1** has been deposited in the Cambridge Structural Database (accession number 2348357, http://www.ccdc.cam.ac.uk/).

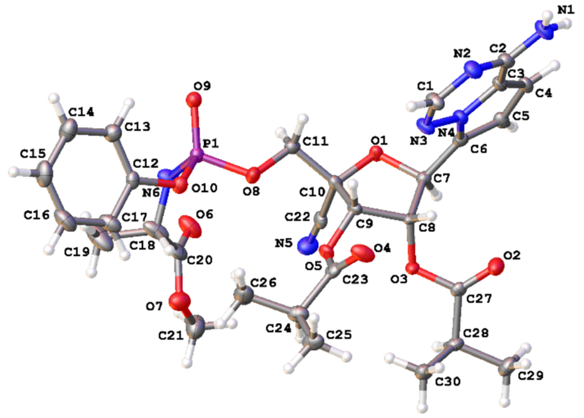


**Crystal data and structure refinement for 1**.

| | | |
|---|---|---|
| Empirical formula | $C_{30}H_{37}N_6O_{10}P$ | |
| Formula weight | 672.62 | |
| Temperature | 100.0 K | |
| Wavelength | 0.71073 Å | |
| Crystal system | Trigonal | |
| Space group | $P3_2$ | |
| Unit cell dimensions | a = 9.381(3) Å | α= 90° |
| | b = 9.381(3) Å | β= 90° |
| | c = 33.188(10) Å | γ = 120° |
| Volume | 2529.3(17) Å$^3$ | |
| Z | 3 | |
| Density (calculated) | 1.325 Mg/m$^3$ | |
| Absorption coefficient | 0.145 mm$^{-1}$ | |
| F(000) | 1062 | |
| Crystal size | 0.32 x 0.3 x 0.28 mm$^3$ | |
| Theta range for data collection | 1.841 to 27.246°. | |
| Index ranges | -12<=h<=9, -12<=k<=12, -42<=l<=42 | |
| Reflections collected | 23593 | |
| Independent reflections | 7210 [R(int) = 0.0613] | |
| Completeness to theta = 25.242° | 100.0 % | |
| Absorption correction | Semi-empirical from equivalents | |
| Max. and min. transmission | 0.2611 and 0.2223 | |
| Refinement method | Full-matrix least-squares on F$^2$ | |
| Data / restraints / parameters | 7210 / 1 / 434 | |
| Goodness-of-fit on F$^2$ | 1.043 | |
| Final R indices [I>2sigma(I)] | R1 = 0.0475, wR2 = 0.0990 | |
| R indices (all data) | R1 = 0.0593, wR2 = 0.1059 | |
| Absolute structure parameter | -0.02(7) [abs. stereochem. determined] | |
| Extinction coefficient | n/a | |

Largest diff. peak and hole 0.451 and -0.353 e.Å$^{-3}$

#### HPLC AND NMR SPECTRAL DATA

##### COMPOUND 20 HPLC AND NMR

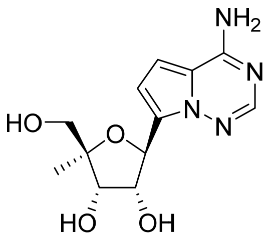


Method Info : Sample Bank Method - 2-98%B with 8.5 min gradient, A=Water + 0.1% TFA, B= Acetonitrile + 0.1% TFA; 1.5 mL/min; Column: Phenomenex Kinetex C18, 2.6u 100A, 4.6 x 100 mm; Instrument 1290II

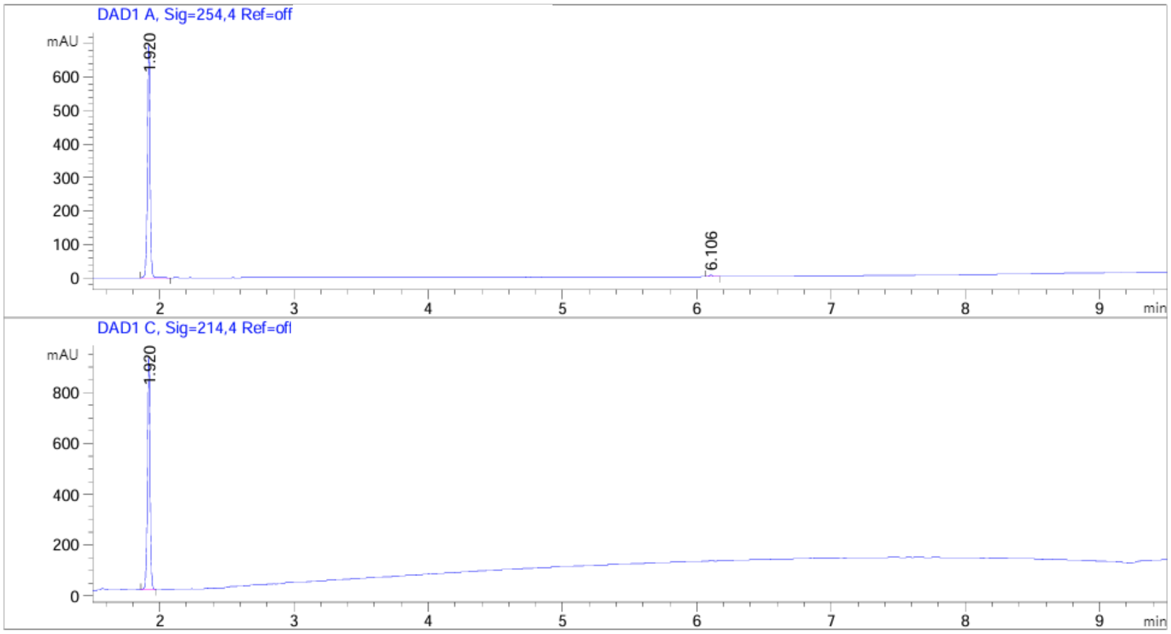


| Peak # | RetTime [min] | Type | Width [min] | Area [mAU*s] | Height [mAU] | Area % |
|---|---|---|---|---|---|---|
| 1 | 1.920 | BB | 0.0202 | 936.54102 | 697.06189 | 99.1791 |
| 2 | 6.106 | BB | 0.0237 | 7.75160 | 4.96347 | 0.8209 |

Totals : 944.29262 702.02536

Signal 2: DAD1 C, Sig=214,4 Ref=off

| Peak # | RetTime [min] | Type | Width [min] | Area [mAU*s] | Height [mAU] | Area % |
|---|---|---|---|---|---|---|
| 1 | 1.920 | BB | 0.0200 | 1219.02466 | 915.82794 | 100.0000 |

Totals : 1219.02466 915.82794

1H NMR (400 MHz, D2O)

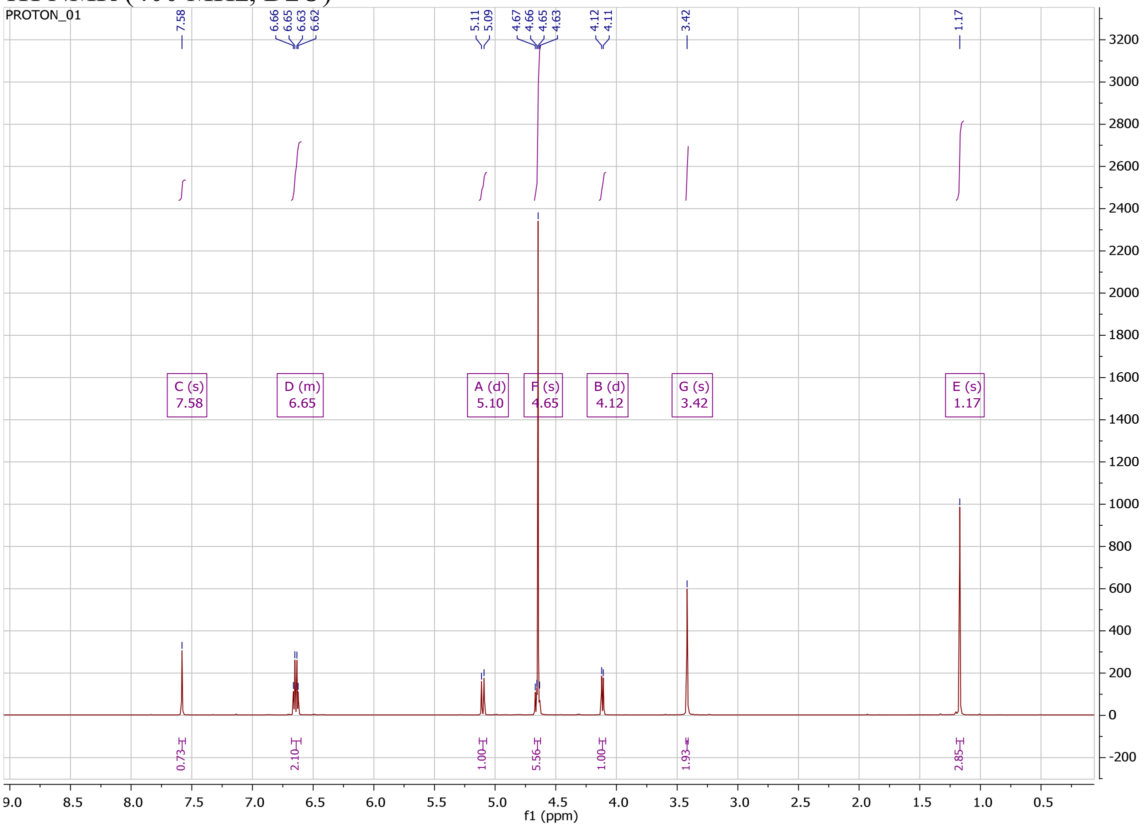

#### COMPOUND 21 HPLC AND NMR

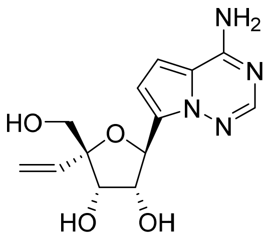

Method Info : Sample Bank Method - 2-98%B with 8.5 min gradient, A=Water + 0.1% TFA, B= Acetonitrile + 0.1% TFA; 1.5 mL/min; Column: Phenomenex Kinetex C18, 2.6u 100A, 4.6 x 100 mm; Instrument 1290II

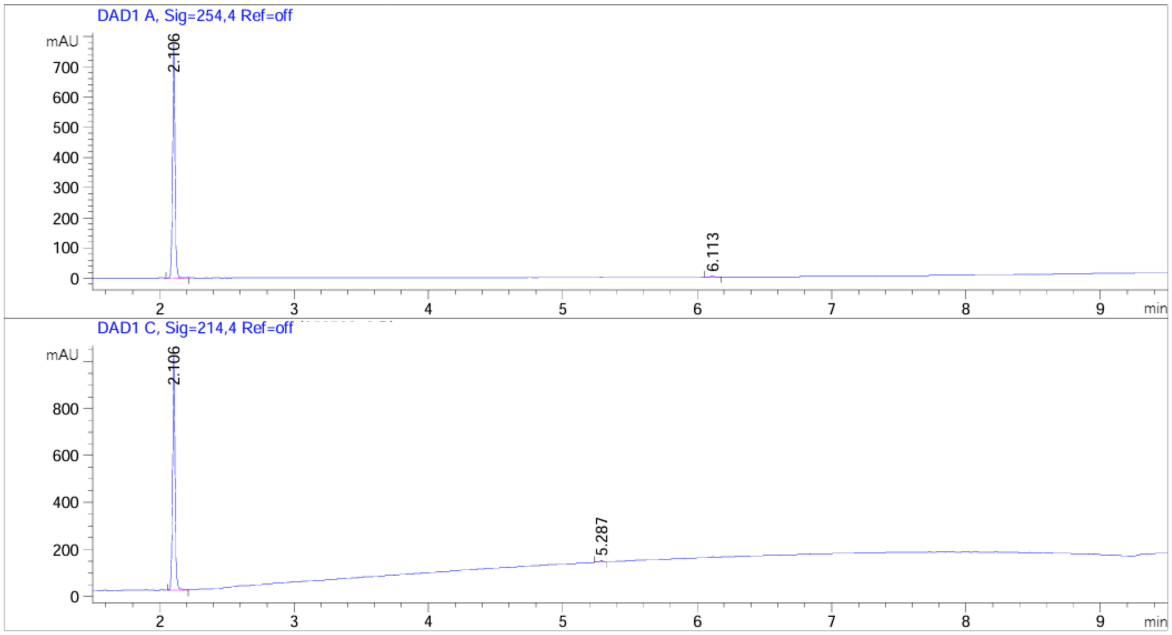


| Peak # | RetTime [min] | Type | Width [min] | Area [mAU*s] | Height [mAU] | Area % |
|---|---|---|---|---|---|---|
| 1 | 2.106 | BB | 0.0188 | 986.80505 | 776.66522 | 99.4111 |
| 2 | 6.113 | BB | 0.0296 | 5.84548 | 2.89469 | 0.5889 |
| Totals : | | | | 992.65053 | 779.55991 | |

Signal 2: DAD1 C, Sig=214,4 Ref=off

| Peak # | RetTime [min] | Type | Width [min] | Area [mAU*s] | Height [mAU] | Area % |
|---|---|---|---|---|---|---|
| 1 | 2.106 | BB | 0.0188 | 1268.41113 | 996.59094 | 99.2932 |
| 2 | 5.287 | BB | 0.0253 | 9.02949 | 5.30745 | 0.7068 |
| Totals : | | | | 1277.44063 | 1001.89839 | |

$^{1}H$ NMR (400 MHz, $CD_3CN$)

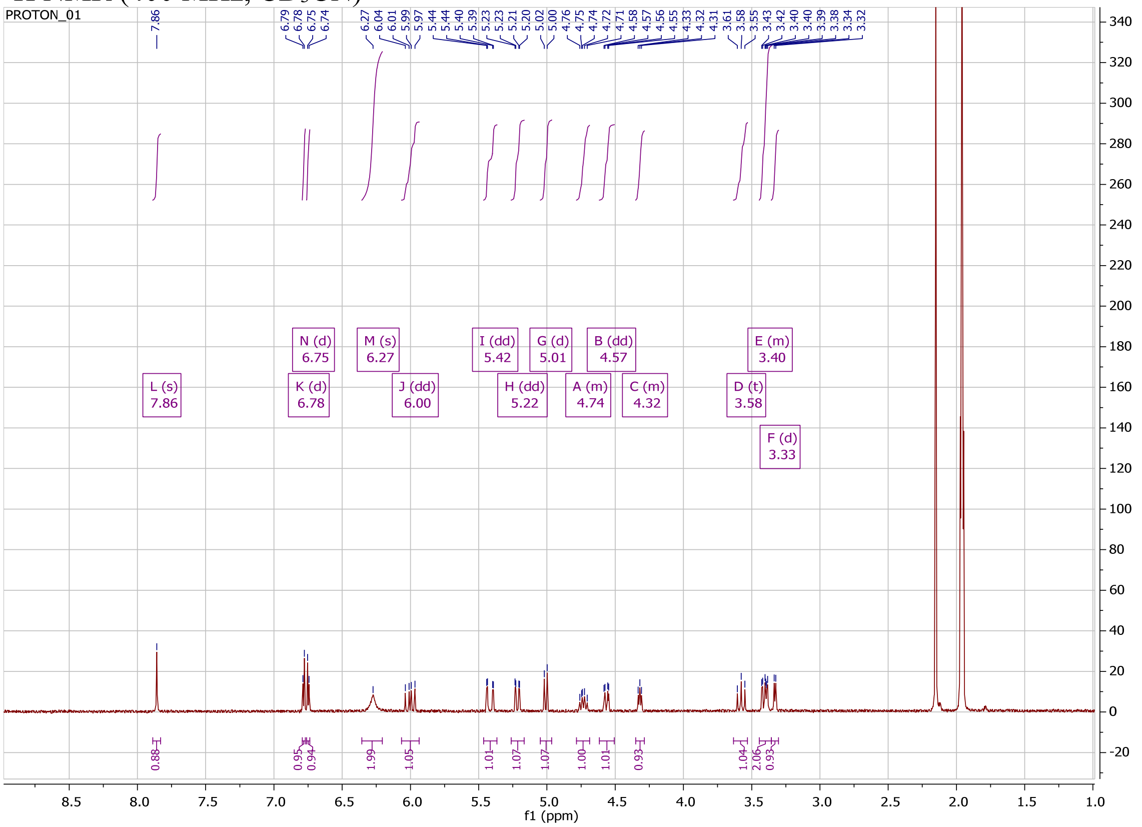


#### COMPOUND 2 HPLC AND NMR

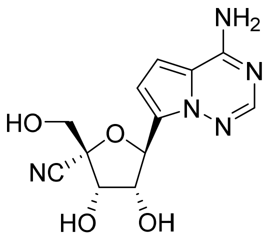


Method Info : Sample Bank Method - 2-98%B with 8.5 min gradient, A=Water + 0.1% TFA, B= Acetonitrile + 0.1% TFA; 1.5 mL/min; Column: Phenomenex Kinetex C18, 2.6u 100A, 4.6 x 100 mm; Instrument 1290II

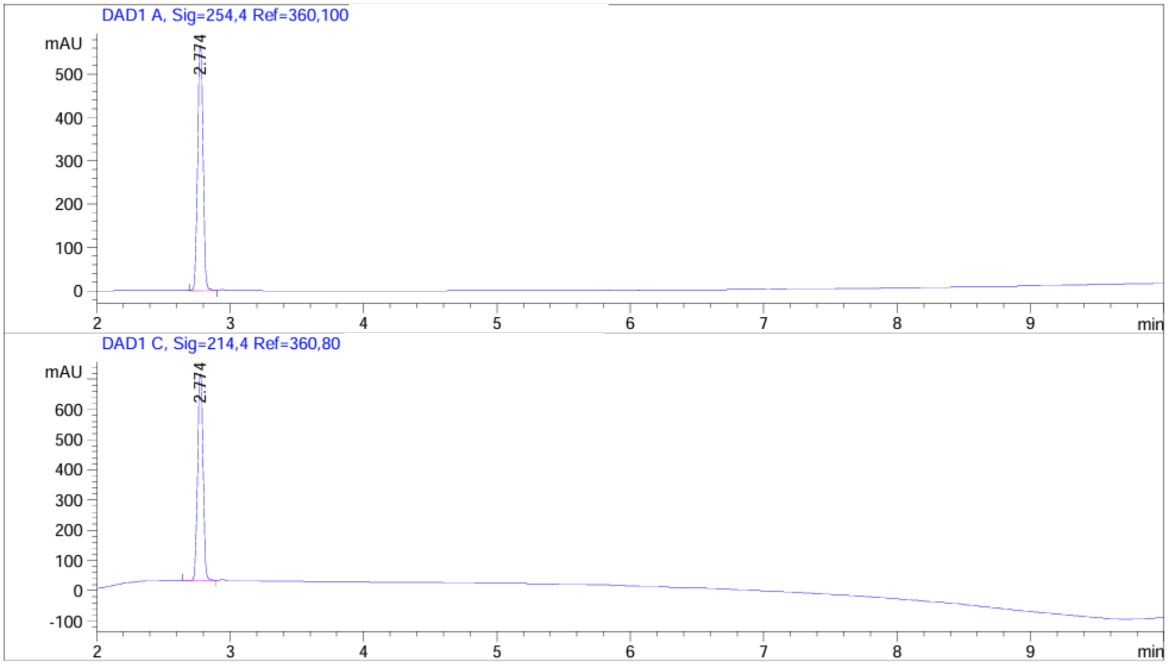


| Peak # | RetTime [min] | Type | Width [min] | Area [mAU*s] | Height [mAU] | Area % |
|---|---|---|---|---|---|---|
| 1 | 2.774 | VV | 0.0447 | 1600.53650 | 571.34607 | 100.0000 |

Totals : 1600.53650 571.34607

Signal 2: DAD1 C, Sig=214,4 Ref=360,80

| Peak # | RetTime [min] | Type | Width [min] | Area [mAU*s] | Height [mAU] | Area % |
|---|---|---|---|---|---|---|
| 1 | 2.774 | BB | 0.0446 | 1928.45337 | 689.29449 | 100.0000 |

Totals : 1928.45337 689.29449

[1]H NMR (400 MHz, $CD_3OD$)

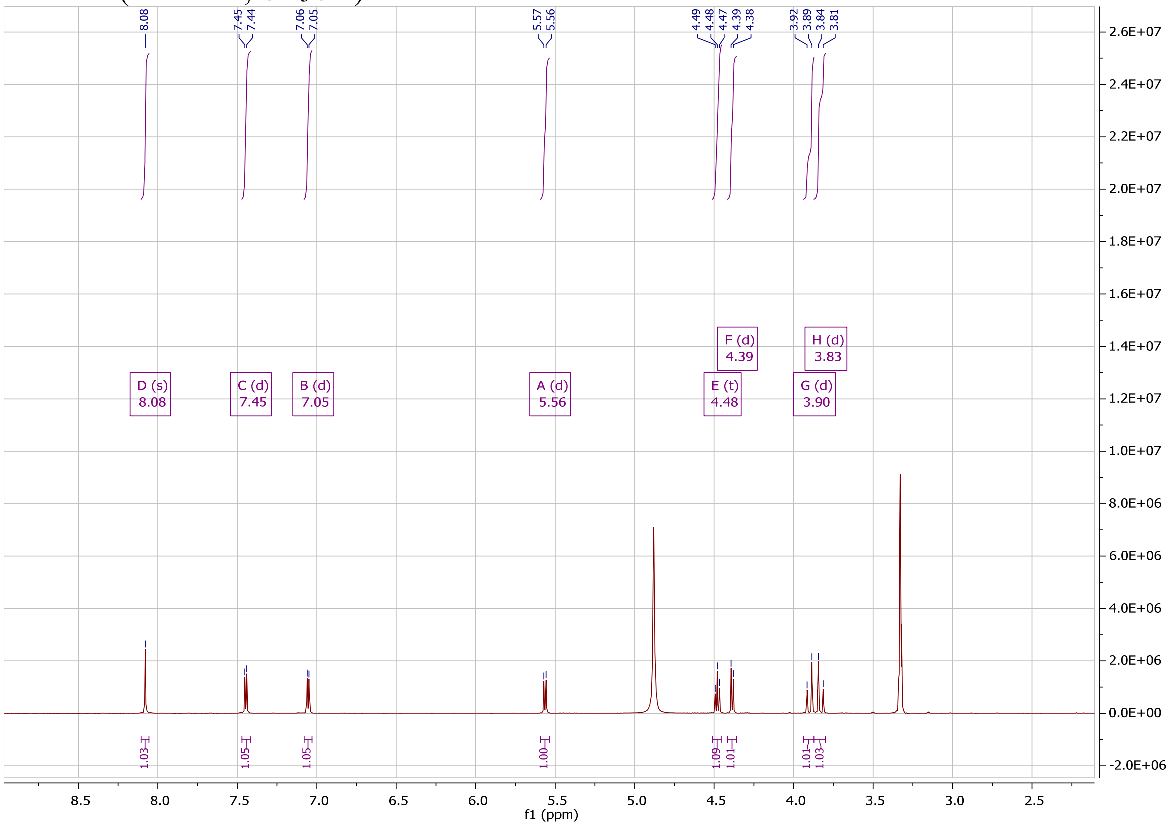


#### COMPOUND 22 HPLC AND NMR

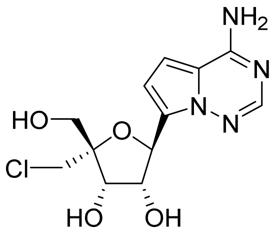


Method Info : Sample Bank Method - 2-98%B with 8.5 min gradient, A=Water + 0.1% TFA, B= Acetonitrile + 0.1% TFA; 1.5 mL/min; Column: Phenomenex Kinetex C18, 2.6u 100A, 4.6 x 100 mm; Instrument 1290II

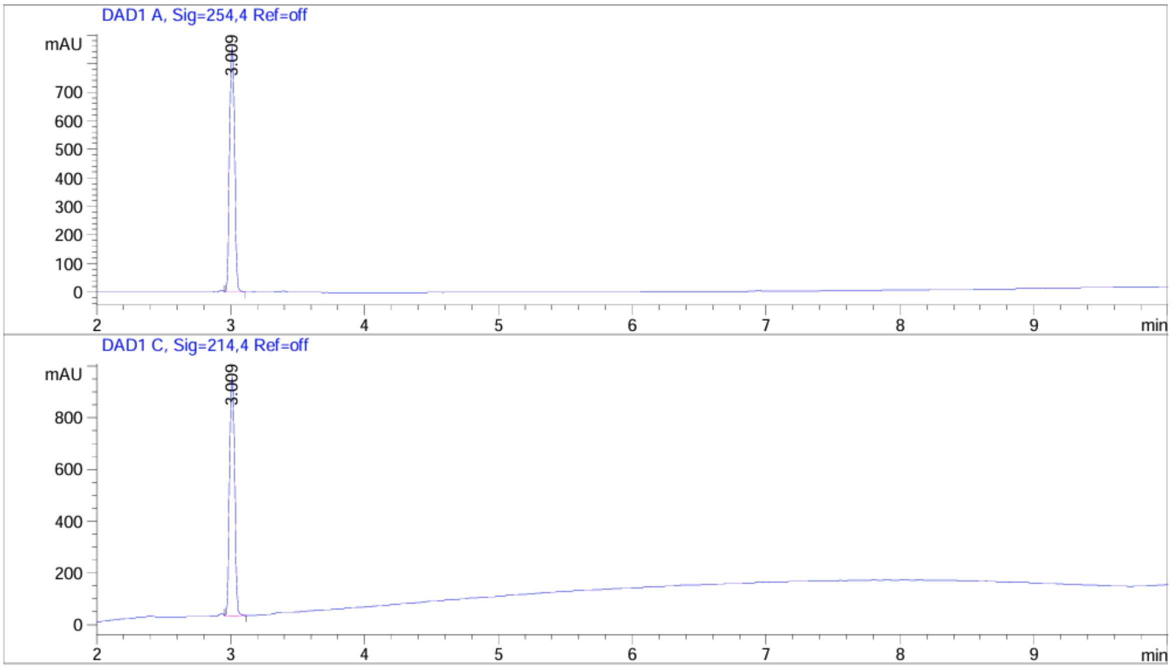


| Peak # | RetTime [min] | Type | Width [min] | Area [mAU*s] | Height [mAU] | Area % |
|---|---|---|---|---|---|---|
| 1 | 3.009 | VB | 0.0423 | 2241.82324 | 862.94659 | 100.0000 |
| Totals : | | | | 2241.82324 | 862.94659 | |

Signal 2: DAD1 C, Sig=214,4 Ref=off

| Peak # | RetTime [min] | Type | Width [min] | Area [mAU*s] | Height [mAU] | Area % |
|---|---|---|---|---|---|---|
| 1 | 3.009 | VB | 0.0425 | 2434.58618 | 930.50122 | 100.0000 |
| Totals : | | | | 2434.58618 | 930.50122 | |

$^1H$ NMR (400 MHz, $D_2O$)

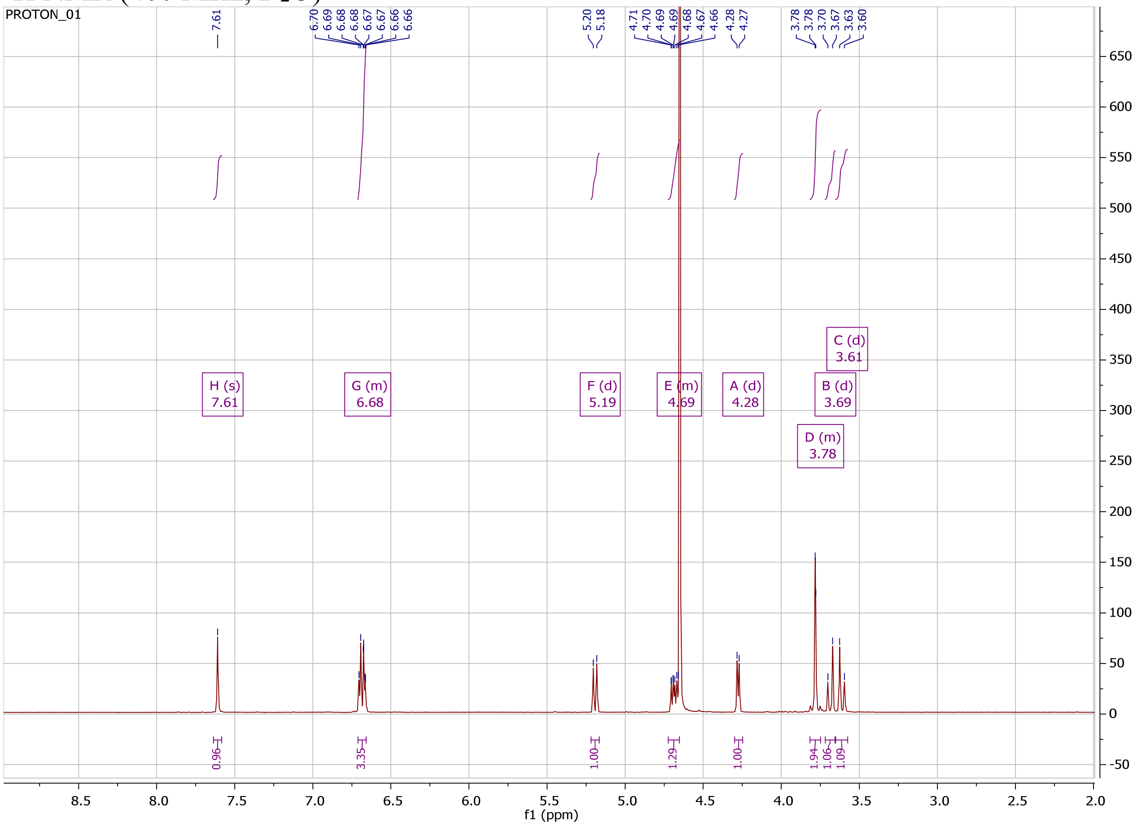


#### COMPOUND 26 HPLC AND NMR

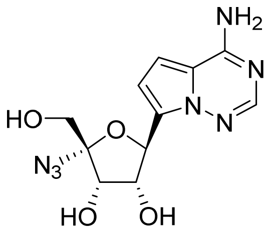


Method Info : Sample Bank Method - 2-98%B with 8.5 min gradient, A=Water + 0.1% TFA, B= Acetonitrile + 0.1% TFA; 1.5 mL/min; Column: Phenomenex Kinetex C18, 2.6u 100A, 4.6 x 100 mm; Instrument 1290II

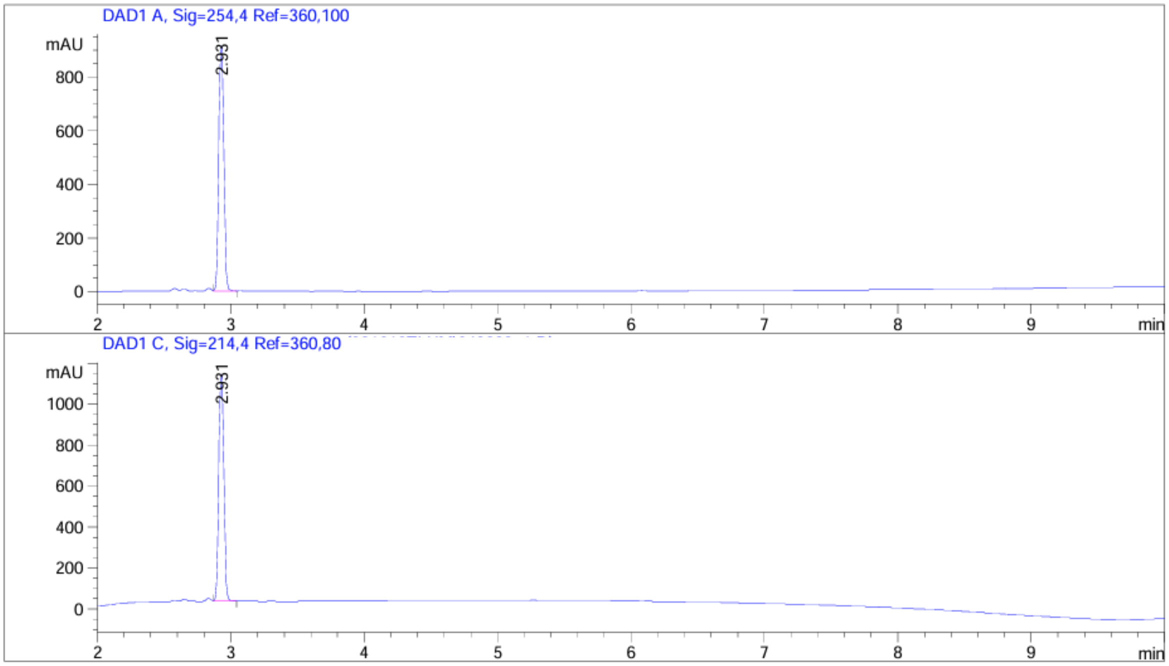


| Peak # | RetTime [min] | Type | Width [min] | Area [mAU*s] | Height [mAU] | Area % |
|---|---|---|---|---|---|---|
| 1 | 2.931 | VV | 0.0427 | 2429.00488 | 922.62347 | 100.0000 |
| Totals : | | | | 2429.00488 | 922.62347 | |

Signal 2: DAD1 C, Sig=214,4 Ref=360,80

| Peak # | RetTime [min] | Type | Width [min] | Area [mAU*s] | Height [mAU] | Area % |
|---|---|---|---|---|---|---|
| 1 | 2.931 | VB | 0.0427 | 2937.01587 | 1117.48340 | 100.0000 |
| Totals : | | | | 2937.01587 | 1117.48340 | |

[1]H NMR (400 MHz, $CD_3OD$)

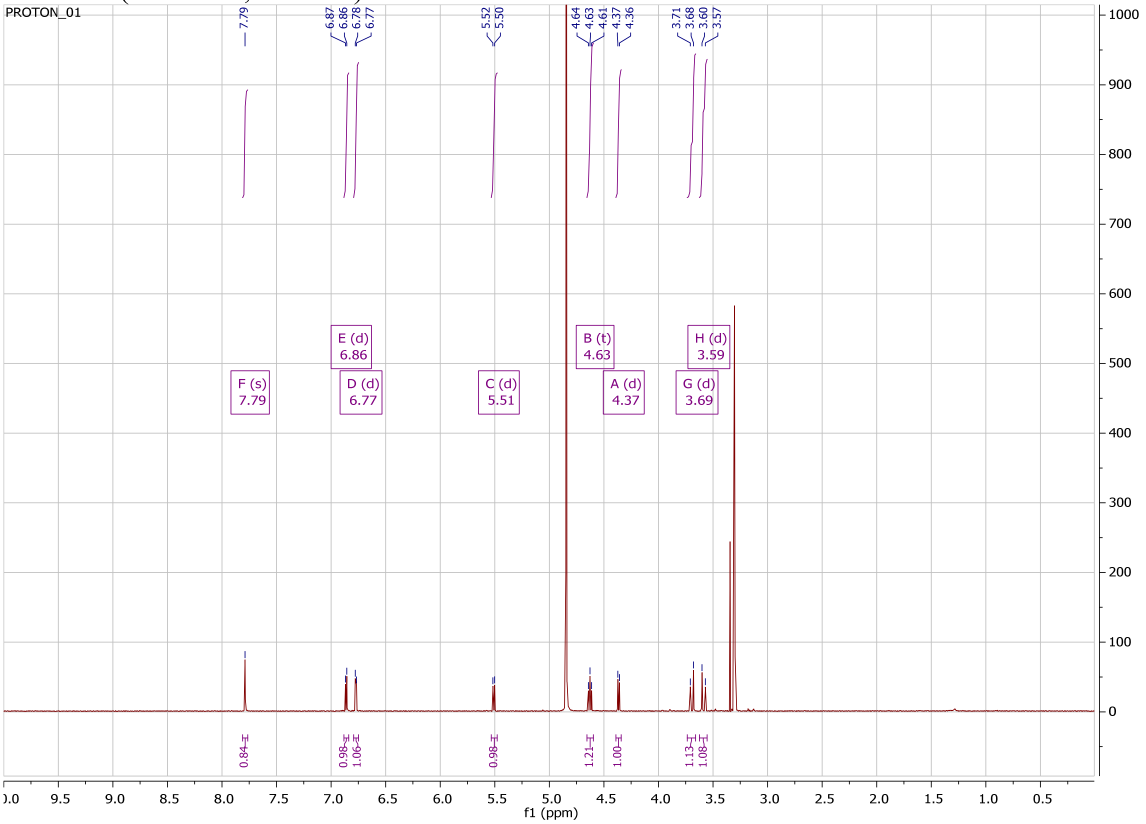


#### COMPOUND 28A HPLC AND NMR

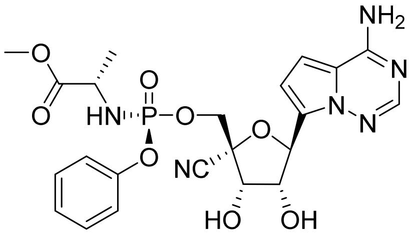

Method Info : Sample Bank Method - 2-98%B with 8.5 min gradient, A=Water + 0.1% TFA, B= Acetonitrile + 0.1% TFA; 1.5 mL/min; Column: Phenomenex Kinetex C18, 2.6u 100A, 4.6 x 100 mm; Instrument 1290II

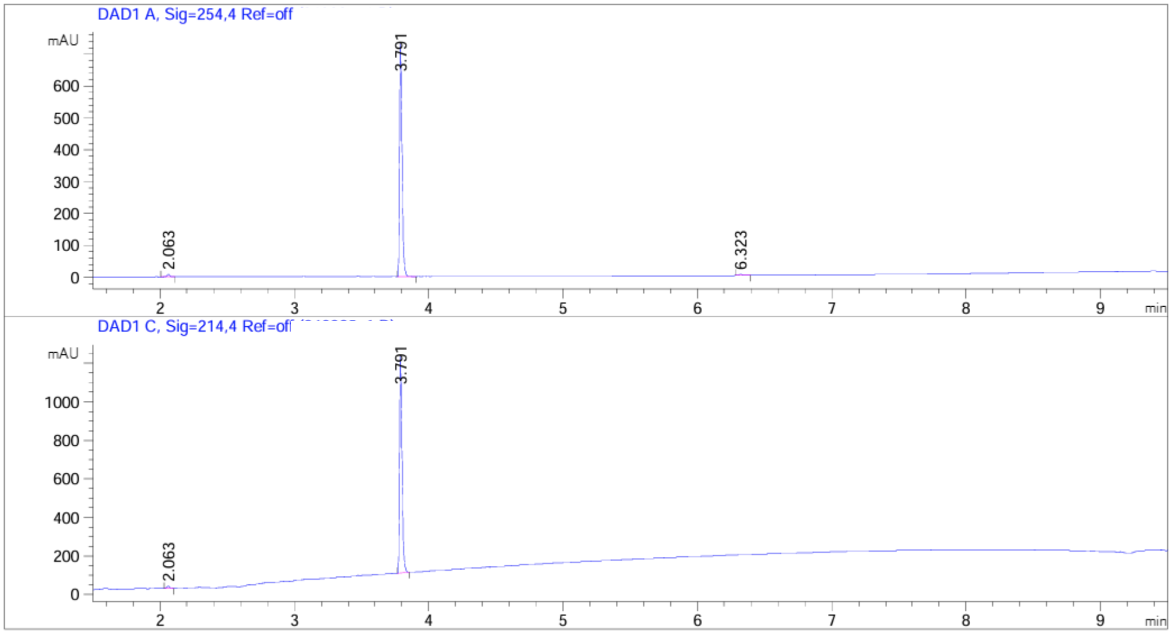


| Peak # | RetTime [min] | Type | Width [min] | Area [mAU*s] | Height [mAU] | Area % |
|---|---|---|---|---|---|---|
| 1 | 2.063 | BB | 0.0199 | 12.12666 | 9.18861 | 1.2562 |
| 2 | 3.791 | VB | 0.0201 | 947.10504 | 730.75836 | 98.1097 |
| 3 | 6.323 | BB | 0.0252 | 6.12151 | 3.63215 | 0.6341 |
| Totals : | | | | 965.35321 | 743.57912 | |

Signal 2: DAD1 C, Sig=214,4 Ref=off

| Peak # | RetTime [min] | Type | Width [min] | Area [mAU*s] | Height [mAU] | Area % |
|---|---|---|---|---|---|---|
| 1 | 2.063 | BB | 0.0210 | 17.78373 | 12.58752 | 1.2138 |
| 2 | 3.791 | BB | 0.0200 | 1447.37439 | 1124.32922 | 98.7862 |
| Totals : | | | | 1465.15812 | 1136.91674 | |

$^{1}H$ NMR (400 MHz, $CD_3OD$)

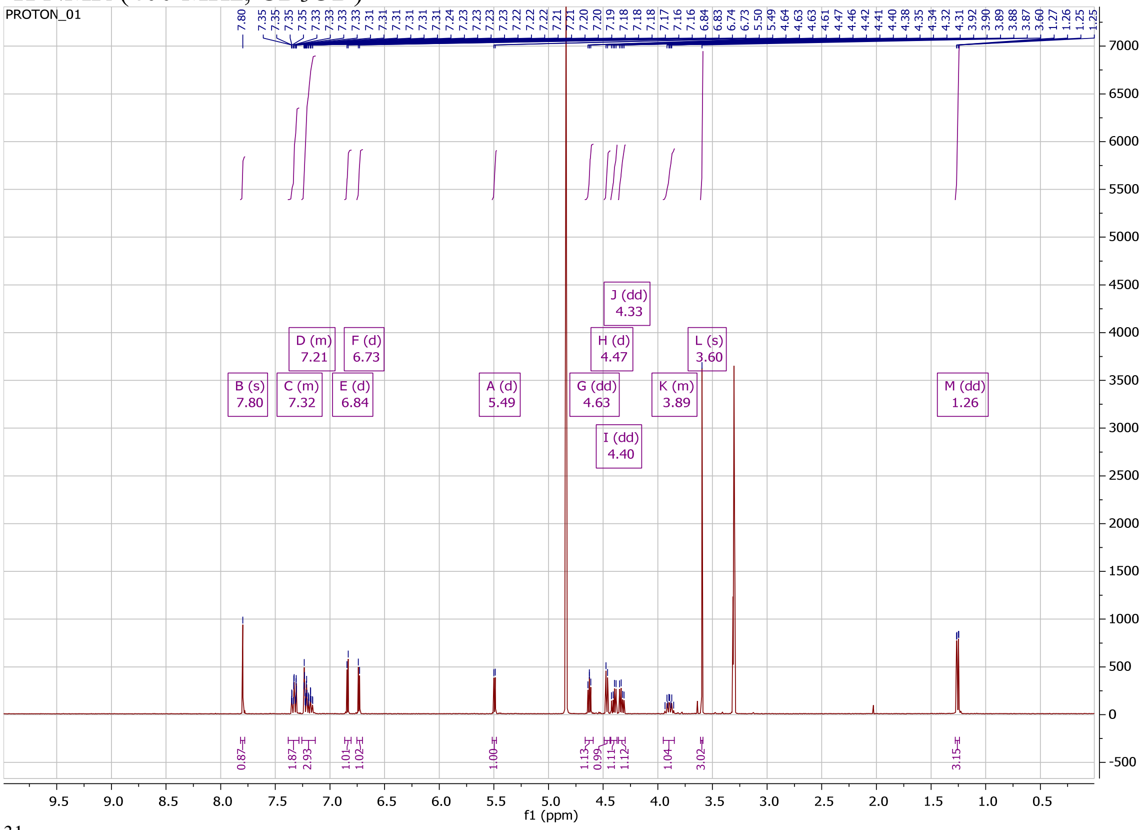

$^{31}P$ NMR (162 MHz, $CD_3OD$)

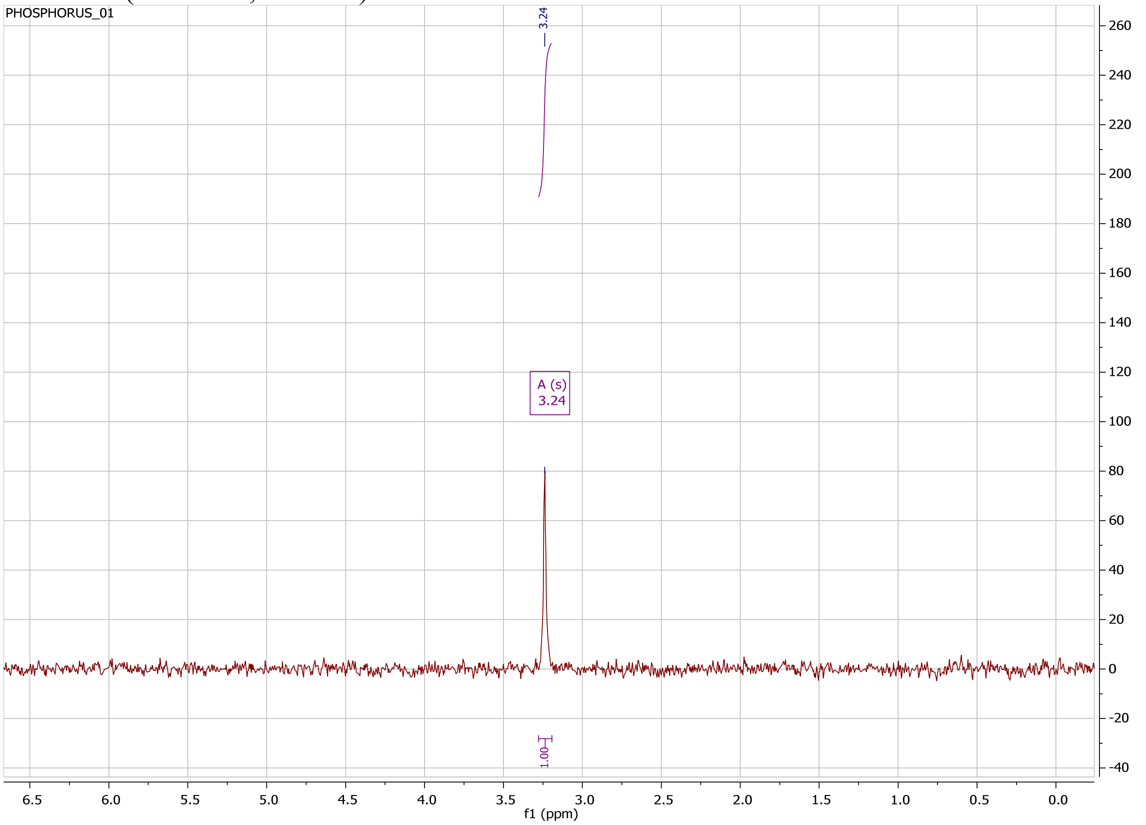

#### COMPOUND 28B HPLC

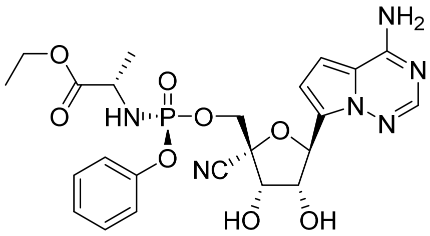

Method Info : Sample Bank Method - 2-98%B with 8.5 min gradient, A=Water + 0.1% TFA, B= Acetonitrile + 0.1% TFA; 1.5 mL/min; Column: Phenomenex Kinetex C18, 2.6u 100A, 4.6 x 100 mm; Instrument 1290II

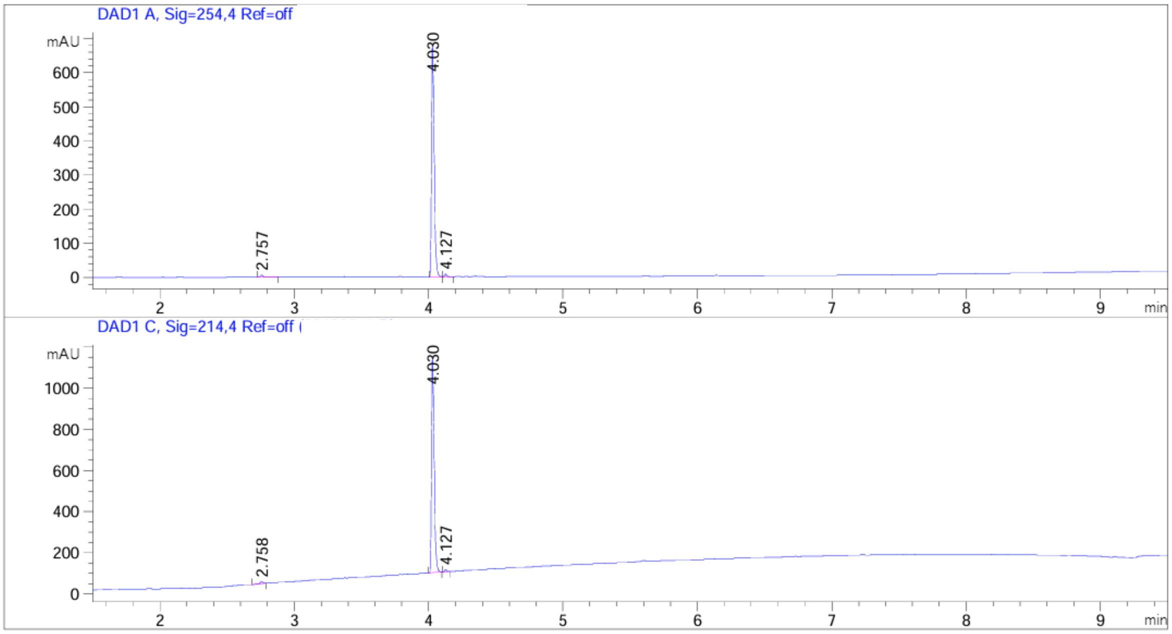


| Peak # | RetTime [min] | Type | Width [min] | Area [mAU*s] | Height [mAU] | Area % |
|---|---|---|---|---|---|---|
| 1 | 2.757 | BB | 0.0202 | 8.23746 | 6.10624 | 0.8702 |
| 2 | 4.030 | BV | 0.0209 | 929.64349 | 683.37170 | 98.2104 |
| 3 | 4.127 | VB | 0.0173 | 8.70258 | 7.64878 | 0.9194 |
| Totals : | | | | 946.58354 | 697.12672 | |

Signal 2: DAD1 C, Sig=214,4 Ref=off

| Peak # | RetTime [min] | Type | Width [min] | Area [mAU*s] | Height [mAU] | Area % |
|---|---|---|---|---|---|---|
| 1 | 2.758 | BB | 0.0204 | 13.79172 | 10.11230 | 0.9544 |
| 2 | 4.030 | BB | 0.0207 | 1419.93494 | 1052.35889 | 98.2579 |
| 3 | 4.127 | BB | 0.0162 | 11.38425 | 10.88701 | 0.7878 |
| Totals : | | | | 1445.11091 | 1073.35820 | |

#### COMPOUND 28C HPLC AND NMR

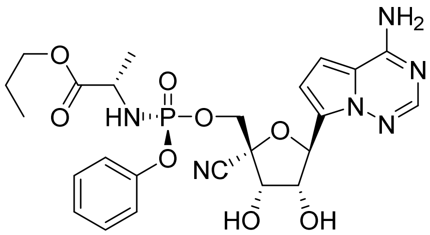

Method Info : Sample Bank Method - 2-98%B with 8.5 min gradient, A=Water + 0.1% TFA, B= Acetonitrile + 0.1% TFA; 1.5 mL/min; Column: Phenomenex Kinetex C18, 2.6u 100A, 4.6 x 100 mm; Instrument 1290II

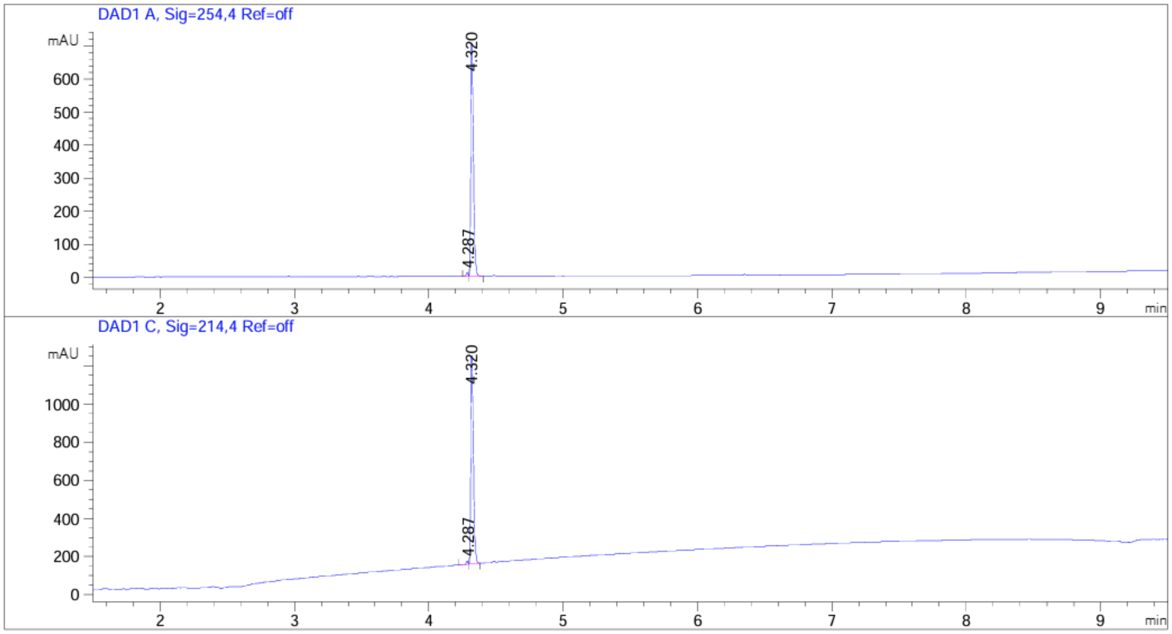


| Peak # | RetTime [min] | Type | Width [min] | Area [mAU*s] | Height [mAU] | Area % |
|---|---|---|---|---|---|---|
| 1 | 4.287 | BV | 0.0154 | 10.43566 | 10.69378 | 1.0763 |
| 2 | 4.320 | VB | 0.0209 | 959.11749 | 704.90320 | 98.9237 |

Totals : 969.55316 715.59698

Signal 2: DAD1 C, Sig=214,4 Ref=off

| Peak # | RetTime [min] | Type | Width [min] | Area [mAU*s] | Height [mAU] | Area % |
|---|---|---|---|---|---|---|
| 1 | 4.287 | BV | 0.0167 | 17.93153 | 16.41781 | 1.2026 |
| 2 | 4.320 | VB | 0.0208 | 1473.17334 | 1090.84985 | 98.7974 |

Totals : 1491.10487 1107.26766

$^{1}H$ NMR (400 MHz, $CD_3OD$)

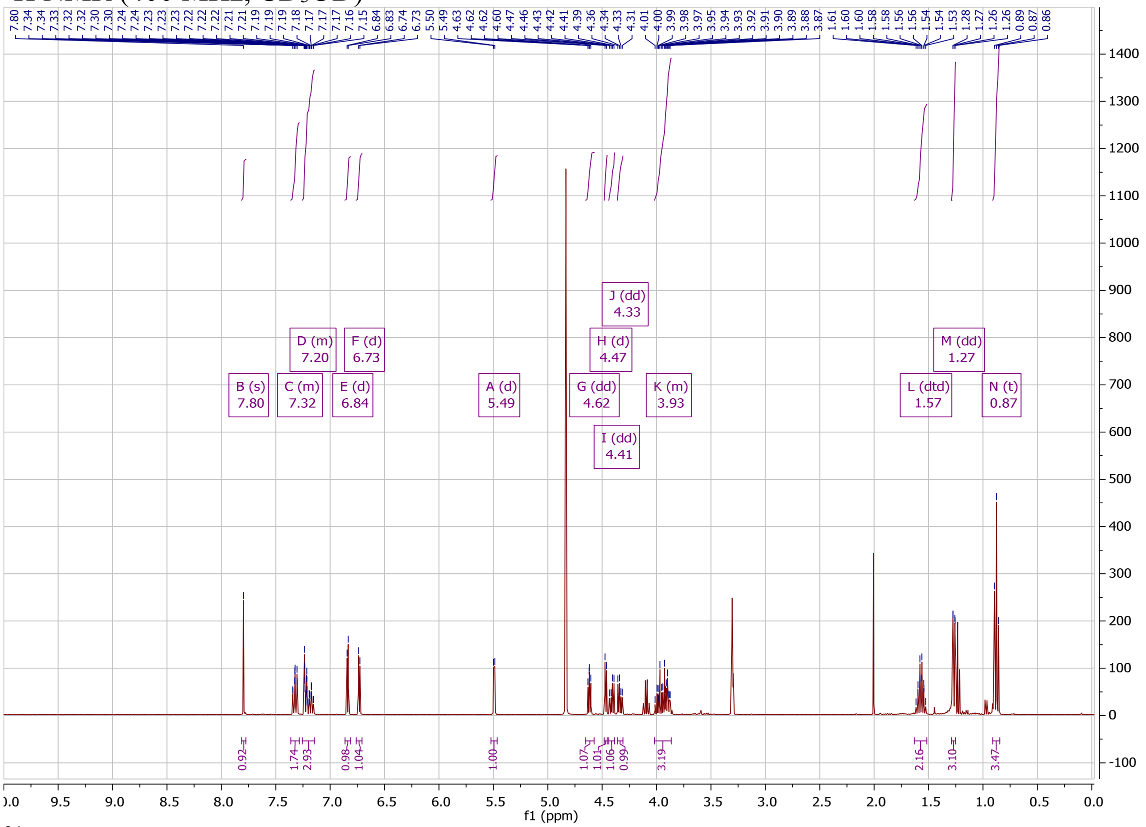

$^{31}P$ NMR (162 MHz, $CD_3OD$)

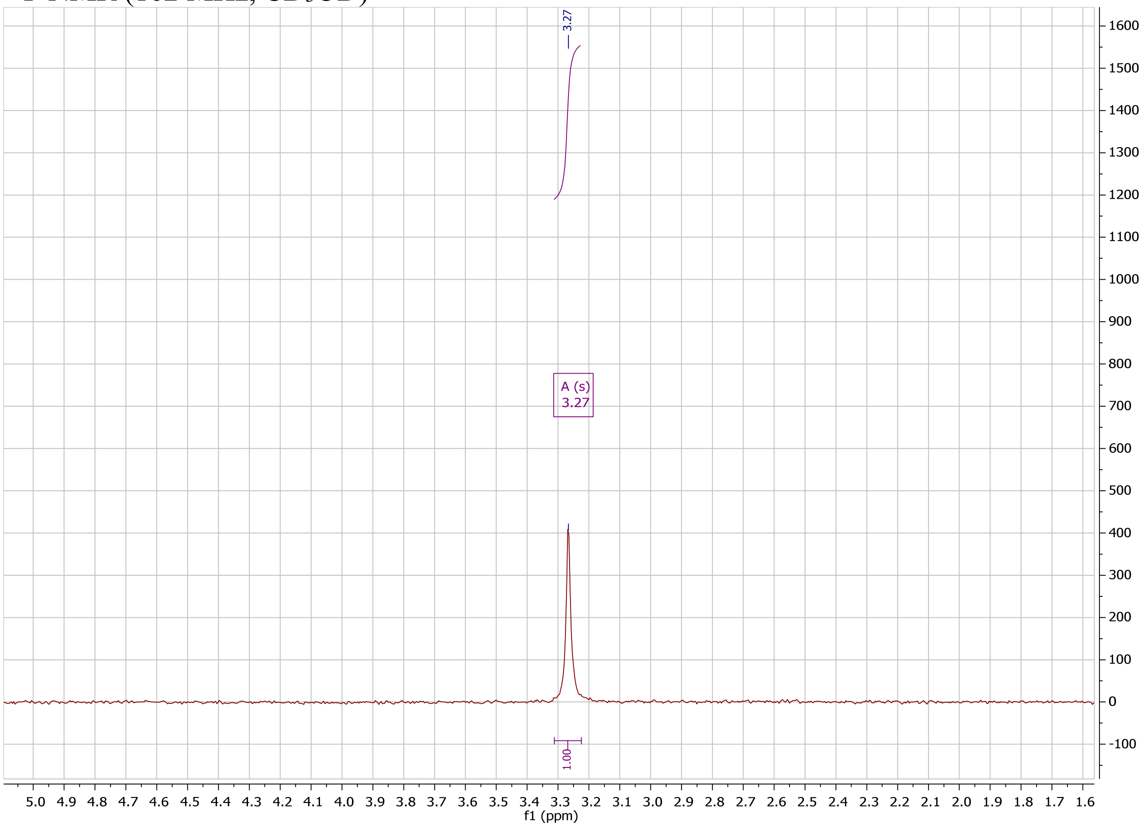

### COMPOUND 28D HPLC

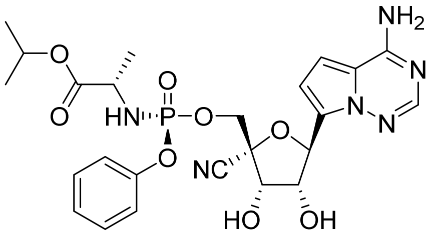

Method Info : Sample Bank Method - 2-98%B with 8.5 min gradient, A=Water + 0.1% TFA, B= Acetonitrile + 0.1% TFA; 1.5 mL/min; Column: Phenomenex Kinetex C18, 2.6u 100A, 4.6 x 100 mm; Instrument 1290II

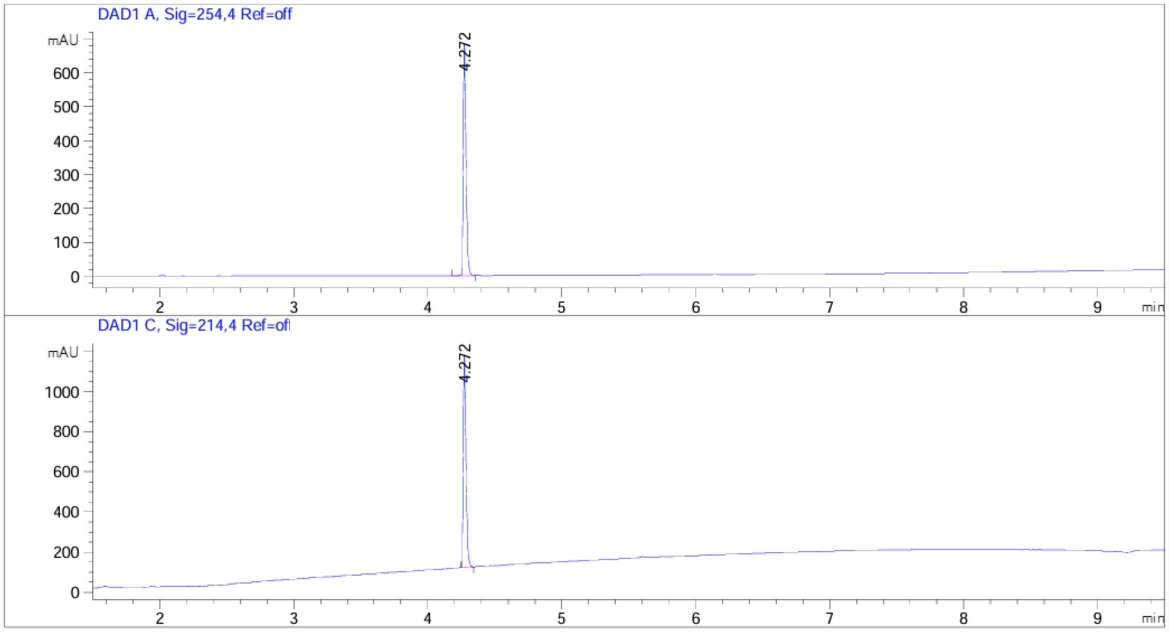


| Peak # | RetTime [min] | Type | Width [min] | Area [mAU*s] | Height [mAU] | Area % |
|---|---|---|---|---|---|---|
| 1 | 4.272 | BV | 0.0219 | 968.25305 | 686.94458 | 100.0000 |

Totals : 968.25305 686.94458

Signal 2: DAD1 C, Sig=214,4 Ref=off

| Peak # | RetTime [min] | Type | Width [min] | Area [mAU*s] | Height [mAU] | Area % |
|---|---|---|---|---|---|---|
| 1 | 4.272 | BB | 0.0218 | 1478.15076 | 1060.56506 | 100.0000 |

Totals : 1478.15076 1060.56506

#### COMPOUND 28E HPLC

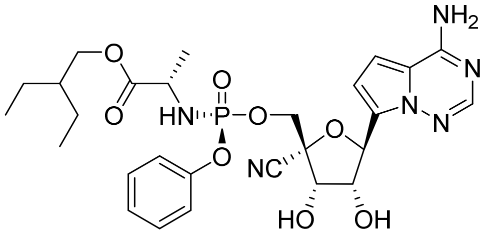

Method Info : Sample Bank Method - 2-98%B with 8.5 min gradient, A=Water + 0.1% TFA, B= Acetonitrile + 0.1% TFA; 1.5 mL/min; Column: Phenomenex Kinetex C18, 2.6u 100A, 4.6 x 100 mm; Instrument 1290II

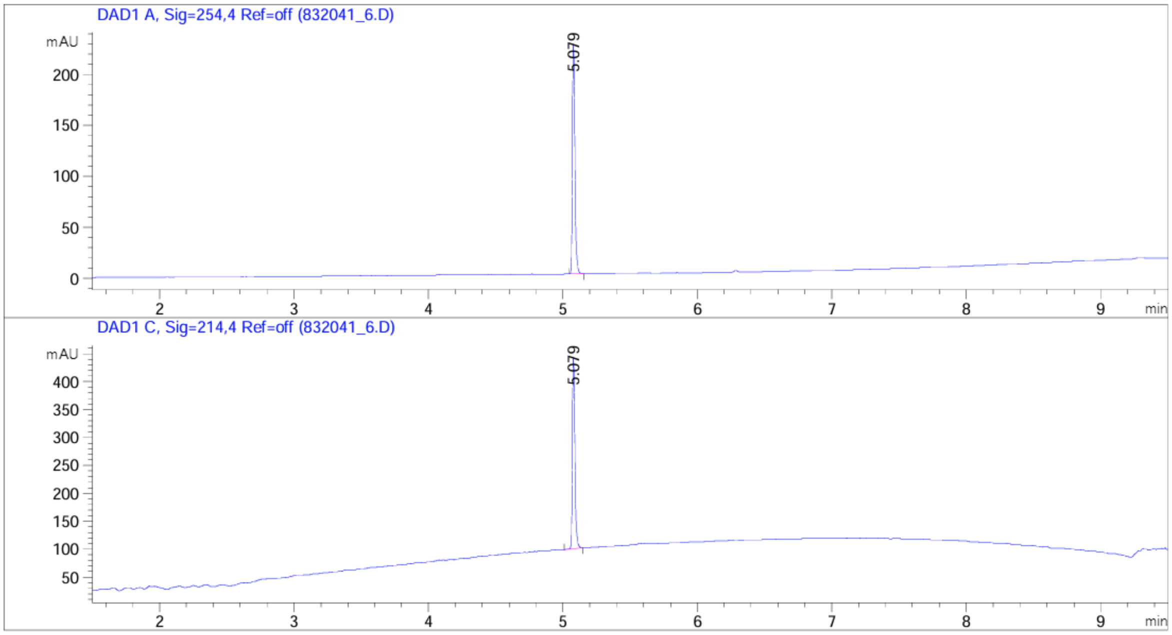


| Peak # | RetTime [min] | Type | Width [min] | Area [mAU*s] | Height [mAU] | Area % |
|---|---|---|---|---|---|---|
| 1 | 5.079 | BB | 0.0192 | 285.44534 | 226.35095 | 100.0000 |

Totals : 285.44534 226.35095

Signal 2: DAD1 C, Sig=214,4 Ref=off

| Peak # | RetTime [min] | Type | Width [min] | Area [mAU*s] | Height [mAU] | Area % |
|---|---|---|---|---|---|---|
| 1 | 5.079 | BB | 0.0192 | 433.49649 | 344.00388 | 100.0000 |

Totals : 433.49649 344.00388

#### COMPOUND 28F HPLC AND NMR

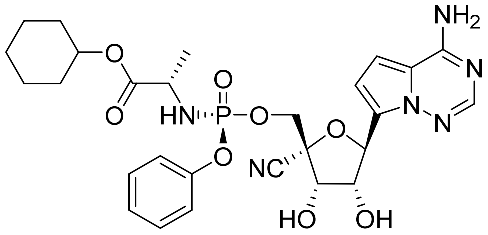

Method Info : Sample Bank Method - 2-98%B with 8.5 min gradient, A=Water + 0.1% TFA, B= Acetonitrile + 0.1% TFA; 1.5 mL/min; Column: Phenomenex Kinetex C18, 2.6u 100A, 4.6 x 100 mm; Instrument 1290II

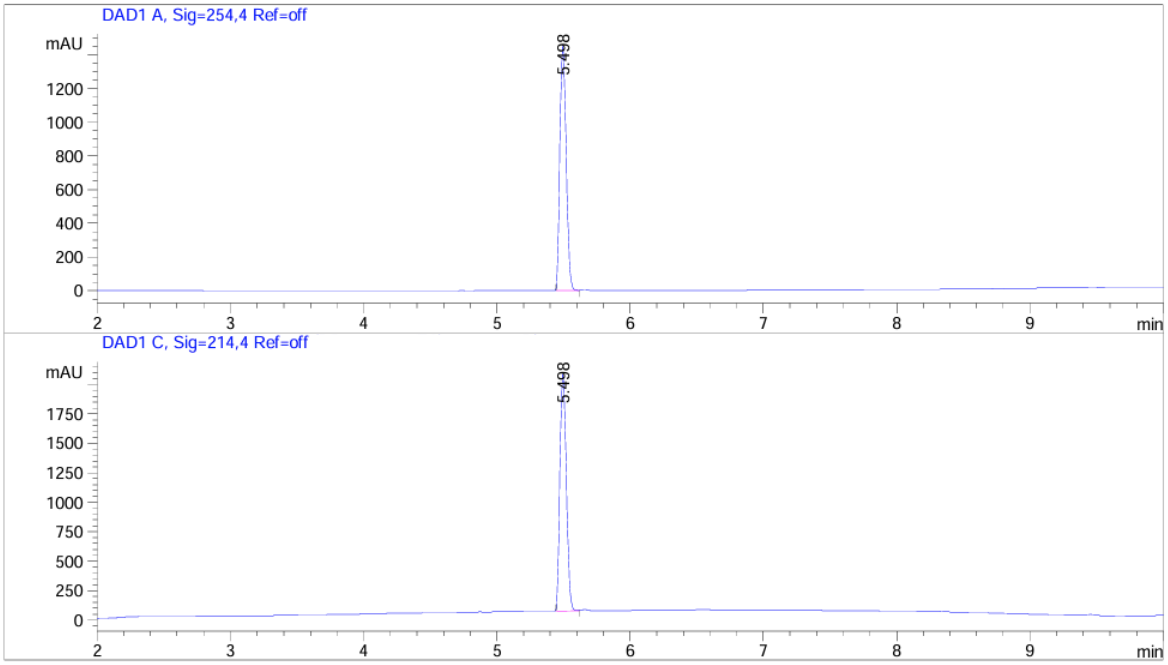


| Peak # | RetTime [min] | Type | Width [min] | Area [mAU*s] | Height [mAU] | Area % |
|---|---|---|---|---|---|---|
| 1 | 5.498 | BV | 0.0522 | 4765.96387 | 1456.16187 | 100.0000 |

Totals : 4765.96387 1456.16187

Signal 2: DAD1 C, Sig=214,4 Ref=off

| Peak # | RetTime [min] | Type | Width [min] | Area [mAU*s] | Height [mAU] | Area % |
|---|---|---|---|---|---|---|
| 1 | 5.498 | BV | 0.0528 | 6716.92188 | 2020.75269 | 100.0000 |

Totals : 6716.92188 2020.75269

[1]H NMR (400 MHz, $CD_3OD$)

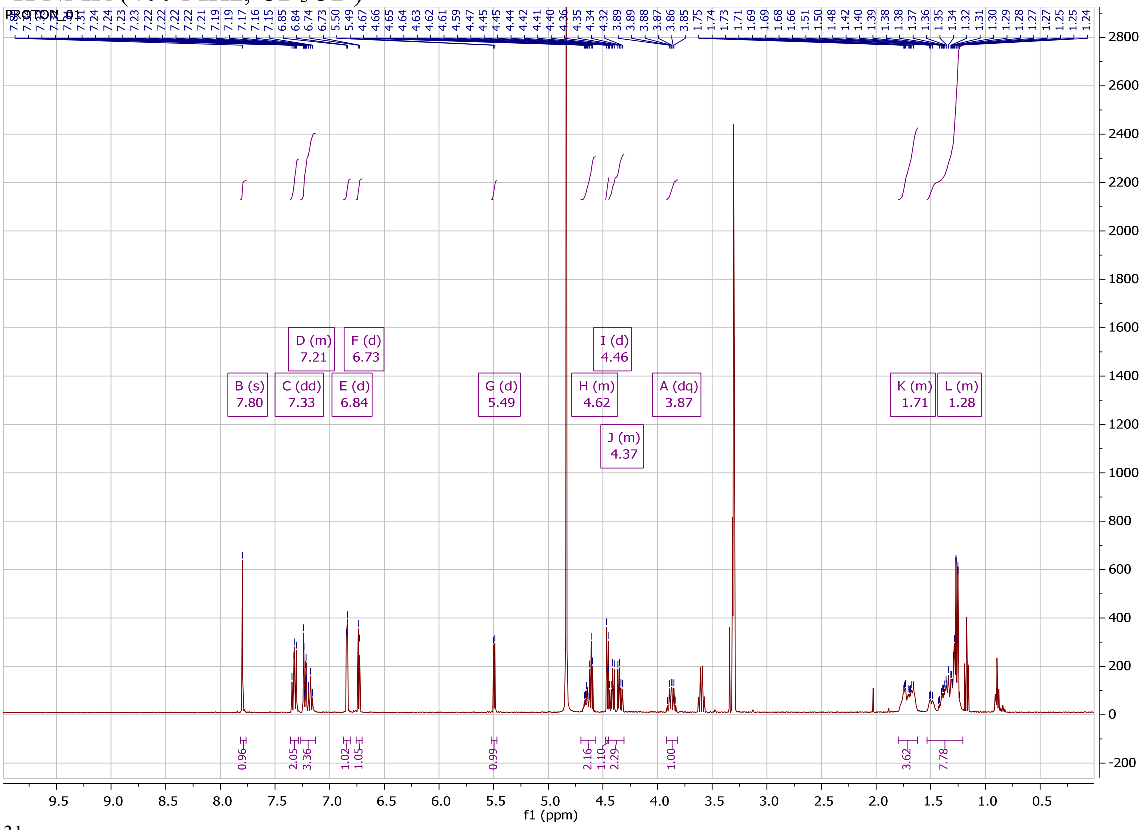

[31]P NMR (162 MHz, $CD_3OD$)

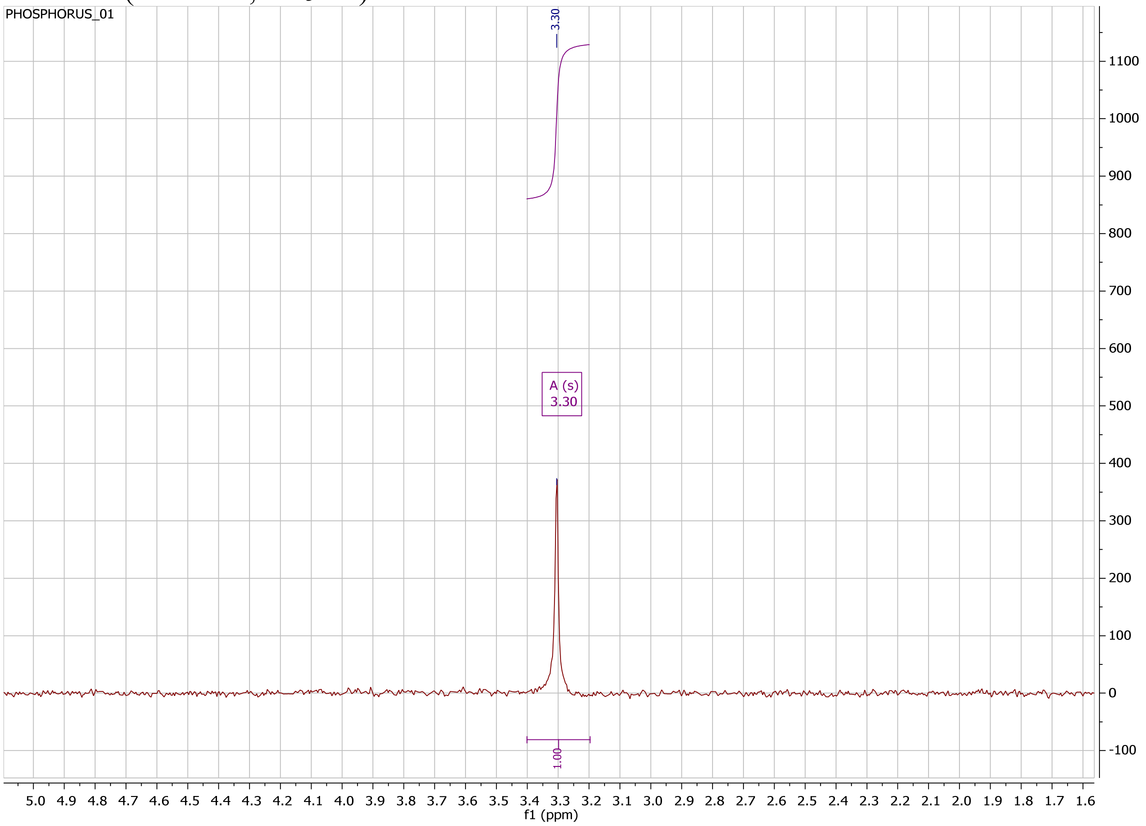

#### COMPOUND 28F-*R*P HPLC AND NMR

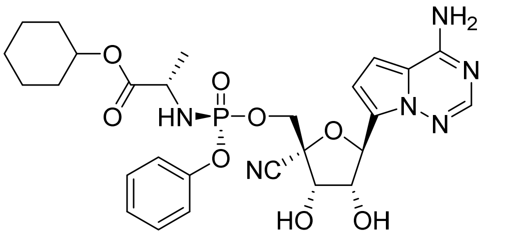


Method Info : Sample Bank Method - 2-98%B with 8.5 min gradient, A=Water + 0.1% TFA, B= Acetonitrile + 0.1% TFA; 1.5 mL/min; Column: Phenomenex Kinetex C18, 2.6u 100A, 4.6 x 100 mm; Instrument 1290II

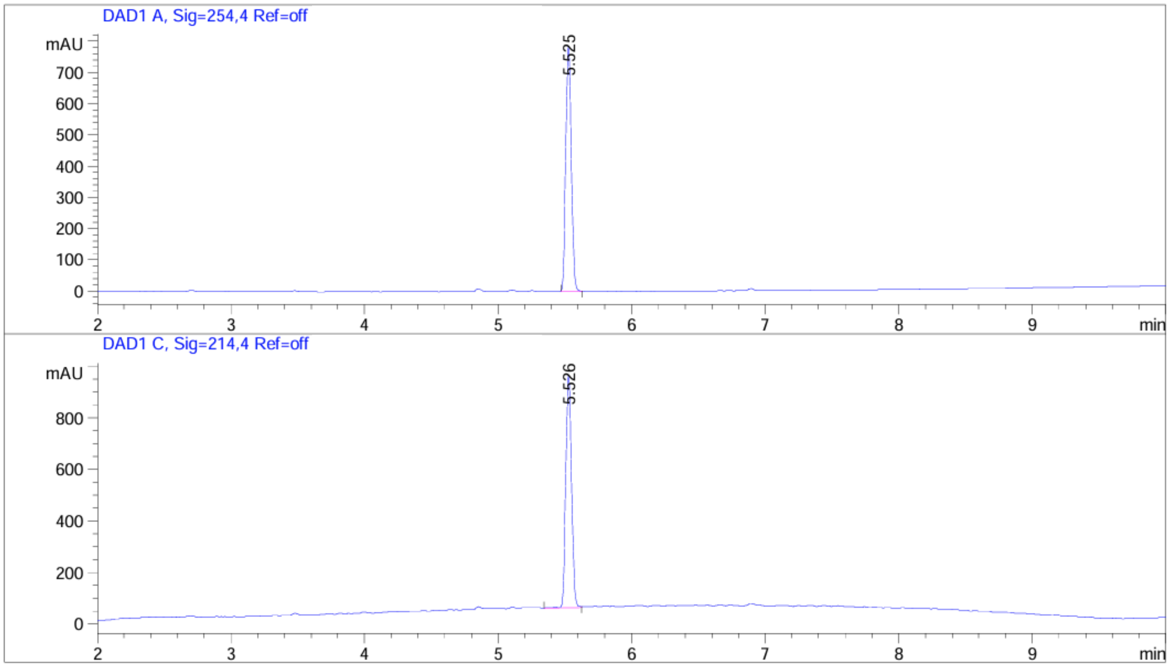


| Peak # | RetTime [min] | Type | Width [min] | Area [mAU*s] | Height [mAU] | Area % |
|---|---|---|---|---|---|---|
| 1 | 5.525 | BB | 0.0460 | 2289.56152 | 785.31079 | 100.0000 |

Totals : 2289.56152 785.31079

Signal 2: DAD1 C, Sig=214,4 Ref=off

| Peak # | RetTime [min] | Type | Width [min] | Area [mAU*s] | Height [mAU] | Area % |
|---|---|---|---|---|---|---|
| 1 | 5.526 | BB | 0.0493 | 2742.62573 | 905.18005 | 100.0000 |

Totals : 2742.62573 905.18005

[1]H NMR (400 MHz, CD3OD)

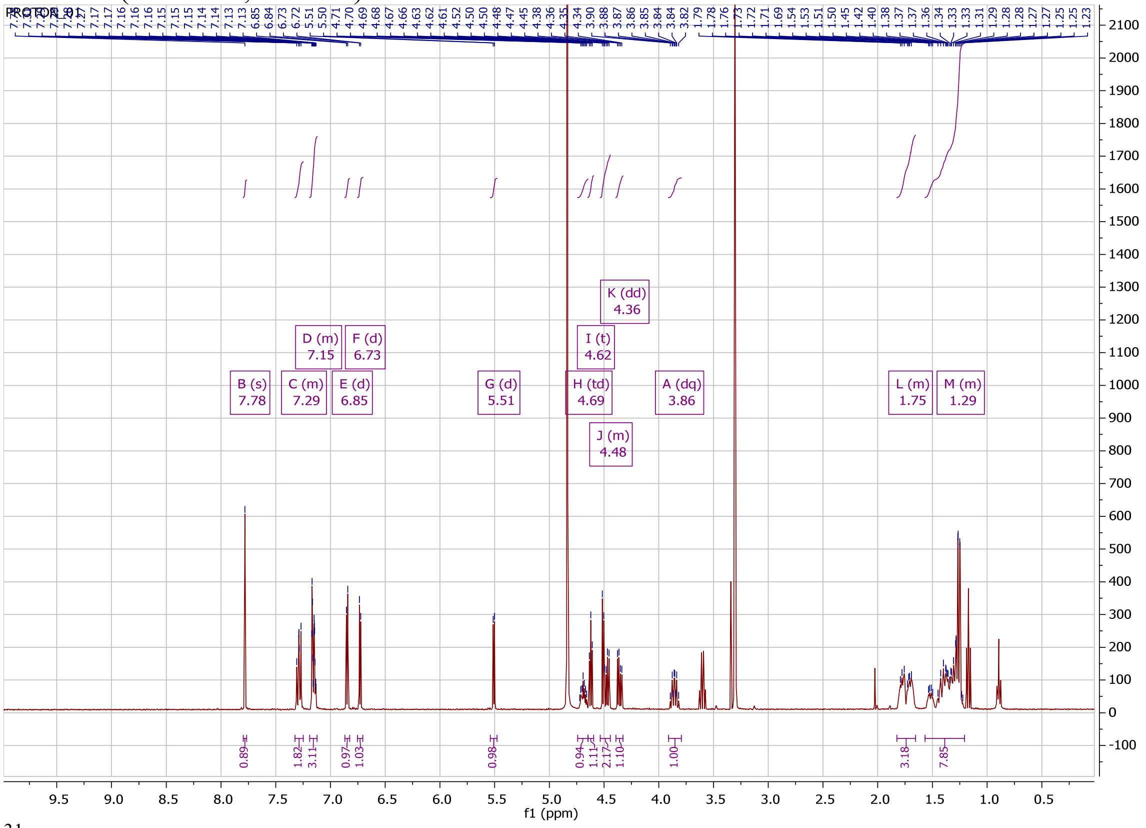

[31]P NMR (162 MHz, CD3OD)

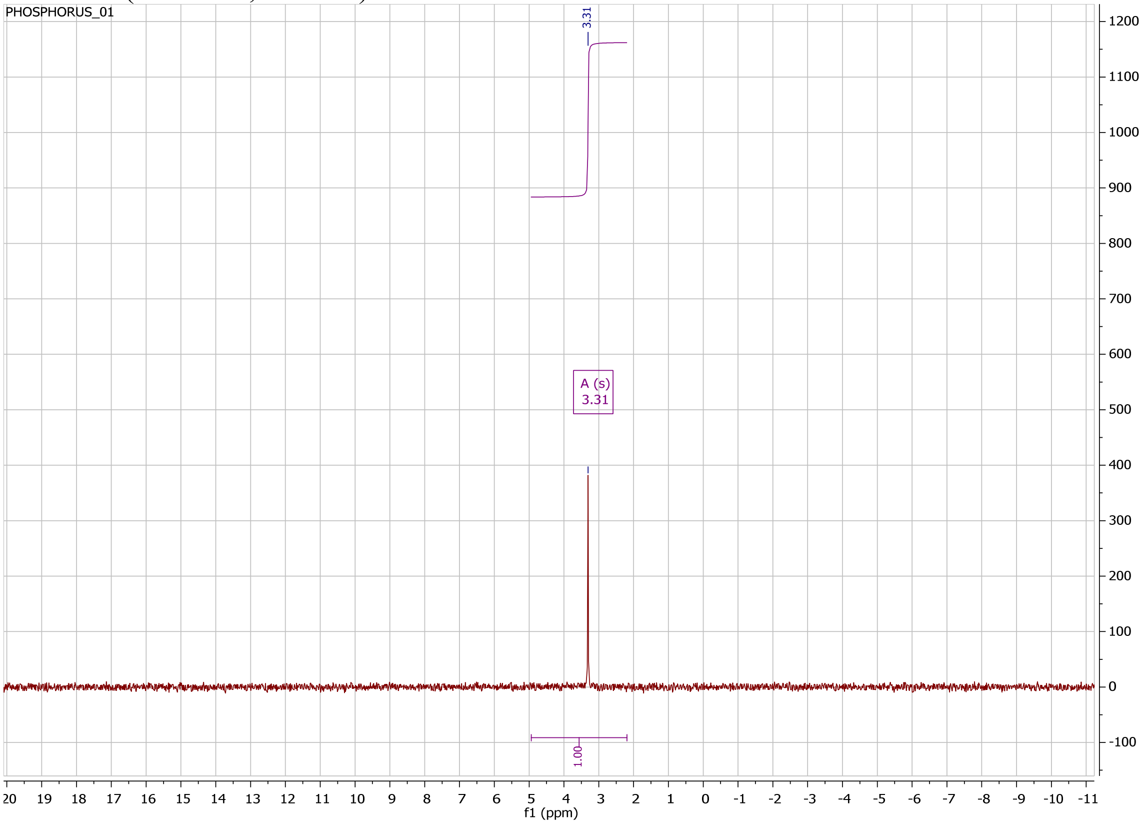

**COMPOUND 28G HPLC**

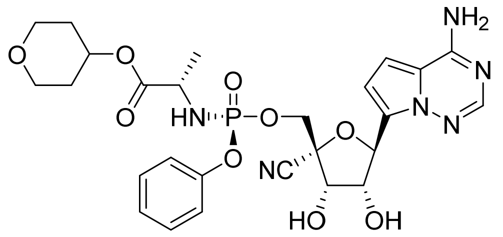

Method Info : Sample Bank Method - 2-98%B with 8.5 min gradient, A=Water + 0.1% TFA, B= Acetonitrile + 0.1% TFA; 1.5 mL/min; Column: Phenomenex Kinetex C18, 2.6u 100A, 4.6 x 100 mm; Instrument 1290II

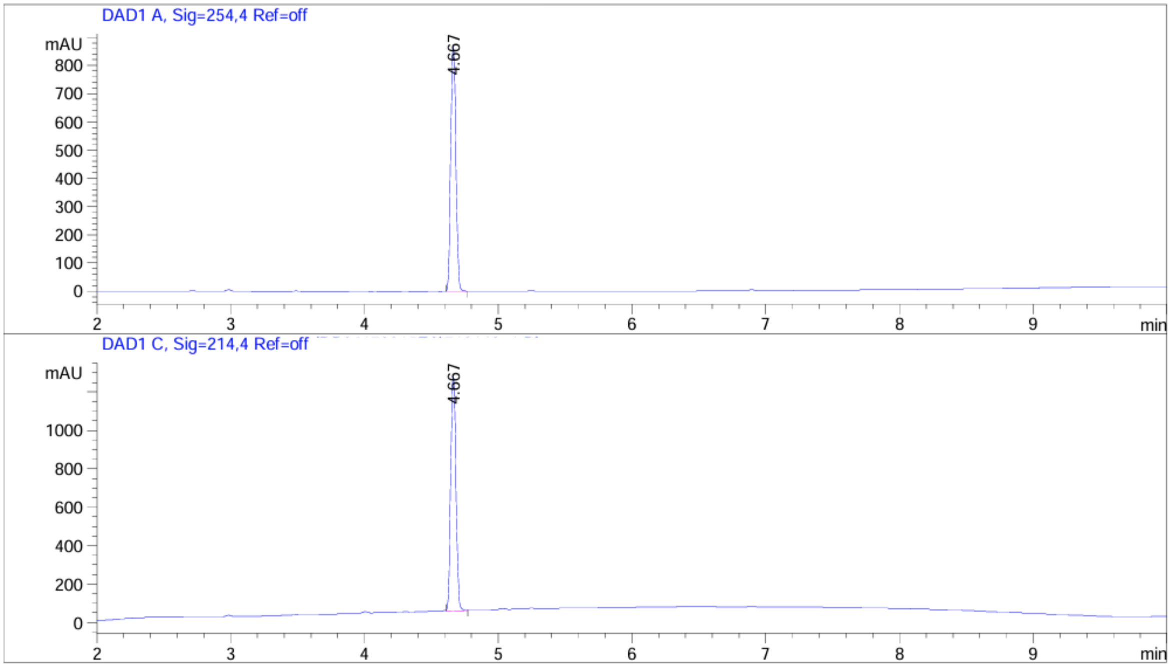

| Peak # | RetTime [min] | Type | Width [min] | Area [mAU*s] | Height [mAU] | Area % |
|---|---|---|---|---|---|---|
| 1 | 4.667 | BB | 0.0436 | 2355.36499 | 870.73718 | 100.0000 |

Totals : 2355.36499 870.73718

Signal 2: DAD1 C, Sig=214,4 Ref=off

| Peak # | RetTime [min] | Type | Width [min] | Area [mAU*s] | Height [mAU] | Area % |
|---|---|---|---|---|---|---|
| 1 | 4.667 | BB | 0.0437 | 3341.28564 | 1228.52222 | 100.0000 |

Totals : 3341.28564 1228.52222

#### COMPUND 1 HPLC AND NMR

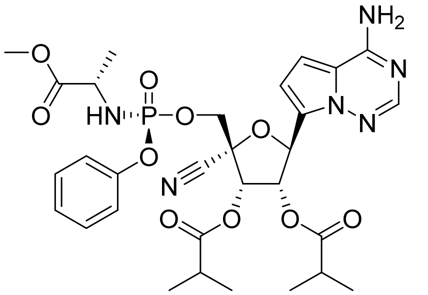

Method Info : Sample Bank Method - 2-98%B with 8.5 min gradient, A=Water + 0.1% TFA, B= Acetonitrile + 0.1% TFA; 1.5 mL/min; Column: Phenomenex Kinetex C18, 2.6u 100A, 4.6 x 100 mm; Instrument 1290II

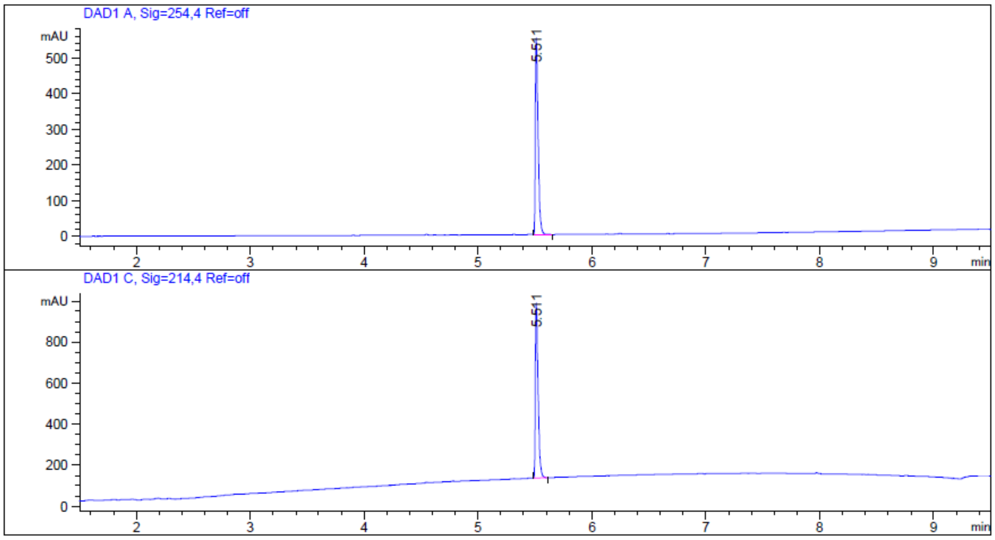


| Peak # | RetTime [min] | Type | Width [min] | Area [mAU*s] | Height [mAU] | Area % |
|---|---|---|---|---|---|---|
| 1 | 5.511 | BB | 0.0264 | 940.01471 | 549.95306 | 100.0000 |
| Totals : | | | | 940.01471 | 549.95306 | |

Signal 2: DAD1 C, Sig=214,4 Ref=off

| Peak # | RetTime [min] | Type | Width [min] | Area [mAU*s] | Height [mAU] | Area % |
|---|---|---|---|---|---|---|
| 1 | 5.511 | BB | 0.0258 | 1443.86108 | 852.43799 | 100.0000 |
| Totals : | | | | 1443.86108 | 852.43799 | |

$^{1}H$ NMR (400 MHz, $CD_3OD$)

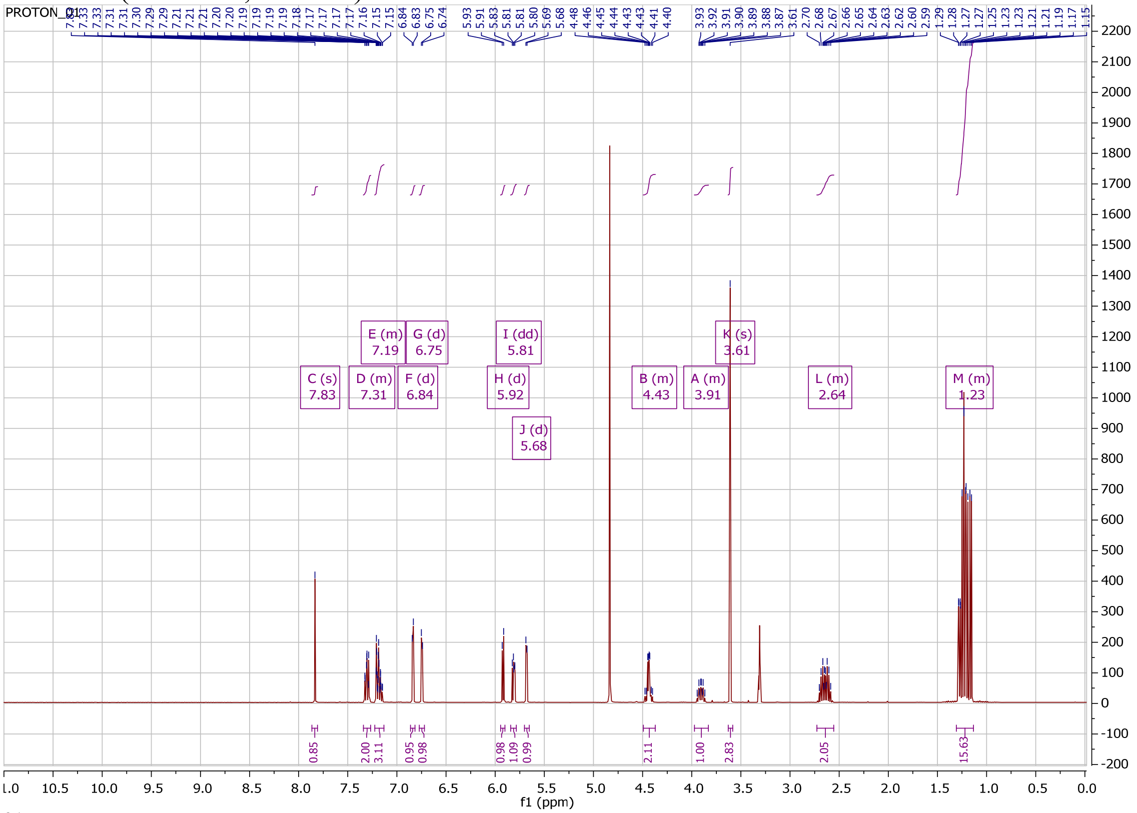

$^{31}P$ NMR (162 MHz, $CD_3OD$)

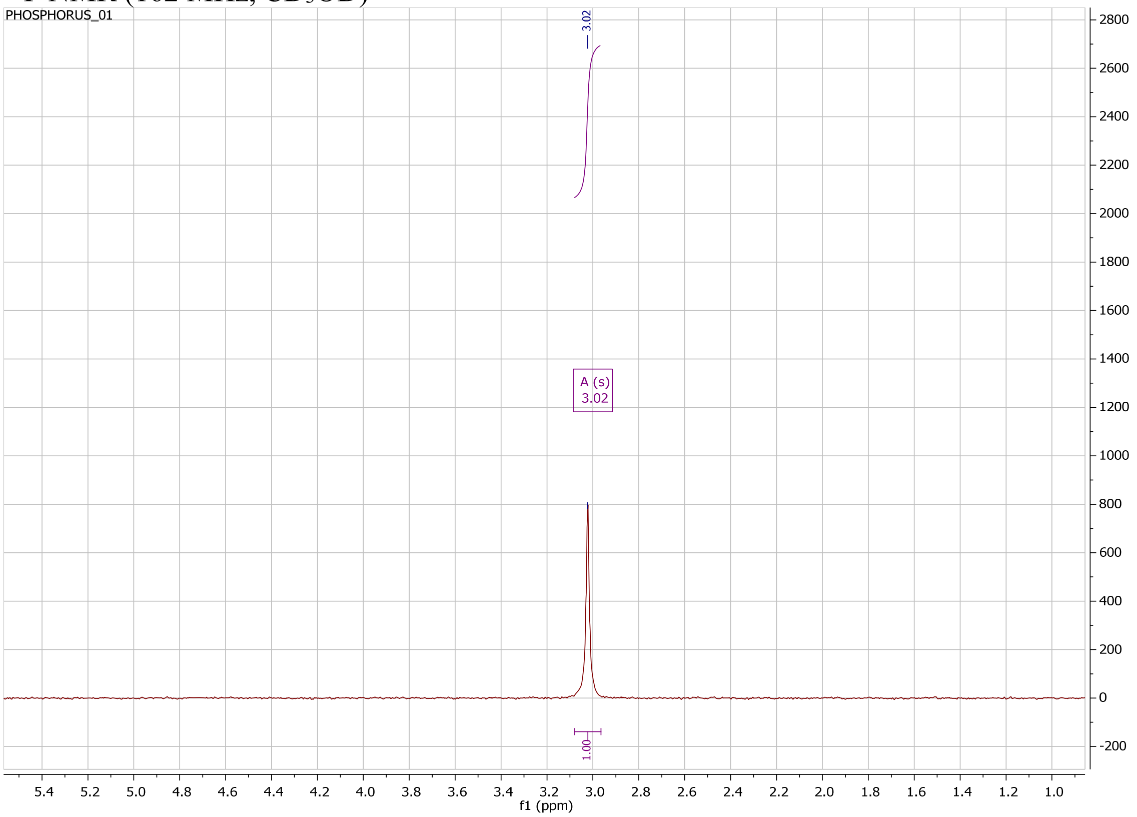

#### COMPOUDN 29AL-*RP* HPLC AND NMR

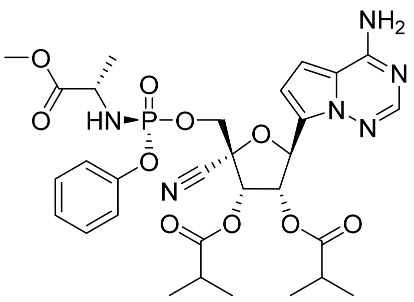

Method Info : Sample Bank Method - 2-98%B with 8.5 min gradient, A=Water + 0.1% TFA, B= Acetonitrile + 0.1% TFA; 1.5 mL/min; Column: Phenomenex Kinetex C18, 2.6u 100A, 4.6 x 100 mm; Instrument 1290II

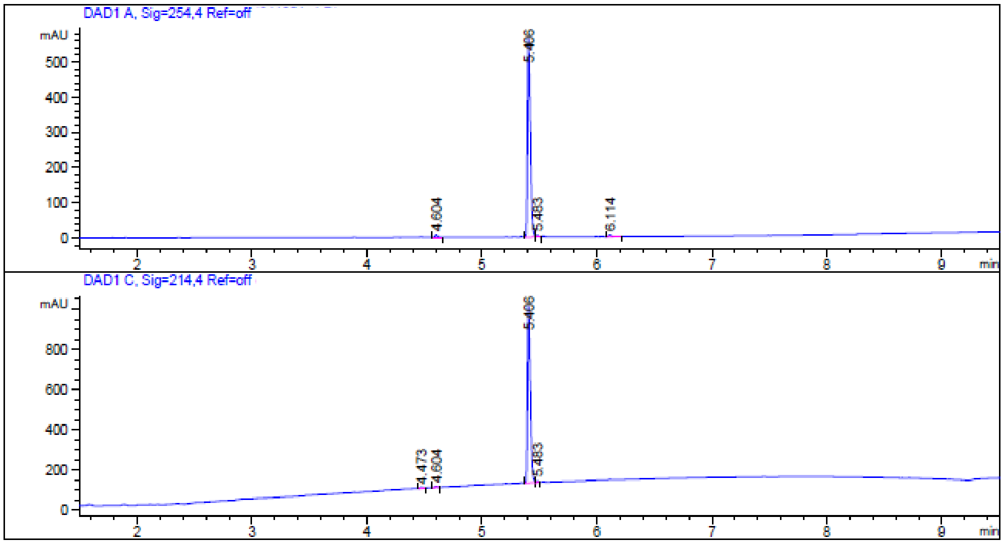


| Peak # | RetTime [min] | Type | Width [min] | Area [mAU*s] | Height [mAU] | Area % |
|---|---|---|---|---|---|---|
| 1 | 4.604 | BB | 0.0202 | 9.17336 | 7.02890 | 0.9637 |
| 2 | 5.406 | BV | 0.0252 | 927.03345 | 564.31952 | 97.3845 |
| 3 | 5.483 | VB | 0.0189 | 6.56893 | 5.31225 | 0.6901 |
| 4 | 6.114 | BB | 0.0272 | 9.15565 | 5.04966 | 0.9618 |

Totals : 951.93138 581.71034

Signal 2: DAD1 C, Sig=214,4 Ref=off

| Peak # | RetTime [min] | Type | Width [min] | Area [mAU*s] | Height [mAU] | Area % |
|---|---|---|---|---|---|---|
| 1 | 4.473 | BB | 0.0212 | 7.28327 | 5.09495 | 0.4994 |
| 2 | 4.604 | BB | 0.0215 | 7.67163 | 5.43754 | 0.5261 |
| 3 | 5.406 | BV | 0.0250 | 1433.64978 | 880.87109 | 98.3099 |
| 4 | 5.483 | VB | 0.0189 | 9.69230 | 7.85642 | 0.6646 |

Totals : 1458.29697 899.26000

$^{1}H$ NMR (400 MHz, $CD_3OD$)

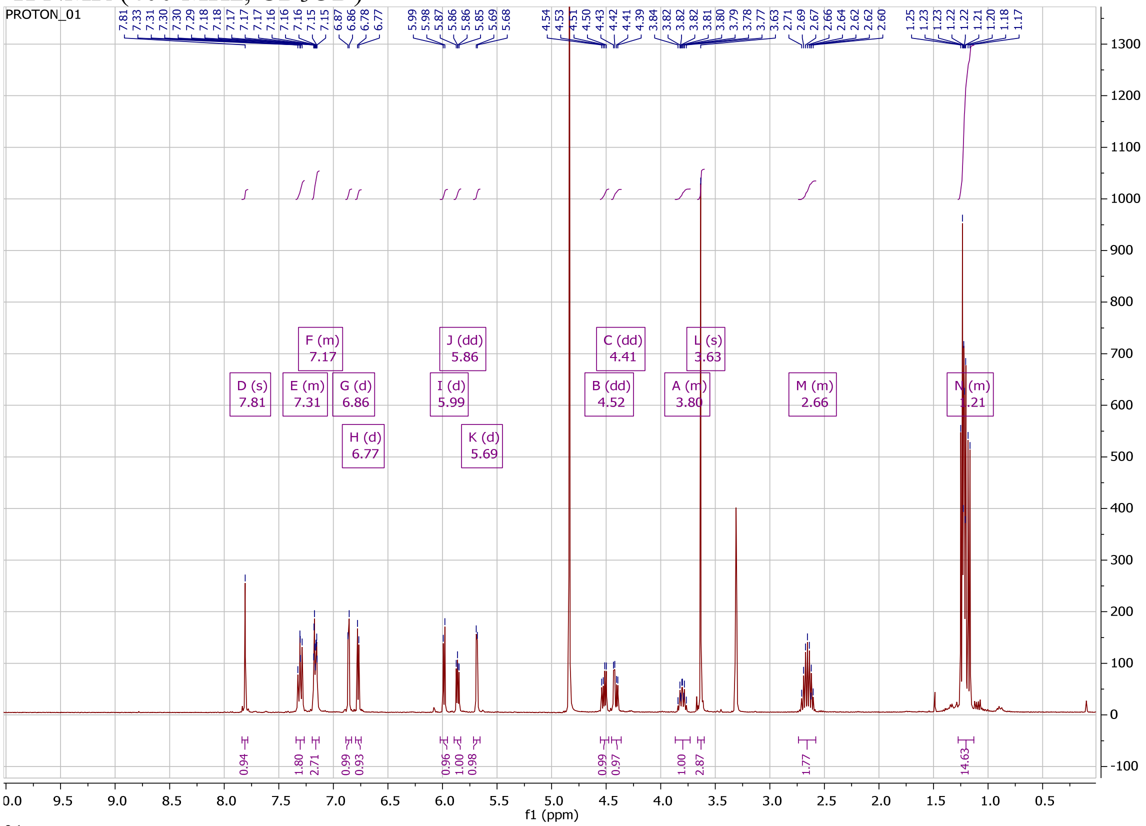

$^{31}P$ NMR (162 MHz, $CD_3OD$)

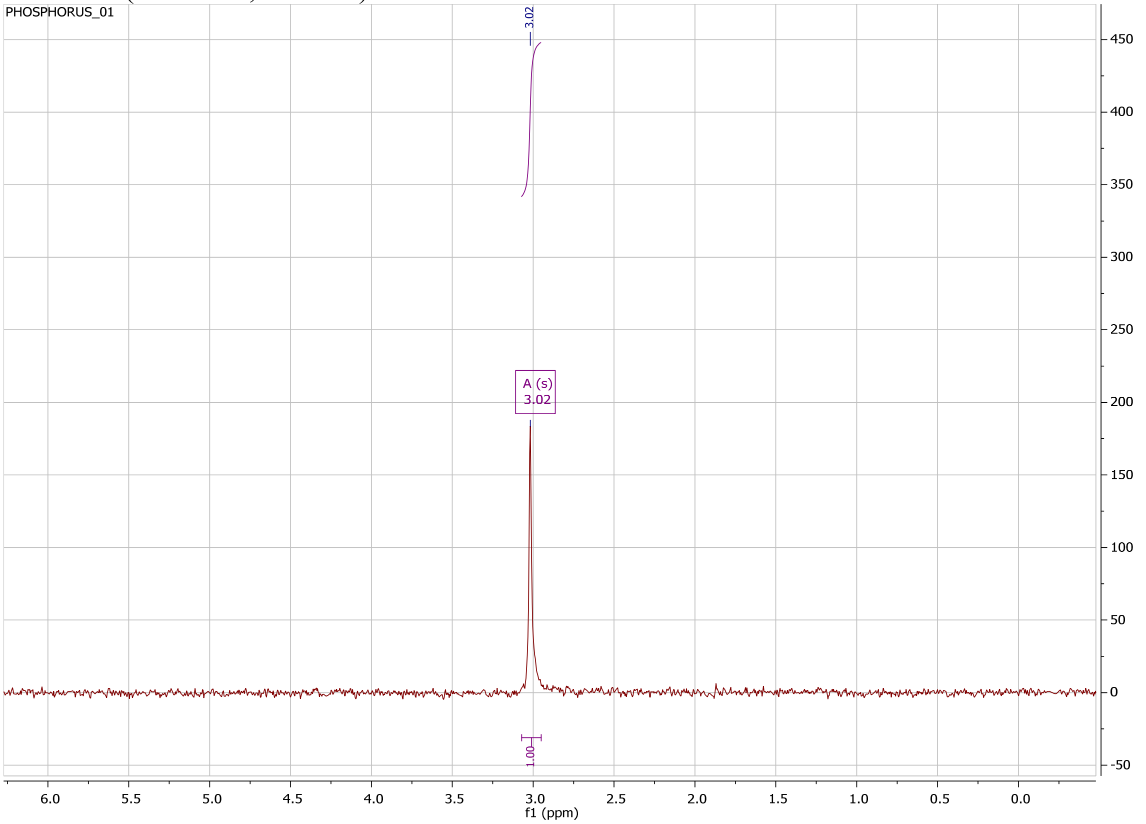

#### COMPOUND 29AM HPLC AND NMR

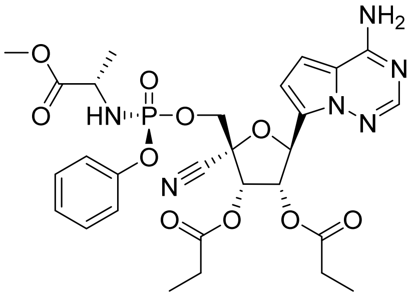

```
Method Info      : Sample Bank Method - 2-98%B with 8.5 min gradient, A=Water + 0.1% TFA, B=
                   Acetonitrile + 0.1% TFA; 1.5 mL/min; Column: Phenomenex Kinetex C18, 2.6u
                   100A, 4.6 x 100 mm; Instrument 1290II
```

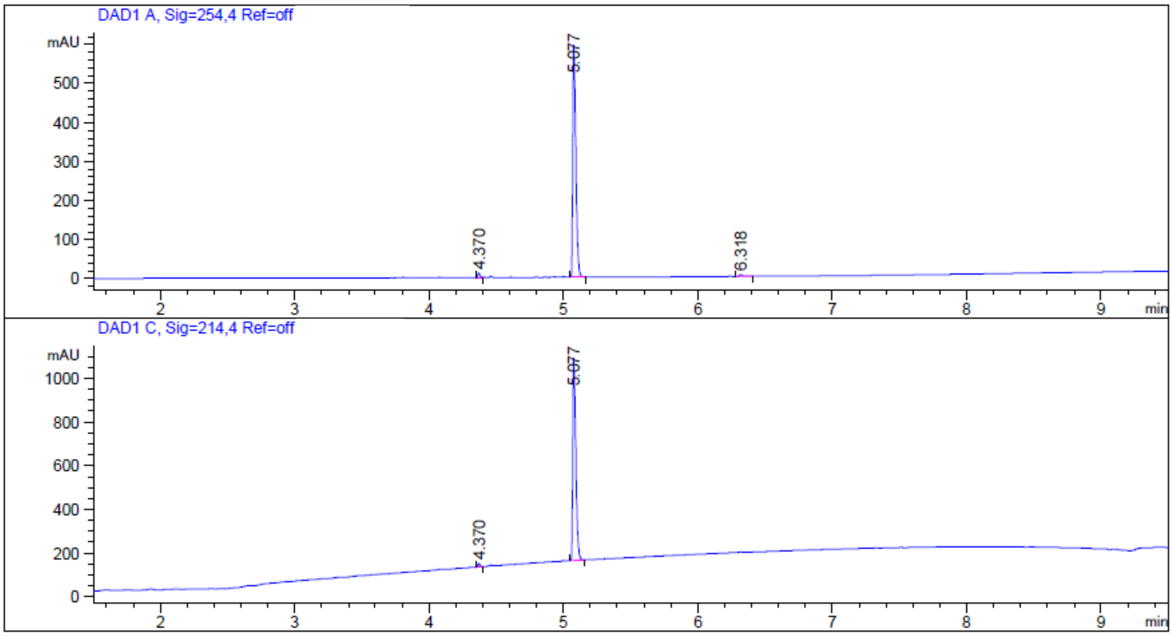


| Peak # | RetTime [min] | Type | Width [min] | Area [mAU*s] | Height [mAU] | Area % |
|---|---|---|---|---|---|---|
| 1 | 4.370 | BB | 0.0163 | 11.60877 | 10.96916 | 1.2022 |
| 2 | 5.077 | BB | 0.0245 | 945.77637 | 595.62177 | 97.9461 |
| 3 | 6.318 | BB | 0.0259 | 8.22346 | 4.69529 | 0.8516 |

Totals : 965.60859 611.28621

Signal 2: DAD1 C, Sig=214,4 Ref=off

| Peak # | RetTime [min] | Type | Width [min] | Area [mAU*s] | Height [mAU] | Area % |
|---|---|---|---|---|---|---|
| 1 | 4.370 | BB | 0.0164 | 17.48951 | 16.49323 | 1.1800 |
| 2 | 5.077 | BB | 0.0244 | 1464.62024 | 930.71680 | 98.8200 |

Totals : 1482.10975 947.21003

$^{1}H$ NMR (400 MHz, $CD_3OD$)

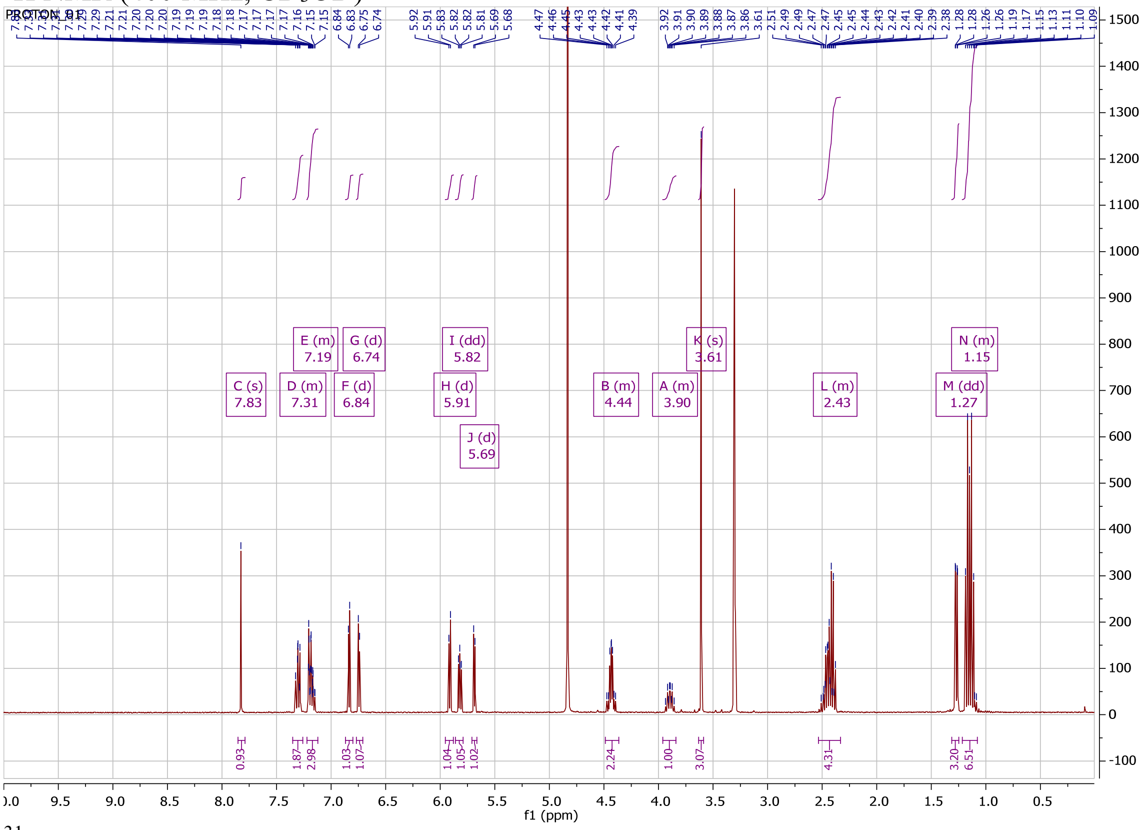

$^{31}P$ NMR (162 MHz, $CD_3OD$)

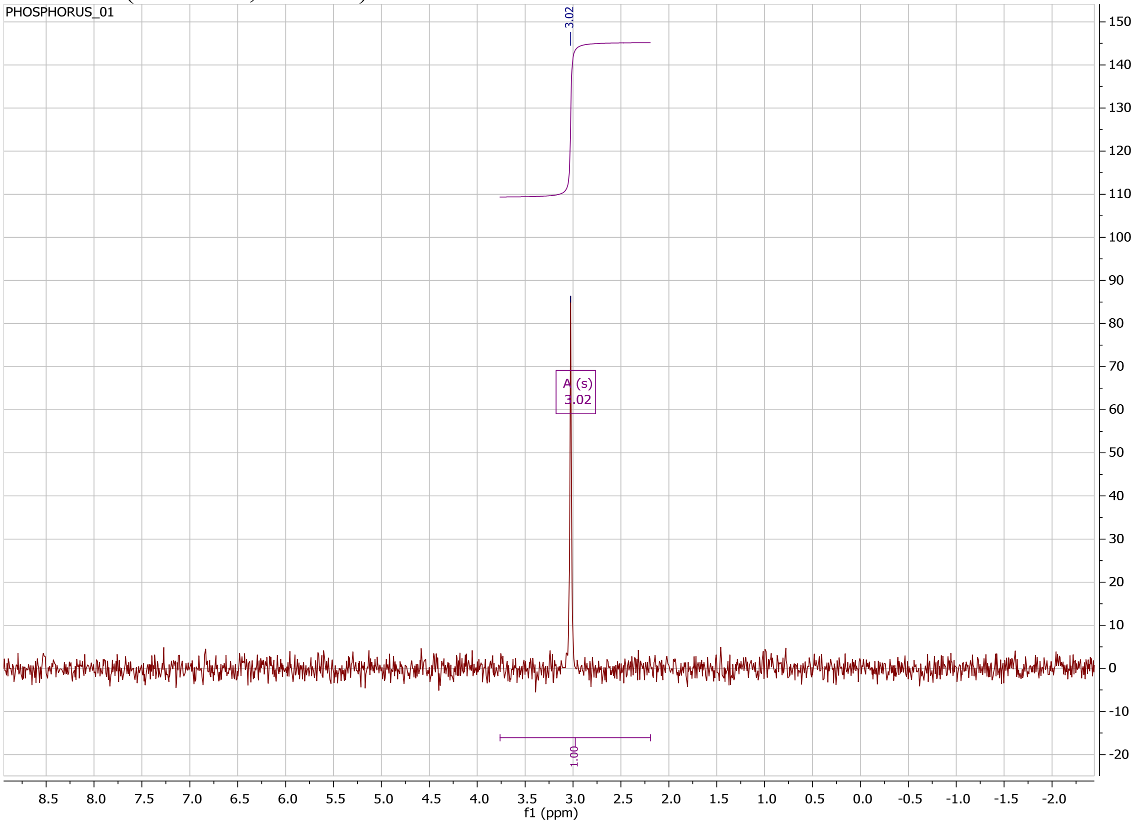

#### COMPOUND 29AN HPLC AND NMR

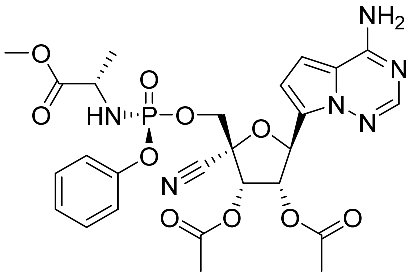

Method Info : Sample Bank Method - 2-98%B with 8.5 min gradient, A=Water + 0.1% TFA, B= Acetonitrile + 0.1% TFA; 1.5 mL/min; Column: Phenomenex Kinetex C18, 2.6u 100A, 4.6 x 100 mm; Instrument 1290II

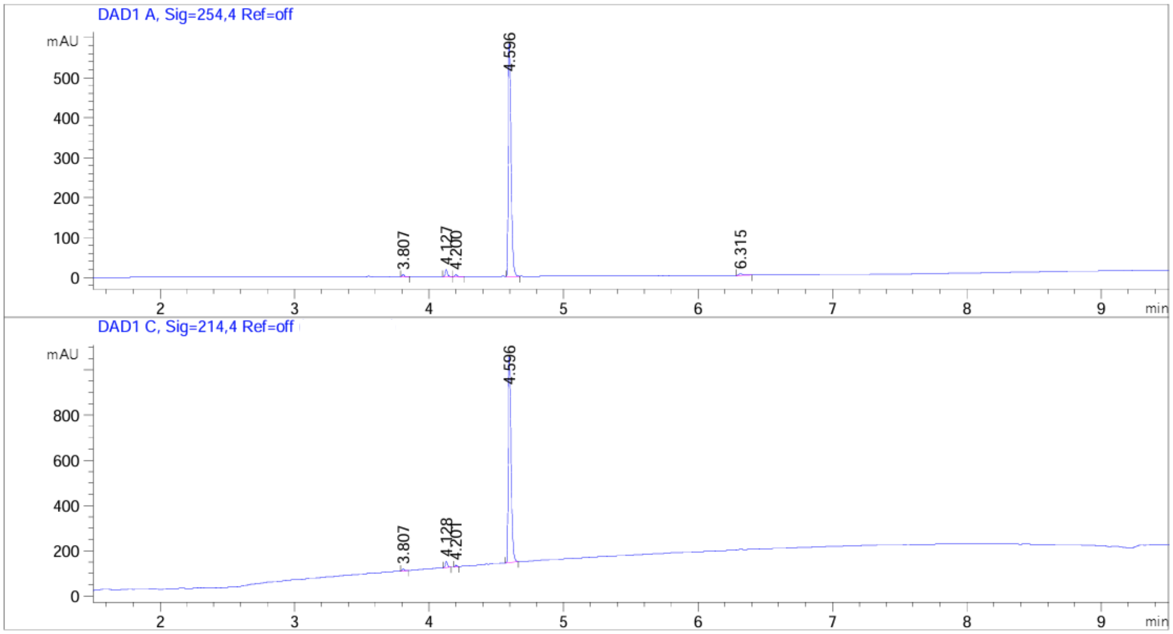


| Peak # | RetTime [min] | Type | Width [min] | Area [mAU*s] | Height [mAU] | Area % |
|---|---|---|---|---|---|---|
| 1 | 3.807 | BB | 0.0164 | 5.32209 | 5.00901 | 0.6007 |
| 2 | 4.127 | BB | 0.0167 | 19.30626 | 17.67804 | 2.1791 |
| 3 | 4.200 | BB | 0.0169 | 5.25874 | 4.75303 | 0.5936 |
| 4 | 4.596 | BB | 0.0225 | 848.03418 | 582.73889 | 95.7175 |
| 5 | 6.315 | BB | 0.0255 | 8.05524 | 4.70684 | 0.9092 |

Totals : 885.97651 614.88581

Signal 2: DAD1 C, Sig=214,4 Ref=off

| Peak # | RetTime [min] | Type | Width [min] | Area [mAU*s] | Height [mAU] | Area % |
|---|---|---|---|---|---|---|
| 1 | 3.807 | BB | 0.0188 | 9.94831 | 8.13479 | 0.7321 |
| 2 | 4.128 | BB | 0.0166 | 28.68180 | 26.60925 | 2.1107 |
| 3 | 4.201 | BB | 0.0156 | 6.88546 | 6.95347 | 0.5067 |
| 4 | 4.596 | BB | 0.0223 | 1313.36438 | 911.72516 | 96.6505 |

Totals : 1358.87994 953.42266

$^{1}H$ NMR (400 MHz, $CD_3OD$)

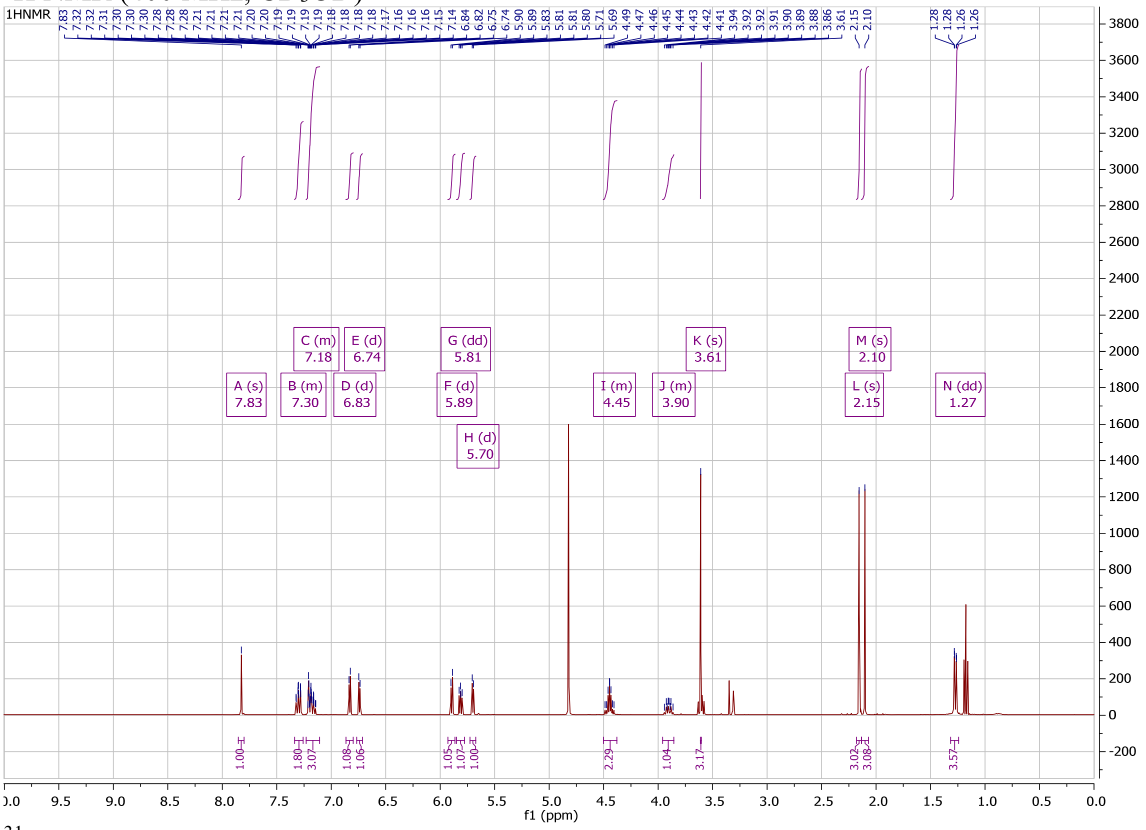

$^{31}P$ NMR (162 MHz, $CD_3OD$)

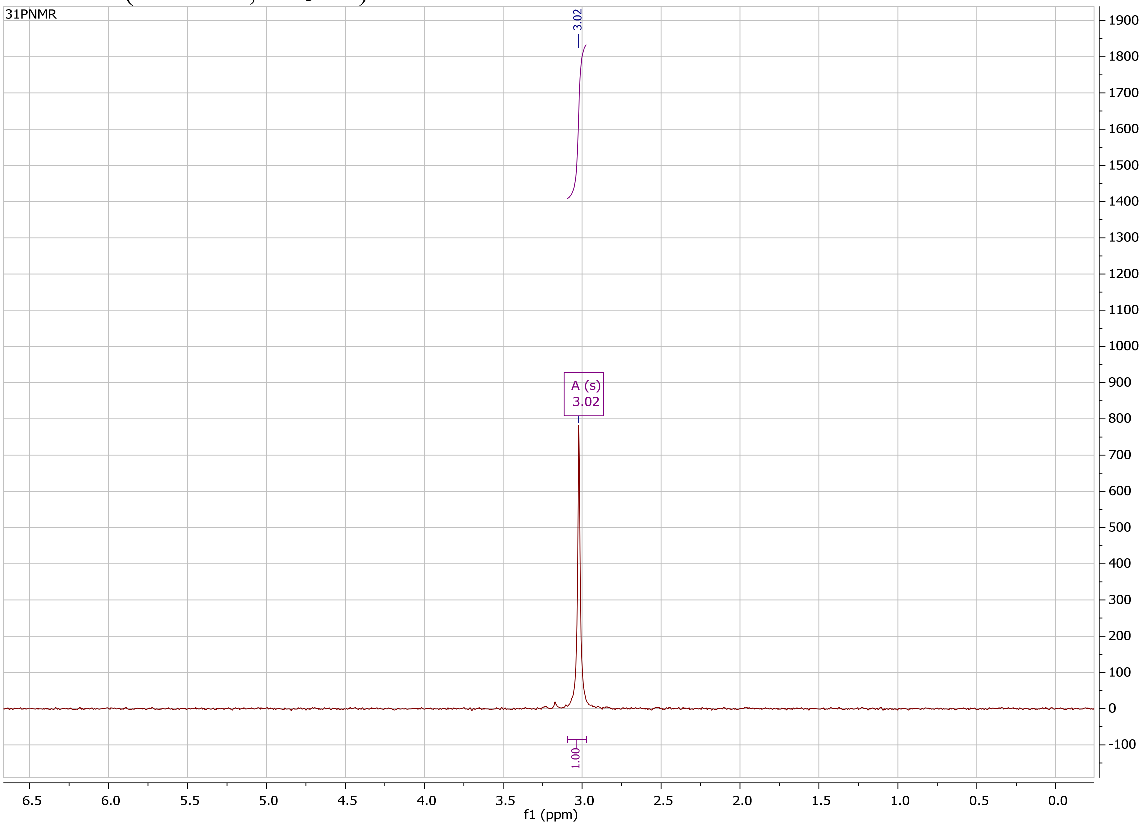

### COMPOUND 29CL HPLC AND NMR

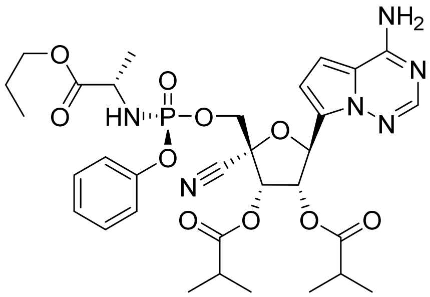


Method Info : Sample Bank Method - 2-98%B with 8.5 min gradient, A=Water + 0.1% TFA, B= Acetonitrile + 0.1% TFA; 1.5 mL/min; Column: Phenomenex Kinetex C18, 2.6u 100A, 4.6 x 100 mm; Instrument 1290II

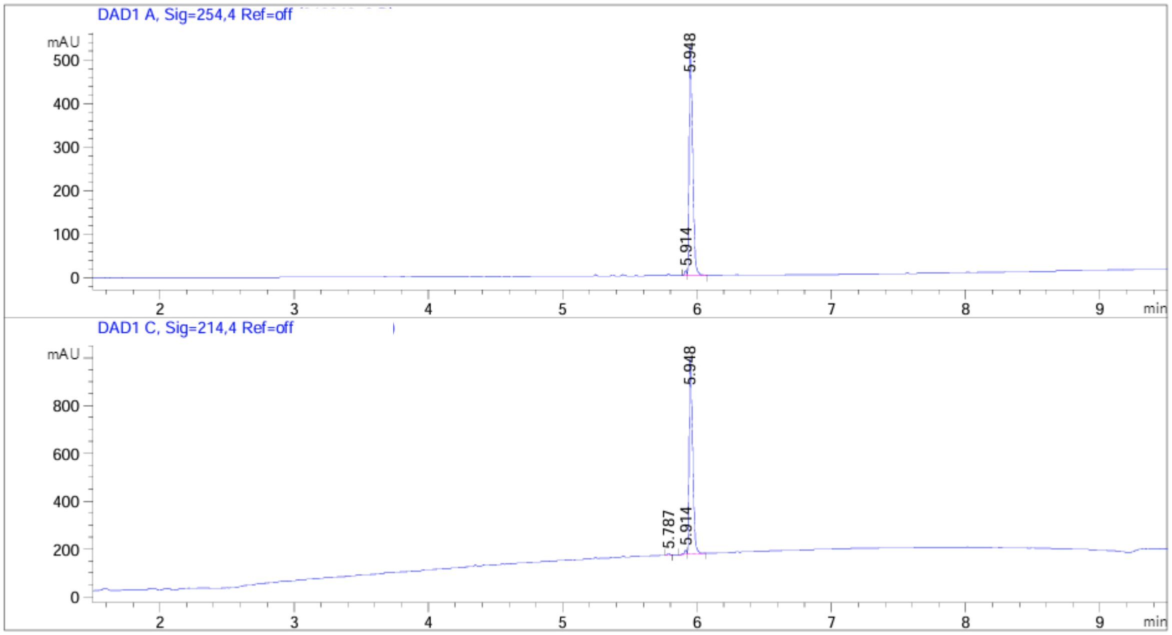


| Peak # | RetTime [min] | Type | Width [min] | Area [mAU*s] | Height [mAU] | Area % |
|---|---|---|---|---|---|---|
| 1 | 5.914 | BV | 0.0163 | 11.72219 | 11.08505 | 1.2116 |
| 2 | 5.948 | VB | 0.0285 | 955.79773 | 530.33435 | 98.7884 |

Totals : 967.51992 541.41940

Signal 2: DAD1 C, Sig=214,4 Ref=off

| Peak # | RetTime [min] | Type | Width [min] | Area [mAU*s] | Height [mAU] | Area % |
|---|---|---|---|---|---|---|
| 1 | 5.787 | BB | 0.0178 | 6.06997 | 5.30480 | 0.4058 |
| 2 | 5.914 | BV | 0.0166 | 18.06433 | 16.76927 | 1.2078 |
| 3 | 5.948 | VB | 0.0278 | 1471.56360 | 824.15332 | 98.3864 |

Totals : 1495.69789 846.22739

[1]H NMR (400 MHz, $CD_3OD$)

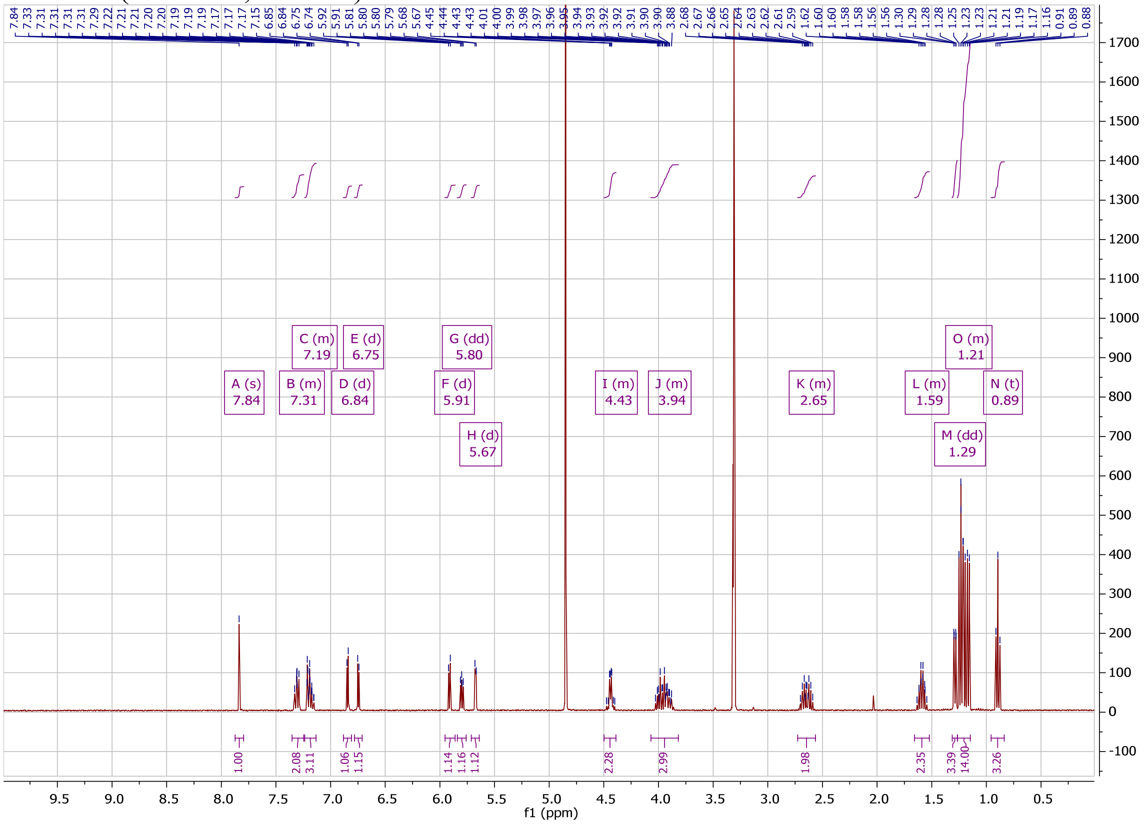

[31]P NMR (162 MHz, $CD_3OD$)

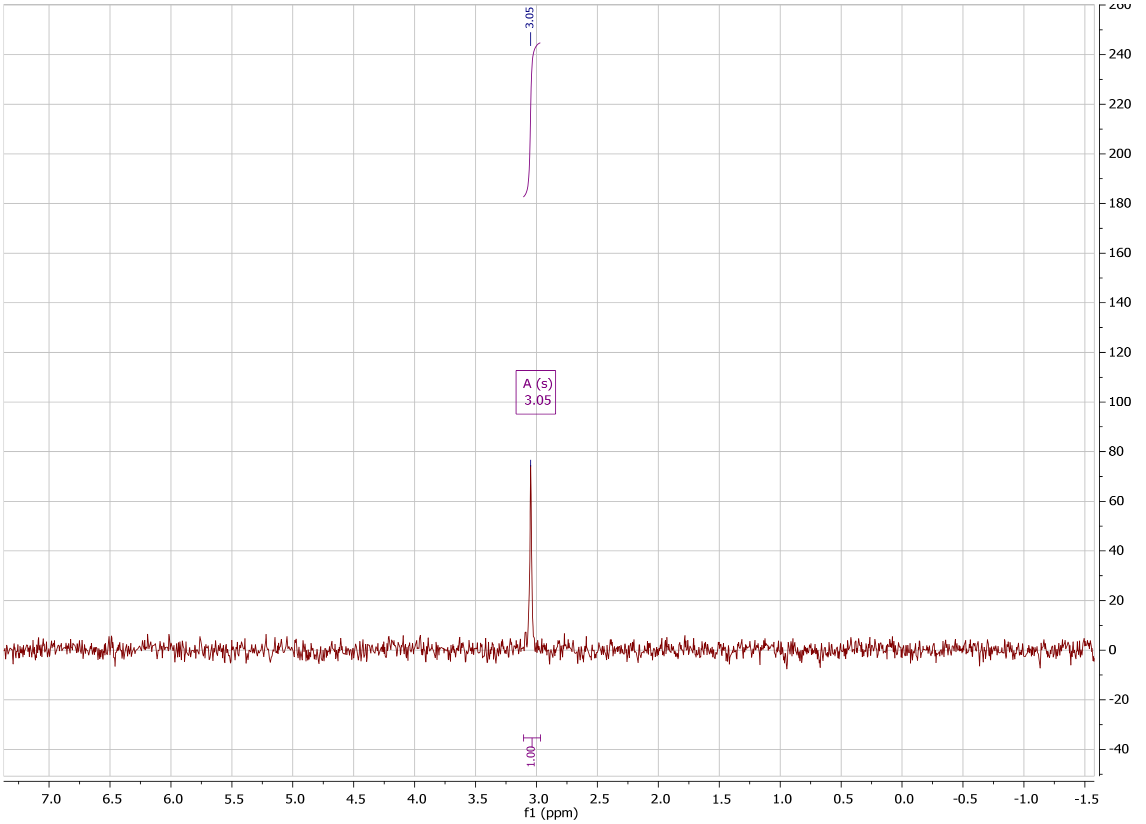

#### COMPOUND 29CM HPLC AND NMR

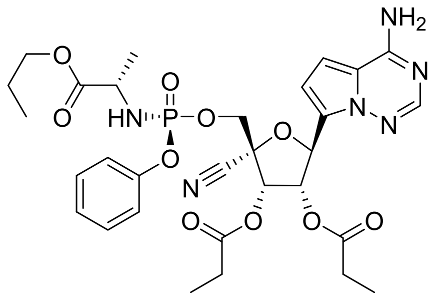

```
Method Info      : Sample Bank Method - 2-98%B with 8.5 min gradient, A=Water + 0.1% TFA, B=
                   Acetonitrile + 0.1% TFA; 1.5 mL/min; Column: Phenomenex Kinetex C18, 2.6u
                   100A, 4.6 x 100 mm; Instrument 1290II
```

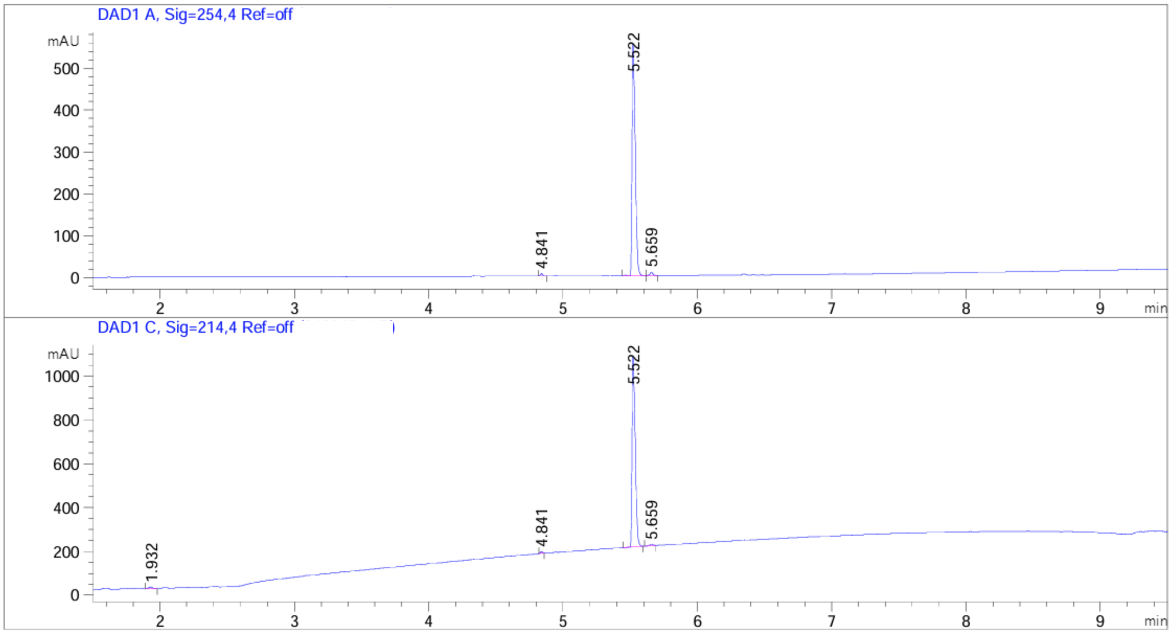


| Peak # | RetTime [min] | Type | Width [min] | Area [mAU*s] | Height [mAU] | Area % |
|---|---|---|---|---|---|---|
| 1 | 4.841 | BB | 0.0172 | 6.37103 | 5.86168 | 0.6787 |
| 2 | 5.522 | BV | 0.0264 | 919.29144 | 553.43988 | 97.9286 |
| 3 | 5.659 | VB | 0.0218 | 13.07410 | 8.80509 | 1.3927 |

Totals : 938.73657 568.10665

Signal 2: DAD1 C, Sig=214,4 Ref=off

| Peak # | RetTime [min] | Type | Width [min] | Area [mAU*s] | Height [mAU] | Area % |
|---|---|---|---|---|---|---|
| 1 | 1.932 | BB | 0.0288 | 15.05377 | 7.70043 | 1.0308 |
| 2 | 4.841 | BB | 0.0155 | 8.07816 | 8.21799 | 0.5531 |
| 3 | 5.522 | BB | 0.0257 | 1425.67957 | 867.99982 | 97.6224 |
| 4 | 5.659 | BB | 0.0223 | 11.59026 | 7.39041 | 0.7936 |

Totals : 1460.40176 891.30865

$^{1}H$ NMR (400 MHz, $CD_3OD$)

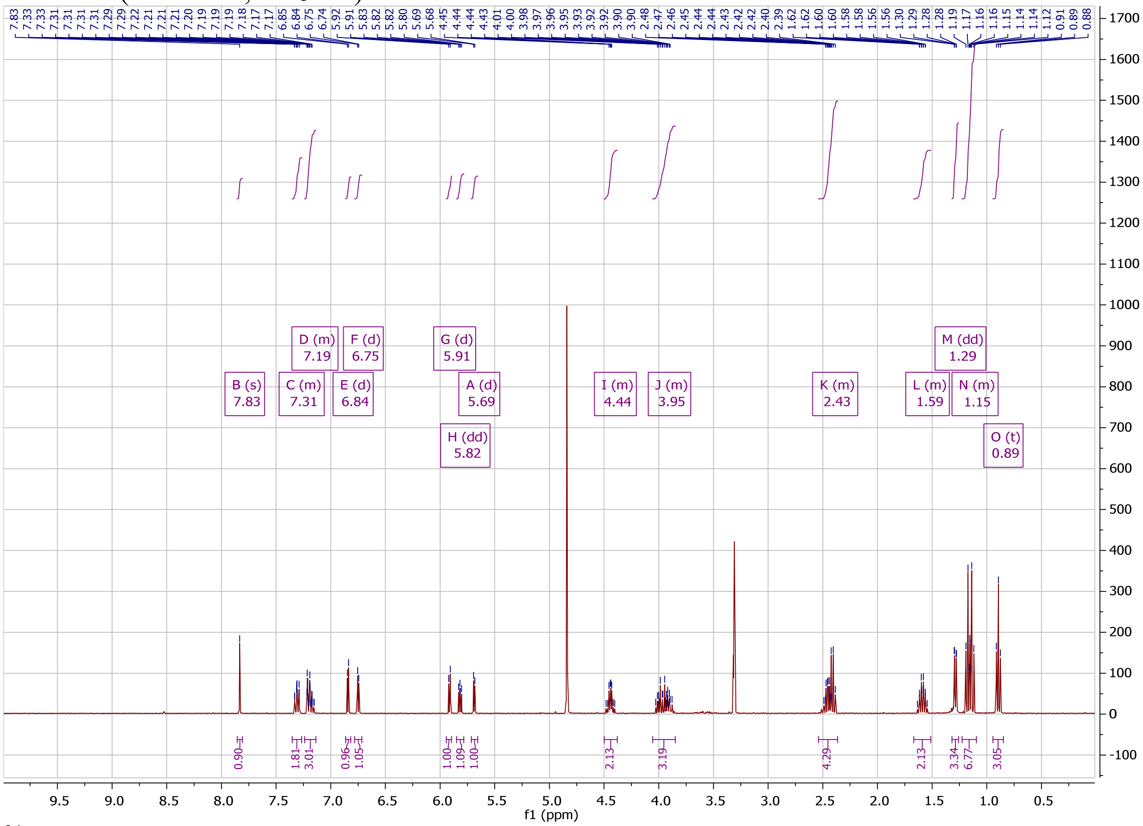

$^{31}P$ NMR (162 MHz, $CD_3OD$)

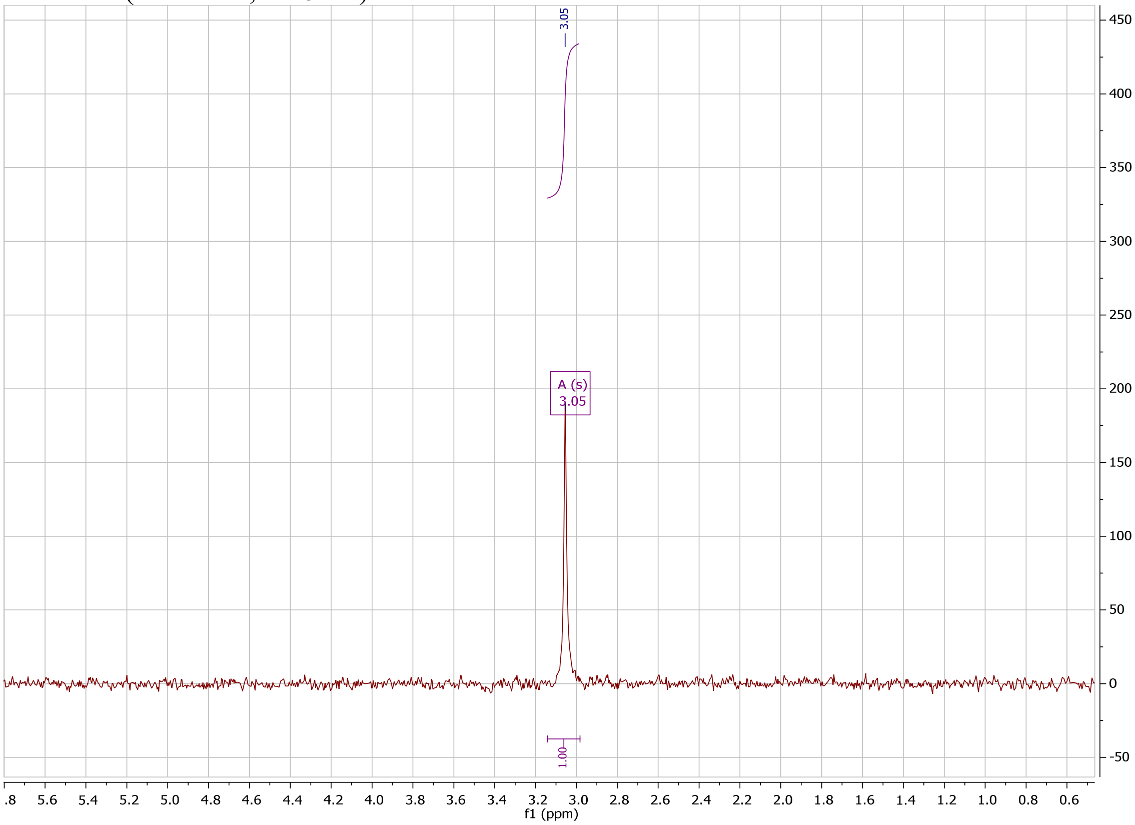

#### COMPOUND 29FL HPLC AND NMR

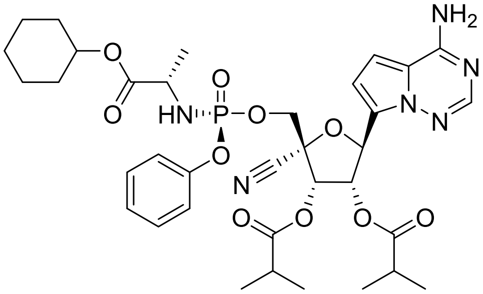


Method Info : Sample Bank Method - 2-98%B with 8.5 min gradient, A=Water + 0.1% TFA, B= Acetonitrile + 0.1% TFA; 1.5 mL/min; Column: Phenomenex Kinetex C18, 2.6u 100A, 4.6 x 100 mm; Instrument 1290II

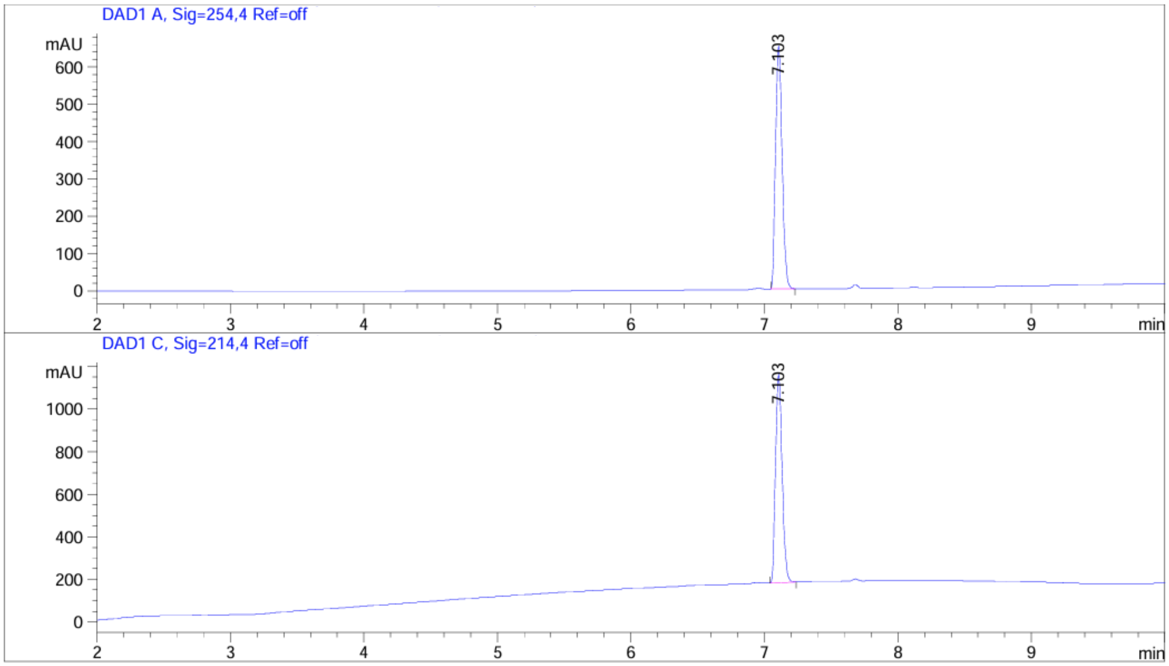


| Peak # | RetTime [min] | Type | Width [min] | Area [mAU*s] | Height [mAU] | Area % |
|---|---|---|---|---|---|---|
| 1 | 7.103 | BB | 0.0538 | 2228.34717 | 652.61346 | 100.0000 |
| Totals : | | | | 2228.34717 | 652.61346 | |

Signal 2: DAD1 C, Sig=214,4 Ref=off

| Peak # | RetTime [min] | Type | Width [min] | Area [mAU*s] | Height [mAU] | Area % |
|---|---|---|---|---|---|---|
| 1 | 7.103 | BB | 0.0540 | 3354.05176 | 979.44244 | 100.0000 |
| Totals : | | | | 3354.05176 | 979.44244 | |

$^{1}H$ NMR (400 MHz, $CD_3OD$)

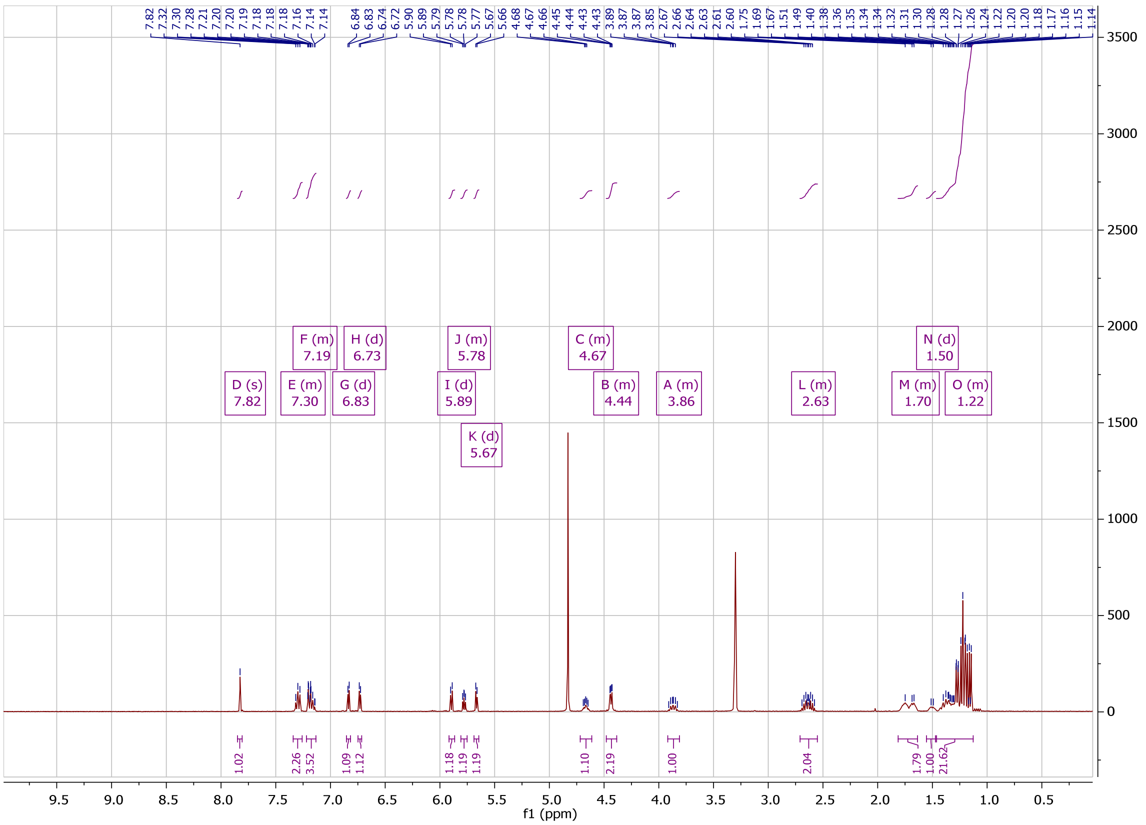

$^{31}P$ NMR (162 MHz, $CD_3OD$)

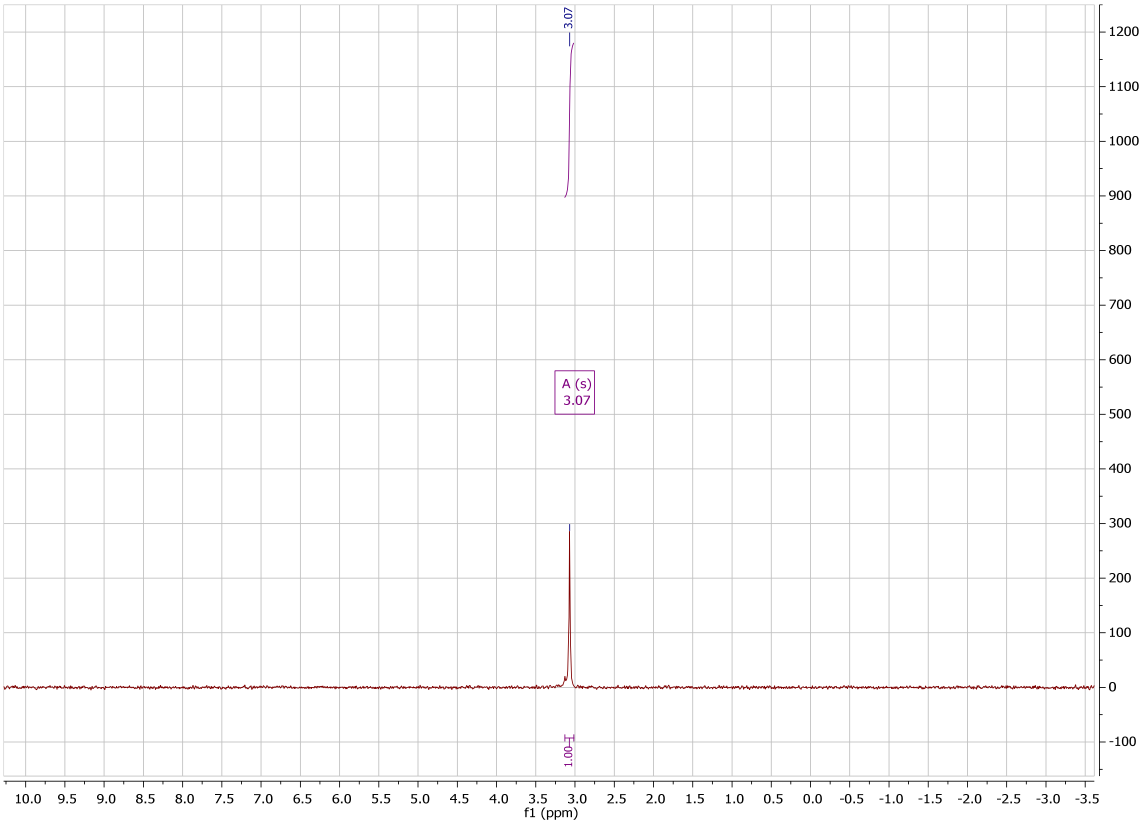

#### COMPOUND 29FM HPLC

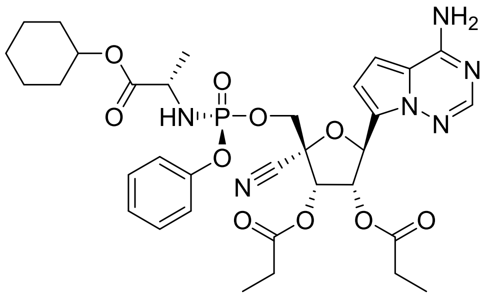

```
Method Info      : Sample Bank Method - 2-98%B with 8.5 min gradient, A=Water + 0.1% TFA, B=
                   Acetonitrile + 0.1% TFA; 1.5 mL/min; Column: Phenomenex Kinetex C18, 2.6u
                   100A, 4.6 x 100 mm; Instrument 1290II
```

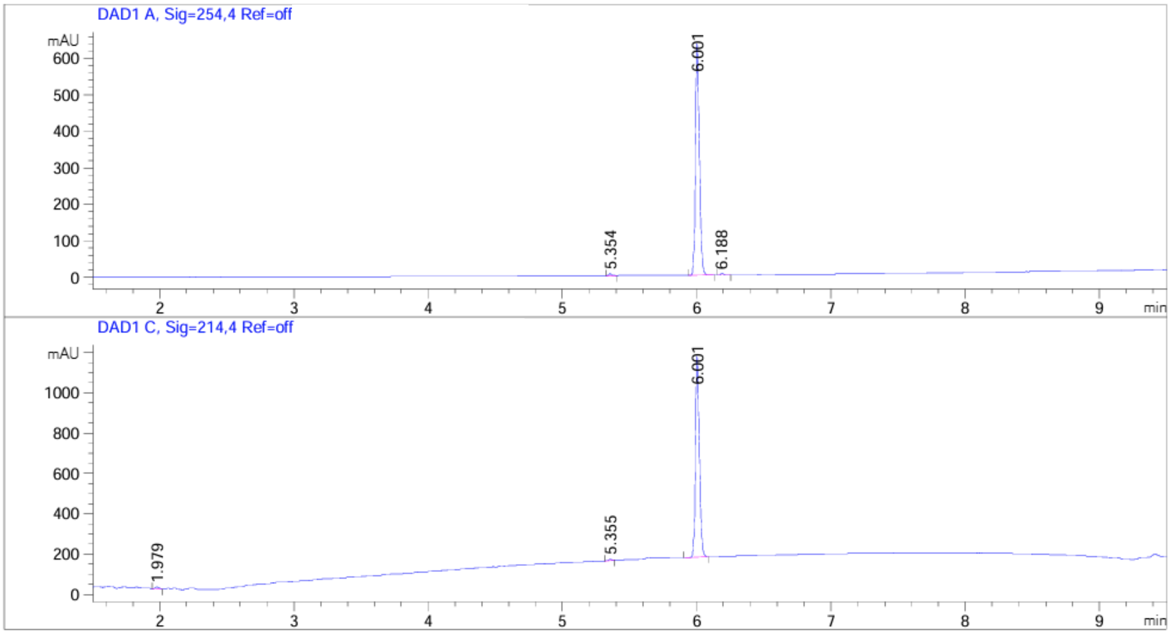


| Peak # | RetTime [min] | Type | Width [min] | Area [mAU*s] | Height [mAU] | Area % |
|---|---|---|---|---|---|---|
| 1 | 5.354 | BB | 0.0206 | 6.88677 | 5.15973 | 0.5373 |
| 2 | 6.001 | BB | 0.0303 | 1268.75769 | 634.52209 | 98.9895 |
| 3 | 6.188 | BB | 0.0259 | 6.06466 | 3.55404 | 0.4732 |

Totals : 1281.70913 643.23586

Signal 2: DAD1 C, Sig=214,4 Ref=off

| Peak # | RetTime [min] | Type | Width [min] | Area [mAU*s] | Height [mAU] | Area % |
|---|---|---|---|---|---|---|
| 1 | 1.979 | BB | 0.0360 | 18.46542 | 8.45231 | 0.9247 |
| 2 | 5.355 | BB | 0.0197 | 10.11086 | 7.74943 | 0.5063 |
| 3 | 6.001 | BB | 0.0301 | 1968.31165 | 995.82825 | 98.5690 |

Totals : 1996.88792 1012.02998

### COMPOUND 29FN HPLC

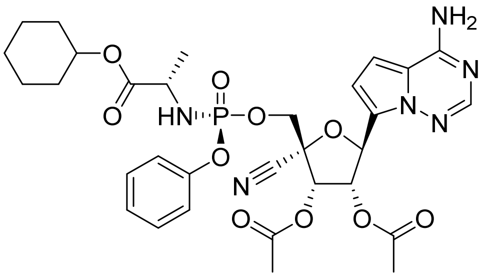


Method Info : Sample Bank Method - 2-98%B with 8.5 min gradient, A=Water + 0.1% TFA, B= Acetonitrile + 0.1% TFA; 1.5 mL/min; Column: Phenomenex Kinetex C18, 2.6u 100A, 4.6 x 100 mm; Instrument 1290II

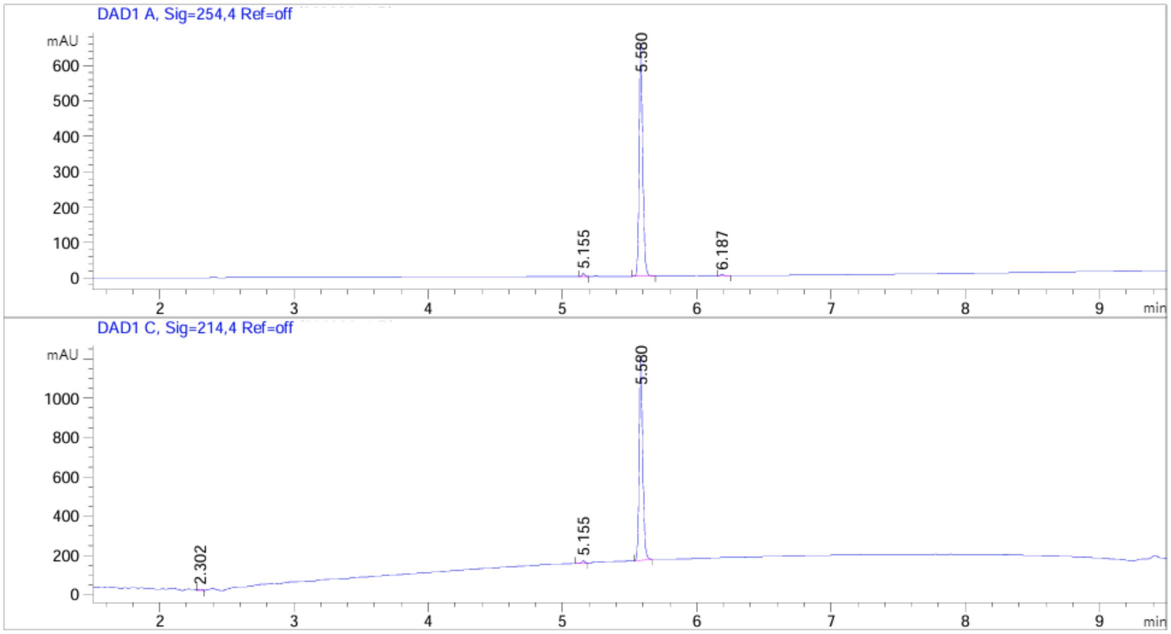


| Peak # | RetTime [min] | Type | Width [min] | Area [mAU*s] | Height [mAU] | Area % |
|---|---|---|---|---|---|---|
| 1 | 5.155 | BB | 0.0192 | 10.25042 | 8.14044 | 0.8285 |
| 2 | 5.580 | BB | 0.0282 | 1220.95825 | 656.30499 | 98.6800 |
| 3 | 6.187 | BB | 0.0258 | 6.08170 | 3.59041 | 0.4915 |

Totals : 1237.29037 668.03584

Signal 2: DAD1 C, Sig=214,4 Ref=off

| Peak # | RetTime [min] | Type | Width [min] | Area [mAU*s] | Height [mAU] | Area % |
|---|---|---|---|---|---|---|
| 1 | 2.302 | BB | 0.0270 | 8.93636 | 5.48401 | 0.4618 |
| 2 | 5.155 | BB | 0.0199 | 16.69020 | 12.65653 | 0.8625 |
| 3 | 5.580 | BB | 0.0280 | 1909.45569 | 1035.71753 | 98.6757 |

Totals : 1935.08225 1053.85807

[1] Bastien, N.; Normand, S.; Taylor, T.; Ward, D.; Peret, T. C. T.; Boivin, G.; Anderson, L. J.; Li, Y. Sequence Analysis of the N, P, M, and F Genes of Canadian Human Metapneumovirus Strains. *Virus Res.* **2003**, *93*, 51-62.
[2] Mossel, E. C.; Huang, C.; Narayanan, K.; Makino, S.; Tesh, R. B.; Peters, C. J. Exogenous ACE2 Espression Allows Refractory Cell Lines to Support Severe Acute Respiratory Syndrome Coronavirus Replication. *J. Virol.* **2005**, *79*, 3846-3850.
[3] Pitts, J.; Li, J.; Perry, J. K.; Du Pont, V.; Riola, N.; Rodriguez, L.; Lu, X.; Kurhade, C.; Xie, X.; Camus, G.; Manhas, S.; Martin, R.; Shi, P.-Y.; Cihlar, T.; Porter, D. P.; Mo, H.; Maiorova, E.; Bilello, J. P. Remdesivir and GS-441524 Retain Antiviral Activity Against Delta, Omicron, and Other Emergent SARS-CoV-2 Variants. *Antimicrob. Agents Chemother.* **2022**, *66*, e00222-22.
[4] Mason, S. W.; Lawetz, C.; Gaudette, Y.; Do, F.; Scouten, E.; Lagace, L.; Simoneau, B.; Liuzzi, M. Polyadenylation-dependent screening assay for respiratory syncytial virus RNA transcriptase activity and identification of an inhibitor. *Nucleic Acids Res.* **2004**, *32*, 4758-4767.
[5] Johnson A. A.; Tsai, Y.; Graves, S. W.; Johnson, K. A. Human Mitrochondrial DNA Polymerase Holoenzyme: Reconstitution and Characterization. *Biochemistry*, **2000**, *39*, 1702-1708.
[6] Graves, S. W.; Johnson, A. A.; Johnson, K. A. Expression, Purification, and Initial Kinetic Characterization of the Large Subunit of the Human Mitochondrial DNA Polymerase. *Biochemistry*, **1998**, *37*, 6050-6058.
[7] Warren, T. K.; Jordan, R.; Lo, M. K.; Ray, A. S.; Mackman, R. L.; Soloveva, V.; Siegel, D.; Perron, M.; Bannister, R.; Hui, H. C.; Larson, N.; Strickley, R.; Wells, J.; Stuthman, K. S.; Van Tongeren, S. A.; Garza, N. L.; Donnelly, G.; Shurtleff, A. C.; Retterer, C. J.; Gharaibeh, D.; Zamani, R.; Kenny, T.; Eaton, B. P.; Grimes, E.; Welch, L. S.; Gomba, L.; Wilhelmsen, C. L.; Nichols, D. K.; Nuss, J. E.; Nagle, E. R.; Kugelman, J. R.; Palacios, G.; Doerffler, E.; Neville, S.; Carra, E.; Clarke, M. O.; Zhang, L.; Lew, W.; Ross, B.; Wang, Q.; Chun, K.; Wolfe, L.; Babusis, D.; Park, Y.; Stray, K. M.; Trancheva, I.; Feng, J. Y.; Barauskas, O.; Xu, Y.; Wong, P.; Braun, M. R.; Flint, M.; McMullan, L. K.; Chen, S.-S.; Fearns, R.; Swaminathan, S.; Mayers, D. L.; Spiropoulou, C. F.; Lee, W. A.; Nichol, S. T.; Cihlar, T.; Bavari, S. Therapeutic Efficacy of the Small Molecule GS-5734 Against Ebola Virus in Rhesus Monkeys. *Nature* **2016**, 531, 381-385.
[8] Xu, Y.; Barauskas, O.; Kim, C.; Babusis, D.; Murakami, E.; Kornyeyev, D.; Lee, G.; Stepan, D.; Perron, M.; Bannister, R.; Schultz, R. E.; Sakowicz, R.; Porter, D.; Cihlar, T.; Feng, J. Y. Off-Target In Vitro Profiling Demonstrates that Remdesivier, is a Highly Selective Antiviral Agent. *Antimicrob. Agents Chemother.* **2021**, *65*, e02237-20.
[9] Mackman, R. L.; Hui, H. C.; Perron, M.; Murakami, E.; Palmiotti, C.; Lee, G.; Stray, K.; Zhang, L.; Goyal, B.; Chun, K.; Byun, D.; Siegel, D.; Simonovich, S.; Du Pont, V.; Pitts, J.; Babusis, D.; Vijjapurapu, A.; Lu, X.; Kim, C.; Zhao, X.; Chan, J.; Ma, B.; Lye, D.; Vandersteen, A.; Wortman, S.; Barrett, K. T.; Toteva, M.; Jordan, R.; Subramanian, R.; Bilello, J. P.; Cihlar, T. Prodrugs of a 1'-CN-4-Aza-7,9-dideazaadenosine C-Nucleoside Leading to the Discovery of Remdesivir (GS-5734) as a Potent Inhibitor of Respiratory Syncytial Virus with Efficacy in the African Green Monkey Model of RSV. *J. Med. Chem.* **2021**, *64,* 5001−5017.
[10] Vermillion, M. S.; Murakami, E.; Ma, B.; Pitts, J.; Tomkinson, A.; Rautiola, D.; Babusis, D.; Irshad, H.; Siegel, D.; Kim, C.; Xiaofeng, Z.; Niu, C.; Yang, J.; Gigliotti, A.; Kadrichu, N.; Bilello, J. P.; Ellis, S.; Bannister, R.; Subramanian, R.; Smith, B.; Mackman, R. L.; Lee, W. A.; Keuhl, P. J.; Hartke, J.; Cihlar, T.; Porter, D. P. Inhaled Remdesivir Reduces Viral Burden in a Nonhuman Primate Model of SARS-CoV-2 Infection. *Sci. Transl. Med.* **2021**, *14*, eabl8282.
